# Temporal and Functional Relationship between Synaptonemal Complex Morphogenesis and Recombination during Meiosis

**DOI:** 10.1101/2024.01.11.575218

**Authors:** Jasvinder S. Ahuja, Rima Sandhu, Lingzhi Huang, Franz Klein, G. Valentin Börner

## Abstract

During prophase of meiosis I, programmed double strand breaks (DSBs) are processed into crossovers, a critical requirement for segregation of homologous chromosomes (homologs) and genome haploidization in sexually reproducing organisms. Crossovers form via homologous recombination in close temporospatial association with morphogenesis of the synaptonemal complex (SC), a proteinaceous structure that connects paired homologs along their length during the pachytene stage. Synapsis and recombination are a paradigm for the interplay between higher order chromosome structure and DNA metabolism, yet their temporal and functional relationship remains poorly understood. Probing linkage between these processes in budding yeast, we show that SC assembly is associated with a distinct threshold number of unstable D-loops. The transition from bona fide paranemic D-loops to plectonemic DSB single end invasions (SEIs) is completed during midpachynema, when the SC is fully assembled. Double Holliday junctions (dHJs) form at the time of desynapsis and are resolved into crossovers during diplonema. The SC central element component Zip1 shepherds recombination through three transitions, including DSB first end strand exchange and second end capture, as well as dHJ resolution. Zip1 mediates SEI formation independent of its polymerization whereas precocious Zip1 assembly interferes with double Holliday junction resolution. Together, our findings indicate that the synaptonemal complex controls recombination while assembled but also beyond its disassembly, possibly by establishing spatial constraints at recombination sites.

## Introduction

Meiosis is the specialized cell division that reduces the chromosome number by half, generating haploid gametes from precursor cells that are typically diploid ^1^. Meiosis I genome reduction depends on pairing of homologous chromosomes (homologs) during prophase, which facilitates crossover formation. Crossovers together with sister cohesion along chromosome arms provide transient linkage between homologs thereby mediating their bipolar attachment to opposite spindle poles, as required for homolog segregation ^2,3^.

Meiotic recombination entails the programmed formation of DSBs and their processing via homologous recombination into crossovers and non-crossovers, with the intact homolog providing a template for DSB repair. Flanking chromosome arms are exchanged during crossover formation, but remain parental in the case of non-crossovers ^4^. DSB formation involves the covalent attachment of the transesterase Spo11 to both DSB 5’ ends, followed by 5’ end resection by a machinery that includes the Mre11-Rad50-Xrs2 complex as well as Mre11-cofactor Sae2 (a.k.a. Com1) ^5–10^. A subset of DSBs progress to crossovers, with resected single stranded 3’ ends undergoing two consecutive rounds of strand exchange, first generating single end invasions (SEIs), which next capture the cognate second DSB end giving rise to double Holliday junctions (dHJs) ^11–13^. Double Holliday junctions are preferentially resolved into crossovers ^14–16^. Another subset of DSBs progress to non-crossovers which arise from localized DNA synthesis along the homolog followed by displacement of the invading strand and religation with the opposite DSB end, without formation of stable joint molecule (JM) DNA intermediates ^14,17,18^. A minority of DSBs utilize the sister chromatid as repair template, a process that is prominent during early meiosis, before homologs have completed pairing ^19–21^. Similar to crossovers, intersister exchange involves formation of intersister (IS-)SEIs as well as IS-dHJs ^22^.

Placement of crossovers along the genome is patterned in several ways: every homolog pair receives at least one, the ‘obligatory crossover’ independent of size, as required for homolog segregation ^23^. Additional crossovers are maximally spaced by a process called crossover interference which reduces crossover frequency in a given interval when an adjacent interval along the same chromosome carries a crossover ^24^. Finally, crossover homeostasis ensures that crossovers are favored over non-crossovers when DSB abundance is low and/or homolog bias is compromised ^20,25^.

In most organisms, meiotic recombination takes place in temporospatial association with extensive changes of chromosome structure. In the budding yeast *S. cerevisiae* where recombination progression and chromosome morphogenesis can be monitored in the same cell population, chromosomes enter prophase I coupled in a pairwise manner, though independent of homology ^26,27^. Upon DSB formation, coupling interactions are replaced by homologous pairing, a process that is completed as cells enter pachynema ^20,28–30^. Contemporaneous with homolog pairing, axial elements appear along the length of closely associated sister chromatids followed by assembly of the central region of the synaptonemal complex (SC) ^31^. SC assembly is complete at the pachytene stage, when axial (a.k.a. lateral) elements of homologs are connected along their lengths by the zipper-like SC central region. Rapid SC disassembly is initiated in late pachynema and completed in diplonema, followed by diffuse stage and diakinesis ^32^.

Transverse filament protein Zip1 is a key component of the yeast SC ^33^. In the context of assembled SC, Zip1 homodimers, formed via its central coiled-coil region, are arranged in a head-to-head manner, with Zip1 C-termini pointing towards the coaligned homolog axes, and N-termini interlacing at the SC central region ^34^. Zip1 is initially recruited to the pericentromeric region where it mediates homology-independent centromere coupling ^26,27^. It subsequently polymerizes along the length of homolog pairs, starting from some or all future crossover sites, preferentially associating with DNA segments that are quasi-periodically spaced along chromosome axes ^34–36^. During leptonema, the presumed E3 ligase Zip3, a member of the functionally defined ZMM group of proteins, colocalizes with Zip1 at centromeres ^37,38^. During pachynema, Zip3 foci localize along homolog pairs with numbers and patterned spacing expected for future crossover sites ^36,37,39,40^.

Homologs enter prophase I in a parental configuration and emerge from it as recombinants, yet the temporal and functional relationship between synapsis and recombination remains poorly understood. A foundational study showed that in yeast, DSBs are formed after completion of premeiotic S-phase, followed by appearance of electrodense leptotene SC structures ^32^. In the same study, DSBs were found to persist into zygonema, followed by an extensive gap, during which recombination was subsequently shown to progress to joint molecule intermediates identified as SEIs and dHJs ^11,13,32^. Crossovers appear sometimes between the end of pachynema and the onset of metaphase I ^32^. Evolutionary conservation of recombination proteins and of chromosome architecture suggests that organisms other than yeast progress through the same recombination steps during SC morphogenesis ^41^. Yet, recombination intermediates presently are not detectable at the DNA level in organisms other than yeast, and have instead been inferred from cytological markers, with unclear stage specificity ^32,42–50^. These limitations have left unanswered questions about coordination between SC morphogenesis and recombination progression.

In addition to their temporal relationship, the functional interplay between meiotic recombination and chromosome morphogenesis also is poorly understood. Both processes appear highly interdependent in many organisms ^15,51–55^. Homolog pairing depends on DSB formation and (some) interhomolog strand exchange ^30,56–58^. SC assembly additionally requires proteins with functions in processing DSBs along the patterned crossover pathway ^15,59,60^. Conversely, mutants defective at recombination steps downstream from dHJ formation are largely normal for pairing and synapsis ^61,62^.

To gain a mechanistic understanding how the synaptonemal complex controls recombination, we have monitored chromosomal Zip1 recruitment and recombination progression in the same yeast population. Probing linkage between these processes by exposing meiotic cultures to several conditions, we show that Zip1 polymerizes along chromosome arms when DSB strand exchange intermediates reach a fixed threshold number. We also reveal an unexpected temporal relationship between synapsis and the two DSB strand exchange steps: Stable SEI intermediates are formed during midpachynema, whereas dHJs are formed after pachytene exit, followed by their resolution into crossovers during diplonema. Finally, we show that Zip1 mediates three transitions in recombination, shepherding DSBs through both first end and second end strand exchange, as well as through Holliday junction resolution.

## Results

### Temporal relationship between synapsis and recombination during wild-type meiosis

#### Incubation conditions modulate synapsis progression

Functional linkage between SC morphogenesis and recombination can be probed by exposing wild-type cells to different incubation conditions potentially resulting in temporal uncoupling of coincidental associations. Here, we induced meiosis in the synchronously sporulating SK1 strain at 30°C, the temperature most widely used in meiosis studies, as well as 23°C, and 33°C ^11,15^. Culture-specific variabilities were minimized by splitting a single G_1_-arrested premeiotic culture at the time of transfer to meiosis medium (t = 0 h) followed by incubation of subcultures at 23°C and 30°C. For 33°C, meiotic cultures were incubated for the first three hours at 26°C, followed by a shift to 33°C (for rationale see Methods in Ahuja et al. ^29^). Aliquots taken at the indicated time points were analyzed for meiotic recombination progression, Zip1 localization, nuclear divisions, crossover frequencies and interference as well as spore viability.

All three conditions are compatible with normal meiosis, as indicated by spore viabilities >95% and nuclear divisions in >90% of cells. Meiotic divisions occur with different timing, with 50% of cells having completed meiosis I at ∼6.9 h, 8.6 h and 9.3 h at 30°C, 23°C and 33°C, respectively (Fig. S1C; Table S1). Synchrony also varies somewhat, with the time required for a culture to go from 0% to 80% of bi- or tetranucleate cells requiring between four (30°C) and six hours (33°C) (Fig. S1C). In addition to temperature, ionic strength also affects the timing and synchrony of meiotic divisions. Accordingly, protoplasts undergoing in medium supplemented with potassium chloride undergo meiotic divisions with a ∼ one hour delay and a time interval of 5.5 h for the transition from 0% to 80% at 30°C (Table S1) ^32^.

Chromosomal localization of SC protein Zip1 was determined by immunodecorating surface-spread nuclei with anti-Zip1 antibody followed by immunofluorescence deconvolution microscopy (Fig. 1A-iii). In addition to duplicate WT cultures for each condition, strains carrying mutant prophase I-exit factor *NDT80* were analyzed in parallel (below) ^63^. For each culture, images of DAPI-stained, unselected nuclei were acquired with identical settings. Over 6,900 images of individual nuclei were pooled and sorted solely based on morphological similarities without knowledge of genotype, time point, or incubation temperatures [TC72; Table S2]. Scoring criteria derived from subcultures at 23°C and 30°C were later applied to the WT culture incubated at 33°C [TC55] ^29^.

**Figure 1.**
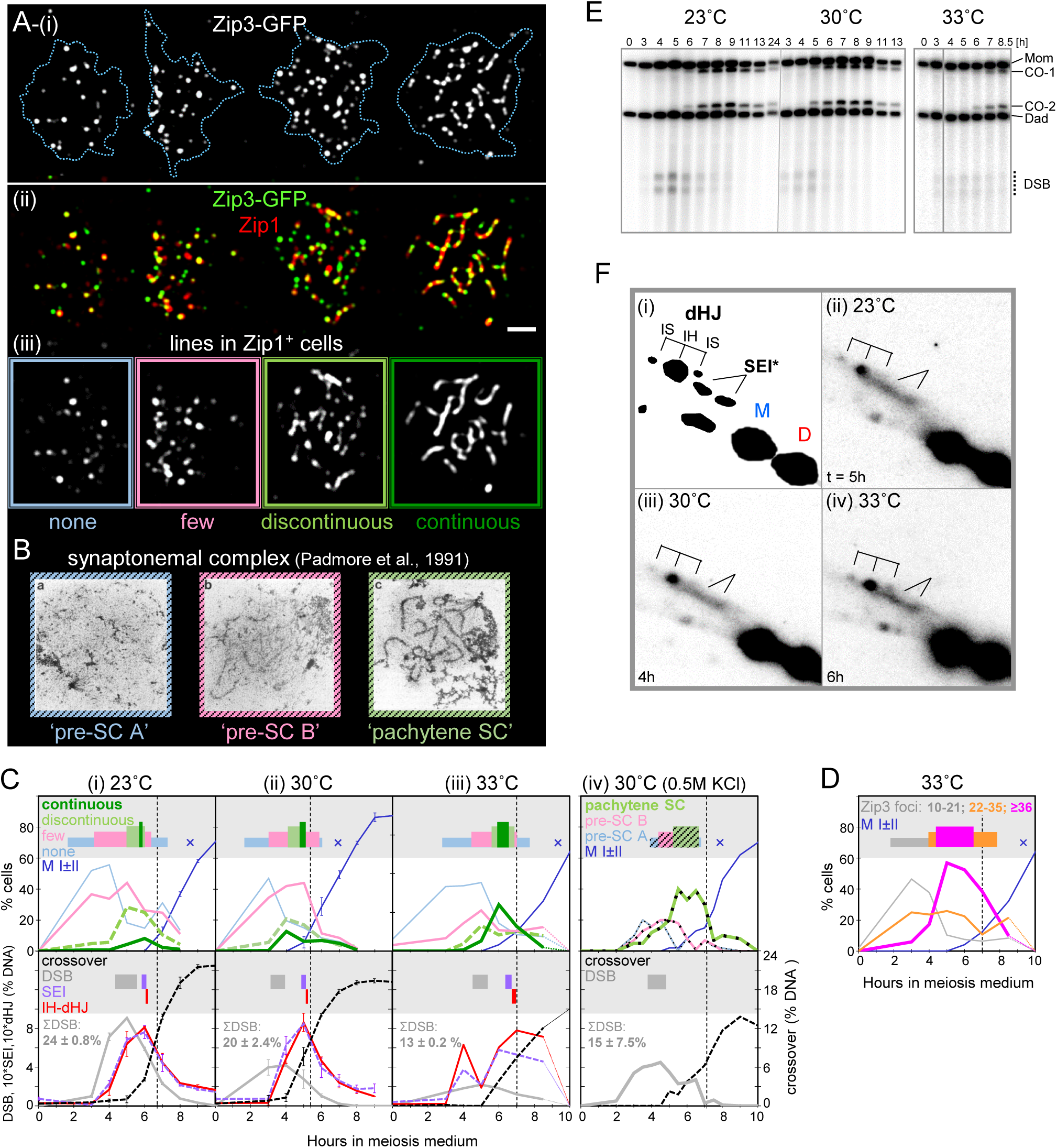
Incubation Conditions Modulate Synapsis and Recombination Kinetics. **(A)** Zip3-GFP and Zip1 localization in representative nuclei (TC55; 33°C). (i) Zip3-GFP. Bar 1 μm. (ii) Merge of Zip3-GFP (green) and Zip1 (red). (iii) Morphology of Zip1 lines: no Zip1 lines (“none”, leptotene or diplotene; blue), “few” lines (zygotene or diplotene; pink), “discontinuous” lines with underlying linear structure (early or late pachytene, light green), and “continuous” lines along most of the chromosome complement (mid-pachytene, dark green). Color coding from image frames is also used in Figs. 1B,C and Fig. 2. **(B)** Electron microscopy (EM) images of synaptonemal complex precursors “pre-SC A” (blue, leptotene) and “pre-SC B” (pink, zygotene), as well as “pachytene SC” (light green) (images from Padmore et al. (1991) ^32^. **(C)** Parallel analyses of SC morphogenesis (top panels) and recombination progression (bottom panels) in cultures incubated at (i) 23°C, (ii) 30°C, (iii) 33°C, and (iv) 30°C with 0.5 M KCl added to meiosis medium ^32^. Stages of Zip1/synapsis (top) and recombination (bottom), with subpanels in white and grey showing steady state levels as well as entry and exit times, respectively (see Methods). Dashed vertical lines indicate time of 50% crossover formation. For Zip1 analysis in (i) and (ii), images from two parallel cultures with nearly identical kinetics and efficiencies were pooled (also see Fig. S1 and Table S1). For recombination analyses, averages of intermediates and products from the same two cultures were used (Fig. S1). Parallel cultures are derived from two premeiotic cultures split at t = 0 h and incubated at 23°C or 30°C (TC72), respectively. At 33°C, a strain also carrying *zip3-GFP*/*ZIP3, rec8-HA/REC8* and *pdr5Δ* was analyzed (TC55; see Methods). For “TC11” (iv), only DSBs and COs were analyzed ^32^. Total DSB accumulation (ΣDSB) was determined in *sae2Δ* cultures incubated at 23°C (TC39, n = 2; TC16, n = 4), at 33°C (TC40, n = 2) or at 30°C (n = 3, data from Martini et al. (2006) ^25^), and in a *rad50S* background for the *HIS4LEU2* variant analyzed in TC11 ^32^. **(D)** Quantitation of Zip3-GFP classes at 33°C in the culture shown in 1C-iii (also see 1A-i). Dashed vertical line indicates 50% crossover formation. **(E)** One-dimensional gel Southern blot analysis of DSBs and COs at recombination hotspot *HIS4LEU2* at three temperatures. **(F)** (i) Schematic 2D gel indicating migration positions of parental DNA fragments (M,D), IH-SEIs, double Holliday junctions between homologs (IH-dHJ), and between sister chromatids (IS-dHJ; top left), (ii – iv) representative 2D gel Southern blot excerpts at the indicated temperatures one hour before JM peak levels (for complete blots see Figure S3A).

Despite a wide range of staining patterns, nuclei fell into five discernible classes (see gallery view in supplemental PDF P1): At very early and very late time points, mononucleate cells lack detectable Zip1 staining (‘Zip1^-^‘; not shown). Zip1-positive nuclei (‘Zip1^+^’) are characterized by the frequency and continuity of Zip1 lines: Nuclei may contain 10-25 Zip1 foci but no linear extensions (‘none’; Fig. 1A-iii), a small number of linear extensions together with a majority of foci (‘few’ lines), partial Zip1 lines along an underlying linear pattern (‘discontinuous’ lines), or Zip1 lines of mostly uniform intensities (‘continuous’ lines) ^15,29^. The aforementioned classes could also contain intensely staining Zip1 ‘polycomplex’ (PC) aggregates in chromatin-free regions, which were scored separately from chromosomal Zip1 localization (below; PDF S1). Where available, Zip3 foci were scored without knowledge of Zip1 localization status (Fig. 1A-ii).

Frequencies and kinetics of the five Zip1 classes are remarkably similar in WT cultures incubated under the same conditions (Fig. S1B). At early times, nuclei with Zip1 in foci (blue) appear at high levels (Fig. 1C; upper panels). Next, nuclei with a small number of short linear Zip1 structures reach peak levels (pink; Fig. 1C; Fig. S1B). Thereafter, nuclei with discontinuous Zip1 lines appear, followed by continuous Zip1 lines, consistent with a precursor-product relationship between these four morphologies. Zip1^+^ nuclei containing few or no Zip1 lines are also frequent at later time points. For example, at t = 7 h (23°C) more than half of Zip1^+^ cells contain either few or no lines (Fig. 1C). These cells are mostly at various stages of SC disassembly rather than representing cells that have initiated meiosis late, as cells with Zip1 lines are essentially absent at later time points (Fig. S1B). Thus, Zip1 disassembly proceeds via stages that are morphologically indistinguishable from those of Zip1 assembly. Reproducible detection only at 23°C of a distinct late peak of the ‘no lines’ class likely is related to a better separation of certain meiotic stages under these conditions. Importantly, midpachytene cells characterized by continuous Zip1 lines constitute an unambiguous class. For simplicity, nuclei with no, few, or discontinuous Zip1 lines obtained from earlier time points henceforth are referred to as leptotene, zygotene, and early pachytene, respectively. At later time points, Zip1^+^ nuclei with discontinuous lines are referred to as late pachytene, whereas those with few or no Zip1 lines are considered as diplotene. Notably, late stages of prophase I including diplonema and diakinesis are morphologically poorly defined in yeast. Given their similarities for SC and recombination progression (below; Fig. S1A-C), data from parallel cultures incubated at 23°C or 30°C, respectively, were pooled, further minimizing undue effects of lower nucleus counts in evanescent stages (Fig. 1C-i,ii; Table S2).

#### Kinetics of synapsis

Zip1 classes appear in the same order at the three temperatures, although at different levels. Importantly, at higher compared to lower temperatures, midpachytene nuclei are more frequent, with a corresponding reduction of zygotene and early/late pachytene nuclei (Fig. 1C, upper panels), as previously observed anecdotally ^15^. We conclude that higher temperatures shorten earlier, but extend later stages of prophase I, excluding a simple linear relationship between incubation conditions and meiotic progression.

Zip1 classes were subjected to cumulative analysis to determine the temporal order of SC morphogenesis relative to recombination progression as well as meiotic divisions. In this analysis, the lifespan of each non-cumulative stage is determined by dividing the area under the steady-state curve by the number of active cells ^20,32^. To obtain a cumulative entry curve, steady-state values from earlier time points interpolated at one-lifespan increments are added to a given steady-state measurement. The exit curve is provided by the entry curve shifted by one life span. By convention, the time when the cumulative curve reaches 50% of its maximum value is used as average entry into a given stage, whereas average exit corresponds to the entry time plus one lifespan (for details see Supplemental Excel S1, Info page).

Fifty percent entry and exit times are indicated by colored rectangles in the grey area above steady-state curves (Fig. 1C, i-iii; for color coding see image frames in Fig. 1A-iii), and cumulative curves as well as calculations are shown in Fig. S2 and Supplemental Excel E1, respectively. Cumulative analysis was first carried out for midpachynema which exhibits an unambiguous morphology, with entry and exit times indicating completion of SC assembly and onset of SC disassembly, respectively (Fig. 1C-i). For all other stages that may either be derived from nuclei undergoing SC assembly or disassembly, modified cumulative analysis was carried out by determining combined entry and exit times for that stage and any stage exhibiting more linear Zip1 structures (see Methods for details).

Three conclusions emerge from this analysis: First, under the three conditions, leptotene lifespans are within a similar range (1.4 to 1.7 h; Table S1), whereas zygotene and early pachytene are shortened at higher compared to the lower temperature (from 1.8 h to 1.1 h and from 0.7 h to 0.3 h, respectively). Second, the duration of midpachynema is progressively increased at higher temperatures, from 11 min (23°C) to 20 min (30°C), and to 37 min (33°C). Accordingly, midpachynema occupies 3% (23°C), 8% (30°C), and 13% (33°C) of Zip1’s time of association with chromatin (Table S1). Third, the diplotene stage [i.e., Zip1 staining with few (pink) or no lines (blue)] is also shortened at higher temperatures, from 1.3 h at 23°C to 0.9 h at 30°C and 33°C (Table S1). We infer that at higher temperatures, cells require less time for SC assembly and disassembly, yet they spend more time with fully assembled, stable SC.

Published EM data were subjected to the same modified cumulative analysis to compare chromosomal Zip1 recruitment with EM-defined SC classes (TC11 in Padmore et al., 1991 ^32^). Earlier, three nucleus classes with electrodense structures were scored ^32^, including two pre-pachytene stages, i.e. (i) nuclei with linear axes, but few tripartite SC structures (‘pre-SC A’; Fig. 1B, striped blue frame), (ii) tripartite SC central element of varying length along no more than 20% of the average yeast chromosome (‘pre-SC B’, pink frame) as well as (iii) ‘pachytene SC’ exhibiting tripartite SC structures along most of the chromosome complement (striped green frame). Correspondences in chromosome morphologies and kinetics suggest that EM-defined ‘pre-SC A’ and ‘pre-SC B’ classes are equivalent to the Zip1 leptotene and zygotene classes defined here, respectively, whereas ‘pachytene SC’ comprises early, mid and late pachynema [compare Fig. 1A-iii (green frames) with Fig. 1B (striped green)]. While standard and modified cumulative analyses provide similar entry and exit times (see Table S1), total lifespans of EM-detected morphologies are considerably shorter (∼2.8 h) than those obtained for Zip1^+^ classes (≥ 4.4 h), due to lower abundance in EM analysis of leptotene and absence of diplotene cells. Such differences may arise because sparse Zip1 loading detected by immunofluorescence does not generate electrodense structures, but differences in incubation conditions could also play a role.

#### Kinetics of recombination

Recombination was monitored at the well-characterized *HIS4LEU2* DSB hotspot by physical analyses of intermediates and products ^64^. At this locus, a DSB is induced essentially in every cell in a ∼300 bp zone ^65,66^. XhoI restriction site polymorphisms flanking the DSB site allow detection of parental DNA fragments (Mom, Dad) as well as DSBs and crossovers by one-dimensional (1D) gel Southern blot analysis (Fig. 1E). Single end invasions (SEIs) and double Holliday junctions (dHJs) are detected in the same DNA samples by two-dimensional (2D) gel analysis (Fig. 1F; e.g. ^64^).

DSB peak levels are highest at 23°C (∼9% of hybridization signal), with a progressive decrease at higher temperatures to 4% (30°C) and 2% (33°C) (Fig. 1C, grey curves in lower panels; Fig. S1A). Effects on DSB steady state levels are due only in part to decreased recombination initiation at higher temperatures: In the processing-defective *sae2Δ* background, unresected DSBs accumulate to 24 ± 0.8% (23°C), 20 ± 2.4% (30°C) and 13 ± 0.2 % (33°C; Table S1; ^25^). Crossovers are also decreased at higher temperatures, though to a lesser extent (Fig. 1E; Fig. S1A). Notably, despite decreased DSB and CO levels, both SEIs (purple) and dHJs (red) reach similar peak levels at 33°C and lower temperatures. Thus, at higher temperatures, JM lifespans are increased and DSB lifespans are decreased.

Cumulative curves of recombination intermediates were generated from averages of steady-state measurements from parallel cultures, taking into account temperature effects on recombination initiation (Fig. S1A). This analysis indicates a longer DSB lifespan at 23°C compared to 30°C and 33°C (grey rectangles in bottom panels Fig. 1C). At 23° and 30°C, SEIs (purple) and dHJs (red) exhibit similar lifespans, but they are increased at 33°C (Fig. 1C; Table S1). Consistent with earlier results ^11^, cumulative analysis places SEI exit immediately prior to entry into dHJs under all three conditions (Fig. 2A, i-iii). A precursor-product relationship between SEIs and dHJs is further supported by a higher proportion of IH-SEIs among total IH joint molecules at earlier versus later time points (see below, Fig. 5). Our current as well as earlier analyses further reveal distinct gaps between DSBs and SEIs and between dHJs and crossovers (Fig. 2A-i to iii; see Fig. 2F in Börner et al. ^15^). Such gaps may arise when recombination intermediates become undetectable by Southern blot analysis, e.g. due to DNA synthesis at DSB ends, and/or branch migration of (some) dHJs prior to their resolution ^18^. Effects of incubation conditions on recombination kinetics are further emphasized by doubling of the time between DSB exit and CO formation, from 0.9 h to 2.1 h, when meiosis occurs at 30°C at lower versus higher ionic strength (compare Fig. 2A-ii versus 2A-iv; Table S1).

**Figure 2.**
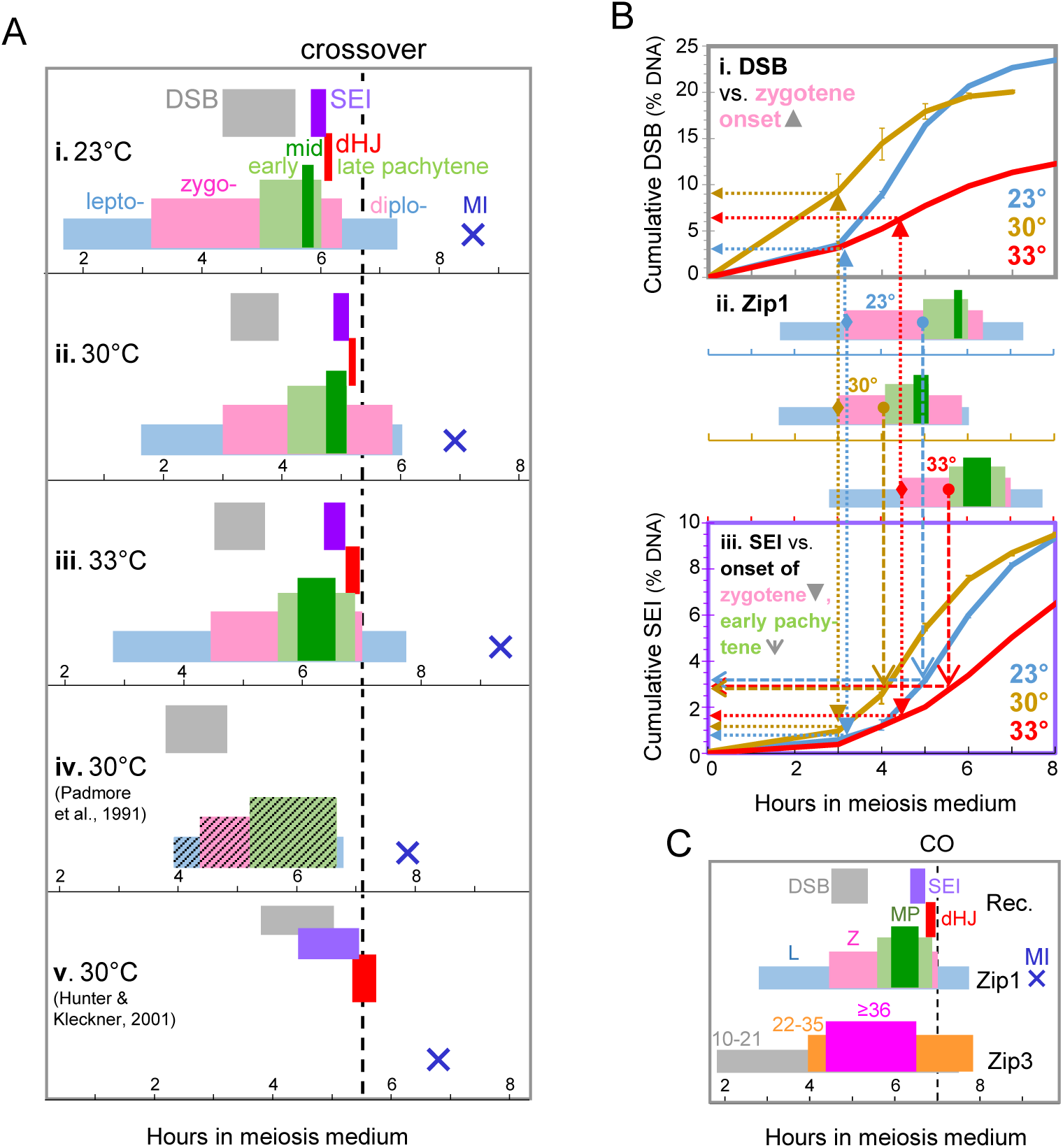
Temporal Relationship between Synapsis and Recombination during Wild-Type Meiosis. **2A** Parallel analyses of recombination and synapsis in the same cultures. Rectangles indicate the time when 50% of cells have entered into or exited from the indicated stages of recombination (DSBs, SEIs, IH-dHJs at *HIS4LEU2*) or synapsis (leptotene, zygotene, early/mid/late pachytene as well as diplotene). “Diplotene” comprises all Zip1 positive post-pachytene nuclei (pink or blue). The dashed vertical line indicates the time of 50% crossover formation which serves as shared reference time for all conditions. Blue Xs mark the time when 50% of cells have completed the meiosis I cell division. Lifespans (i) to (iv) are taken from Fig. 1, (v) shows recombination kinetics for a variant *HIS4LEU2* hotspot previously analyzed (data from Hunter and Kleckner (2001) ^11^). For non-cumulative data see Fig. S3B; for details see Table S1. **2B** Association between transitions in SC assembly and cumulative levels of recombination intermediates. (i) Cumulative DSB levels. (ii) Synapsis stages from Fig. 2A,i-iii. (iii) Cumulative SEI levels. Filled vertical arrows pointing up and down connect 50% zygotene entry times (filled diamonds) with cumulative DSB (i) and SEI curves (iii), respectively. Open arrows pointing down connect 50% early pachytene entry times (circles) with cumulative SEI curves (iii). Horizontal arrows indicate cumulative levels of (i) DSBs and (iii) SEIs at the corresponding cytological transition stages. Error bars indicate range. **2C** Temporal relationship between recombination, synapsis and Zip3 recruitment (33°C). Horizontal rectangles represent stages of recombination, Zip1 as well as Zip3 recruitment. Data are from Fig. 1C,D.

### Synapsis initiation and extension are correlated with threshold numbers of DSB strand exchanges

Several issues need to be considered regarding the kinetics of recombination and synapsis: First, whereas nucleus-wide events are monitored in our synapsis analysis, potentially exhibiting some intranuclear asynchrony, recombination analysis is limited to *HIS4LEU2*, presently the only DSB hotspot that allows robust detection of all recombination intermediates and products. Yet, *HIS4LEU2* likely is representative for DSB timing, as nucleus-wide Rad51 foci and the DSBs at an alternative hotspot exhibit comparable kinetics when analyzed in the same cells (see Figs. 2E and 5H, respectively, in Joshi et al. ^21^). Second, despite some uncertainties for SEIs, quantitative assumptions for IH-dHJs as CO-specific intermediates are well supported ^11^. Third, due to preferential recovery of DNA from cells that have not yet completed spore formation, levels of recombination intermediates and crossover products may be over- and underestimated, respectively, at late time points ^32^. Thus, JMs and COs may form somewhat earlier and later, respectively, than suggested by our analysis. Finally, cumulative analysis provides average 50% entry- and exit times, but would not be able to capture e.g., multiple waves of recombination intermediates or products.

Our analysis of SC morphogenesis and recombination in the same cell population confirms some earlier conclusions but also provides new insights about their relative timing. Entry into crossovers occurs during diplonema, as clearly discernible in cultures sampled at high frequency during later time points (23°C and 30°C; crossovers are indicated by the vertical dashed line; Fig. 2A-i,ii). Accordingly, pachytene exit is separated from CO entry by 41 min (23°C) and 18 min (30°C). Consistent with these findings, crossovers appear 32 min after pachytene exit in a meiotic culture surveyed at 30°C by EM ^32^ (Fig. 2A-iv).

DSB entry, by contrast, is not associated with a single stage of SC morphogenesis, coinciding with zygotene entry at 30° and 33°C, while occurring later at 23°C (Fig. 2A, i-iii). DSB exit is also not associated with a distinct synapsis stage, occurring during early pachytene entry at 30° and 33°C, but towards the end of early pachytene at 23°C (Fig. 2A,i-iii). This lack of temporal correlation is unexpected as genetic analysis indicates dependence of synapsis on recombination in yeast (Introduction) ^67^. One explanation for these findings is that transitions in SC assembly are correlated with absolute threshold numbers of recombination intermediate(s), rather than with a particular stage of recombination. To examine this possibility, cumulative numbers of DSBs and SEIs at key transitions in SC assembly were determined (Fig. 2B). No correlation was apparent between DSBs and SC assembly, as cumulative DSB levels vary widely at the time of zygotene onset (see upward arrows between zygotene onset and cumulative DSB curves in Fig. 2B-i,ii). Conversely, both onset of zygotene and of early pachytene are correlated with distinct absolute SEI numbers: At the three temperatures, cumulative SEI numbers cluster around 1.2% and 3.1% of DNA at zygotene and early pachytene entry, respectively (see dotted and dashed vertical arrows, respectively, in Fig. 2B-ii,iii). Thus, transitions in Zip1 polymerization are correlated with a distinct threshold number of SEIs, with the first threshold coinciding with the leptotene-zygotene transition and the next higher SEI number coinciding with the zygotene-early pachytene transition. Most likely, an unstable SEI precursor (pre-SEI), possibly involving a paranemic interaction between resected DSB and template DNA, is associated with one or both transitions (Discussion). Under conditions where DSBs form at high levels (23°C), pre-SEI numbers sufficient for triggering SC assembly are achieved earlier, yet DSBs may continue to form thereafter, resulting in a shift of average DSB entry times to a later cytological stage (see Fig. 2A-i).

### Double Holliday junctions are formed and resolved during diplonema

Under the three conditions examined here, entry into SEIs occurs during midpachynema suggesting that crossover-specific, stable DSB first end strand exchange is completed only when homologs are fully and stably synapsed (Fig. 2A). Strikingly, double Holliday junctions are formed and resolved during diplonema, after pachytene exit. Accordingly, entry into dHJs coincides with the exit from mid- or late pachynema at both 23°C and 30°C (Fig. 2A-i,ii). At 33°C, dHJ entry and exit also occur after the midpachytene stage, despite undersampling of later time points (Fig. 2A-iii), and at t = 8.5 h, Zip1 lines have essentially disappeared whereas dHJs remain close to peak levels (Figs. 1C-iii). It is noteworthy that all homolog pairs are synapsed during midpachynema, eliminating uncertainties about the synapsis status of the chromosome carrying the recombination hotspot. We conclude that DSB first-end strand exchange occurs during mid-pachynema, when homolog pairs are most intimately associated, whereas DSB second-end strand exchange and formation as well as resolution of dHJs occur during diplonema.

A previous study had inferred that SEIs and dHJs form earlier relative to SC morphogenesis, a conclusion obtained from overlaying and proportionally shortening data from cultures progressing through meiosis with different kinetics ^11^. By contrast, the earlier SEI and dHJ data were obtained under very similar salt conditions as those used here, and cultures completed 50% meiotic divisions within 7 min (Table S1). When the earlier SEI and dHJ data are superimposed with our synapsis analysis at 30°C, SEIs coincide with pachynema and dHJs as well as crossovers appear during diplonema, consistent with our conclusion (compare Figs 2A-ii with 2A-v). [Note that the *HIS4LEU2* hotspot variant analyzed earlier^11^ exhibits longer SEI and dHJ lifespans compared to the version used here (see Fig. S2B in Joshi et al. ^21^ and Oh et al. ^64^ for parallel analyses), likely contributing to an earlier SEI onset and overlap of IH-dHJs with crossovers (Fig. 2A-v; compare Fig. 1C-ii with Fig. S2B; ^11,64^]. Taken together, these data indicate that DSB first-end strand exchange is stabilized during midpachynema, whereas DSB second-end capture and double Holliday junction formation are completed after the SC has disassembled, followed by bulk crossover formation during diplonema.

### Zip3 is recruited to future crossover sites at DSB formation and persists into diplonema

Zip3 associates with centromeres when meiosis is initiated, but later localizes along paired homologs with numbers and distribution expected for crossovers ^36,37,39^. Zip3 further is critical for the formation of SEIs, dHJs and crossovers ^15^. To determine the timing of crossover site designation relative to recombination progression and synapsis, Zip3-GFP focus numbers were monitored in the same culture at 33°C also assayed for recombination and Zip1 recruitment (above). Zip3-GFP nucleus classes were defined as containing 10-20 foci, suggesting association with the 16 pairs of yeast centromeres, 21-35 foci, indicating an intermediate Zip3 focus number, or ≥ 36 foci, indicating Zip3 association with some or all interference-distributed crossover sites (Fig. 1A-i,ii; Fig. S3C) ^29^. 36 foci were chosen as cutoff for the class with the highest number of Zip3 foci since essentially all midpachytene nuclei contain ≥ 36 foci (below; Fig. S3D).

Nuclei containing 10-20 Zip3 foci appear at t = 3 h, followed by nuclei with 21-35 foci which remain detectable at substantial frequencies throughout the experiment. Nuclei with ≥ 36 foci are prominent from 5 h to 7 h (Fig. 1D). Modified cumulative analysis was carried out by first determining the 50% entry and exit times for the unambiguous nucleus class with ≥ 36 Zip3 foci, followed by calculation of lifespans for stages with fewer Zip3 foci combined with those with more foci (see above). Accordingly, cells enter the stages with 10-20, 21-35 and ≥ 36 Zip3 foci at 1.8 h, 4.0 h and 4.4 h, respectively (Fig. 1D; 2C).

Temporal comparison of Zip1 and Zip3 classes results in the following conclusions. First, Zip3 foci at centromeres appear slightly before or together with Zip1 foci (leptotene), consistent with earlier reports ^38^ (Fig. 1A-ii, 2C). Recruitment of Zip3 prior to Zip1 is suggested by detection of 20% of nuclei with 10-20 Zip3 foci that lack any Zip1 staining at t = 3 h (Fig. S3D). Second, Zip3 focus numbers increase to ≥ 36 by t = 4.5 h, exhibiting a characteristic string-of-beads interference distribution while cells are still at the leptotene stage (Fig. 2C) ^68^. Consistent with an increase of Zip3 focus numbers to maximum levels prior to Zip1 polymerization into continuous lines, more than half of Zip1 leptotene nuclei contain intermediate or high numbers of Zip3 foci (Fig. S3D). Zip1 transitions into zygonema and pachynema occur while Zip3 foci remain at maximum numbers (Fig. 3C). Zip3 appears to persist at designated crossover sites throughout midpachynema, as essentially all midpachytene nuclei exhibit maximum Zip3 focus numbers (Fig. 1A; 2D). Following pachytene exit, Zip3 remains detectable along chromosomes in substantial numbers. Accordingly, of the 75 Zip1^+^post-pachytene cells at t = 7 h and 8.5 h (no or few Zip1 lines), 31% exhibit ≥ 36 Zip3 foci and 39% exhibit 22-35 Zip3 foci, indicating that a substantial number of Zip3 foci persist into diplonema.

**Figure 3.**
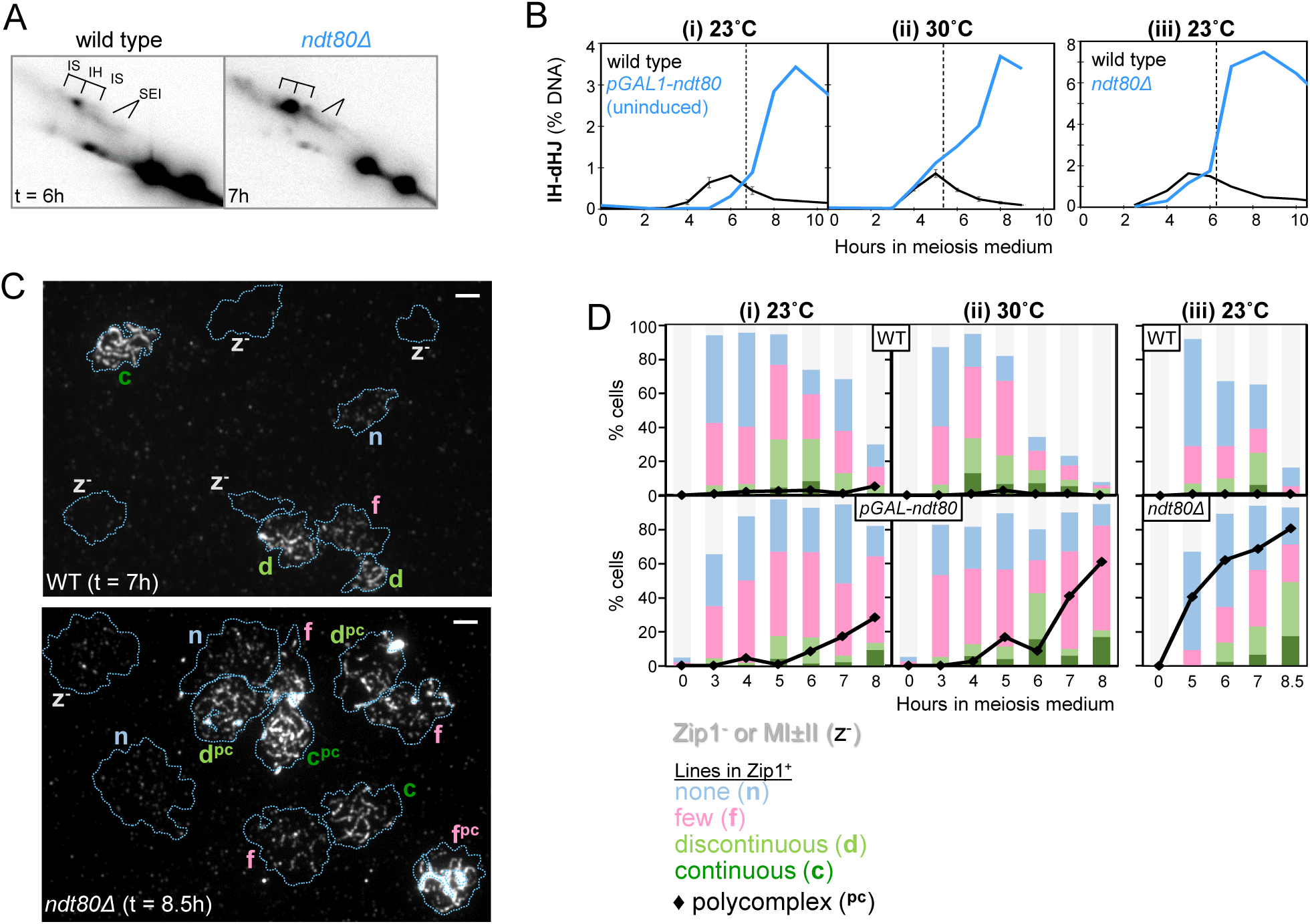
The *ndt80* Mutant Forms Double Holliday Junctions despite Defective Pachytene Entry. **3A** Excerpts from 2D gels at the time of maximum JM levels in *ndt80Δ* (8.5 h) and WT (6 h) at 23°C (TC42). Bar 2 μm. **3B** Quantitation of IH-dHJs in parallel cultures of WT and *ndt80* (uninduced *pGAL1-ndt80*) at (i) 23°C and (ii) 30°C as well as (iii) WT and *ndt80Δ* at 23°C. For complete blots see Fig. S4B. **3C** Representative image frames of maximum levels of synapsis in WT and *ndt80Δ* at 23°C. The outside edges of DAPI-stained chromatin are traced by dotted lines. Letters indicate Zip1 status, for legend see Fig. 3D. Bar 2 μm. **3D** Quantitation of Zip1 classes and polycomplexes in parallel cultures of WT and *ndt80*. Wild-type cultures in (i) and (ii) are the same as in Fig. 1. For effects of beta-estradiol induction on recombination and meiotic progression in the *pGAL1-ndt80* cultures, see Fig. S4C,D.

Surprisingly, entry into the stage with maximum Zip3 foci coincides with bulk DSB formation (Fig. 2C). Zip3 foci persist at high levels until SEI appearance but exhibit reduced numbers when dHJs appear. Zip3 foci remain at intermediate numbers when crossovers appear and they invariably colocalize with persisting Zip1 foci (not shown). Together, these findings indicate that Zip3 is displaced from centromeres at the time of DSB formation and relocalizes to presumed designated crossover sites, prior to Zip1 polymerization, with important implication for the timing when crossover interference is established (Discussion). Zip3 remains associated with future crossover sites until strand exchange and appearance of SEIs but dissociates from most chromosomal positions as the SC disassembles and SEIs progress to dHJs.

### Double Holliday junction formation is uncoupled from pachytene entry in ndt80Δ

An earlier inference that dHJs form in the context of fully assembled SC, and are resolved upon pachytene exit was derived in part from analysis of the *ndt80Δ* mutant where cells reportedly arrest with fully assembled SC and unresolved dHJs ^14,63,69^. To address this apparent discrepancy with our observations, recombination and Zip1 assembly into SC were analyzed in two *ndt80* mutants in parallel with wild-type cultures. Both an *NDT80* deletion (*ndt80Δ*) and an uninduced strain expressing *NDT80* under the control of the β-estradiol-inducible *pGAL1* promoter (*pGAL-ndt80*) were analyzed, at 23° and 30° C. Sample preparation, image capturing as well as image scoring occurred in parallel with wild type (above).

In both *ndt80Δ* and uninduced *pGAL-ndt80* compared to wild type, IH-dHJs appear with normal timing (30°C) or with a delay (23°C), and accumulate to 3- to 4-times peak levels in wild type at late time points, consistent with earlier observations (Fig. 3A,B) ^14^. Crossovers reach ∼50% of wild-type levels in *ndt80* by t = 24 h at both temperatures, with comparable reductions at earlier time points, levels similar to those observed e.g., in *zip1Δ* (Fig. S4A,C; see also Fig. 6). When *pGAL1-ndt80* is induced in aliquots of the same cultures ∼1.5 hours after the IH-dHJs peak in wild type, accumulated dHJs are rapidly resolved, crossovers rise to wild-type levels and meiotic divisions occur in >90% of cells, indicating that *pGAL-ndt80* cultures had entered meiosis with high efficiency and synchrony (Fig. S4C).

Unexpectedly, in both *ndt80Δ* and uninduced *pGAL-ndt80*, nuclei with continuous or discontinuous Zip1 lines do not exceed peak steady state levels in parallel WT cultures, even though most nuclei exhibit Zip1 staining (Fig. 3C,D). Accordingly, the majority of *ndt80* nuclei exhibit Zip1 in foci (blue) or foci with few lines (pink), morphologies typically observed in leptotene or zygotene nuclei, respectively. Moreover, polycomplexes appear in *ndt80* cultures at the time of wild-type SC assembly and reach frequencies of 40% to 80% of cells, compared to ≤ 5% in WT or in a *zip1* point mutation that arrests transiently at pachynema (Fig. 3C,D; below). Thus, in *ndt80Δ* at 8.5 h, >80% of nuclei carry a PC, and fewer than 6% of pachytene nuclei are free of a PC, consistent with earlier reports ^20,63,69,70^. Notably, frequency and timing of polycomplexes in *ndt80* are comparable to those in mutants defective for SC assembly ^15^. We conclude that dHJ formation is uncoupled from pachytene entry in *ndt80* in the SK1 background. Observations from the *ndt80* mutant are therefore not informative with respect to events that normally occur during and after pachynema. Furthermore, these findings suggest that dHJ formation does not depend on normal SC assembly or disassembly, even though SC disassembly normally precedes dHJs formation.

### Crossover interference is modulated by incubation conditions

To ensure that features of meiosis occur normally under the incubation conditions examined here, crossover frequencies and interference were examined in a strain carrying marked intervals along three chromosomes ^64^ (Fig. 4A). Meiotic progression in sub-cultures analyzed at 23°C and 30°C was indistinguishable from that in strains analyzed for recombination and synapsis (Fig. S5A). Genetic map distances also were remarkably similar under the two conditions (Fig. 4B). Only one interval on chromosome III (*CEN3*-*MAT*) exhibits a significant increase in map distance at 23°C versus 30°C. Non-Mendelian segregation/ gene conversion also occurs with similar frequencies, even though larger (i.e., >2-fold) increases tend to occur at 23°C versus 30°C (Fig. S5B). Together, these findings suggest that recombination frequencies are not dramatically different at the two temperatures.

**Figure 4.**
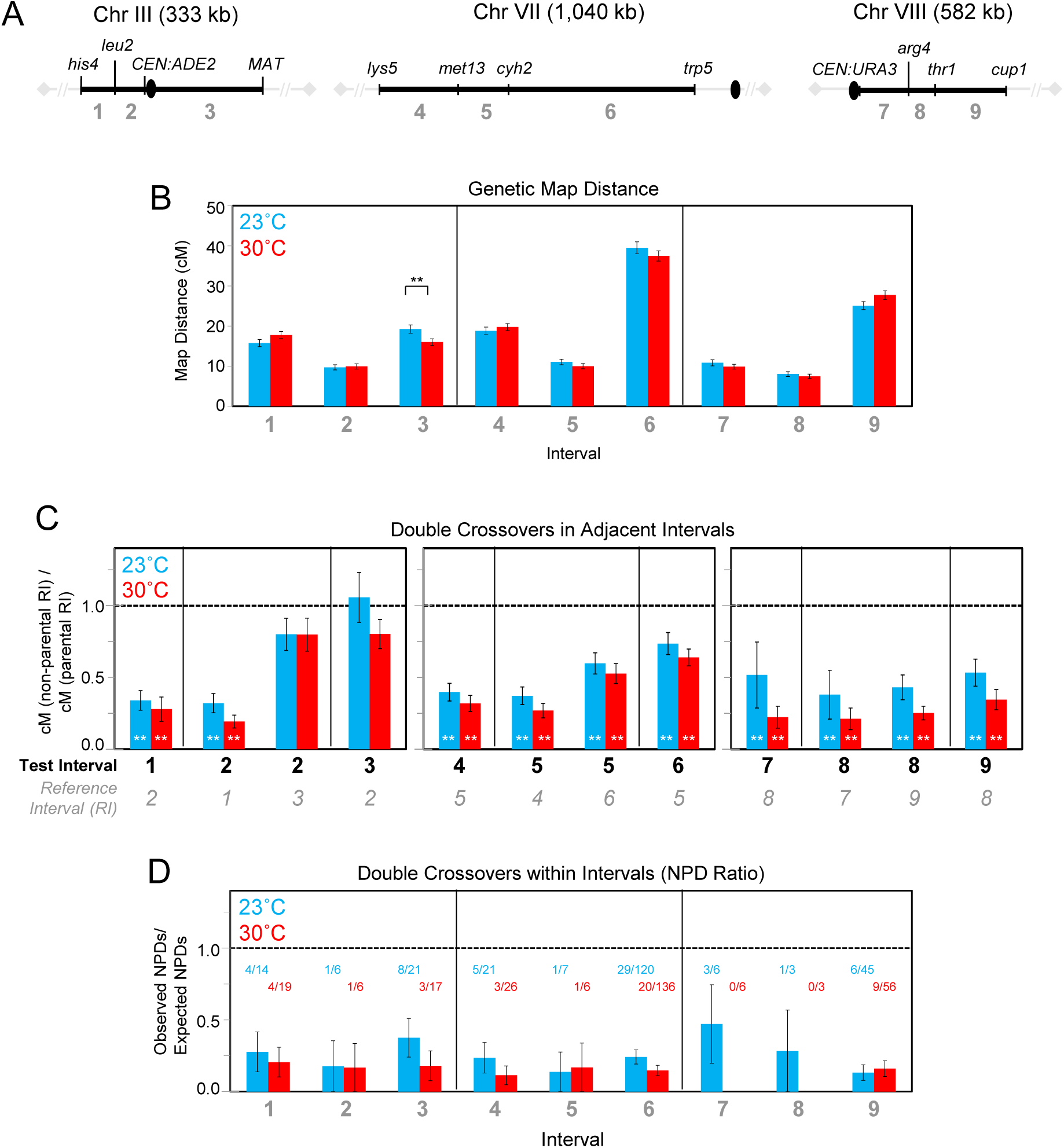
Genetic Map Distances and Interference at 23°C and 30°C. **4A** Maps of nine genetic intervals along yeast chromosomes III, VII, and VIII. Numbers 1 through 9 provide interval names used here and in Table S3. Kilo base pairs indicate physical sizes of chromosomes, black ovals indicate location of centromeres, grey diamonds indicate chromosome ends. **4B** Genetic map distances at 23°C and 30°C derived from meiotic WT cultures incubated under identical conditions as those used for analyses of recombination and Zip1 recruitment (Fig. 1). Map distances were determined using the Perkins equation, standard errors were determined using the Stahl-website. Two stars indicate map distances differing significantly at 23°C compared to 30°C. Spore viabilities were 95.5% (23°C) and 97.7% (30°C), numbers of four viable spore tetrads were n = 1,192 (23°C) and n = 1,176 (30°C). Data for 23°C and 30°C are derived in part from the same premeiotic culture split at t = 0 h, except for 722 tetrads at 30°C which came from a parallel isogenic culture. Error bars are SE. **4C** Coefficient of coincidence at 23°C and 30°C determined for a given ‘Test Interval’ (bold) flanked by a ‘Reference Interval’ (RI; italic) exhibiting either parental or non-parental marker configurations. Positive interference is indicated if the genetic distance in a given test interval is significantly lower among tetrads flanked by a recombinant reference interval compared to those flanked by a parental reference interval. A ratio of 1 (dashed line) indicates that the test interval is unaffected by the recombination status of the reference interval. Two white asterisks indicate significant interference. All adjacent interval pairs exhibit significant interference apart from interval pair 2 and 3, and interference tends to be stronger at 30°C compared to 23°C (for details see Table S3). Error bars are SE. **4D** Ratios of observed versus expected double crossovers involving all four chromatids (‘NPD Ratios’) at 23°C and 30°C in intervals 1 through 9. Values in blue and red indicate observed over expected NPD tetrad numbers. Error bars are SE.

Crossover interference was assayed using two approaches ^71,72^. The coefficient of coincidence is obtained from the ratio of ‘observed’ to ‘expected’ crossovers, with ratios below 1 indicating positive crossover interference ^71^. In this analysis, ‘expected’ and ‘observed’ frequencies are determined for a test interval in tetrad subsets that in a reference interval along the same chromosome exhibit parental or recombinant marker configurations, respectively. At both temperatures, significant crossover interference is detected in all but one interval pair (Fig. 4C). Remarkably, however, crossover interference tends to be stronger at 30°C compared to 23°C, as indicated by a lower coefficient-of-coincidence in all interval pairs, even though this difference is significant only for one interval pair (Table S3).

A trend towards weaker crossover interference at 23°C is also apparent when frequencies of four chromatid crossovers are considered, as suggested by higher ratios of observed to expected non-parental ditypes (NPD ratio) (Fig. 4D). Thus, interference is intact at the two temperatures despite apparent shifts in timing between recombination and SC morphogenesis. Stronger interference at 30°C raises the possibility that earlier bulk DSB formation and/or a longer duration of pachynema could make crossover interference more robust.

### Synapsis-defective Zip1-C1 is intact for SEI formation, but impaired for DSB second end capture

Having established the temporal relationship between Zip1 assembly into SC and recombination during wild-type meiosis, we next examined roles of Zip1 recruitment in recombination focusing on two previously identified mutants reportedly defective for chromatin association (*zip1-C1*) or SC disassembly (*zip1-L657A*) ^73,74^. Analysis was carried out at 23°C to ensure maximum resolution for recombination progression (above).

*zip1-C1* carries an internal 34 amino acid deletion in its C-terminal globular domain ^73^. Its phenotype resembles a *zip1* deletion in many respects, including absence of an SC central element despite coaligned homolog axes. Yet, *zip1-C1* exhibits PCs at high frequencies, and spore formation in SK1 is reduced to 36% compared to >90% in *zip1Δ* ^73^. Accordingly, fewer than 25% of *zip1-C1* cells complete meiotic divisions at 23°C, compared to >70% cells in the parallel *zip1Δ* culture (Fig. 5A; Fig. S6E).

**Figure 5.**
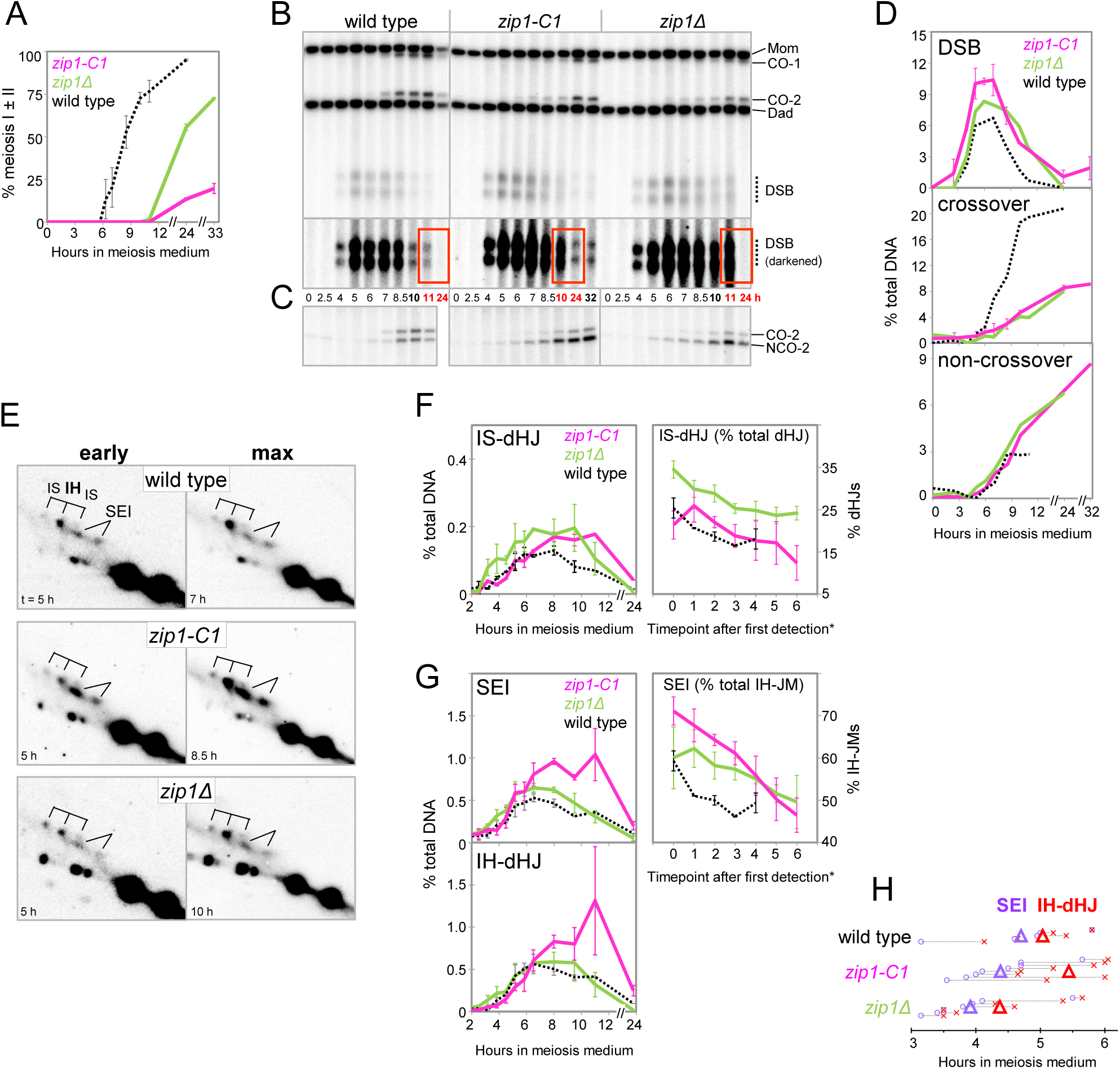
Synapsis-Defective *zip1-C1* Mediates DSB First-End Strand Exchange but is Impaired for DSB Second-End Capture (23°C) **5A** Meiotic nuclear divisions in *zip1-C1* (n = 3), *zip1Δ* (n = 2), and WT (n = 2). Time axis is interrupted twice (after 11 h and 24 h). Error bars indicate range. **5B** One dimensional gel Southern blot analysis of recombination at *HIS4LEU2* in WT, *zip1-C1* and *zip1Δ.* For recombination intermediates and products see Figure 1D. Lower panel, darker adjustment of DSB region of blot. Red boxes highlight late DSBs detected between 10 h and 24 h. **5C** Excerpt from one-dimensional gel Southern blot showing CO-1 and NCO-1 products refractory to NgoMIV in a double digest with XhoI. **5D** Quantitation of DSBs, crossovers (Fig. 5B) and non-crossovers (Fig. 5C) in WT, *zip1-C1*, and *zip1Δ*. Error bars indicate range of duplicate cultures where available (tc16-RaS). **5E** Representative excerpts from 2D gel Southern blot analysis in WT, *zip1-C1* and *zip1Δ,* at the time of first detection (early) and at peak time (max). For intermediates see Figure 1F. For additional blots see Figure S6D. (tc16 RaS). [Note that the longer SEI signal in *zip1-C1* appears somewhat earlier than the shorter one, likely due to earlier DSB formation on the Mom compared to the Dad homolog.] **5F** *Left panel:* Quantitation of IS-dHJs in *zip1-C1* (magenta), *zip1Δ (*green) and WT (dotted) in parallel cultures (n = 3). *Right panel:* Sister bias as indicated by IS-dHJs as percent of total dHJs (i.e., IS-dHJs plus IH-dHJs). Asterisk: Informative JM signals are typically detected first at 4.5 h, 3.2 h and 3.8 h, respectively, in *zip1-C1*, *zip1Δ* and WT (tc17-RiS). Starting at the time of first detection, seven time points were analyzed. **5G** *Left panels:* Quantitation of SEIs (top) and IH-dHJs (bottom) in *zip1-C1*, *zip1Δ* and WT in parallel cultures (n = 3). *Right panel:* Fraction of (IH-)SEIs among total interhomolog joint molecules (SEIs plus IH-dHJs) as an indicator of SEI appearance and persistence. Error bars are SDs. Time points analyzed are the same as in Fig. 5F (tc17-RiS). Asterisk: See Fig. 5G. **5H** Time of formation of SEIs and IH-dHJs in WT, *zip1-C1* and *zip1Δ.* Large triangles indicate average times when SEIs (purple) and IH-dHJs (red) reach 50% of maximum WT levels. The respective times in individual meiotic cultures are indicated by empty circles (SEIs) and Xs (dHJs), connected by dotted lines (WT, n = 5; *zip1-C1*, n = 8; *zip1Δ*, n = 7).

Despite cytological similarities with *zip1Δ*, *zip1-C1* exhibits remarkable features for recombination progression. While DSB maximum levels and kinetics are similar in *zip1Δ* and *zip1*-*C1* in the *sae2Δ* background (Fig. S6A,B), steady-state DSBs in a *SAE2* background are increased in *zip1-C1* (Fig. 5D; Fig. S6C) and a subset of DSBs reproducibly persist beyond 11 h, into time points when DSBs in both *zip1Δ* and wild type have disappeared (Fig. 5B). Importantly, *zip1-C1* exhibits four distinct effects on the DSB strand exchange events that give rise to SEIs and dHJs (Fig. 5E-H). First, the success of stable DSB strand exchange is increased in *zip1-C1* compared to *zip1Δ*, as indicated by substantially higher peak levels of JM intermediates along the interhomolog pathway, with increases in *zip1-C1* of 2.0 (± 0.6) and 2.4 (± 1.1)-fold over WT for SEIs and dHJs, respectively, versus 1.1 (± 0.2) and 1.3 (± 0.3)-fold increases in *zip1Δ* (n = 7; Fig. 5G).

Second, unlike *zip1Δ, zip1-C1* is competent in maintaining the fate of strand exchange intermediates, as indicated by preferential use of the homolog over the sister chromatid (homolog bias). Thus, IS-dHJs appear with WT levels and kinetics in *zip1-C1*, reaching increased levels only when IH-dHJs also accumulate (Fig. 5F,G). Moreover, the fraction of IS-dHJs among total dHJ in *zip1-C1* closely traces WT values of <25%. In *zip1Δ*, by contrast, IS-dHJs are increased especially at early time points, both in absolute terms and relative to IH-dHJs, with IS-dHJs accounting for ∼35% of total dHJs (Fig. 5E,F; Fig. S6D). Notably, frequency of intersister dHJs compared to interhomolog dHJs provides information about the DSB first end strand exchange step as both intermediates are derived from the respective first end strand exchanges.

Third, DSB first end strand exchange in *zip1-C1* occurs with WT kinetics, as indicated by the time when SEIs reach 50% of WT maximum levels at 4.4 (± 0.6) h compared to the WT at 4.7 (± 0.9) h (Fig. 5H). In *zip1Δ*, by contrast, strand exchange occurs precociously along both the interhomolog and the intersister pathways, as indicated by entry into SEIs at 3.9 (± 0.7) h as well as by early appearance of IH-dHJs and IS-dHJs (Fig. 5F, H; Fig. S6D) (n = 7).

Fourth and importantly, *zip1-C1* is impaired for DSB second-end capture, resulting in transient accumulation of IH-SEIs prior to appearance of IH-dHJs. Accordingly, in *zip1-C1* the IH-SEI signal is more prominent than IH-dHJs especially at earlier time points, in contrast to wild type or *zip1Δ* (Fig. 5E, Fig. S6D), and further evident from an increased contribution of IH-SEIs to total IH-JMs in *zip1-C1* at all times (Fig. 5G, right panel). Delayed progression from IH-SEIs to IH-dHJs further results in a pronounced gap between the time when SEIs and IH-dHJs reach half maximum WT levels: Compared to gaps of 0.46 (± 0.40) h in *zip1Δ* and 0.34 (± 0.36) h in wild type, SEIs and IH-dHJs in *zip1-C1* are separated by 1.1 (± 0.6) h (Fig. 5H).

Despite more efficient formation of IH-SEIs and IH-dHJs in *zip1-C1*, crossover and non-crossover levels as well as kinetics in *zip1-C1* closely resemble those in *zip1Δ* (Fig. 5C,D; Fig. S6C). Increases of interhomolog JMs that are not accompanied by increases in crossovers or non-crossovers suggests that *zip1-C1* improves the efficiency of IH strand exchange, but fails to support formation of recombination products, possibly because intermediates decay into other, non-detectable intermediates or products. Conversely, viability among the subset of cells that form tetrads is increased to ∼51% (*zip1-C1*) from ∼34% (*zip1Δ*) (Fig. S6F). Finally, we do not consider as likely that increased JM levels in *zip1-C1* compared to *zip1Δ* are due to a resolution defect, as this should also be associated with a delay in crossover and/or non-crossover formation.

One interpretation of these findings is that unlike *zip1Δ*, *zip1-C1* remains partially competent for mediating progression along the interhomolog pathway, as indicated by timely and efficient SEI formation with normal homolog bias. *zip1-C1* is impaired, however, for the transition from IH-SEIs to IH-dHJs. Importantly, effects of *zip1-C1* on JM fate and kinetics likely represent a normal Zip1 function, as *zip1-C1* is recessive for both meiotic progression and spore viability (Fig. S6E,F), whereas gain-of function mutations usually are dominant. We conclude that Zip1 plays a role not only at the DSB-to-SEI transition, as previously inferred and further corroborated here from analysis of *zip1Δ* ^15^, but also in the ensuing DSB second-end capture. These two Zip1 functions can be separated.

### Zip1-C1 mediates SEI formation independent of association with chromosome axes

To determine how *zip1-C1* affects meiotic recombination, we analyzed Zip1 chromatin association using immunofluorescence and chromatin immunoprecipitation (ChIP-on-chip) analysis. In addition to *zip1-C1*, an internally FLAG-tagged C1 allele was used in some experiments with *zip1-C1-FLAG* sharing key features of the untagged allele (Fig. S7A).

At 23°C, both Zip1-C1 and Zip1-C1-FLAG are expressed as proteins of expected lengths and accumulate to wild-type levels (Figure S7B). Zip1-C1 recruitment to surface-spread nuclei initially occurs with apparently normal patterning and timing (Fig. 6A,B). Accordingly, Zip1^+^ nuclei with no lines (see Fig. 1A) occur at ∼WT frequencies at early times (t = 4 h), although most of them also contain a polycomplex. Nuclei with some (pink) or extensive Zip1 lines (green) become detectable in wild type starting at t = 5 h, yet they are absent in *zip1-C1* (Fig. 6B) [note that likely due to day-to-day variability, the wild-type culture was delayed by 1.5 h in this experiment compared to cultures in Figs. 1 and 5]. Instead, PCs of increasing size become more frequent, until at t = 9 h, a strong PC, apposed to the nucleolus, is present in essentially all nuclei (Fig. 6B, Fig. S7C). Thus, Zip1-C1 is initially recruited to chromatin, yet stable linear Zip1 structures fail to assemble, and most Zip1 localizes to the PC ^73^.

**Figure 6.**
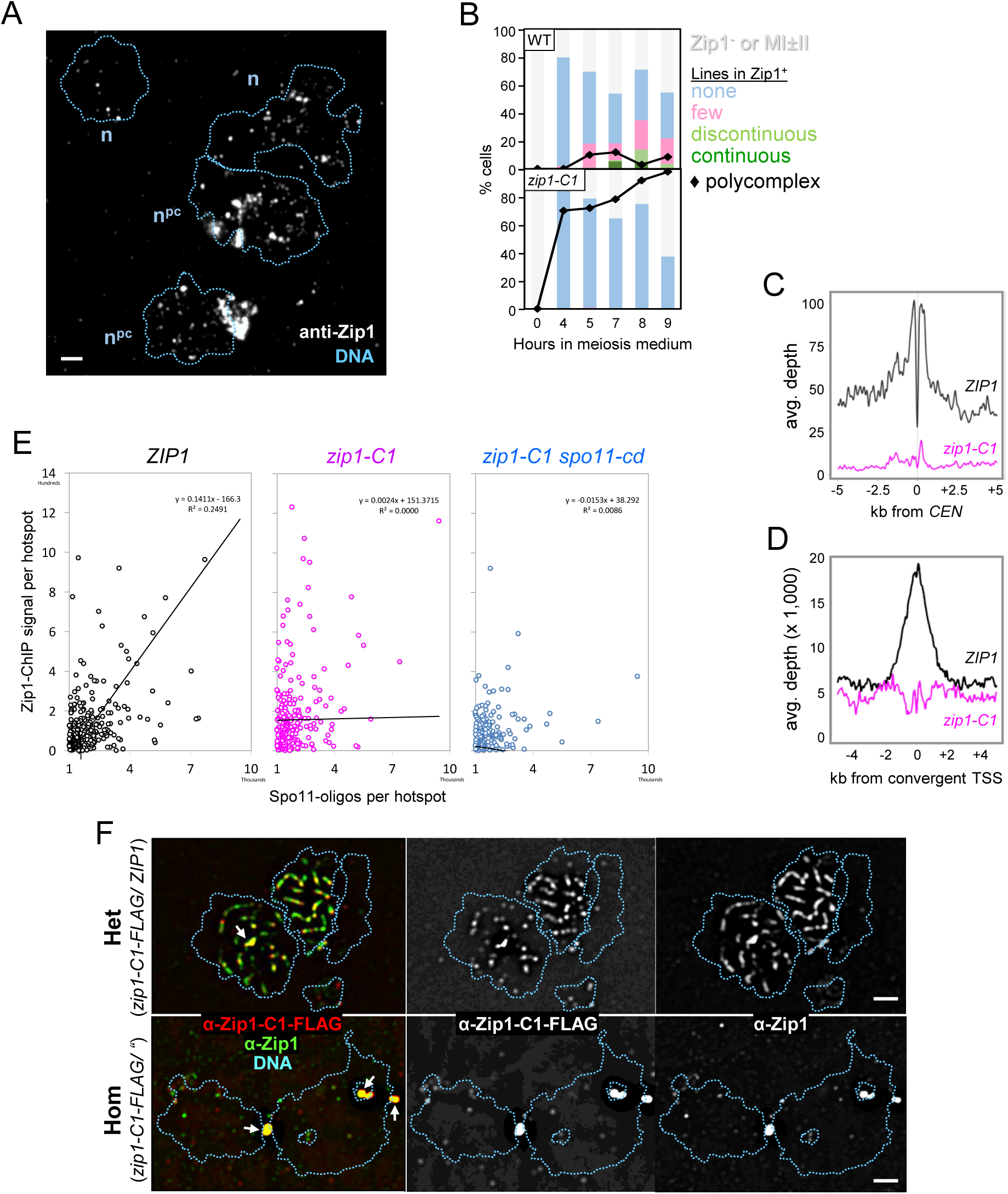
Zip1-C1 Association with Chromatin Independent of Synaptonemal Complex Assembly. **6A** Representative image field of four surface-spread *zip1-C1* nuclei at t = 4 h immunodecorated with anti-Zip1 antibodies containing no lines (n) with or without polycomplex (pc). The outside edges of DAPI-stained chromatin are traced by dotted lines. Bar 1 μm. **6B** Quantitative analysis of Zip1 localization on surface-spread nuclei in parallel WT and *zip1-C1* cultures (RiS tc15). **6C** Zip1 ChIP-Seq-profiles averaged over all chromosomes centered at their centromeres. *ZIP1*, 4 h in meiosis medium (black), *zip1-C1*, 6 h in SPM (magenta), later time point of *zip1-C1* compensates for its delay. **6D** Zip1 ChIP-Seq-profiles averaged over 3,900 convergent transcription sites (cohesin sites), centered at their mid points. *ZIP1* (black) and *zip1-C1* (magenta) after 4 h and 6 h in meiosis medium, respectively. **6E** Association of Zip1 with the 2,000 strongest DSB hotspots along the yeast genome in *ZIP1* (left), *zip1-C1* (middle) and *zip1-C1 spo11-YF* (right). R^2^ of linear regression is 0.25 for *ZIP1*, but 0 for *zip1-C1* or *zip1-C1 spo11-yf*. DSB hotspots are from Pan et al. (2011) ^103^. **6F** Immunofluorescence detection of internally tagged Zip1-C1-FLAG in heterozygous *zip1-C1-FLAG/ ZIP1* (top; n = 4) and homozygous *zip1-C1-FLAG/”* nuclei (bottom; n = 2). (tcRS11); antibodies: Rabbit anti-Zip1, mouse anti-FLAG. Arrows mark polycomplexes. Bar 1 μm.

To gain insights into the chromosomal positions associated with Zip1-C1, ChIP-on-chip analysis was carried out in WT and *zip1-C1,* using antibodies against full-length Zip1 ^35^. Three time points from two *zip1-C1* meiotic cultures were analyzed and compared to wild-type profiles for Zip1 and Rec8. During wild-type meiosis, Zip1 is prominently associated with core sites that also contain Rec8, likely constituting the cytologically-visible meiotic chromosome axis. Core sites include ∼5 kb regions surrounding the 16 centromeres as well as the 3,900 sites of convergent transcription, which are distributed along the yeast genome with semi-regular spacing of ∼20 kb ^35^ (Fig. 6C, D). In addition, we detected a weak positive correlation in wild type between Zip1 signals and DSB hotspots sorted by strength (Fig. 6E, black). In *zip1-C1*, association with both centromeres and convergent transcription start sites is eliminated (Fig. 6C,D), yet Zip1-C1 is detectable at multiple DSB hotspots, although the weak positive correlation between Zip1-C1 and hotspot strength is eliminated (Fig. 6E, red; Fig. S7D). An additional signal reduction in *zip1-C1 spo11-YF* indicates that Zip1-C1 association with DSB hotspots depends on DSB formation (Fig. 6E, right).

ChIP-on chip signals of Zip1-C1 at DSB sites are weak, but reproducible (compare signals from different time point samples in Fig. S7D). Thus, Zip1-C1 retains some ability to interact with chromosomes, a conclusion supported by its incorporation into SC when expressed in presence of wild-type Zip1. Accordingly, while FLAG-tagged Zip1-C1 is detected in a PC when homozygous, it is efficiently incorporated into SC, and PCs are essentially eliminated when Zip1-FLAG-C1 is expressed heterozygously with wild-type Zip1 (Fig. 6F).

Together, these findings have two key implications. First, Zip1 recombination functions at the DSB-to-SEI transition, including homolog bias and timely SEI formation, are mediated by Zip1-C1 independent of association with chromosomal core sites and of polymerization into linear SC. Second, Zip1-C1 appears to retain some capacity of interacting with chromosomal positions, including DSB sites. Zip1-C1 may localize to sites that are transiently or weakly occupied by wild-type Zip1, but such binding could be obscured in wild type by more prominent ChIP-on chip signals. Importantly, even though the DSB-to-SEI transition normally is completed in the context of fully assembled SC (Fig. 2), Zip1-C1 mediates this transition independent of SC polymerization.

### Precocious Zip1 polymerization allows double Holliday junction formation, but not their resolution

We next examined effects on synapsis and recombination of *zip1-L657A* which carries a leucine-to-alanine substitution in Zip1’s coiled-coil region ^74^. In the BR1919 strain background, a *zip1-4LA* mutant carrying *L657A* and three additional leucine-to-alanine mutations undergoes meiotic arrest with fully assembled SC, suggesting a defect in SC disassembly ^74^. In the SK1 strain background at 23°C, however, *zip1-L657A* completes meiosis efficiently, with a progression delay less pronounced than *zip1Δ* (Fig. 7A; Fig. S8A) ^74^.

**Figure 7.**
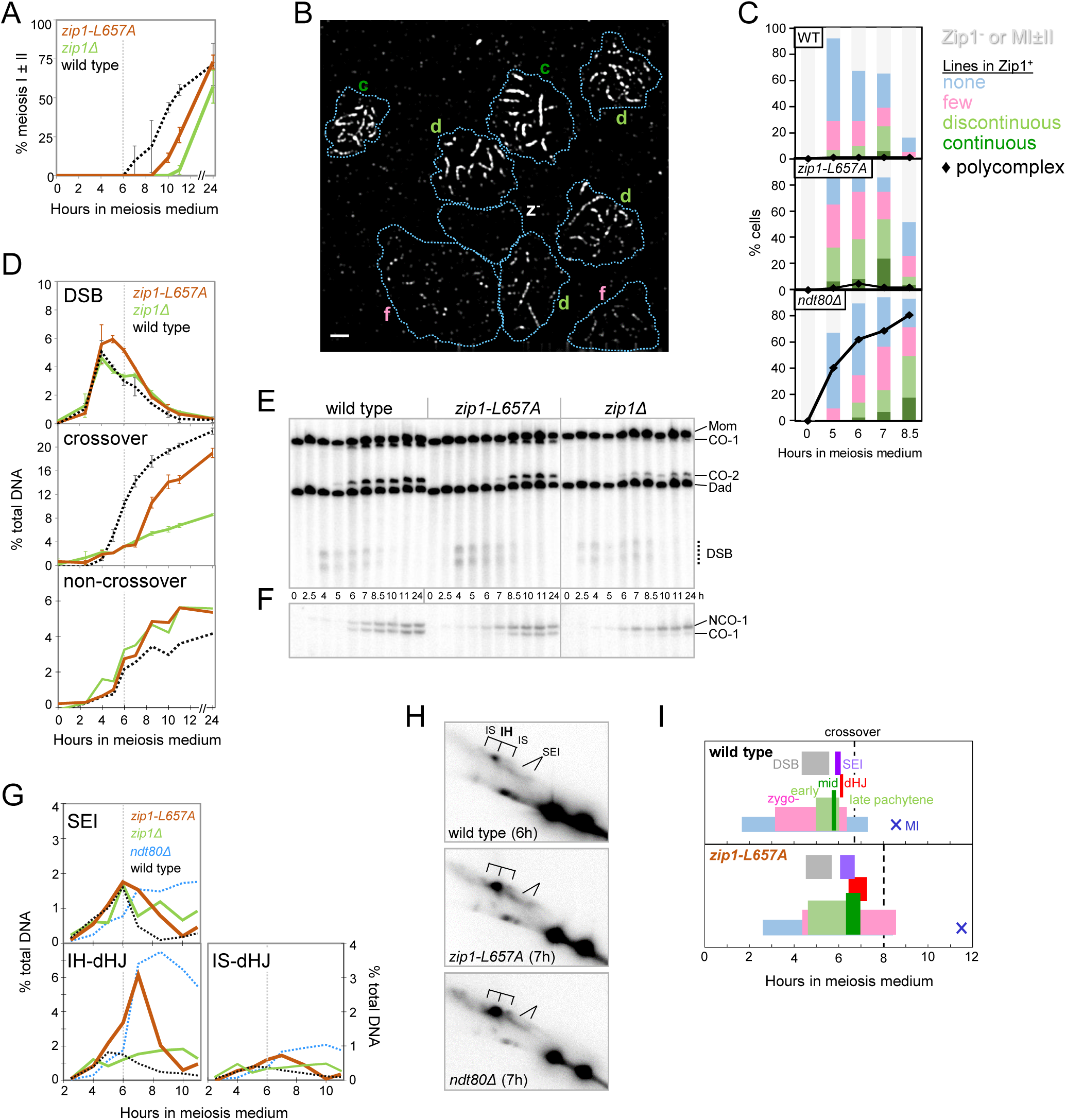
Premature Synapsis in *zip1-L657A* Reveals Zip1 Functions in Holliday Junction resolution (23°C). **7A** Meiotic nuclear divisions in *zip1-L657A*, *zip1Δ* and wild type (n = 2). Error bars indicate range. The dotted grey line here and in subsequent plots indicates t = 6 h. **7B** Zip1 localization to chromatin at t = 7 h in a representative field of *zip1-L657A* nuclei (n = 10). For Zip1 staining classes see Fig. 1A; for representative fields in wild type and *ndt80Δ*, analyzed in the same experiment, see Fig. 3C. Bar 1 μm. **7C** Quantitative analysis of Zip1 classes and polycomplexes in parallel cultures of WT, *zip1-L657A,* and *ndt80Δ* (tc42). **7D** Quantitative analysis of DSBs, crossovers and non-crossovers (Fig. 7E,F) in wild type, *zip1-L657A*, and *zip1Δ*. Error bars indicate range (n = 2) where available. **7E** One dimensional gel Southern blot analysis of recombination at *HIS4LEU2* in wild type, *zip1-L657A*, and *zip1Δ*. For recombination intermediates and products see Fig. 1D. For quantitation, see Fig. 7D. **7F** Excerpt from one-dimensional gel Southern blot showing CO-2 and NCO-2 products refractory to BamHI in a double digest with XhoI in the same DNA samples analyzed in Fig. 7E. For quantitation, see Fig. 7D. **7G** Quantitative analysis of SEIs, IH-dHJs and IS-dHJs in parallel *zip1-L657A*, *zip1Δ,* wild type and *ndt80Δ* cultures. For additional analyses of *ndt80Δ* in the same experiment see Fig. S4. **7H** Representative excerpts from 2D gel Southern blot analyses at time points with maximum JM levels of *zip1-L657A, ndt80Δ* and wild type. For recombination intermediates see Fig. 1F. For complete blots see Figure S8. **7I** Temporal relationship between recombination and synapsis in *zip1-L657A* and wild type. Wild-type data are from the culture shown in Fig. 2A (tc72) due to higher frequency of sampling. For details, see Fig. 2A.

To examine SC status as well as recombination in *zip1-L657A* under conditions of meiotic progression, nuclear spreads were analyzed in parallel with wild type, *zip1Δ* and *ndt80Δ*, with the latter serving as positive control for pachytene arrest (above). In wild type, pachytene nuclei (green) typically reach peak levels of ∼30% between 6 h and 7 h and remain detectable for approximately two hours (Fig. 7C; Table S2). By contrast, >30% of *zip1-L657A* nuclei are in pachytene already by t = 5 h, and they remain at high levels for approximately 4 hours (Fig. 7B,C,I). Furthermore, midpachytene nuclei (dark green) in *zip1-L657A* reach peak levels 3-times above wild type (Fig. 7B,C). Conversely, Zip1-positive nuclei that lack linear Zip1 structures (blue) are 2 to 3-fold underrepresented in *zip1-L657A* compared to wild type. In *zip1-L657A* compared to *ndt80Δ*, linear Zip1 staining classes (green) also appear much earlier, and PCs are essentially absent in *zip1-L657A* (Fig. 7C). Finally, most linear Zip1 structures have disappeared in *zip1-L657A* by t = 8.5 h, while they persist in *ndt80Δ*.

Together, these findings indicate that after a shortened leptotene stage, *zip1-L657A* cells precociously progress to zygotene and pachytene, resulting in a dramatic increase in the frequency of these stages. Exit from Zip1-staining occurs with some additional delay in *zip1-L657A* as indicated by disappearance of Zip1^+^ nuclei somewhat later than in wild type. Moreover, *zip1-L657A* differs markedly from *ndt80Δ,* as indicated both by the more efficient entry into the pachytene stage and the absence of PCs.

We next examined recombination in the same cultures. DSBs form efficiently in *zip1-L657A*, as indicated by WT-like DSB peak levels in the *SAE2* background (Fig. 7D,E; Fig. S8B,D), and accumulation in a *sae2Δ* background to 80% of *ZIP1* levels (Fig. S8E). DSBs further remain detectable for a longer time in the *SAE2* background, possibly due to a longer persistence at the DSB and/or SEI stages (below; Fig. 7D,E; Fig. S8D). Yet, DSB strand exchange occurs with normal timing and kinetics in *zip1-L657A*, as indicated by the appearance of substantial levels of SEIs and dHJs, as well as formation of both crossovers and non-crossovers (Fig. 7D,E; Fig. S8B-C).

Importantly, while SEI steady state levels are only moderately increased in *zip1-L657A*, IH-dHJs reach maximum levels 3- to 4-fold above WT peak levels, similar to persisting IH-dHJ levels in the parallel *ndt80Δ* culture (Fig. 7G,H). Accordingly, the lifespan of IH-dHJs is increased from 0.13 in wild type to 0.8 h in *zip1-L657A* (Fig. 7I). Consistent with delayed IH-dHJ resolution, crossovers appear with a delay of 2-3 hours in *zip1-L657A*, reaching 80% of WT levels, while they are reduced to 40% in *zip1Δ* (Fig. 7D,E). Resolution of dHJs in *zip1-L657A* is specifically impaired along the interhomolog pathway, as IS-dHJs accumulate to a lesser extent. Finally, spore viability in *zip1-L657A* is only moderately reduced compared to the WT (75% versus >90%; n > 390), but much higher than in *zip1Δ* (34%, above). Importantly, these data show that intermediate recombination defects and product formation in *zip1-L657A* are distinct from those in *zip1Δ* or *zip1-C1* (Fig. 7D-H; Fig. 5).

Analyses of recombination and SC assembly in the same cell population indicate that all cytological stages occur prematurely in *zip1-L657A* relative to recombination: While bulk DSBs still appear during zygonema, entry into SEIs and into IH-dHJs is advanced compared to wild type, occurring in early and midpachynema, respectively (Fig. 7I; above). Moreover, pachynema lasts 2.3 h in *zip1-L657A* compared to 0.9 h in wild type, indicating a 2.5-fold increase in the duration of this stage.

One explanation of these findings is that in *zip1-L657A*, Zip1 polymerizes prematurely, and normal controls of DSB first and second end strand exchange are defective, resulting in premature formation of SEIs and dHJs, but a delay in their resolution. Thus, whereas wild-type cells form dHJs only after SC disassembly, dHJs in *zip1-L657A* form efficiently in the context of assembled SC but fail to undergo timely resolution. Thus, assembled SC may interfere with IH-dHJ resolution although other cause-effect relationships cannot be excluded. We further do not consider as likely that delayed dHJ resolution is the cause of the SC disassembly defect in *zip1-L657A*. Notably, precocious SC assembly is manifest in *zip1-L657A* prior to dHJ accumulation, making it a more likely candidate for the primary defect. Finally, it is unlikely that defective meiotic progression is responsible for distinct recombination defects in the three *zip1* mutants analyzed here. Notably, both *zip1Δ* and *zip1-L657A* complete meiosis with similar efficiencies and timing, yet they exhibit very different recombination defects.

## Discussion

In the current work, we have defined the temporal and functional relationship between synaptonemal complex morphogenesis and recombination during yeast meiosis. We show that SC polymerization starts when a threshold number of interhomolog strand exchange intermediates, most likely paranemic D-loops, have been formed. Once SC is fully assembled, these intermediates are stabilized into plectonemic single end invasions. Double Holliday junctions appear as the SC disassembles, followed by their resolution into crossovers during diplonema. SC transverse filament protein Zip1 shepherds recombination intermediates through three transitions, both in the context and after disassembly of the SC, including DSB first as well as second end strand exchanges and double Holliday junction resolution. Even though SEIs normally form in the context of assembled SC, Zip1 mediates their formation independent of its polymerization or sustained association with major chromosomal binding sites.

### Unstable D-loop Intermediates License Synapsis and are Stabilized by Full Length SC

The present study yields novel insights into the temporal coordination between DSB first-end strand exchange and SC morphogenesis, with important implications for their functional interdependence. In budding yeast, SC assembly is dependent on recombination initiation, yet DSBs are not consistently associated with a single SC stage arguing against direct linkage between these stages (Fig. 2A). Instead, Zip1 polymerization into linear SC segments coincides with a distinct threshold number of crossover-specific interhomolog strand exchange intermediates. We consider as likely that an unstable interhomolog SEI precursor, rather than SEIs themselves, triggers SC initiation and extension. Notably, unstable protein-mediated side-by-side DNA interactions typically proceed formation of plectonemic D-loops mediated by base pairing ^75^. While such paranemic interactions are undetectable by standard 2D gel analysis ^11^, they likely share numbers and kinetics of SEIs, albeit shifted to an earlier time by one pre-SEI lifespan.

It is unlikely that SEIs themselves trigger synapsis, as this would predict two SEI waves, one prior to and the other following synapsis completion, a scenario incompatible with a precursor-product relationship of SEIs with dHJs. Our data also do not support a role of non-crossover formation in SC assembly. At 23°C, non-crossovers appear after completion of SC assembly, concurrent with dHJ entry ^14,20^. A proposed role of SEIs in zygotene entry ^11^ is not supported when events are monitored in the same culture. Finally, our findings are not at odds, with a model where synapsis mediates termination of DSB formation ^76,77^: Although DSBs persist into early pachynema under some conditions in our experiments, these breaks would have been formed prior to extensive SC polymerization (Fig. 2A-i).

Our data support the following scenario: Subsequent to DSB formation and 5’ strand resection, an early cohort of DSBs completes homology search and undergoes paranemic interactions with the homolog template. Such paranemic interactions may persist for a long time or involve alternative partner molecules, including the sister chromatid ^18^. One strand exchange along a given homolog pair may initiate SC initiation, detectable as the leptotene-to-zygotene transition on a per nucleus basis, whereas the zygotene-to-early pachytene transition may be licensed by an additional pre-SEI along the same homolog. Assuming that (pre-)SEIs are crossover-specific intermediates and that at 30°C, 95 crossovers are formed per yeast meiosis ^16^, these findings suggest that cells enter zygonema when ∼12 future crossover sites have progressed to a pre-SEI intermediate, whereas ∼29 pre-SEIs are associated with progression to early pachynema. These estimates roughly correspond to one pre-SEI per homolog (16) and one pre-SEI per chromosome arm (32), respectively. Whereas the first transition may be reversible, the second would likely result in stable synapsis.

A currently unknown event subsequently stabilizes discontinuous, early pachytene SC into continuous midpachytene Zip1 lines, with stable homolog juxtaposition facilitating the pre-SEI to SEI transition. Appearance of continuous SC at the midpachytene stage could indicate that gaps in the SC have been filled, or more likely, that Zip1 association becomes stabilized along the length of homolog axes. Notably, SC morphogenesis involves continuous turnover of transverse filament proteins such as Zip1 ^78–80^. The transition from early to midpachytene observed here is reminiscent of a switch in *C. elegans* from a highly dynamic to a more stable SC conformation ^80–82^. SEIs may form as consequence of stabilized homolog juxtaposition, but a reverse cause-effect relationship or parallel outcomes cannot be excluded. Notably, SC-mediated global axis juxtaposition is not indispensable for (some) stable strand exchange, as suggested by substantial SEI formation in synapsis-defective Zip1-C1 variant (Fig. 5; below).

### Double Holliday Junctions are Formed and Resolved into Crossovers during Diplonema

The widely held notion that double Holliday junctions are generated during pachynema and resolved into crossovers at pachytene exit stems from analyses of *ndt80* mutant meiosis, together with kinetically adjusted cytological and recombination data ^11,69^. Our analysis challenges this model, revealing that crossovers appear up to 40 minutes after pachytene exit when monitored in the same wild-type culture (Fig. 2A-I; Table S2). If crossovers form during wild-type diplonema, it follows that dHJs, as crossover precursor, also persist well into diplonema, consistent with IH-dHJ resolution post pachytene exit (Fig. 2A). Additional evidence indicates that dHJs are not only resolved but also formed during diplonema. First, dHJ entry follows pachytene exit under conditions of optimal resolution of later time points (Fig. 2A-i,ii,iv). Second, using crossover timing as a reference point and factoring in an average IH-dHJ lifespan of less than 10 minutes (at 23°C and 30°C), dHJ entry also falls after pachytene exit. Moreover, some dHJs may generate (class II) non-crossovers ^83^, further shortening the lifespan of crossover-bound dHJs and placing their formation later into diplonema.

While dHJ formation and resolution occur post SC disassembly, the SC’s impact on both transitions remains noteworthy. Zip1 post-SC disassembly roles in dHJ formation and resolution are suggested both by impaired transition from SEIs to dHJs in *zip1-C1* (Fig. 5), and transient dHJ accumulation in *zip1-L657A* (Fig. 7). Zip1 may control post-SC recombination steps by sustained association with specific chromosome positions, potentially guiding dHJ formation and/or resolution, a notion supported by Zip1 retention at Zip3-marked crossover sites in post pachytene nuclei. Zip1 may further establish spatial constraints during pachynema that outlast SC disassembly, and that are conducive to diplotene dHJ resolution. Consistent with a change in SC architecture in *zip1-L657A*, axis protein Red1 is retained in continuous lines along the lengths of homolog axes, and homologs are separated by a widened gap in the related *zip1-4LA* mutant ^76^ (not shown). While IH-dHJs normally form during diplonema, substantial SC may interfere with their resolution when they emerge precociously under certain mutant conditions. Accordingly, both *ndt80Δ* and *zip1-L657A* display dHJ formation despite faulty pachytene entry, followed by failures in timely dHJ resolution.

### Zip1 Controls Recombination at Three Transitions

The impact of SC central element proteins, particularly Zip1, on the outcome of recombination has been extensively studied, yet understanding their roles at intermediate recombination steps has proven challenging. Our findings show that Zip1 guides recombination intermediates through three crucial transitions along the crossover pathway. First, at the DSB-to-SEI transition, Zip1 ensures preferential recombination with the homolog and efficient formation of IH-SEIs, functions impaired in *zip1Δ* but (partially) intact in *zip1-C1* (Fig. 5F) ^15^. Second, Zip1 mediates the SEI-to-IH-dHJ transition, as suggested by delayed appearance of the latter intermediate in *zip1-C1* (Fig. 5G). Third, Zip1 plays a role in the resolution of IH-dHJs into crossovers, as indicated by transient IH-dHJ accumulation and their delayed resolution in *zip1-L657A* (Fig. 7).

Our findings indicate that Zip1 exerts some of its functions independent of assembly into SC, as suggested by SEI formation in *zip1-C1* despite a >100-fold shortening of the lifespan of linear Zip1 structures (Fig. 5C). This provides direct evidence for a polymerization-independent Zip1 function during SEI formation, as further indicated by substantially higher levels of dHJ and crossover formation in *zip1* mutants where synapsis is limited to individual chromosomes or undetectable compared to *zip1Δ* ^84,85^. Notably, Zip1 influences recombination even when SC formation is eliminated due to the absence of axis protein Red1 ^86^.

Zip1’s impact may occur through direct effects on recombination or by recruiting other recombination factors. Stalling of recombination at the SEI stage in *zip1-C1* resembles defective DSB second end capture in *rad52* ^12^, as well as hypomorphic *spo11* ^20^. Moreover, *zip1-4LA* fails to recruit axis remodeler Pch2 to meiotic chromosomes, and like in *zip1-L657A*, IH-dHJs accumulate in *pch2Δ* and the pachytene stage is extended, albeit not under conditions examined here ^76,87^.

### Interdependence of Synapsis and Recombination

Despite the evolutionary conservation of recombination genes and SC morphology, the extent to which temporal and functional coordination exists between these processes has remained unclear ^88^. Importantly, the timing of cytological markers relative to their activity along meiotic chromosomes involves uncertainties. While during both yeast meiosis and mouse spermatogenesis cytological markers of DSBs and/or strand exchange appear prior to and are displaced following SC assembly ^45,76,89,90^, these patterns are not universal. Accordingly, in *C. elegans*, *Arabidopsis*, and human oogenesis, DSB markers persist into pachynema ^46,47,91^. Moreover, proteins crucial for crossover formation including Hei10, are limited to pachytene chromosomes in male mice ^92^, yet they persist into diplonema in plants (*Arabidopsis*) and in the fungus *Sordaria*, as also observed for COSA-1 in *C. elegans* ^48,50,93^. Similarly, during male mouse meiosis, MLH1 foci are displaced from the SC at pachytene exit, yet they persist into diplonema during oogenesis ^90^. Finally, detection of (some) crossovers and non-crossovers at the time of pachynema during the first synchronous wave of mouse spermatogenesis is complicated by their continued increases during diplonema and overlaps with subsequent waves of spermatogenesis ^89^. The present investigation yields vital insights into these events, serving as a benchmark for comparative studies in other organisms.

Mutant analyses across diverse organisms also suggest varied functions of the synaptonemal complex in recombination. In yeast, the SC mediates multiple recombination transitions, from strand exchange and resolution of double Holliday junctions to terminating DSB formation (above) ^15,77^. Additionally, Zip1 blocks an alternative recombination pathway lacking crossover interference and patterned crossover placement ^94^. These findings imply that the SC influences the formation and placement of crossovers by guiding progression along the patterned crossover pathway and blocking alternate pathway(s). An SC role in DSB strand exchange is suggested by observations in mouse and *C. elegans*, where in the absence of transverse filament proteins recombination stalls at intermediate stages, as indicated by the persistence of Rad51 foci ^46,95^. Conversely, elimination of *Arabidopsis* ZYP1a/b yields extra crossovers, defective crossover interference, and absence of obligatory crossovers on some homologs, but no defect in recombination progression ^54,96^. Thus, the use of alternative recombination pathway(s) in the absence of transverse filament proteins is not always accompanied by intermediate recombination defects.

The role of the SC in the non-random placement of crossovers is also subject of debate (Introduction). Appearance of patterned Zip3 foci at or before the zygotene stage, as observed here, suggests that crossover designation occurs prior to and independent of SC formation, consistent with appearance of interference-spaced late recombination nodules before SC assembly in *Sordaria* (Fig. 2C) ^15,39,68,97^. Notably, Zip3 is critical for SEI formation and may associate with crossover-designated pre-SEIs at or shortly after DSB entry (Fig. 2C) ^15^. Conversely, the interference distribution of cytological crossover markers and/or crossovers is impaired in absence of the SC in *Arabidopsis* and *C. elegans*, potentially indicating species-specific aspects in the establishment or maintenance of patterned crossover placement ^98^.

In summary, our study has unveiled the temporal and functional relationship between all known recombination intermediates and products and assembly as well as disassembly of the synaptonemal complex. Our findings provide a benchmark for a mechanistic understanding of the intricate relationship between these processes in yeast and other organisms.

## Acknowledgments

We thank Rachel Soucek, Hanna Henley, and Steven Zimmerman for help with experiments; Monique Zetka, Judith Yanowitz, Aaron Severson, Amy MacQueen, Nancy Kleckner, Jesus Monge Neria as well as other Börner lab members for discussion; Scott Keeney, Neil Hunter, Angelika Amon, and Shirleen Roeder for antibodies, strains and plasmids. Research reported in this publication was supported by NIH grant R01GM141698 (to G.V.B.) and S10OD025252 (to Bibo Li), as well as SFB F 8807-B from the Austrian Science Fund (to F.K).

## Methods

### Strain construction

For strains see Table S4. All strains are isogenic SK1 strains, and the indicated alleles were confirmed by Southern blot analyses and/or sequencing. The *zip1-C1* internal deletion strain RSY152 was generated by targeting for integration at *ZIP1* MfeI-digested *URA3*-marked integrating plasmid pTP6 ^73^ which carries the *zip1-C1* construct (gift from Shirleen Roeder). Following transformation into haploid VBY826 (*HIS4::LEU2, ura3, nuc1:HYGMX4*), uracil prototrophic transformants were confirmed by Southern blot analysis with HpaI. The resulting strain VBY1036 which carries the *zip1-C1* internal deletion followed by a *zip1* variant lacking its promoter region and start codon was incubated on 5-FOA to select for popout of the *URA3* marker. The *zip1-C1* internal deletion was confirmed by Southern blot analysis. The resulting strain was mated to hotspot containing strain VBY1056 (*his4X*::*LEU2*-*URA3*) and haploid spore colonies carrying both *HIS4::LEU2* hotspot versions but devoid of *nuc1::HYGMX4* were identified and mated giving rise to RSY152.

A FLAG epitope was introduced into *zip1-C1* at amino acid position 705 via the following steps: Following introduction of a single FLAG-encoding 27bp segment via overlapping PCR ^99^, a PfoI-AatII fragment was excised from the PCR product and transferred into BS plasmid pVB28 which carries full length *ZIP1* with *URA3* inserted into its 5’ intergenic space at PstI. The *ZIP1* region containing the C1 internal deletion was PCR amplified from pTP6 ^73^, inserted between AatII and SacI, to generate pJas55D. A SacI-SalI fragment containing *URA3-zip1-FLAG@705-C1* was transformed into recipient strain RSY128 (*HIS4LEU2, ura3, zip1::KANMX4*), and the resulting uracil prototrophic, G418 sensitive strain was crossed with VBY1056 to generate homozygous RY124 (*URA3-zip1-FLAG@705-C1/”*) as well as heterozygous RY31.

Similarly, a *URA3*-marked version of *zip1-L657A* was generated by replacing in pVB28 the BlpI-AatII fragment carrying the *L657A* point mutation from plasmid NMB201 ^74^ (gift from Shirleen Roeder). The resulting XbaI fragment carrying *URA3-zip1-L657A* was transformed into recipient strain RSY128, and uracil prototrophic, G418 sensitive transformants were confirmed by Southern blot analysis. The *URA3-zip1-L657A* construct was subsequently combined with both versions of the *HIS4LEU2* hotspot in diploid strain RSY257.

### Synchronous meiotic cultures

Synchronous meiotic time courses were performed as described ^100^. Briefly, strains frozen in 25% glycerol at −80°C were patched on YPG plates (1% Bacto yeast extract, 2% Bacto peptone, 3% glycerol, 2% agar) and incubated at 30°C for ∼16 hrs. Patches were streaked for single colonies on YPD plates (1% Bacto yeast extract, 2% Bacto peptone, 2% dextrose, 2% agar) and incubated at 30°C for ∼54 h. Single colonies were inoculated in 4 ml liquid YPD and incubated on a roller drum at 30°C for 26 h. Saturated cultures were inoculated at dilutions ranging between 1:100 to 1:200 in 150 ml pre-warmed YPA (1% Bacto yeast extract, 2% Bacto peptone, 1% potassium acetate) and incubated with vigorous shaking at 30°C for 13.5 h. Following growth and G1 arrest in YPA, cultures exhibiting ODs of ∼1.2-1.5 were selected. Cultures were washed and resuspended in meiosis medium (0.5 % potassium acetate, 0.02 % raffinose, 1:1,000 antifoam) and incubated at the indicated temperatures. Transfer to meiosis medium was set to t = 0 h. Meiotic cultures at 23°C and 30°C were incubated at the respective temperature starting at t = 0 h in appropriately calibrated media. Cultures analyzed at 33°C were incubated at 26°C for three hours, followed by an increase of the incubator temperature to 33°C, resulting in warming of the medium to 33°C within ∼30 min.

### Meiotic progression

Aliquots between 0.1 to 0.5 ml were removed at various time points over the duration of the time course. Aliquots were mixed at a 1:1 ratio with fixative (80% ethanol, 100 mM sorbitol, 0.5 mM EDTA) and stored at −20°C. Fixated cell aliquots were stained with DAPI dye (1 μg/ ml) and the number of nuclei per cell was counted at each time point using a fluorescence microscope. At least 100 cells per time point of each culture were counted.

### Tetrad dissection and interference analysis

Samples were collected at t = 24 h following transfer of cells to liquid meiosis medium. Tetrads were analyzed from the same cultures used for physical analysis and progression analysis. Cells were digested with zymolyase for 30 min at 37°C followed by dissection of yeast asci. Plates were incubated at 30°C for ∼3 days, until spore colonies were visible. Spore viability was calculated by dividing the number of viable spores by the total number of spores dissected. Calculations of genetic map distances, modified coincidence analysis and NPD ratios were performed as described by https://elizabethhousworth.com/StahlLabOnlineTools/ and by Malkova et al. (2004) ^71^.

### Meiotic chromatin spreads and immunocytology

Aliquots (1 ml to 10 ml) from synchronized meiotic cultures were collected at the indicated time points and chromatin spreads were prepared as described ^101^. Briefly, cells were washed in Tris buffer and spheroplasted using zymolyase 20T. After spheroplasting, cells were washed and resuspended in MES buffer. Spheroplasted cells were crosslinked on pre-cleaned glass slides using 3% paraformaldehyde/ sucrose solution. After crosslinking, cells were lysed using 1% lipsol. Lysed cells were spread on the slide using a glass serological pipette. Spreads were allowed to dry overnight in the fume hood.

For immunocytology, slides carrying spread meiotic cells were washed once with TBS. Non-specific binding of antibody was minimized by blocking spreads with 1% BSA in a moist chamber for 10 minutes. Following blocking, appropriate primary antibodies in TBS/BSA solution were used to immunodecorate samples. For Zip1 detection, rabbit anti-Zip1p antibody was used (gift from Scott Keeney, Memorial Sloan Kettering, NY). Zip3-GFP was detected using goat anti-GFP (Rockland). Spreads were incubated with primary antibodies overnight at 4°C. Slides were subsequently washed with TBS and probed with appropriate secondary antibodies. Secondary antibody incubation was performed for 2 hours at room temperature. All incubations were carried out in a moist chamber. Following secondary antibody incubation, slides were washed with TBS and stained with DAPI dye (1 μg/ ml). Spreads were then mounted in ProLong Diamond.

### Image acqusition and scoring of Zip1 classes

Using a 100x 1.47NA objective, fields of between one and 20 surface spread nuclei were identified based on DAPI staining and image stacks were acquired using a Deltavision imaging system, at appropriate wave lengths. Following deconvolution, individual non-overlapping nuclei were cropped to size from larger images and Z-sections were saved as tiff files with unique identifiers. In-focus Z-sections were identified in the DAPI channel using the “Find_Focused_Slices” ImageJ plugin. Zip1 classes from a given experiment were scored without knowledge of time point, genotype or incubation temperature using photographed nuclear spreads. Using ImageJ, each image was contrast adjusted with 15% saturated pixels (“Enhance Contrast”, “saturated = 0.15”). All channels from the same Z-section were saved in separate folders, and the Zip1 channel was set to min-max values of 200 and 1,200, respectively. Background staining was masked by setting the minimum displayed value to 100 on a 16-bit image.

A list was prepared comprising filenames corresponding to the nuclei to be scored (see PDF S1). From this list, sets of 200 nuclei were opened at a time, without knowledge of genotype, incubation condition or time point. Excised nuclei were individually presented to the scorer and nuclei with similar Zip1 morphologies were assigned to one of 12 distinct positions on the computer screen, during several rounds of iterative sorting. Following sorting, the screen location (X,Y) of each image was stored in a log file. Results from this first round of scoring were integrated into a single list, and nuclei from this list underwent two additional rounds of sorting, thus reaching a continuum of morphologies with each pile having essentially identically stained nuclei. The process of file opening from a list and logging the screen location of sorted images was automated using JavaScript. Cells containing two or more nuclei adhere poorly to glass slides and therefore were omitted from scoring. To account for cells containing divided nuclei, an appropriate number of divided cells obtained from DAPI counts of intact cells was added to the cells scored to obtain the fraction in each class of total nuclei.

### Physical Analysis

#### Crosslinking and DNA extraction

Aliquots from meiotic cultures were pelleted and crosslinked using trioxsalen and exposure for 10 minutes on a long wave (360 nm) UV light box. For DNA extraction, cells were spheroplasted using zymolyase and processed using the SDS lysis method ^100^. RNase A, proteinase K and phenol chloroform treatment were also performed as described. Following phenol chloroform extraction and ethanol precipitation, DNA pellets were resuspended in TE buffer and allowed to rehydrate overnight at 4°C prior to restriction digestion. For analysis of DSBs, crossovers, and joint molecules, ¼ of DNA was digested with 60 U XhoI in 80 μl overnight at 37°C. To detect non-crossovers, DNA digested overnight with XhoI was further digested for ∼5 h with HF-BamHI or NgoMIV (New England Biolabs). Digested DNA was precipitated using standard ethanol/ sodium acetate precipitation.

#### One-dimensional and two-dimensional gel electrophoresis

To detect DSBs, COs, and NCOs, appropriately restriction digested DNA was run on 0.6 % agarose gels without ethidium bromide at 2 V/cm for 26 hours. Southern blotting was performed using alkaline transfer to nylon membrane (Amersham N+). For joint molecule detection, 2D gel electrophoresis was performed as described ^100^. Briefly, XhoI digested DNA was separated on a 0.4 % Seakem Gold agarose gel without ethidium bromide at 35 V for 17.5 h. Following staining with ethidium bromide, gel slices ranging from 12 to 3 kb were excised as indicated by a comigrating molecular weight marker using long wave UV light. Gel slices were placed horizontally on a separate gel tray. Following lane excision, 0.8 % agarose containing ethidium bromide was poured around the gel slices in a cold room. Electrophoresis was performed in the second dimension at 5 V/cm for ∼5 h in a cold room. Following electrophoresis, Southern blotting was performed using SSC transfer of DNA to uncharged nylon membrane. After transfer, the membrane was crosslinked with UV light.

Membranes were hybridized with “probe 4” for the *HIS4::LEU2* hotspot ^100^ which was ^32^P-dCTP labelled using the Prime-It II Random Primer Labeling Kit (Stratagene). Following hybridization, membranes were exposed to imaging plates and scanned using a Typhoon phosphoimager. Non-overexposed scans of 1D and 2D Southern blots were used for quantitation using Quantity One software (BioRad).

### Cumulative and modified cumulative analyses

Cumulative analysis for recombination intermediates, Zip1 and Zip3 staining as well as of meiotic progression was carried out using the previously described CumKin 2.0 Macro ^20^. This macro converts steady-state levels into cumulative values as follows (for details see Info tab in Supplemental Excel S1): First, the lifespan of a given intermediate is determined by dividing the area under the steady state curve by the total number of events. Second, starting from the peak time point of the respective steady state curve, steady state levels at one-lifespan intervals are interpolated in both directions. The first non-zero steady state value at an interpolated time point is transferred into the cumulative curve as is. Proceeding in one lifespan steps, cumulative values are obtained by adding to the steady state level at each interpolated time point all steady state levels from previous interpolated time points. The time when the resulting cumulative curve reaches 50% of its maximum value provides the average 50% entry time, whereas the 50% exit time corresponds to that time point plus one lifespan. Cumulative blots show cumulative levels of each intermediate at the taken rather than the interpolated time points.

For cumulative analysis of recombination intermediates and products several considerations further are relevant: The total number of DSBs was obtained from maximum DSB levels detected in *sae2/com1Δ* or *rad50S* backgrounds under the respective conditions, i.e. 23% (23°C), 20% (30°C), and 13% (33°C) (see Table S1). For cumulative analysis of IH-SEIs and IH-dHJs, the following assumptions were made: First, IH-SEIs and IH-dHJs progress exclusively to crossovers ^15^, and all crossovers are formed via IH-SEIs and IH-dHJs. Second, as one IH-dHJ molecule is resolved into two crossover bands, their total number corresponds to the number of crossovers under the respective conditions, i.e., 22% (23°C), 20% (30°C), and 15% (33°C). Third, the majority of SEI signals are derived from IH-SEIs, and IS-SEIs contribute negligibly to the SEI signal, consistent with absence of the crescent-shaped signal characteristic for IS-SEIs ^22^. Fourth, the two well-resolved IH-SEI signals typically detected on 2D blots [SEI-3 (7.4 kb) and SEI-4 (8.7 kb) ^11^] are representative of two additional IH-SEIs (SEI-1 and SEI-2) that occur at *HIS4LEU2*, but are not well resolved. Accordingly, we assume that we roughly detect half of IH-SEIs corresponding to half the number of IH-dHJs.

DSB and all JM signals were set to zero one hour after the last time point analyzed (t = 10 h or 12 h), thereby avoiding unrealistic lifespan extensions from persisting intermediates at <10% of peak levels. Such apparently persisting intermediates are attributable to the preferential extraction of DNA from a minority of cells unable to complete meiosis from which DNA is extracted more easily than from cells engulfed by a thick spore wall. For crossovers, the CumKin 2.0 Macro was also used.

For cytological stages including Zip1 and Zip3 staining patterns, cumulative analysis was further modified in several ways. Notably, only the stage with the most extensive staining (Zip1 continuous lines or ≥ 36 Zip3 foci) is unambiguous, whereas steady state curves for all other stages represent composites of assembly and disassembly curves due to the morphological similarities between these stages. At the same time, for cytological analysis it is ascertained that all intermediate stages of assembly and disassembly are quantitatively detected. We therefore carried out standard cumulative analysis for the cytological stages with the most extensive staining, followed by modified cumulative analysis for other stages where lifespans as well as entry and exit times for combined stages were determined. For example, for the early pachytene (EP) and late pachytene stages (LP), the primary steady state curves for ‘continuous’ and ‘discontinuous’ line stages were summed up, the lifespan for the combined stages was calculated, and used to determine the combined cumulative entry curve for cells containing discontinuous or continuous lines, which corresponds to the early pachytene entry curve. Shifting this curve by the combined lifespan to the right provided the exit curve from late pachynema. For the ‘few’ Zip1 line stage, this approach was further modified by calculating the entry curve using the steady state curve and calculated lifespan for ‘few’ lines, whereas the exit from that stage was obtained by shifting this curve to the right by the combined lifespan obtained from the sum of few, discontinuous and continuous lines. This approach delays the 50% entry time for zygonema by 0.4 h (23°C) and 0.6 h (30°C) but avoids calculation of entry curves using combined lifespans which at 3.2 h (23°C), 2.9 h (30°C), and 2.6 h (33°C) understate the temporal resolution of our analysis (for spreadsheets with calculated entry curves see Supplemental Excel S1). Finally, for leptotene entry, the intrinsically cumulative Zip1+ entry curve was used as is, a curve that corresponds to nuclei of meiotically active cells (∼95%). The exit curve from diplotene was obtained by shifting the Zip1+ entry curve to the right by the lifespan of all Zip1+ stages. To ensure comparability between our data and those from earlier EM analysis ^32^, the same modified cumulative analysis was used for recalculating entry and exit curves for that experiment, providing very similar results for 50% entry and exit times obtained from the earlier analysis (see Table S1). Notably, only our approach is truly cumulative as the cumulative curve never dips below levels at an earlier time point as apparent in the original analysis^32^.

For the meiotic culture analyzed at 33°C, no 10-hour time point was available due to culture volume limitations. Maximum crossover levels for that time point were obtained from four cultures performed with the same strain under identical conditions (tc51-5,6 and tc55-1,2), and levels for recombination intermediates were set to zero at t = 10 h, consistent with extrapolation of their descending levels from the peak levels and levels observed in other experiments performed under identical conditions.

### Chromatin immunoprecipitation and bioinformatics

Chromatin immunoprecipitation was carried out largely as described using affinity purified antibody #214 raised in rabbit against full length yeast Zip1 ^35^.

#### Noise Filtering

Micro arrays were read using Affymetrix software (GCOS) and exported as CEL files. These were converted to text files using TAS (Tiling analysis software) from Affymetrix, computing ChIP-Seq/WCE. Coordinates represent S288C reference genome R64-2-1. The resulting data were noise filtered by sliding window smoothing using a bandwidth of 500 bp. Data were processed using R version 2.6.2 (http://www.r-project.org/) and displayed using Microsoft Excel. Cumulative plots at DSB sites and at sites of convergent transcription were produced as described ^102^ with DSB signals taken from ^103^

#### Profile Normalization

Following noise filtering, the signal intensities of profiles were normalized using NCIS ^104^ so that the 10% lowest values fall below 1 (decile normalization). The lowest 10% binding sites are interpreted as background. Decile normalization robustly projected biological repeats on top of each other, as it eliminates experimental variations and conserves the relations between peak sizes. No further normalization was applied.

