## Supplementary material for "Temporal and Functional Relationship between Synaptonemal Complex Morphogenesis and Recombination during Meiosis": Figures S1-S8;Tables S1 to S4;PDF S1;Excel S1

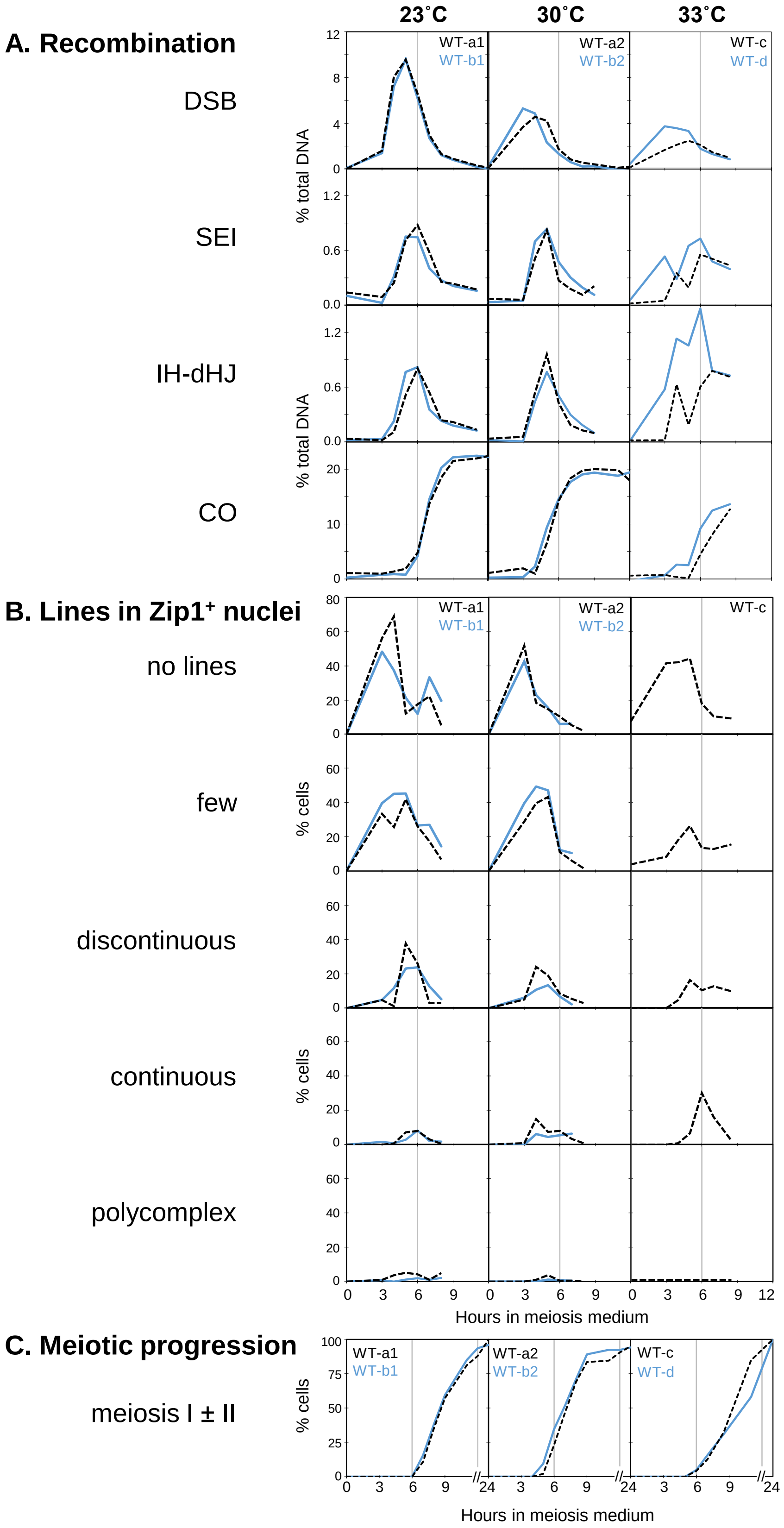

**Figure S1. Reproducibility of Recombination and Zip1 Recruitment during WT Meiosis at Three Temperatures**

**S1A** Analysis of recombination intermediates and products in WT at *HIS4LEU2* at 23˚C, 30˚C and 33˚C. Grey vertical lines indicate t = 6 h. Cultures are from a single experiment (TC72) except for WT at 33˚C (TC55). Cultures designated by the same lower-case letters are derived from a single pre-meiotic culture, with number extensions indicating incubation temperatures 23°C (1) or 30°C (2).

**S1B** Zip1$^{+}$ classes in cultures also analyzed for recombinationv (see Fig. S1A). For numbers of nuclei scored for each time point see Table S2.

**S1C** Meiotic nuclear divisions. Vertical lines indicate the t = 6 h time point, as well as the time scale interruption between 11 h and 24 h.

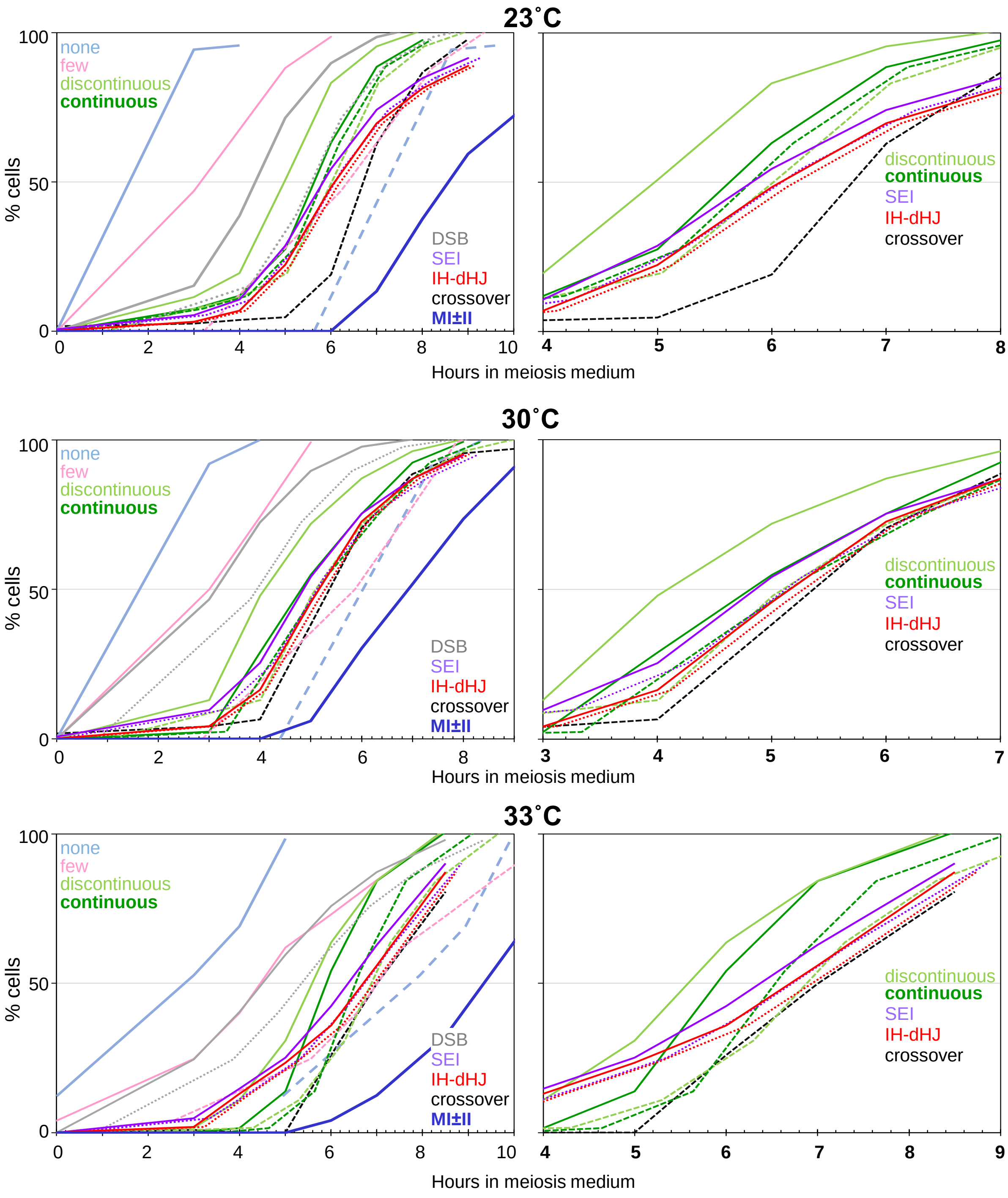

**Figure S2. Cumulative Analysis of Synapsis, Recombination and Meiotic Progression in Wild Type at Three Temperatures**

Continuous lines indicate entry, dotted and dashed lined indicate exit from an intermediate stage. In left panels, cytological stages from leptotene to diplotene and meiotic divisions are shown together with recombination progression. Right panels provide detailed analyses of early, mid and late pachytene as well as single end invasions, double Holliday junctions and crossover recombinants.

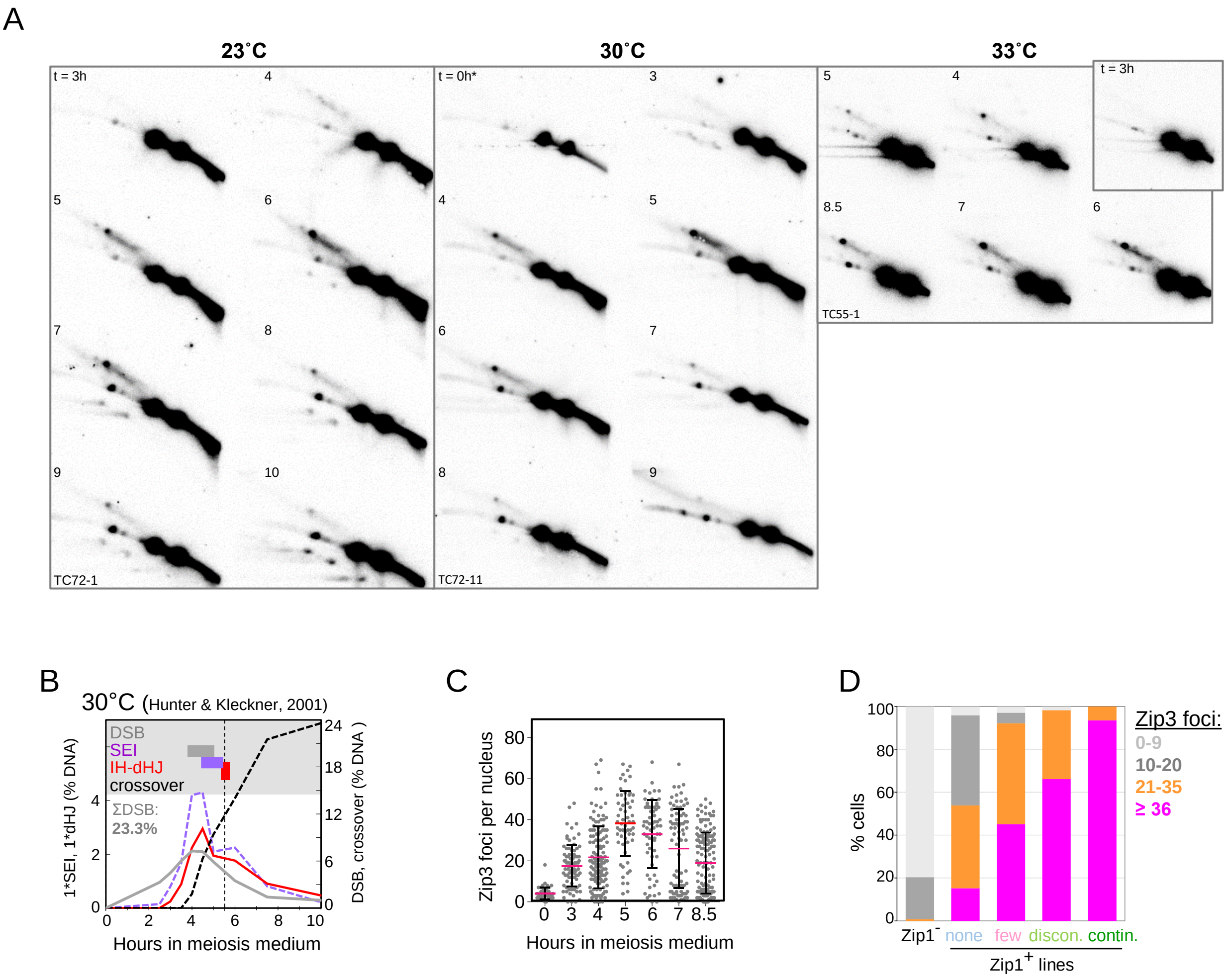

**Figure S3 Recombination at *HIS4LEU2* and Zip3 Recruitment during Wild-Type Meiosis.**

**S3A** Complete 2D gel Southern blot analyses of representative cultures at 23˚C (TC72-1), 30˚C (TC72-11) and 33˚C (TC55) quantitated in Fig. 1C. Asterisk: The 23˚C and 30˚C cultures share the t = 0 h sample derived from the same premeiotic culture.

**S3B** Recombination intermediates and products at a *HIS4LEU2* hotspot variant (*NBamHI/ NBamHI;* data from Hunter and Kleckner, 2001). SEIs and IH-dHJs (left y-axis), DSBs and COs (right y-axis). JM maximum steady state levels are 2 to 3-fold higher compared to the *HIS4LEU2* (*BamHI/NgoMIV*) variant analyzed here, as further suggested by parallel analysis of the two hotspot variants (Fig. S2B in Joshi et al., 2015) and analysis by others (e.g. Oh and Hunter, 2007).

**S3C** Number of Zip3-GFP foci in a meiotic WT culture at 33°C also assayed for Zip1 and recombination (Fig. 1C,D). Data are from a mock culture to which DMSO was added at t = 3 h, 5 h and 7 h (Ahuja et al., 2017). Red lines indicate means, error bars are standard deviations.

**S3D** Zip3-GFP focus numbers in five classes of Zip1 staining nuclei exhibiting no Zip1 (Zip1$^{-}$) or increasing abundance of Zip1 lines (none, few, discontinuous or continuous). All nuclei between 0 h and 8.5 h from Fig. S3C (n = 610) were included in this analysis.

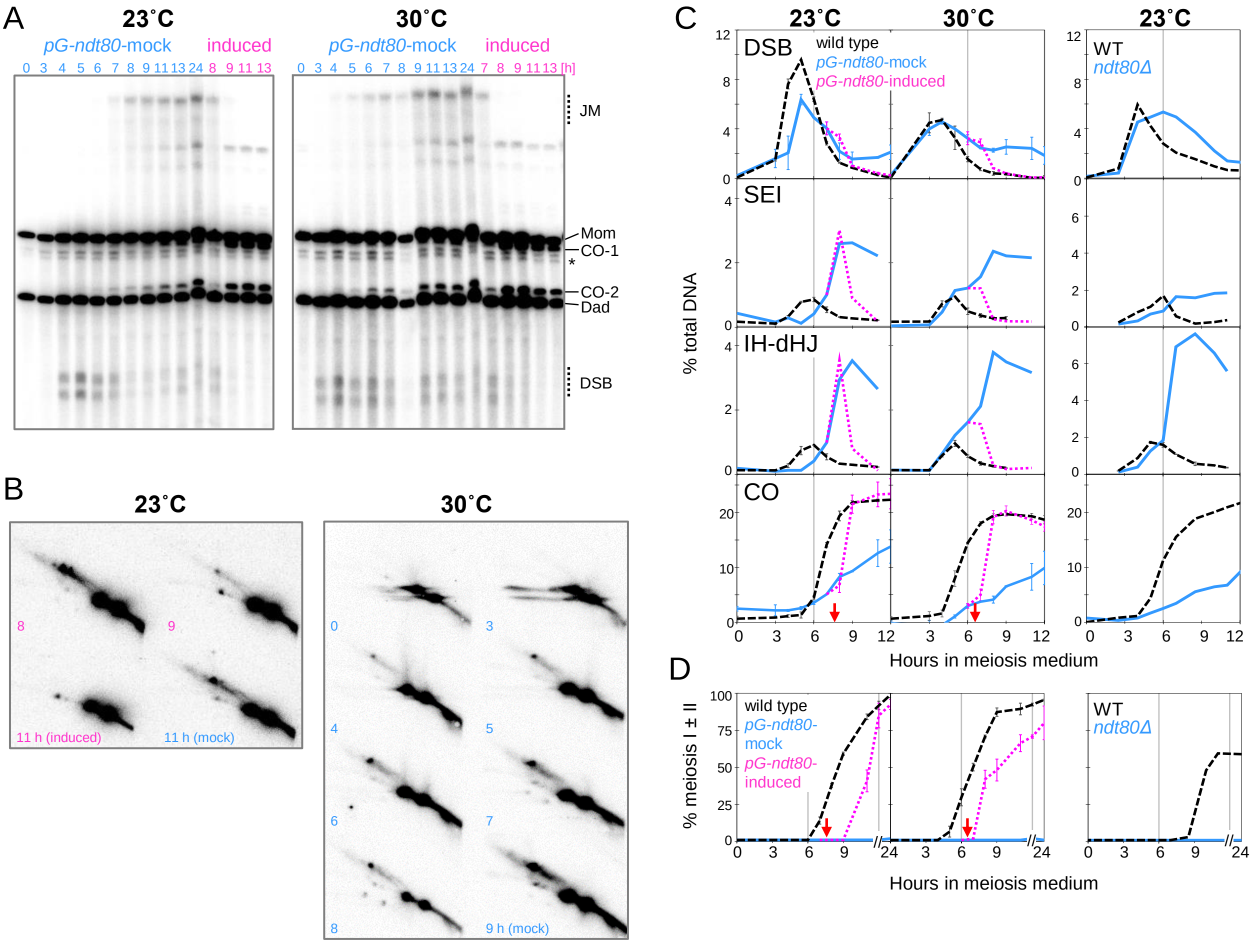

**Figure S4 Recombination and Meiotic Progression in *ndt80* Mutants at 23°C and 30°C.**

**S4A** One dimensional gel Southern blot analyses of mock and β-estradiol induced *pGAL-ndt80* cultures at 23°C (TC72-4/6) and 30°C (TC72-14,16). Asterisk: two non-specific hybridization signals. For parallel WT cultures, see Fig. 1E.

**S4B** 2D gel Southern blot analysis of recombination in induced and mock *pGAL-ndt80* cultures at 23°C (left) and of mock culture at 30°C (right). For 2D gels of parallel WT cultures, see Fig. S2A.

**S4C** Quantitative analysis of recombination intermediates (DSB, SEI, IH-dHJ) and CO products in mock and induced *pGAL-ndt80* cultures at 23°C and 30°C as well as in *ndt80Δ* at 23°C. Error bars indicate range where data from two parallel cultures were analyzed. For 1D and 2D gels of WT see Fig. 1D,E and S3A. Vertical lines indicate t = 6 h. For *pGAL-ndt80*, meiotic subcultures incubated at 23°C and 30°C are derived the same G1-arrested pre-meiotic culture. Subcultures were again split at the time of induction (7.5 h and 6.5 h, respectively, indicated by red arrows) into induced and mock-treated subcultures.

**S4D** Meiotic divisions in WT (black) and *ndt80* mutants (blue) at 23°C and 30°C, as well as induced *pGAL-ndt80* cultures (magenta). Vertical lines indicate t = 6 h and the time scale interruption between 11 h and 24 h, respectively. Red arrows indicate time of induction at 7.5 h (23°C) or at 6.5 h (30°C).

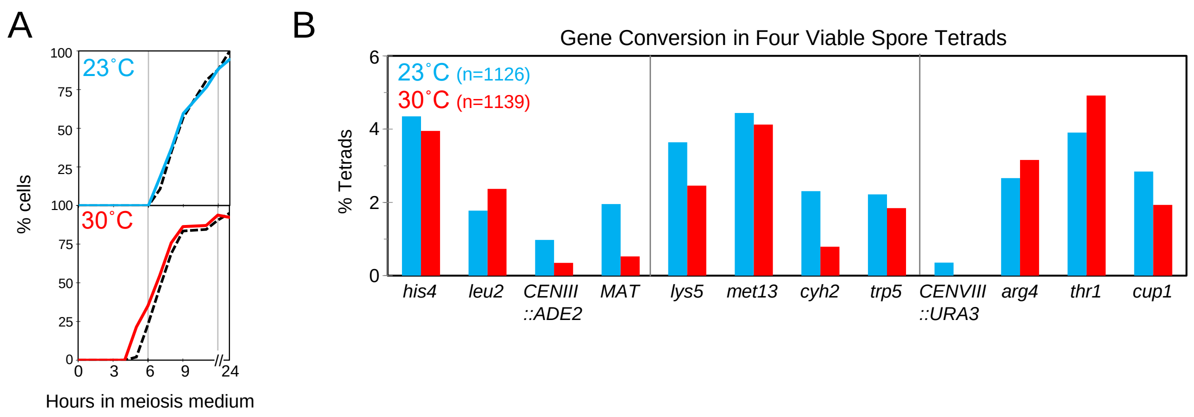

**Figure S5 Meiotic Progression and Gene Conversion at 23˚C and 30˚C.**

**S5A** Comparison of meiotic progression in strains tested for recombination in nine intervals (colored) and for Zip1 recruitment and recombination at *HIS4LEU2* (dashed lines, see Figure 1).

**S5B** Frequencies of tetrads exhibiting non-Mendelian (3:1, or 4:0) marker segregation (gene conversion) at ≤ 4 positions among four viable spore tetrads at 23˚C and 30˚C. Tetrads with gene conversion at ≥ 5 markers were presumed to result from aberrant spore association and were excluded from analysis.

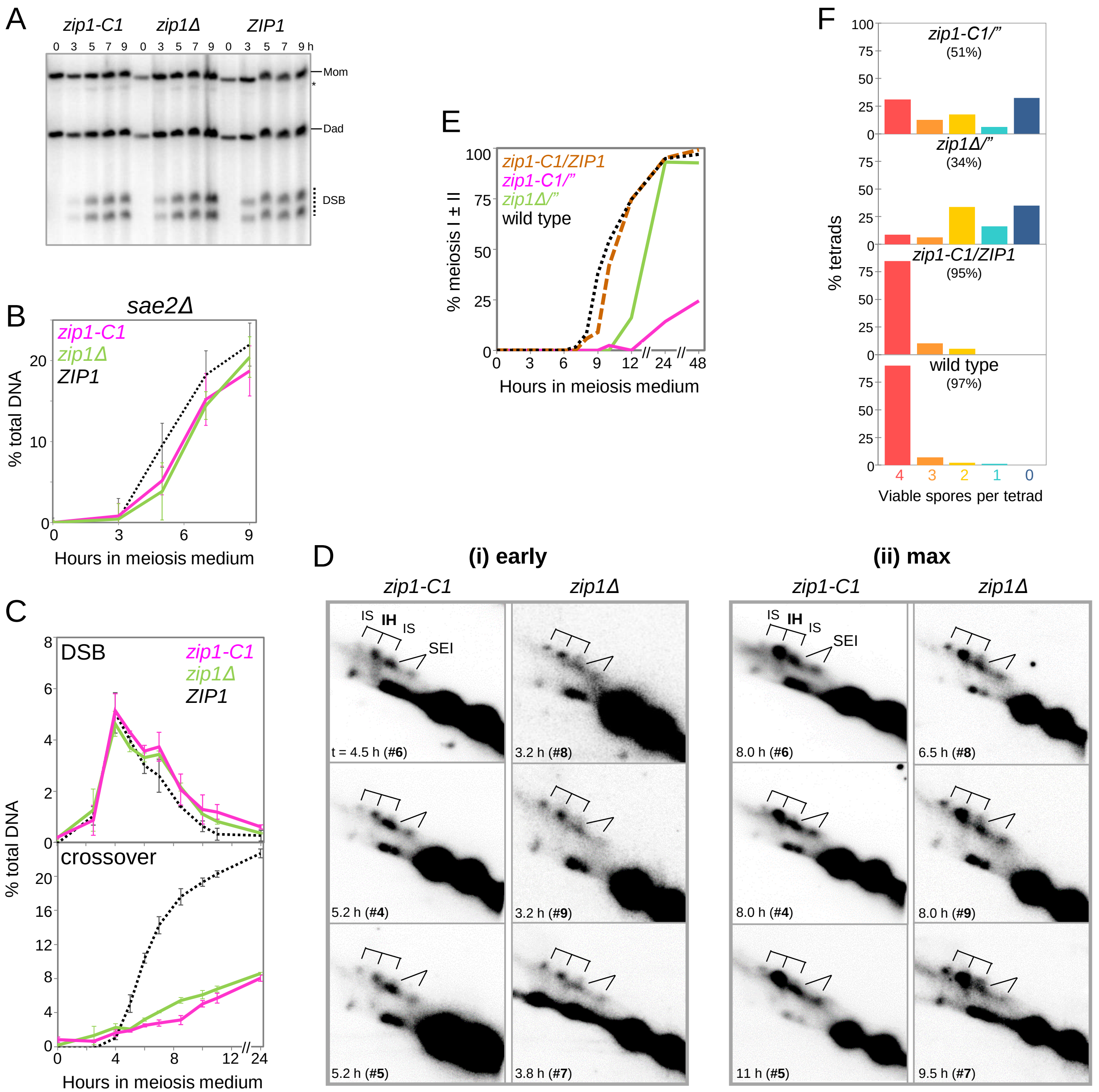

**Figure S6 Recombination in Synapsis-Defective *zip1-C1* (*Δ791-824*) at 23°C.**

**S6A** Representative Southern blot analysis of DSB formation at *HIS4LEU2* in *zip1-C1*, *zip1Δ* and WT in the *sae2Δ* background (tc16-RiS).

**S6B** Quantitative analysis of DSB accumulation in *zip1-C1*, *zip1Δ* and WT in the *sae2Δ* background. Error bars indicate SD (n = 4; tc16-RiS).

**S6C** Quantitation of DSBs and crossovers in WT, *zip1-C1* and *zip1Δ* in a second experiment (tc42-Jas). Error bars indicate ranges for two cultures per genotype.

**S6D** Excerpts from 2D gel Southern blot analysis of parallel meiotic cultures *zip1-C1* (#4, #5, #6) and *zip1Δ* (#7. #8, #9) at the earliest time of reliable detection (i) and at the time of maximum levels (ii). *zip1-C1* and *zip1Δ* cultures are matched pairwise based on the approximate timing of recombination events.

**S6E** Meiotic nuclear divisions in a strain heterozygous for *zip1-C1/ZIP1* compared to homozygous *zip1-C1* and *zip1Δ*. The time scale is interrupted after 12 h and 24 h.

**S6F** Viable spores per tetrads (tc18RS, tc21RS). Following sporulation in liquid meiosis medium, at least 80 tetrads were dissected for each genotype. Percent of spore viability is given in parentheses for each genotype.

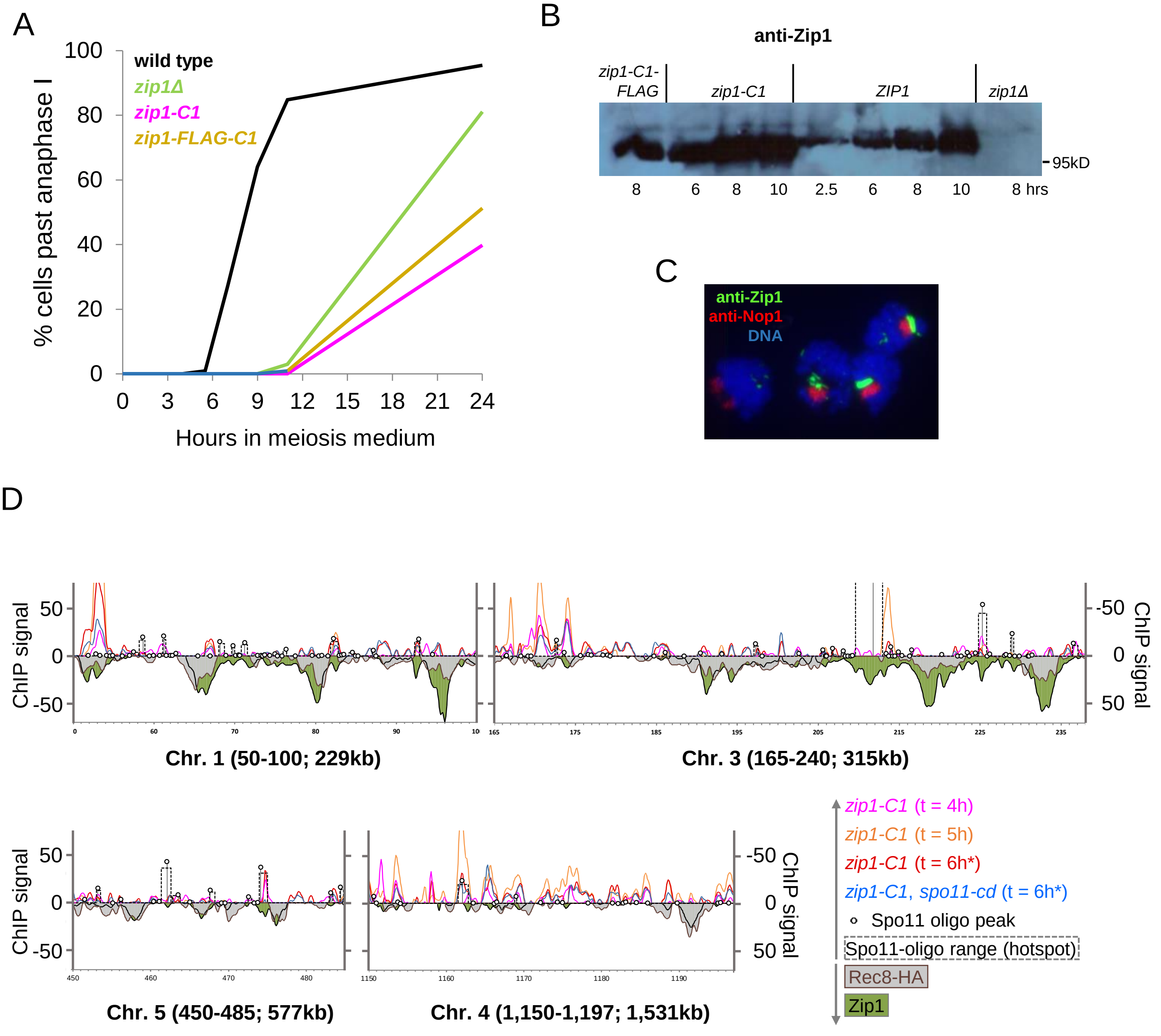

**Figure S7 Association of Zip1-C1 with Chromatin Independent of SC Assembly.**

**S7A** Cells after the first meiotic division in *ZIP1*, *zip1-C1-FLAG*, *zip1-C1* and *zip1Δ*.

**S7B** Western blot analysis of meiotic cultures at 23°C with rabbit anti-Zip1 antibodies at the indicated time points in internally tagged *zip1-C1-FLAG*, *zip1-C1, ZIP1*, and *zip1Δ*.

**S7C** Immunofluorescence detection in four surface-spread nuclei of Zip1 polycomplex (green) apposed to the nucleolus (Nop1, red) in homozygous *zip1-C1* mutants.

**S7D** Chip-on-chip enrichment profiles with anti-Zip1 antibodies at short (1, 3, top) and long chromosomes (4, 5, bottom). Timepoints reflect hours in meiosis medium. *zip1-C1* at time points t = 4 h (magenta), t = 5 h (orange) and t = 6 h (ruby), as well as in a *spo11-cd*, catalytic dead background (blue). Spo11-oligo hotspots are indicated by their local maximum (lollipop), while the dimensions of the hotspots are indicated with dashed lines. For orientation, Zip1 wild-type profiles (green fill) and Rec8 wild-type profiles are plotted to the negative scale. Both bind to core sites, but only Zip1 also localizes to strong hotspots.

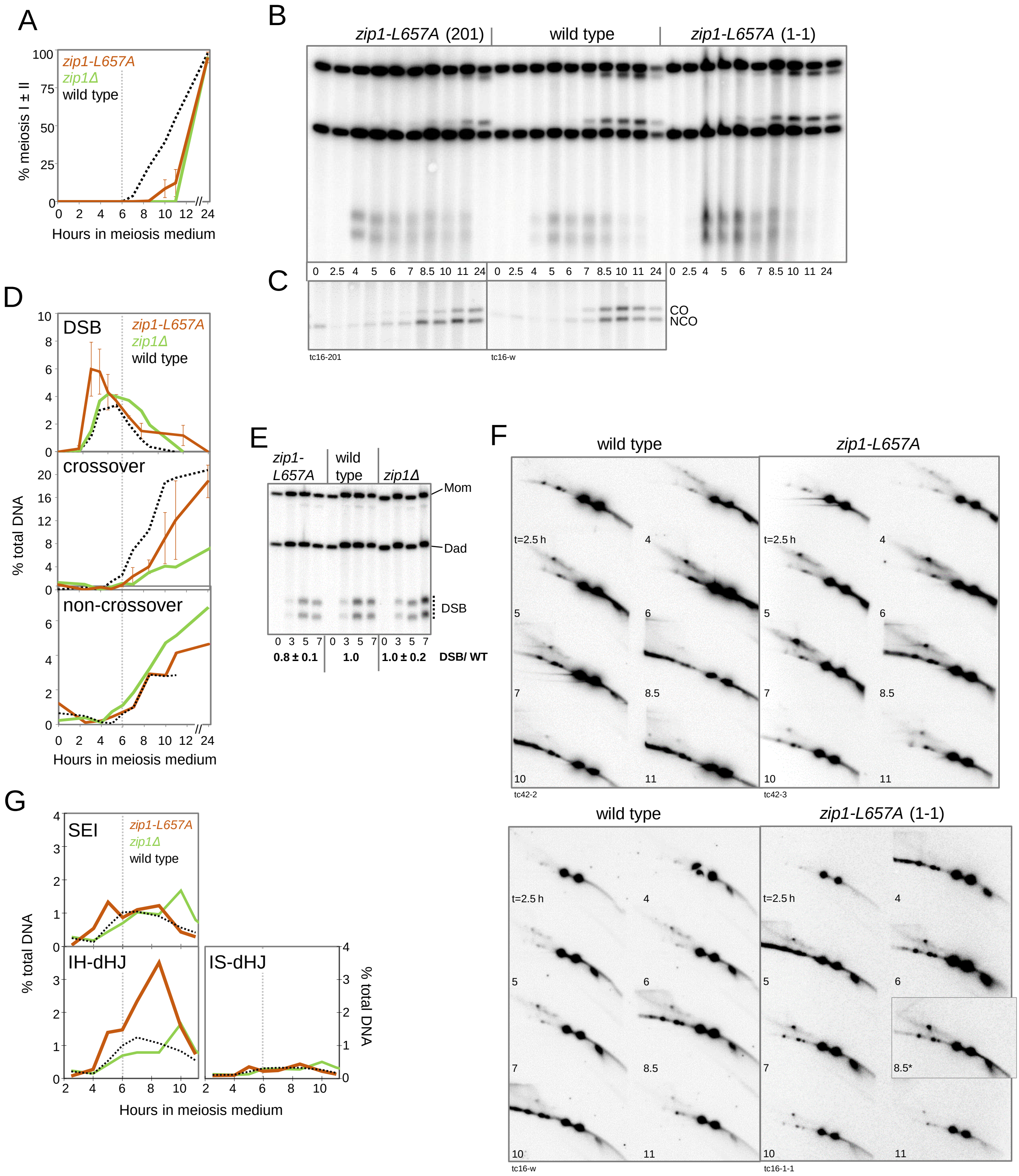

**Figure S8 Effects of the *zip1-L657A* Point Mutation on Meiotic Progression, Synapsis and Recombination.**

**S8A** Meiotic nuclear divisions in *zip1-L657A* (n = 2), *zip1Δ* and wild type. Error bars indicate range. Here and in subsequent plots, the dotted gray line indicates t = 6 h.

**S8B** One dimensional gel Southern blot analysis of DSBs and crossovers at *HIS4LEU2* in *zip1-L657A* and wild type.

**S8C** Excerpt from one-dimensional gel Southern blot showing CO-1 and NCO-1 products refractory to NgoMIV in a double digest with XhoI in the same DNA samples analyzed in Fig. 7B.

**S8D** Quantitative analysis of DSBs, crossovers and non-crossovers in *zip1-L657A*, *zip1Δ* and wild type. Error bars indicate range (n = 2) where available.

**S8E** Representative Southern blot analysis of DSB formation at *HIS4LEU2* in *zip1-C1*, *zip1Δ* and WT in the *sae2Δ* background (tc39).

**S8F** Two sets of complete 2D gel Southern blot analyses of *zip1-L657A* and WT cultures, analyzed in two experiments (tc42, tc16). Blots in the top row are quantitated in Figure 7G, the bottom row is quantitated in Figure S8G. Asterisk, the excerpt from the WT sample t = 8.5 h was cropped and inverted due to loading on the original gel in flipped orientation.

**S8G** Quantitative analysis of SEIs, IH-dHJs and IS-dHJs in parallel *zip1-L657A*, *zip1Δ,* and wild-type cultures. [RaS tc16: WT, 1-1]

### Table S1. Timing of Key Events during Meiosis under Different Incubation Conditions

| | Timing relative to transfer to meiosis medium | | | | | | | | | | | | | | |
|---|---|---|---|---|---|---|---|---|---|---|---|---|---|---|---|
| | 23°C | | | 30°C | | | 33°C | | | 30°C [TC11; Padmore] [(b)] | | | 30°C [TC20; Hunter] [(b)] | | |
| | Entry | Exit | Lifespan | Entry | Exit | Lifespan | Entry | Exit | Lifespan | Entry | Exit | Lifespan | Entry | Exit | Lifespan |
| DSB | 4.34 | 5.57 | 1.23 | 3.13 | 3.95 | 0.84 | 4.51 | 5.37 | 0.85 | 3.79 [3.80] | 4.83 [4.87] | 1.04 [1.07] | 3.80 [3.90] | 5.03 [5.20] | 1.23 [1.30] |
| SEI | 5.83 | 6.09 | 0.26 | 4.87 | 5.13 | 0.27 | 6.08 | 6.39 | 0.31 | n.d. | | | 4.43 [4.60] | 5.46 [5.35] | 1.03 [0.75] |
| IH-dHJ | 6.06 | 6.19 | 0.13 | 5.13 | 5.24 | 0.12 | 6.73 | 6.96 | 0.24 | | | | 5.34 [5.20] | 5.75 [5.70] | 0.40 [0.50] |
| crossovers | 6.71 | | | 5.36 | | | 7.01 | | | 7.11 | | | 5.52 [5.70] | | |
| total DSBs (*sae2Δ* or *rad50S*) | | | 24 ± 0.8% (n=6) | | | 20 ± 2.4% (n = 3) [(e)] | | | 13 ± 0.2% (n = 2) | | | 15 ± 7.5% | | | 23.3% |
| total crossovers | | | 22 ± 0.1% | | | 20 ± 0.3% | | | 14.6 ± 2.0% [(a)] | | | 14% [(c)] | | | 23.6% |
| Zip1+, leptonema (no lines, early) or "preSC-A" | 1.66 | 3.15 | 1.48 | 1.63 | 3.00 | 1.37 | 2.81 | 4.46 | 1.65 | 3.94 [4.00] | 4.38 [4.40] | 0.44 [0.40] | n.d. | | |
| Zip1+, zygonema (few lines, early) or "preSC-B" | 3.15 | 4.98 | 1.83 | 3.00 | 4.09 | 1.09 | 4.46 | 5.59 | 1.13 | 4.38 [4.40] | 5.21 [5.30] | 0.83 [0.90] | | | |
| Zip1+, early pachynema (discontinuoes lines, early) | 4.98 | 5.69 | 0.71 | 4.09 | 4.75 | 0.66 | 5.59 | 5.92 | 0.33 | | | | | | |
| Zip1+, mid pachynema (continous lines) | 5.69 | 5.88 | 0.19 | 4.75 | 5.09 | 0.34 | 5.92 | 6.56 | 0.64 | 5.21 [5.30] | 6.67 [6.60] | 1.46 [1.30] | | | |
| Zip1+, late pachynema (discontinuous)* | 5.88 | 6.02 | 0.14 | 5.09 | 5.10 | 0.01 | 6.56 | 6.88 | 0.33 | | | | | | |
| Zip1+, diplonema (few or no lines) | 6.02 | 7.29 | 1.28 | 5.10 | 6.02 | 0.93 | 6.88 | 7.75 | 0.86 | 6.67 [n.a.] | 6.79 [n.a.] | 0.12 [n.a.] | | | |
| Meiosis I divisions | 8.57 | | | 6.92 | | | 9.34 | | | 7.87 | | | 6.80 | | |
| total duration of Zip1+/electrodense structures | | | 5.63 | | | 4.39 | | | 4.94 | | | 2.85 [2.60] | | | n.d. |

| | Timing relative to crossovers | | | | | | | | | |
|---|---|---|---|---|---|---|---|---|---|---|
| | 23°C | | 30°C | | 33°C | | Padmore | | Hunter | |
| | Entry | Exit | Entry | Exit | Entry | Exit | Entry | Exit | Entry | Exit |
| DSB | -2.36 | -1.13 | -2.23 | -1.42 | -2.50 | -1.64 | -3.32 | -2.28 | -1.72 | -0.49 |
| SEI | -0.87 | -0.61 | -0.49 | -0.23 | -0.93 | -0.62 | n.d. | | -1.09 | -0.06 |
| IH-dHJ | -0.64 | -0.51 | -0.24 | -0.12 | -0.28 | -0.05 | | | -0.18 | 0.23 |
| crossovers | 0.00 | | 0.00 | | 0.00 | | 0.00 | | 0.00 | |
| total DSBs (*sae2Δ* or *rad50S*) | | | | | | | | | | |
| total crossovers | | | | | | | | | | |
| Zip1+, leptonema (no lines, early) or "preSC-A" | -5.04 | -3.56 | -3.73 | -2.37 | -4.20 | -2.55 | -3.17 | -2.73 | n.d. | |
| Zip1+, zygonema (few lines, early) or "preSC-B" | -3.56 | -1.73 | -2.37 | -1.28 | -2.55 | -1.42 | -2.73 | -1.90 | | |
| Zip1+, early pachynema (discontinuoes lines, early) | -1.73 | -1.02 | -1.28 | -0.61 | -1.42 | -1.09 | | | | |
| Zip1+, mid pachynema (continous lines) | -1.02 | -0.83 | -0.61 | -0.27 | -1.09 | -0.45 | -1.90 | -0.44 | | |
| Zip1+, late pachynema (discontinuous)* | -0.83 | -0.69 | n.a. | | -0.45 | -0.13 | | | | |
| Zip1+, diplonema (few or no lines) | -0.69 | 0.59 | -0.27 | 0.66 | n.a. | | -0.44 | -0.32 | | |
| Meiosis I divisions | 1.86 | | 1.55 | | 2.33 | | 0.76 | | 1.28 | |
| total duration of Zip1+/electrodense structures | | | | | | | | | | |

* underestimate

Notes:

(a) Under the current conditions for meiosis at 33°C which entails incubation at 26°C until t = 3h, crossovers reach maximum levels of 14.6 (± 2.0)% SD (n = 4) by t = 8.5 h (see Supplemental Materials, Ahuja et al., 2017), a maximum somewhat lower than the 17.3 (± 2.2)% SD (n = 8) measured after incubation at 30°C for the first two hours followed by a shift to 33°C (Joshi et al., 2015). 15% was used as a conservative estimate for maximum crossover levels. The 50% entry time for crossovers at 33°C would be somewhat later if higher maximum crossover levels was assumed.

*(b)* Values in parenthesis indicate the entry and exit times as provided by Padmore et al. (1991) and Hunter & Kleckner (2001), respectively, which were determined by somewhat different cumulative analyses.

*(c)* 15% were assumed as maximum crossovers, as the graph in Figure 7b (Padmore et al., 1991) indicates that 92% of crossovers were detected, which implies that the maximum crossover number was 15%. Independent of the absolute crossover levels, 50% of the maximum crossover levels (92%) in tc11 is 46% which is reached at t = 7.0 h.

*(d)* All Zip1 entry and exit times were determined using stage-specific, modified cumulative analysis, as described in Experimental Procedures.

*(e)* Data are from Martini et al. (2006).

### Table S2. Images of Nuclei Scored for Zip1 Morphology

| experiment | tc72 | tc72 | tc72 | tc72 | tc72 | tc72 | tc72 | tc72 | | tc55 | | tc42 | tc42 | tc42 | | tc15 | tc15 |
|---|---|---|---|---|---|---|---|---|---|---|---|---|---|---|---|---|---|
| culture | 1 | 3 | 11 | 13 | 4 | 6 | 14 | 16 | | 1 | | 2 | 3 | 5 | | w | c |
| culture in Fig. | | | | | | | | | | | | | | | | | |
| genotype | WT | WT | WT | WT | pG-ndt80 | pG-ndt80 | pG-ndt80 | pG-ndt80 | | WT | | WT | zip1-201 | ndt80D | | WT | zip1-C1 |
| temperature | 23°C | 23°C | 30°C | 30°C | 23°C | 23°C | 30°C | 30°C | | 33°C | | 23°C | 23°C | 23°C | | 23°C | 23°C |
| induction (β-ED) | | | | | | t = 7.5h | | t = 6.5h | | | | | | | | | |
| time points | | | | | | | | | time points | | time points | | | | time points | | |
| 0 | 125 | 100 | see tc72-1 | see tc72-3 | 102 | | see tc72-4 | | 0 | 52 | 0 | 100 | 100 | 100 | 0 | 100 | 100 |
| 3 | 105 | 124 | 118 | 129 | 137 | | 64 | | 3 | 84 | 5 | 242 | 201 | 201 | 4 | 50 | 101 |
| 4 | 164 | 120 | 195 | 65 | 295 | | 206 | | 4 | 127 | 6 | 104 | 102 | 105 | 5 | 99 | 104 |
| 5 | 98 | 177 | 159 | 81 | 151 | | 276 | | 5 | 58 | 7 | 133 | 204 | 222 | 7 | 109 | 102 |
| 6 | 238 | 256 | 154 | 96 | 156 | | 212 | | 6 | 64 | 8.5 | 212 | 202 | 204 | 8 | 100 | 100 |
| 7 | 88 | 245 | 97 | 106 | 217 | | 160 | 167 | 7 | 82 | | | | | 9 | 104 | 104 |
| 8 | 144 | 176 | 154 | 100 | 142 | 144 | 219 | 171 | 8.5 | 144 | | | | | | | |
| 9 | 34 | 100 | | | 115 | 229 | | | | | | | | | | | |
| **total nuclei** | **996** | **1,298** | **877** | **577** | **1,315** | **373** | **1,137** | **338** | **total nuclei** | **611** | **total nuclei** | **791** | **809** | **832** | **total nuclei** | **562** | **611** |
| | | | | | | | | | | | | | | | | | **11,127** |

### Table S3. Crossover interference along three chromosomes in wild type at 23˚C and 30˚C

#### A. Chromosome III

| Genotype; Temperature | Reference Interval / Test Interval | | *his4-leu2* (1): *leu2-CEN3* (2) | *his4-leu2* (1): *CEN3-MAT* (3) | *leu2-CEN3* (2): *his4-leu2* (1) | *leu2-CEN3* (2): *CEN3-MAT* (3) | *CEN3-MAT* (3): *his4-leu2* (1) | *CEN3-MAT* (3): *leu2-CEN3* (2) |
|---|---|---|---|---|---|---|---|---|
| Wild type; 23˚C | P | P:N:T | 576:1:172 | 464:5:268 | 576:4:287 | 564:4:312 | 464:2:216 | 564:1:143 |
| | | cM ± SE | 11.9 ± 0.9 | 20.2 ± 1.2 | 17.9 ± 1.0 | 19.1 ± 1.0 | 16.7 ± 1.1 | 10.5 ± .08 |
| | N | P:N:T | 291:0:24 | 218:2:92 | 173:0:24 | 144:4:60 | 273:2:92 | 316:0:64 |
| | | cM ± SE | 3.8 ± 0.7 | 16.7 ± 1.8 | 6.1 ± 1.2 | 20.2 ± 3.1 | 14.2 ± 1.6 | 8.4 ± .10 |
| | **ratio N/P** | | **.32 ± .07** | **.82 ± .10** | **.34 ± .07** | **1.06 ± .17** | **.85 ± .11** | **.80 ± .11** |
| | **sig (SE) †** | | **S** | **N** | **S** | **N** | **N** | **N** |
| Wild type; 30˚C | P | P:N:T | 517:1:188 | 491:3:208 | 517:3:336 | 607:3:280 | 491:4:237 | 607:1:154 |
| | | cM ± SE | 13.7 ± 0.9 | 16.1 ± 1.1 | 20.7 ± 1.0 | 16.7 ± 1.0 | 17.8 ± 1.2 | 10.5 ± 0.8 |
| | N | P:N:T | 339:0:19 | 241:0:116 | 189:1:18 | 155:0:57 | 211:0:116 | 283:0:57 |
| | | cM ± SE | 2.7 ± 0.6 | 16.3 ± 1.2 | 5.8 ± 1.7 | 13.4 ± 1.5 | 17.7 ± 1.3 | 8.4 ± 1.0 |
| | **ratio N/P** | | **.19 ± .05** | **1.01 ± .10** | **.28 ± .08** | **0.80 ± .10** | **.99 ± .10** | **.80 ± .11** |
| | **sig (SE) †** | | **S** | **N** | **S** | **N** | **N** | **N** |
| | **sig (ratios 23˚C vs 30˚C) ‡** | | **N** | **N** | **N** | **N** | **N** | **N** |

Table S3: Crossover interference for tetrads was analyzed by coincidence analysis as described by Malkova et al. (2004). For each test interval, tetrads were divided into two groups based on the crossover status of the reference interval, carrying parental (P) or non-parental (N) marker configurations, respectively. Map distances for each subset of tetrads were determined in centiMorgan (cM ± standard error) as described in Materials and Methods. The "ratio" of the two map distances indicates the strength of interference between the reference and the test interval: The lower the ratio, the stronger the interference. Standard errors and the significance of differences between ratios at 23˚C and 30˚C were calculated using the application "Analysis of Statistical Significance of Differences Between Two Ratios of Map Distances" from the Stahl website (see Materials and Methods).

† Significance of difference in ratios between reference and test intervals. If the absolute value of the difference between the two map distances is greater than twice the standard error, the difference is considered as significant.

‡ Significances of differences between interference ratios at 23˚C versus 30˚C as calculated by comparing standard errors via a two-tailed test.

### B. Chromosome VII

| Genotype; Temperature | Reference Interval / Test Interval | | *lys5-met13* (4) | | *met13-cyh2* (5) | | *cyh2-trp5* (6) | |
|---|---|---|---|---|---|---|---|---|
| | | | *met13-cyh2* (5) | *cyh2-trp5* (6) | *lys5-met13* (4) | *cyh2-trp5* (6) | *lys5-met13* (4) | *met13-cyh2* (5) |
| Wild type; 23˚C | P | P:N:T | 485:1:186 | 205:19:437 | 485:5:323 | 248:23:546 | 205:0:140 | 248:0:106 |
| | | cM ± SE | 14.3 ± 1.0 | 41.7 ± 1.9 | 21.7 ± 1.2 | 41.9 ± 1.7 | 20.3 ± 1.3 | 15.0 ± 1.2 |
| | N | P:N:T | 328:0:39 | 140:8:210 | 187:0:39 | 106:4:114 | 456:4:214 | 569:1:117 |
| | | cM ± SE | 5.3 ± 0.8 | 36.0 ± 2.5 | 8.6 ± 1.3 | 30.8 ± 2.9 | 17.7 ± 1.2 | 9.0 ± 0.8 |
| | **ratio N/P** | | **.37 ± .06** | **.86 ± .07** | **.40 ± .06** | **.74 ± .08** | **.87 ± .08** | **.60 ± .12** |
| | **sig (SE) †** | | **S** | **N** | **S** | **S** | **N** | **S** |
| Wild type; 30˚C | P | P:N:T | 481:1:176 | 205:13:426 | 481:3:370 | 256:19:583 | 205:1:146 | 256:0:105 |
| | | cM ± SE | 13.8 ± 1.0 | 39.1 ± 1.7 | 22.7 ± 1.0 | 40.6 ± 1.5 | 21.6 ± 1.5 | 14.5 ± 1.2 |
| | N | P:N:T | 373:0:30 | 147:5:244 | 177:0:30 | 105:1:102 | 439:2:247 | 602:1:102 |
| | | cM ± SE | 3.7 ± 0.7 | 34.6 ± 1.9 | 7.3 ± 1.2 | 26.0 ± 2.2 | 18.8 ± 1.1 | 7.7 ± 0.8 |
| | **ratio N/P** | | **.27 ± .05** | **0.88 ± .06** | **.32 ± .13** | **.64 ± .06** | **.87 ± .08** | **.53 ± .07** |
| | **sig (SE) †** | | **S** | **N** | **S** | **S** | **N** | **S** |
| | **sig (ratios 23˚C vs 30˚C)‡** | | **N** | **N** | **N** | **N** | **N** | **N** |

**C. Chromosome VIII**

| Genotype; Temperature | Reference Interval / Test Interval | | *CEN8-arg4* (7) | | *arg4-thr1* (8) | | *thr1-cup1* (9) | |
|---|---|---|---|---|---|---|---|---|
| | | | *arg4-thr1* (8) | *thr1-cup1* (9) | *CEN8-arg4* (7) | *thr1-cup1* (9) | *CEN8-arg4* (7) | *arg4-thr1* (8) |
| Wild type; 23˚C | P | P:N:T | 685:0:154 | 401:6:413 | 685:1:204 | 423:5:443 | 401:1:138 | 423:0:118 |
| | | cM ± SE | 9.2 ± 0.7 | 27.4 ± 1.2 | 11.8 ± 0.8 | 27.2 ± 1.1 | 13.3 ± 1.1 | 10.9 ± 0.9 |
| | N | P:N:T | 205:1:9 | 139:0:69 | 154:2:8 | 118:1:40 | 419:2:67 | 448:1:40 |
| | | cM ± SE | 3.5 ± 1.5 | 16.6 ± 1.6 | 6.1 ± 2.7 | 14.5 ± 2.5 | 8.1 ± 1.2 | 4.7 ± 0.9 |
| | **ratio N/P** | | **.38 ± .17** | **.61 ± .07** | **.57 ± .23** | **.53 ± .09** | **.61 ± .10** | **.43 ± .09** |
| | **sig (SE) †** | | **S** | **S** | **S** | **S** | **S** | **S** |
| Wild type; 30˚C | P | P:N:T | 690:0:149 | 350:8:465 | 690:0:205 | 380:8:494 | 350:0:149 | 380:0:119 |
| | | cM ± SE | 8.9 ± 0.7 | 31.2 ± 1.3 | 11.5 ± 0.7 | 30.7 ± 1.2 | 14.9 ± 1.0 | 11.9 ± 1.0 |
| | N | P:N:T | 205:0:8 | 149:0:61 | 149:0:8 | 119:0:32 | 473:0:61 | 502:0:32 |
| | | cM ± SE | 1.9 ± 0.7 | 14.5 ± 1.6 | 2.6 ± 0.9 | 10.6 ± 1.7 | 5.7 ± 0.7 | 3.0 ± 0.5 |
| | **ratio N/P** | | **.21 ± .08** | **.47 ± .05** | **.22 ± .08** | **.35 ± .06** | **.38 ± .05** | **.25 ± .05** |
| | **sig (SE) †** | | **S** | **S** | **S** | **S** | **S** | **S** |
| | **sig (ratios 23˚C vs 30˚C)‡** | | **N** | **N** | **N** | **N** | **S** | **N** |

### Table S4. Strains Used in this Study

| Name | Genotype | Reference |
|---|---|---|
| VBY 1083 | *ho::hisG/", ura3(Δsma-pst:hisG)/ura3-PM, leu2::hisG)/", his4X::LEU2-(NgoMIV;ori)-URA3/ HIS4::LEU2-(NBam;ori)* | this study |
| VBY 1886 | as VBY1083, but *ZIP3 /zip3-GFP KanMX , rec8-HA KanMX4 /REC8 , pdr5Δ ::HphMX4* / " | Ahuja et al., 2017 |
| RSY 152 | as VBY 1083, but *zip1-C1/"* | this study |
| RSY 154 | as VBY 1083, but *zip1Δ: KanMX4/"* | Joshi et al., 2015 |
| RSY 203 | as VBY 1083, but *zip1-C1/"* ; *sae2Δ* /" | this study |
| RSY 204 | as VBY 1083, but *zip1Δ/"* ; *sae2Δ* /" | this study |
| RSY 257 | as VBY 1083, but *URA3 -zip1:: zip1-(L657A)* /" | this study |
| JSY 451 | as VBY 1083, but *sae2Δ* /" | Joshi et al., 2015 |
| JSY 506 | as VBY 1083, but *zip1-(L657A)::URA3* /"; *sae2Δ* /" | this study |
| JSY 651 | as VBY 1083, but *ndt80Δ* /" | this study |
| JSY 1302 | as VBY 1083, but *pGAL1-ndt80 ::TRP1 ; GAL(848)ER-URA3* | this study |
| JSY 524 | *zip1-C1* /", *promURA::tetR:GFP::LEU2/" ura3::tetx224(336?)-HIS3/"* (ChIP) | this study |
| JSY 525 | *zip1-C1* /" *spo11-HA-Y129F* /" *promURA::tetR:GFP::LEU2/" ura3::tetx224(336?)-HIS3/"* ( ChIP) | this study |
| JSY 526 | *ZIP1 /ZIP1 , promURA::tetR:GFP::LEU2/" ura3::tetx224(336?)-HIS3/"* (ChIP) | this study |
| RY 31 | *ZIP1 /URA3-zip1::zip1-FLAG@705-C1* /"; *promURA ::tetR:GFP ::LEU2* /" *ura3::* tetx224(336?)-*HIS3* /" | this study |
| RY 124 | *URA3-zip1 ::zip1-FLAG@705-C1* /"; *promURA ::tetR:GFP ::LEU2* /" *ura3* ::tetx224(336?)-*HIS3* /" | this study |
| NHY 942 | *MAT* α, *ho ::hisG , ade2Δ , can1R , ura3(ΔSma-Pst) , met13-B , trp5-S , CEN8 ::URA3 , thr1-A , cup1S* | Martini et al., 2006 |
| NHY 943 | *MAT* a, *ho ::hisG , ade2Δ , his4-B , leu2 ::hisG , CEN3 ::ADE2 , lys5-P , ura3(ΔSma-Pst) , cyh2R* | Martini et al., 2006 |

#### PDF S1. Legend for pages 2 to 72

Image filenames are in the format –

**{TC}_{Culture}_{time-point}{image-id}_n_{nucleus_index_on_image}_{channel}**

**TC** – time-course/experiment ID: e.g. tc72

**Culture** – Sporulation culture ID (also see Table S2):
1 & 3 (WT 23ºC)
11 & 13 (WT 30ºC)
4 (*pGAL-ndt80* uninduced 23ºC)
14 (*pGAL-ndt80* uninduced 30ºC)
6 (*pGAL-ndt80* culture 4, 23ºC, induced t = 7.5 h)
16 (*pGAL-ndt80* culture 6, 30ºC, induced t = 6.5 h)

**time-point** – time in hours in sporulation medium
(see color coding below)

The font color of the image-name corresponds to the time point as follows–

0h is "white",
3h is "turquoise",
4h is "orange",
5h is "lavender",
6h is "green",
7h is "pink",
8h is "yellow",
9h is "red"

**im**[illegible]int

**nucleus_index_on_image** – identifies the n^th^ nucleus in the given image

**channel** – green = GFP

**<u>Image classes refer to presence of Zip1 lines:</u>**

continuous lines: Pages 2-4
discontinuous lines: Pages5-11
few lines: Pages 12-29
no lines (none): Pages 30-47
no Zip1 staining: Pages 48-63
PC + continuous lines: Pages 64
PC + discontinuous lines: Pages 65-66
PC + few lines: Pages 67-68
PC + no lines (none): Pages 69-70
PC + no chromatin staining: Page 71

**<u>.tiff conversion to .jpeg in ImageJ</u>**

```
function contr_save(input, output_img, filename) {
	//open image
	open(input + filename);
	//use lookup table for "Grays"
	run("Grays");
	// for contrast set minimum to 50 and maximum to 650
	setMinAndMax(50, 650);
	// save image as .jpeg
	saveAs("Jpeg", output_img + filename);
	close();

}
```

**<u>.jpeg files were compiled into page-wise collages and assembled into pdf R</u>**
We made a custom script to copy .jpeg images into multiple one-page collages using R package "magick" package for opening images and creating collages. These collages were compiled into pdf files using the "pdf" function from "grDevices".

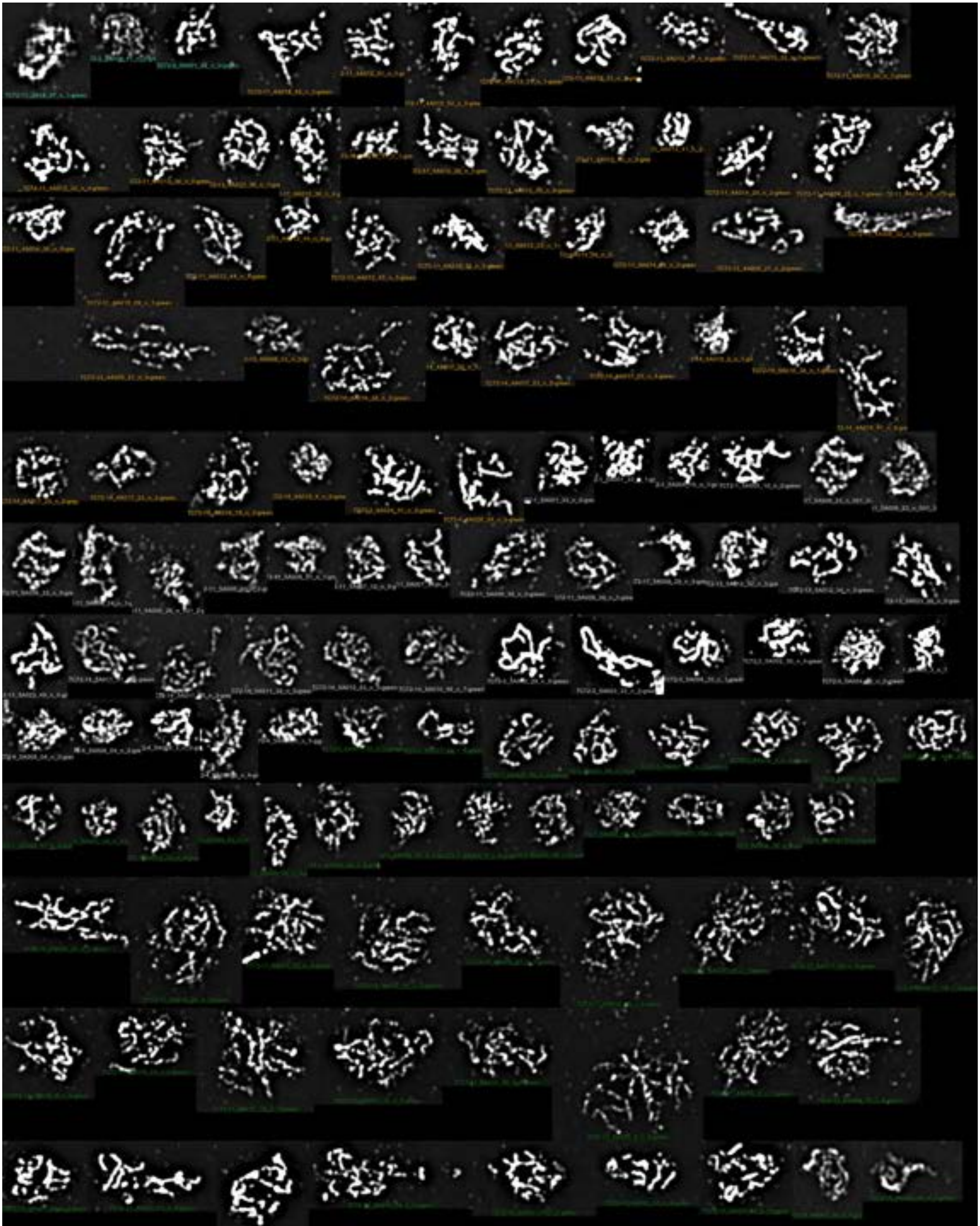

CONTINUOUS-1

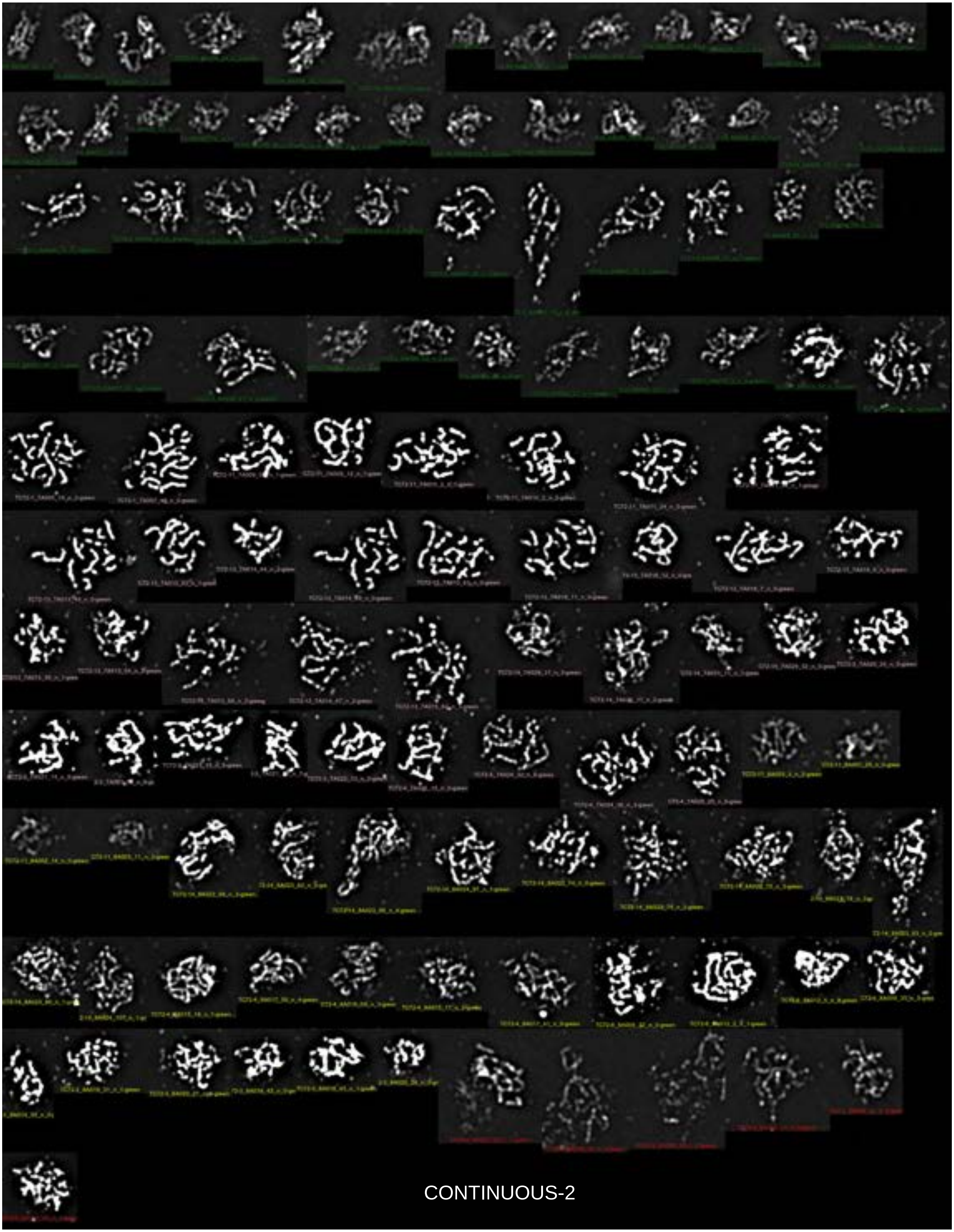

CONTINUOUS-2

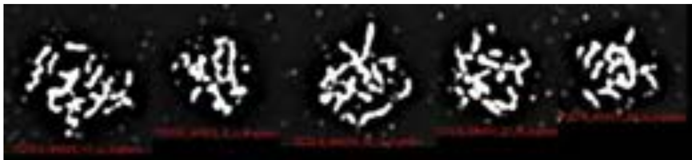

CONTINUOUS-3

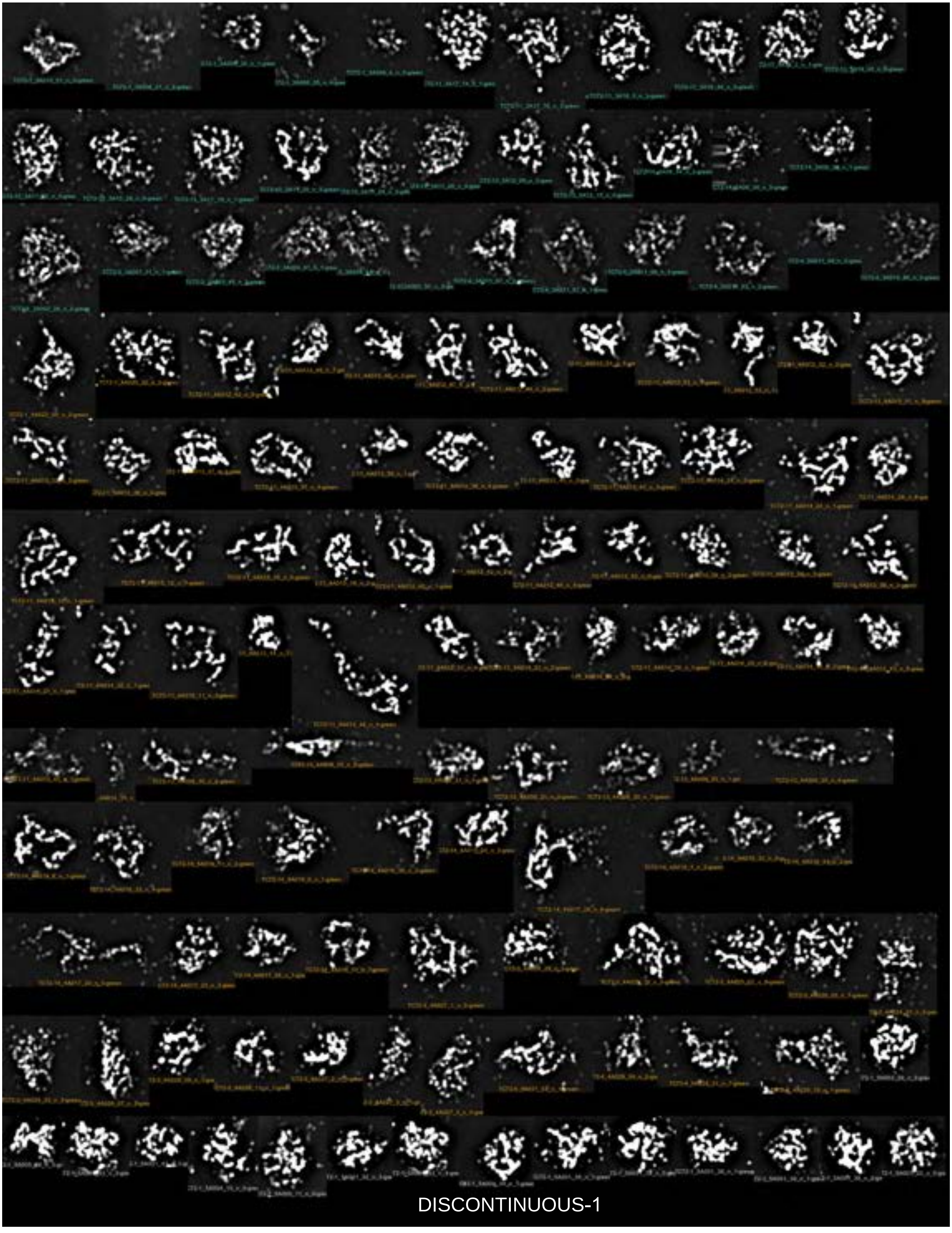

DISCONTINUOUS-1

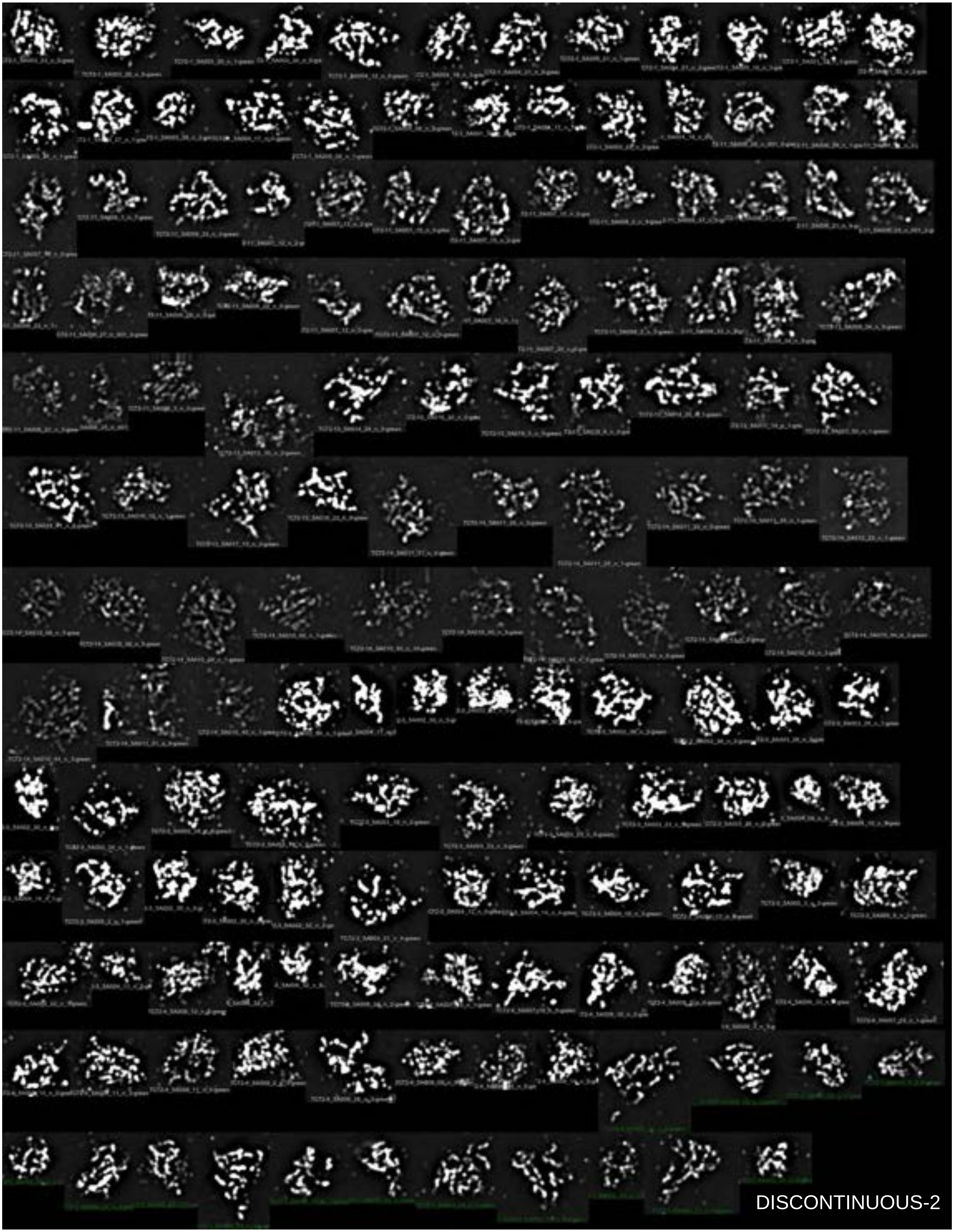
DISCONTINUOUS-2

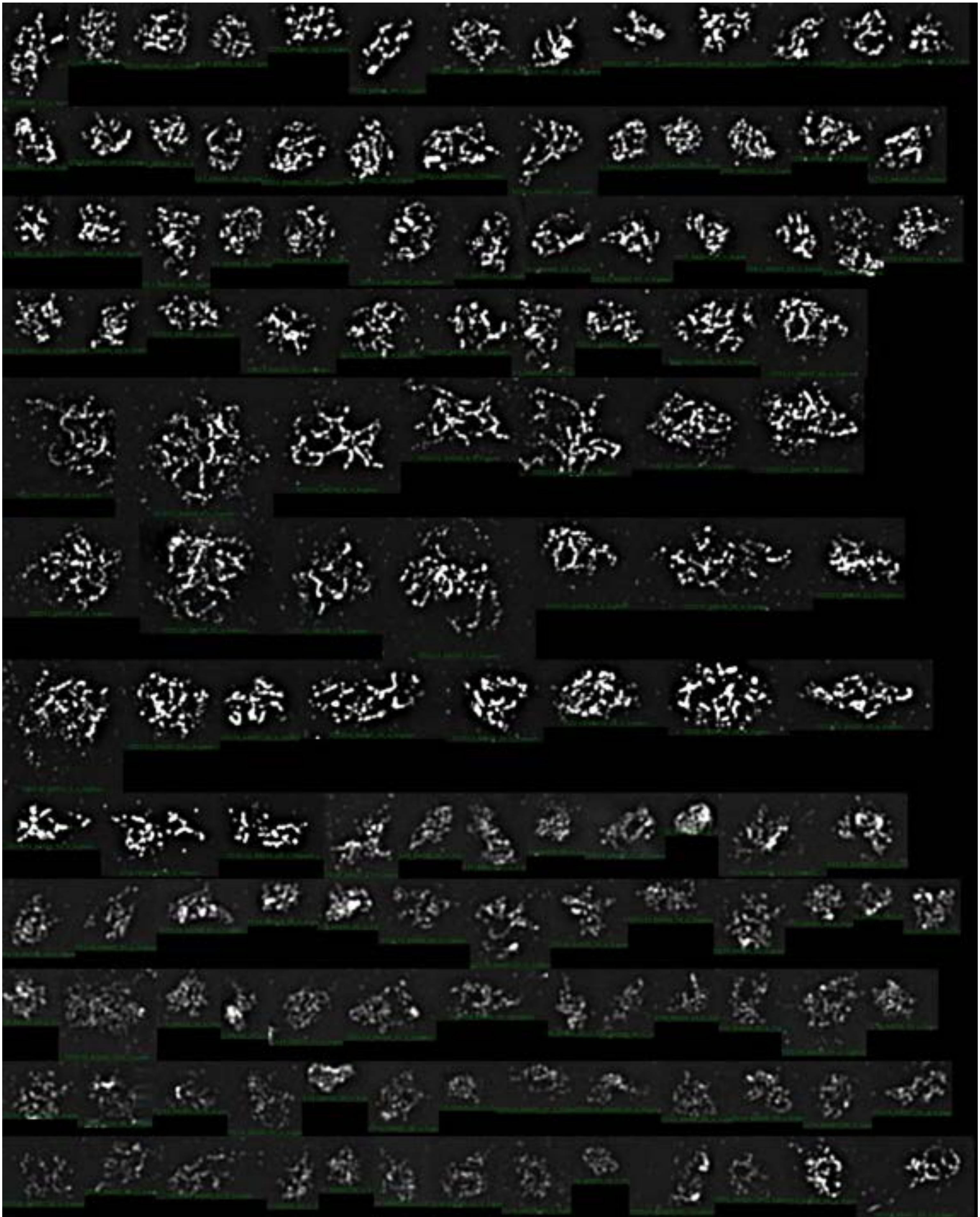

DISCONTINUOUS-3

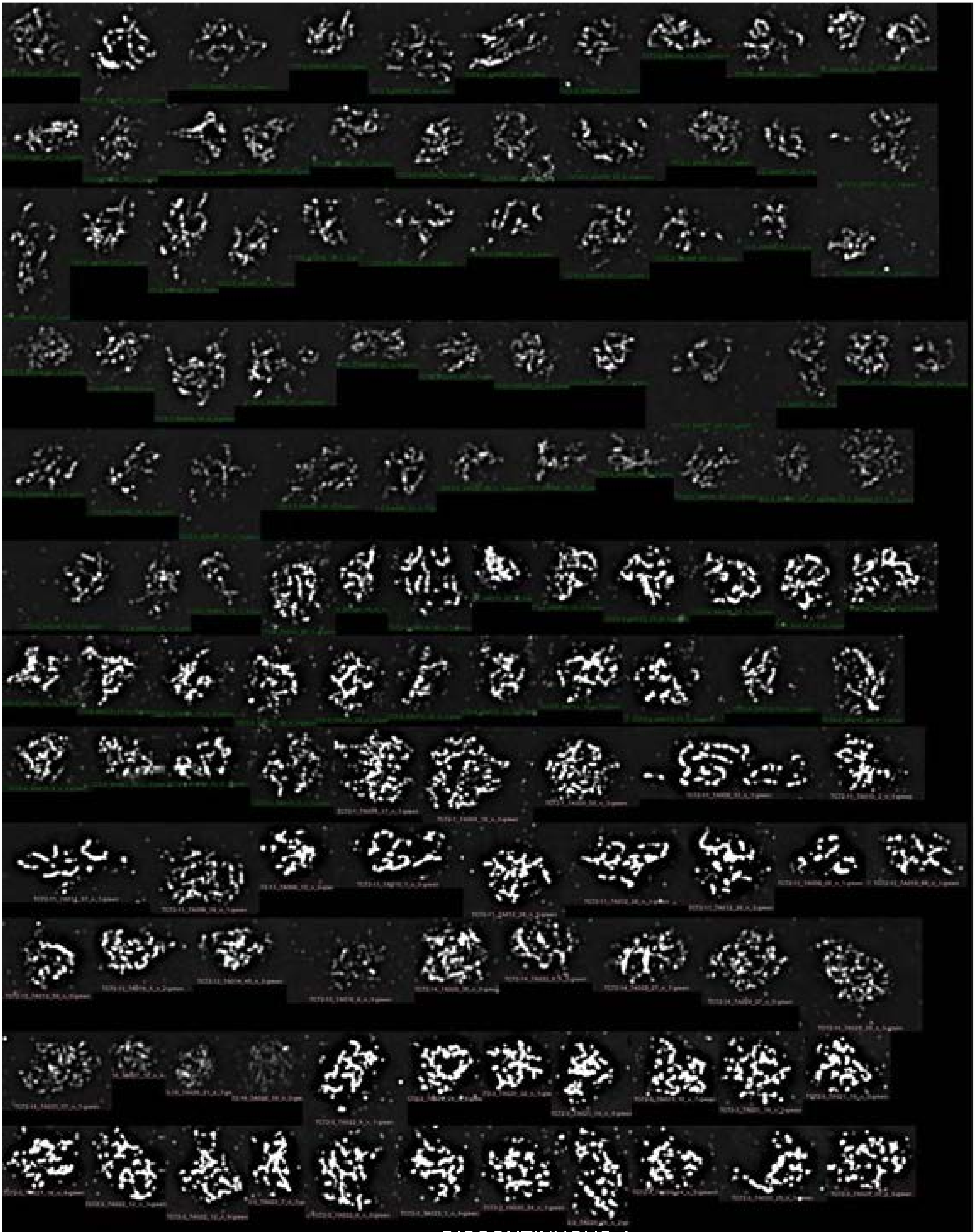

DISCONTINUOUS-4

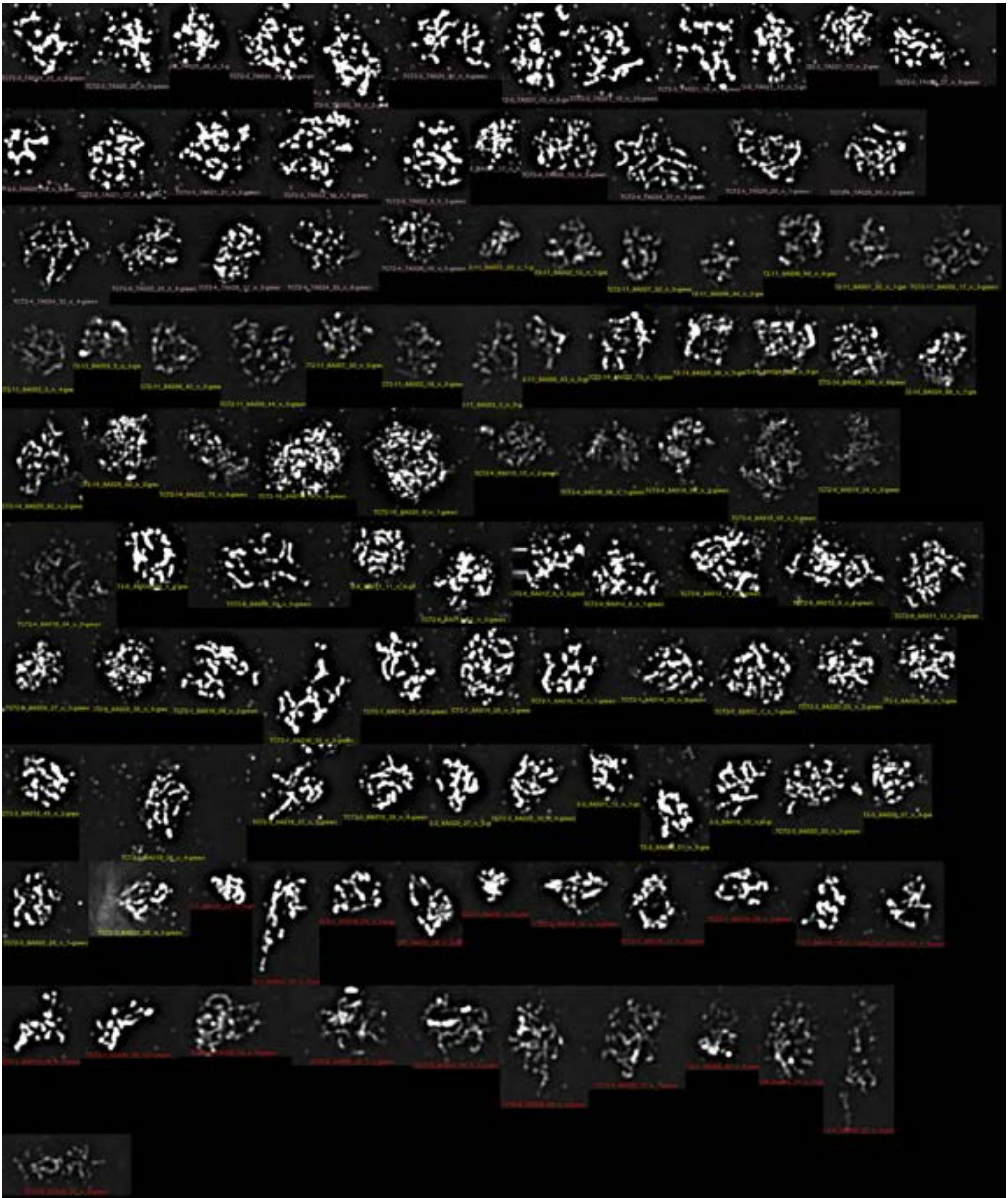

DISCONTINUOUS-5

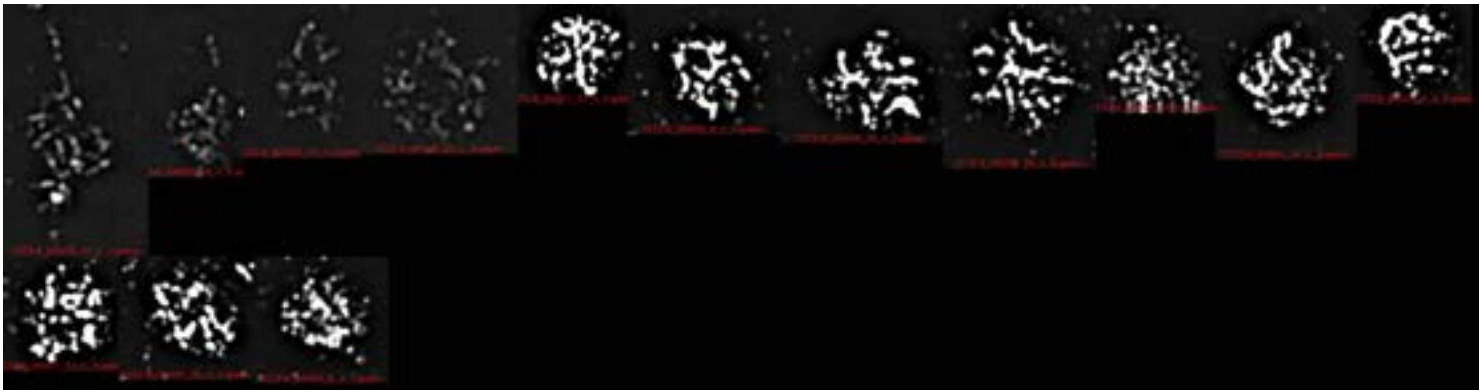

DISCONTINUOUS-6

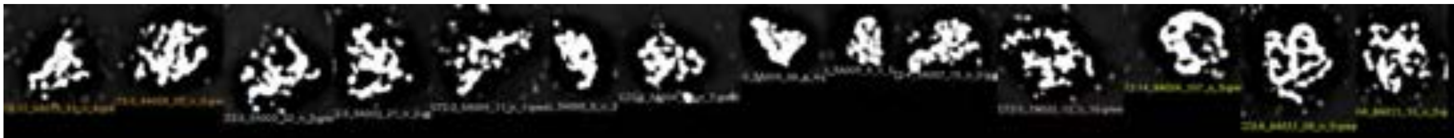

DISCONTINUOUS-7

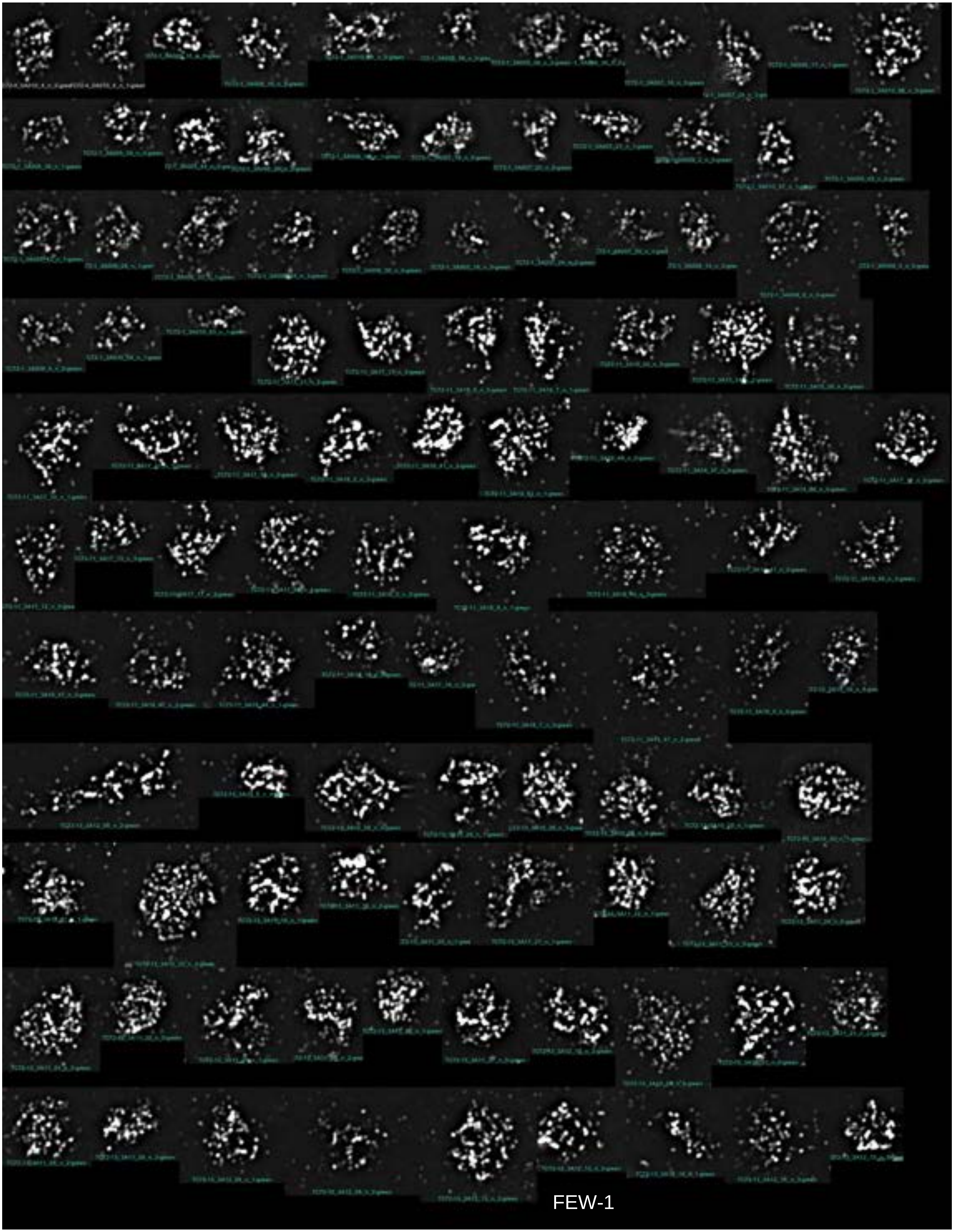

FEW-1

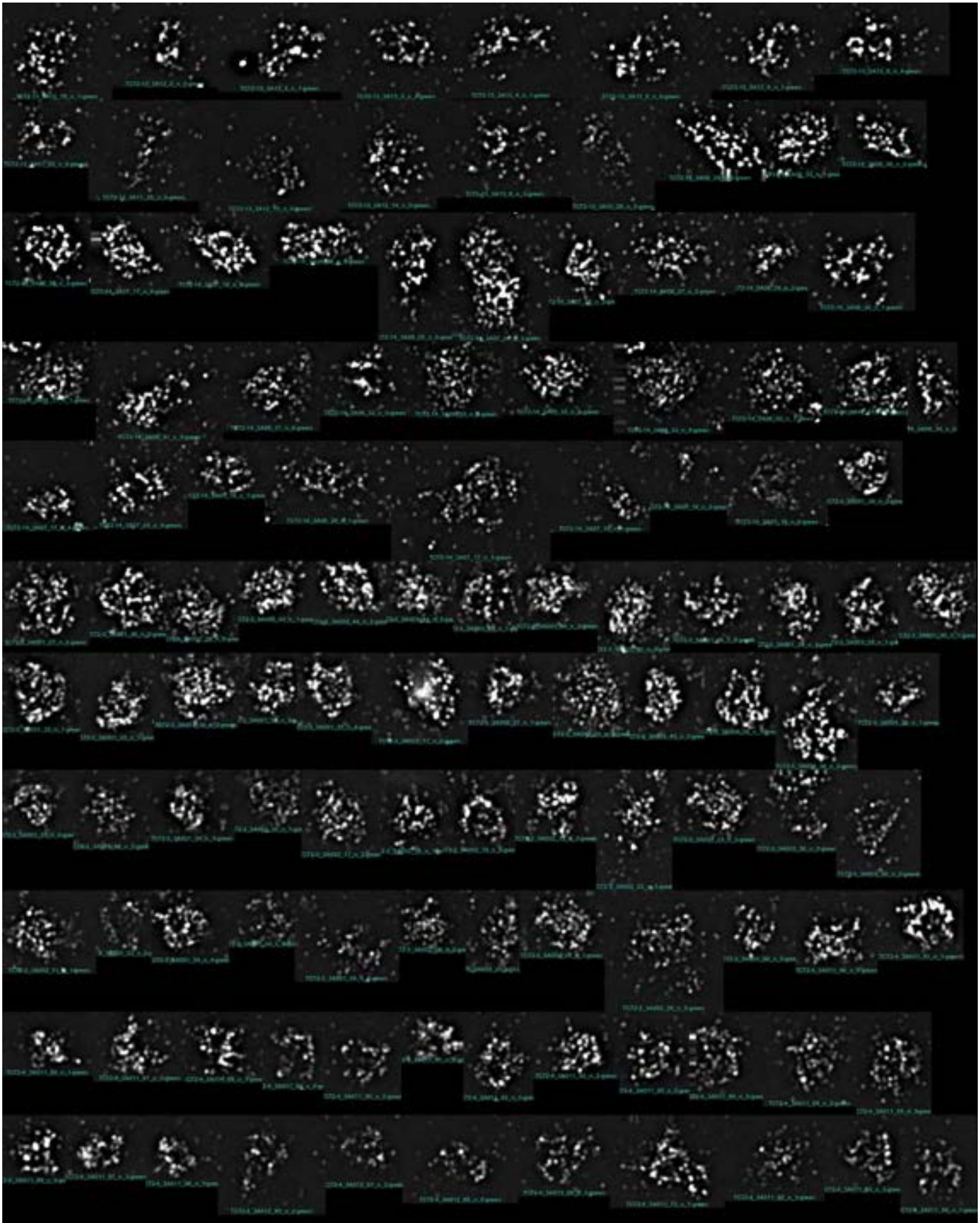

FEW-2

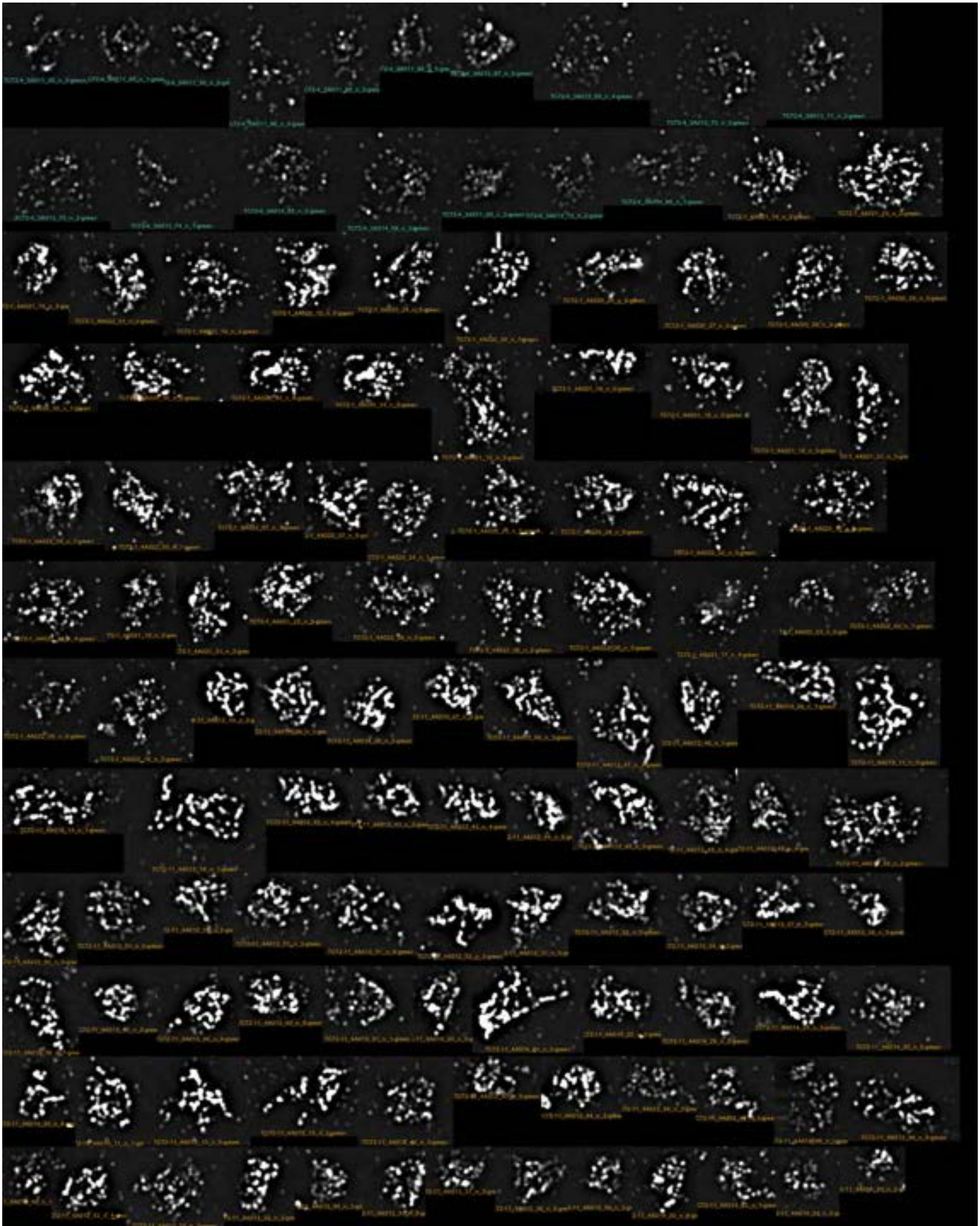

FEW-3

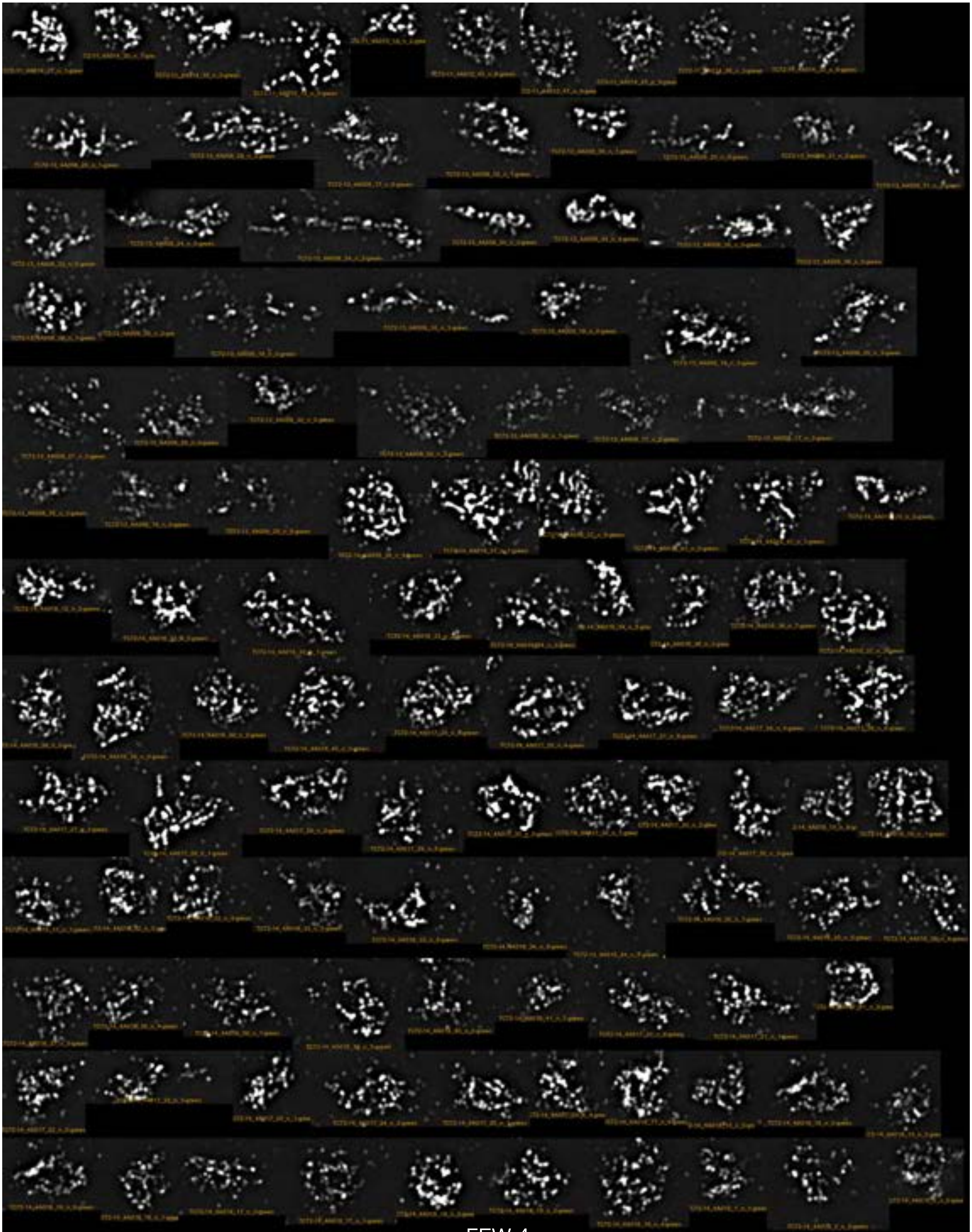

FEW-4

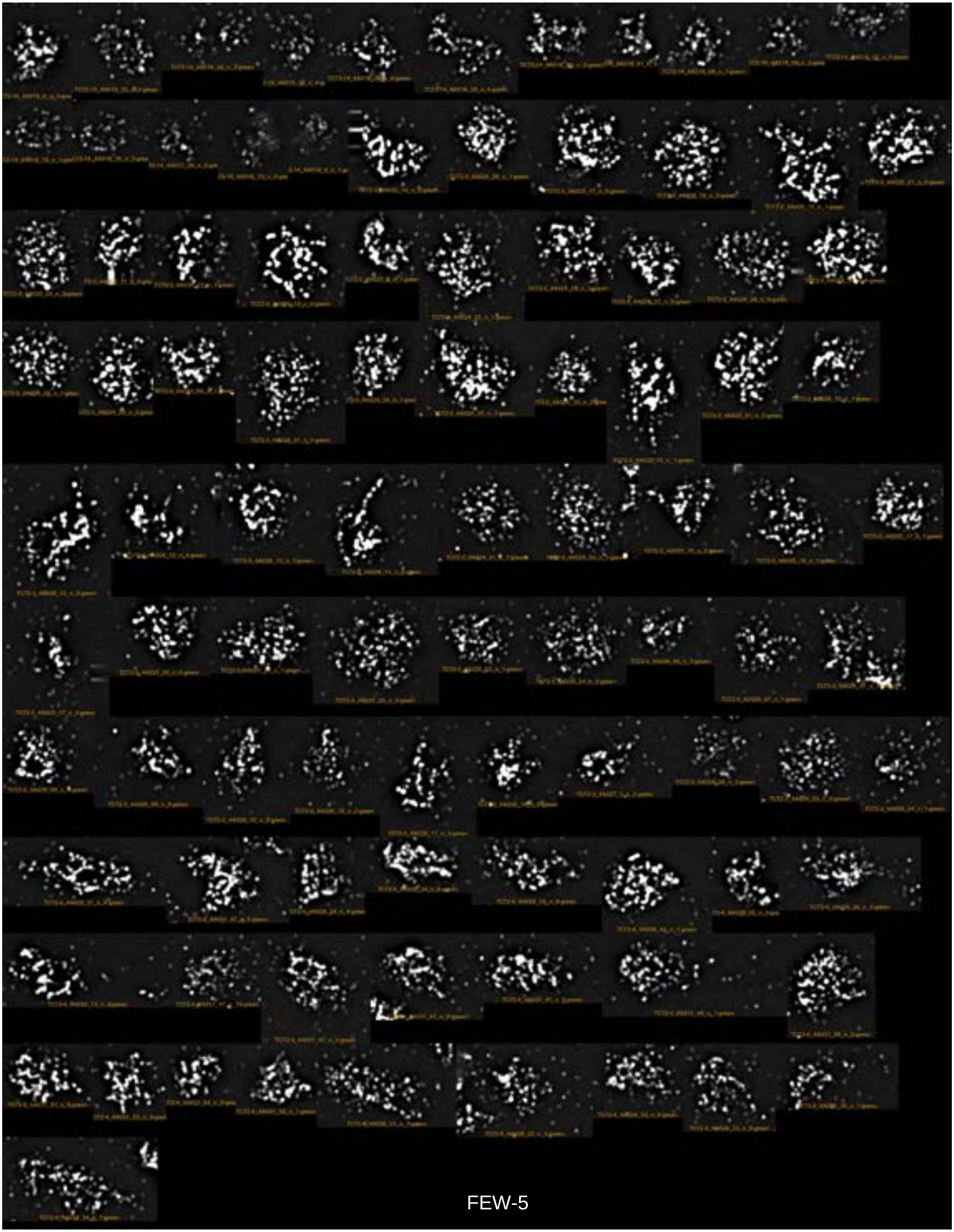

FEW-5

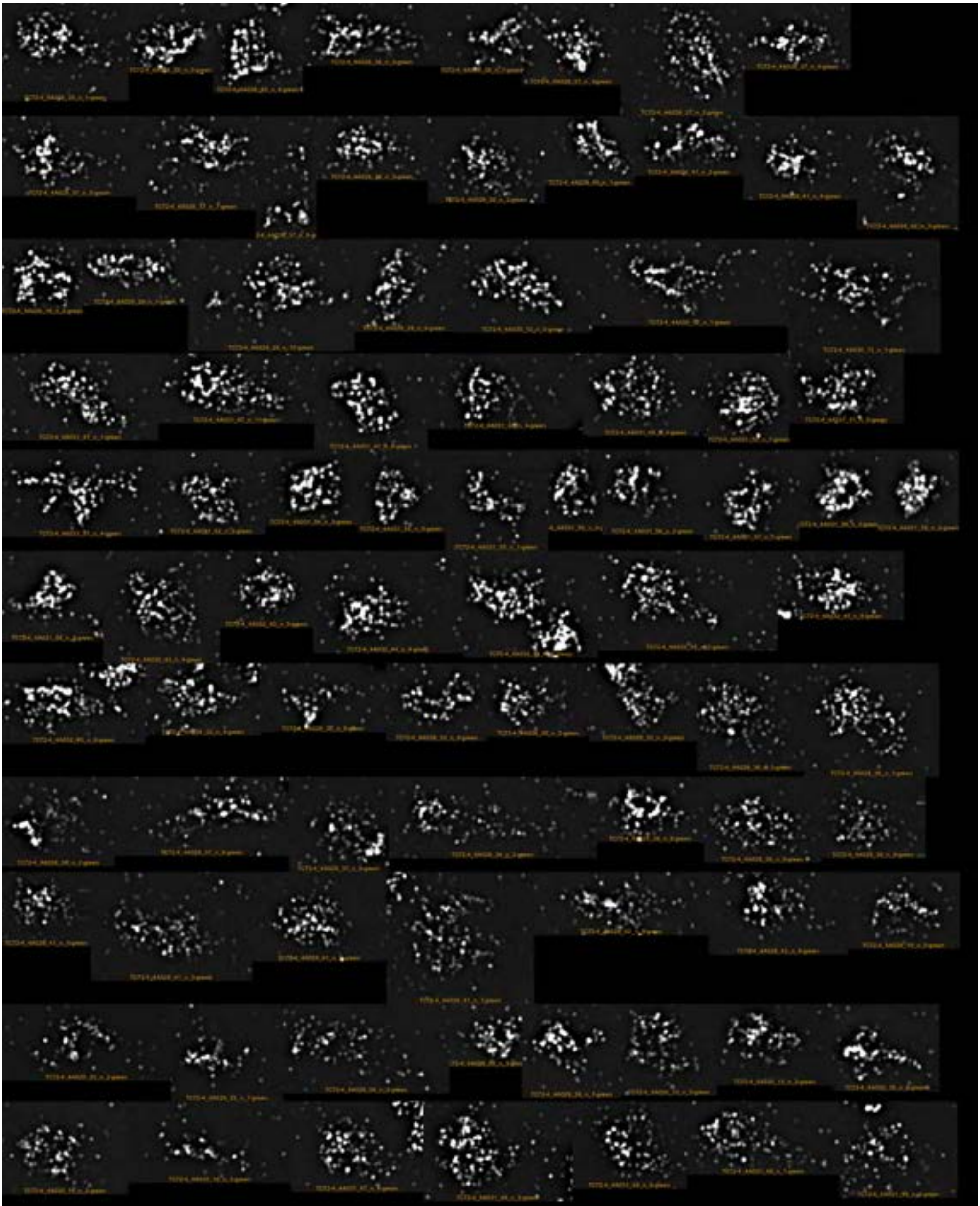

FEW-6

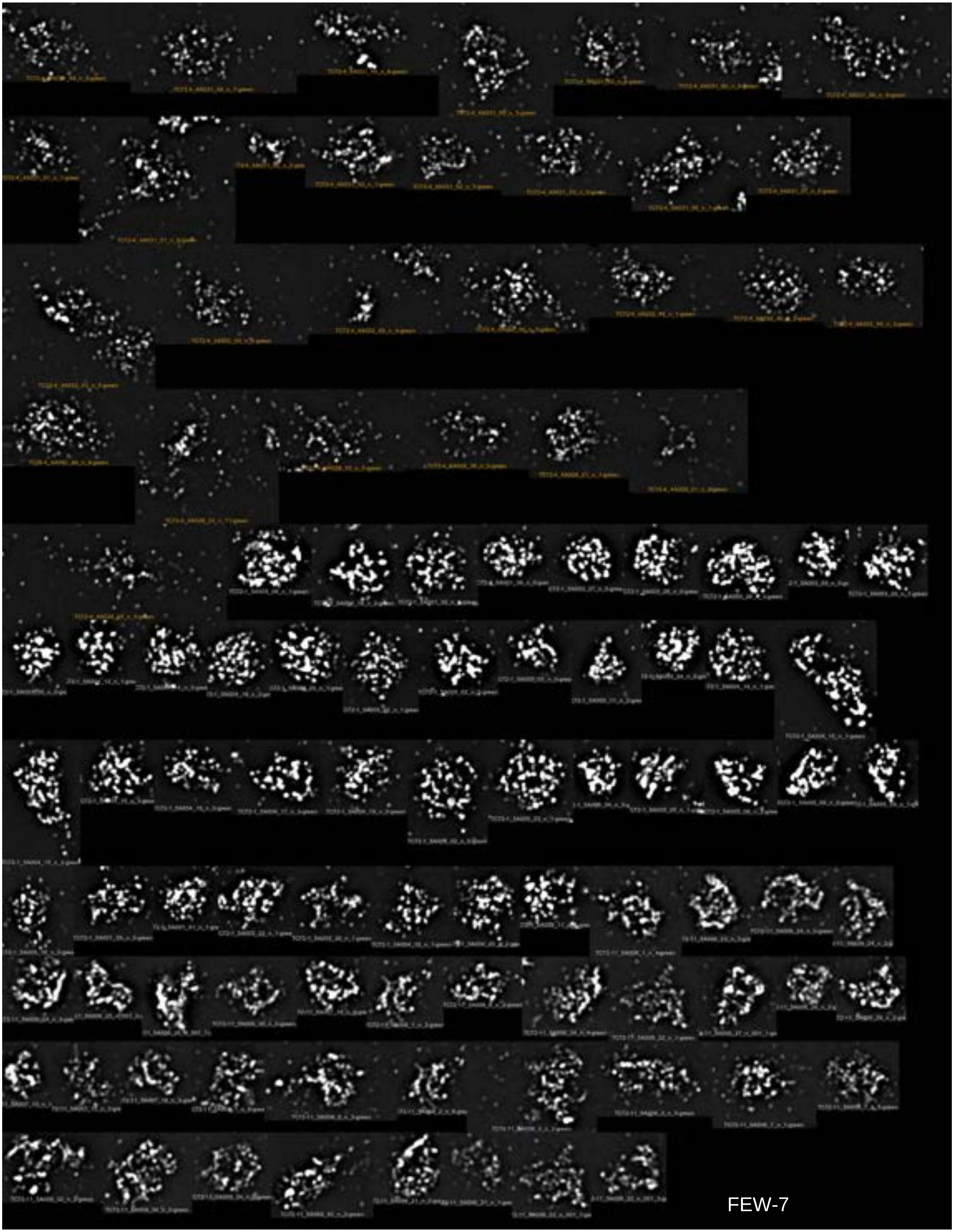

FEW-7

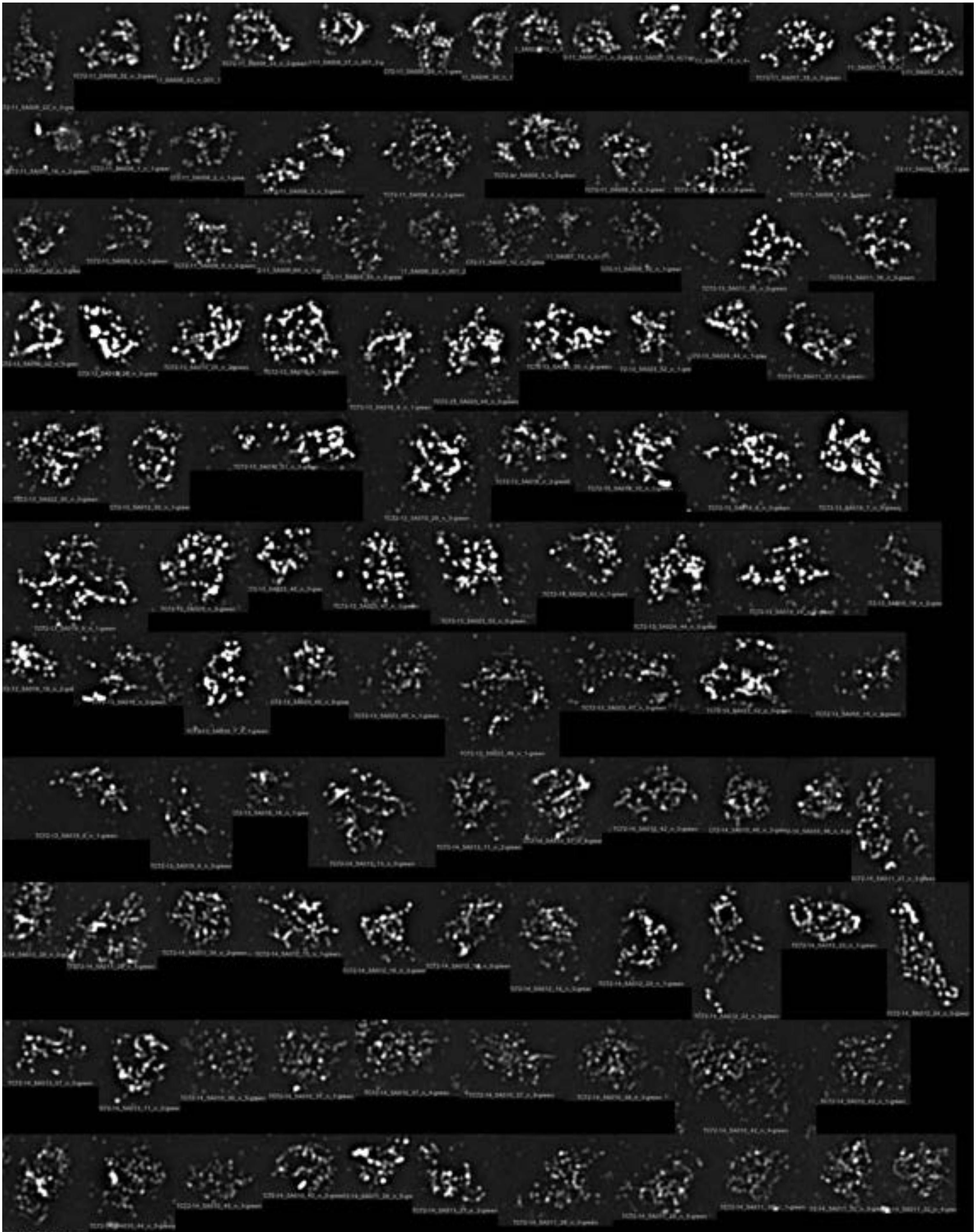

FEW-8

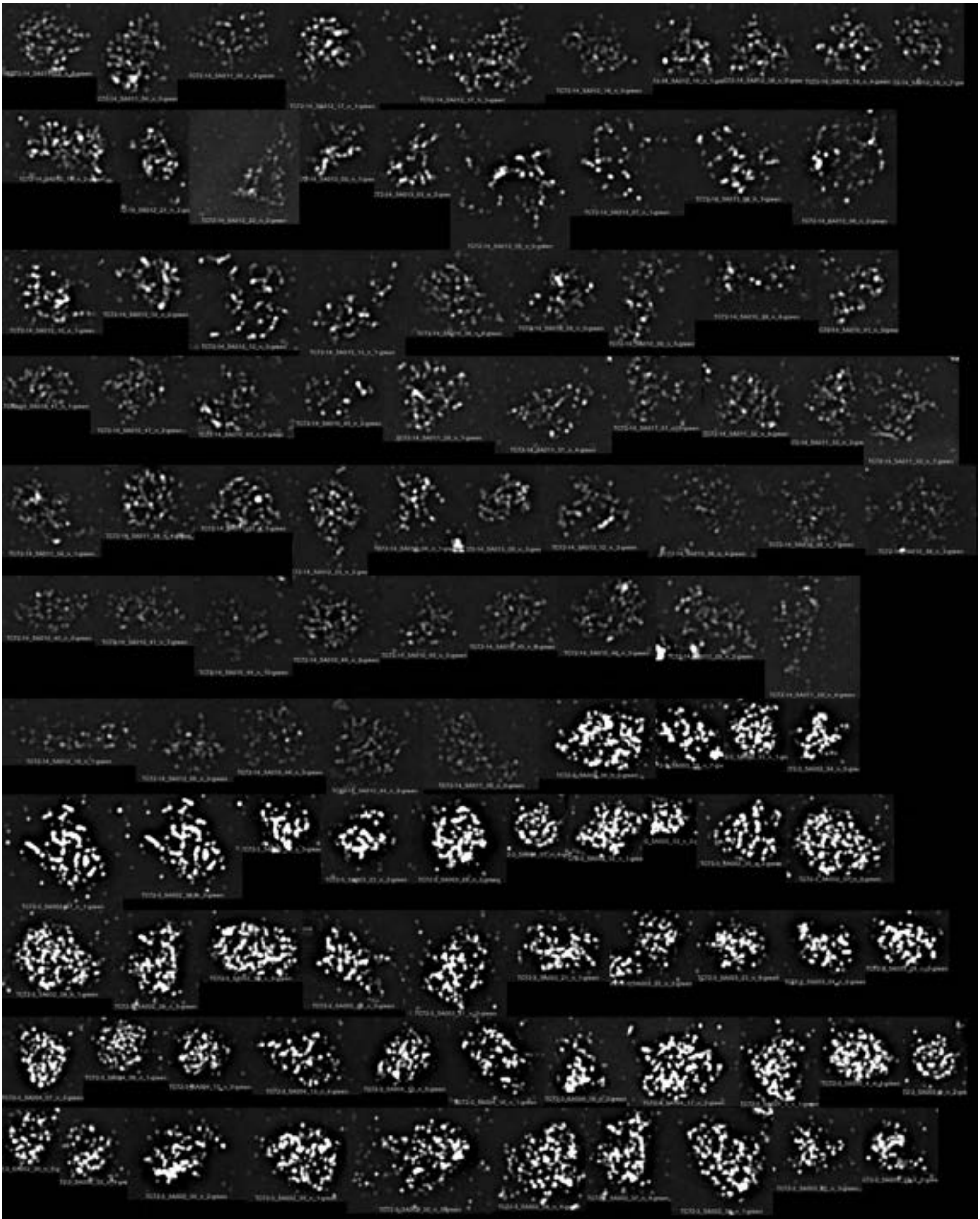

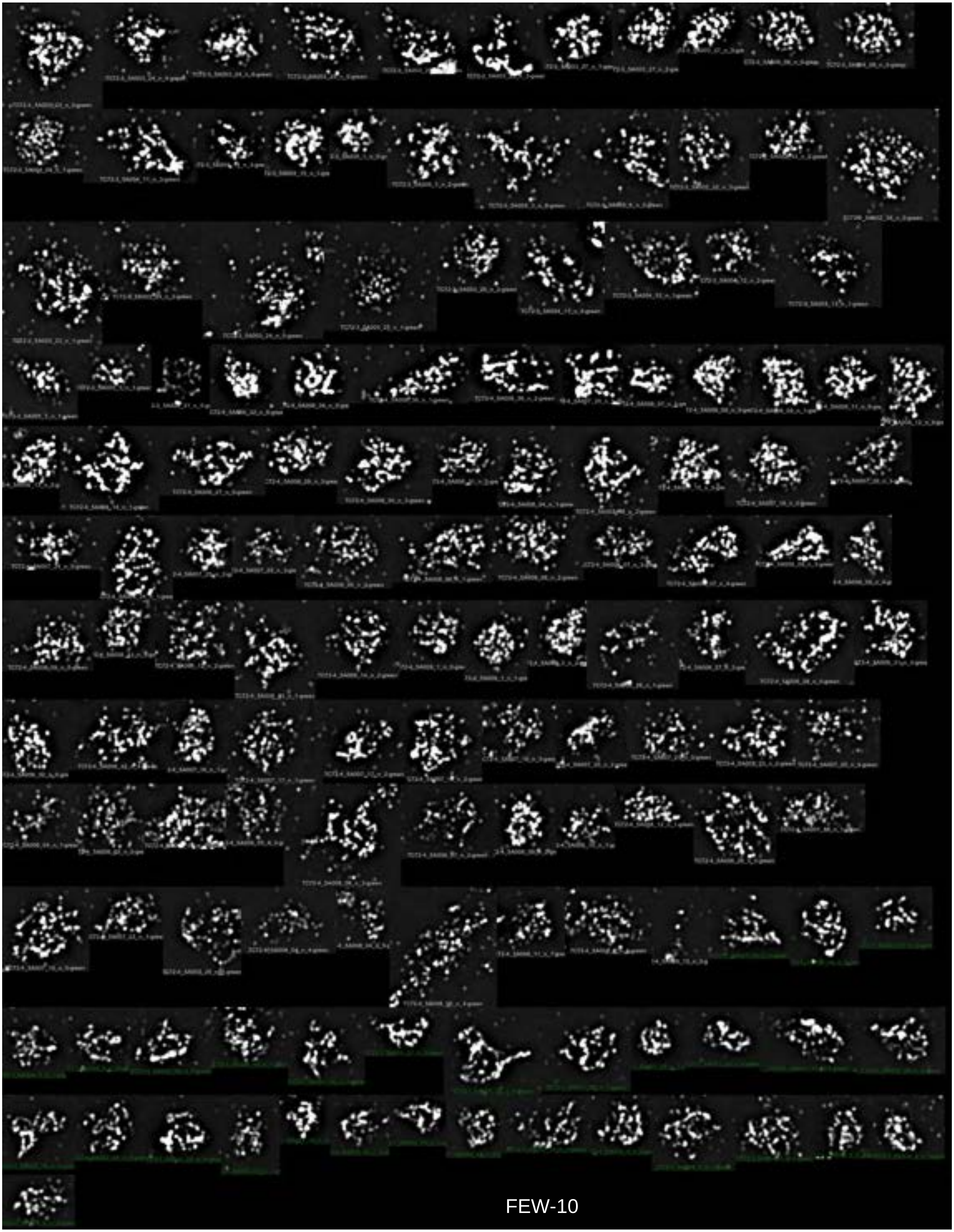

FEW-10

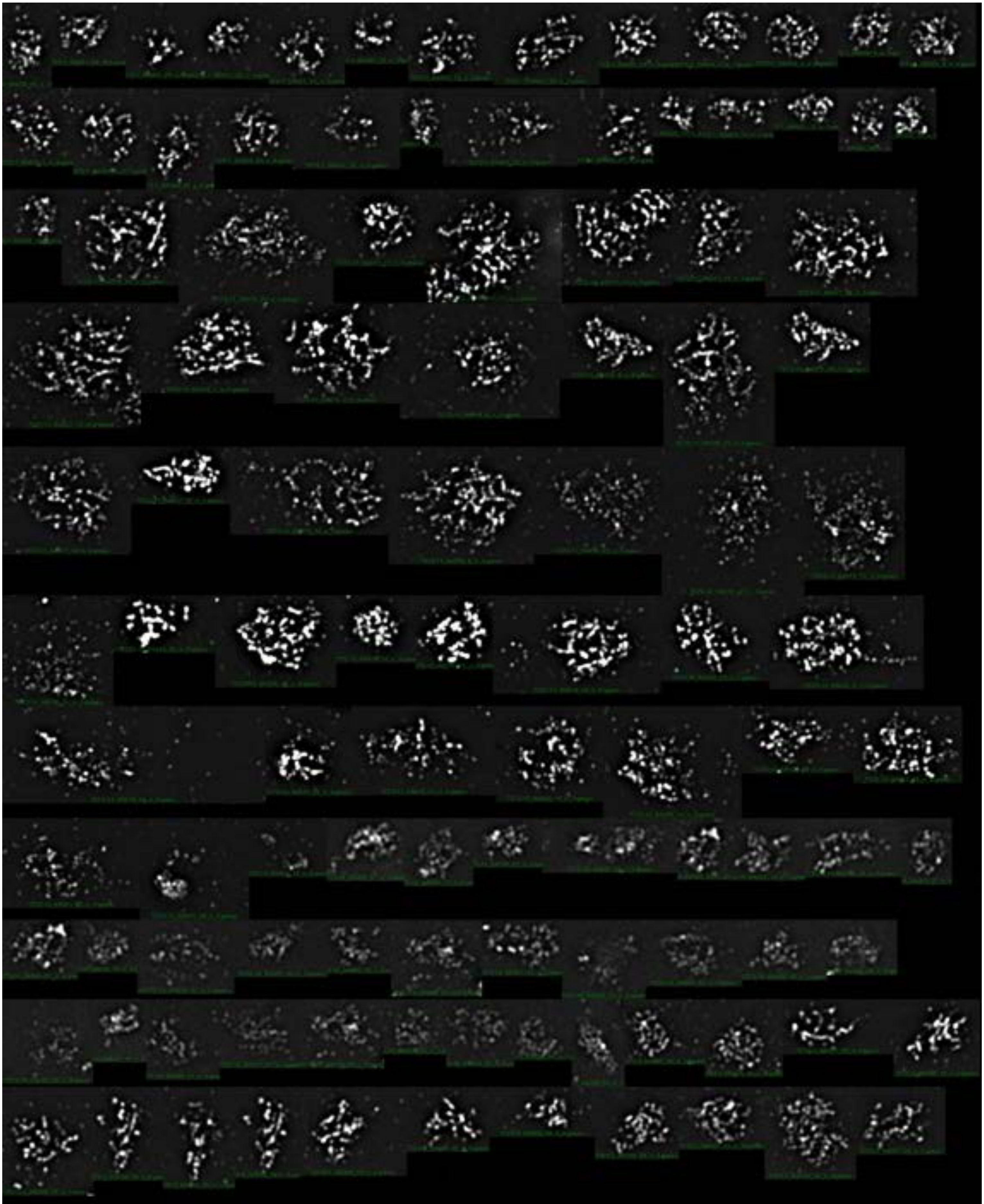

FEW-11

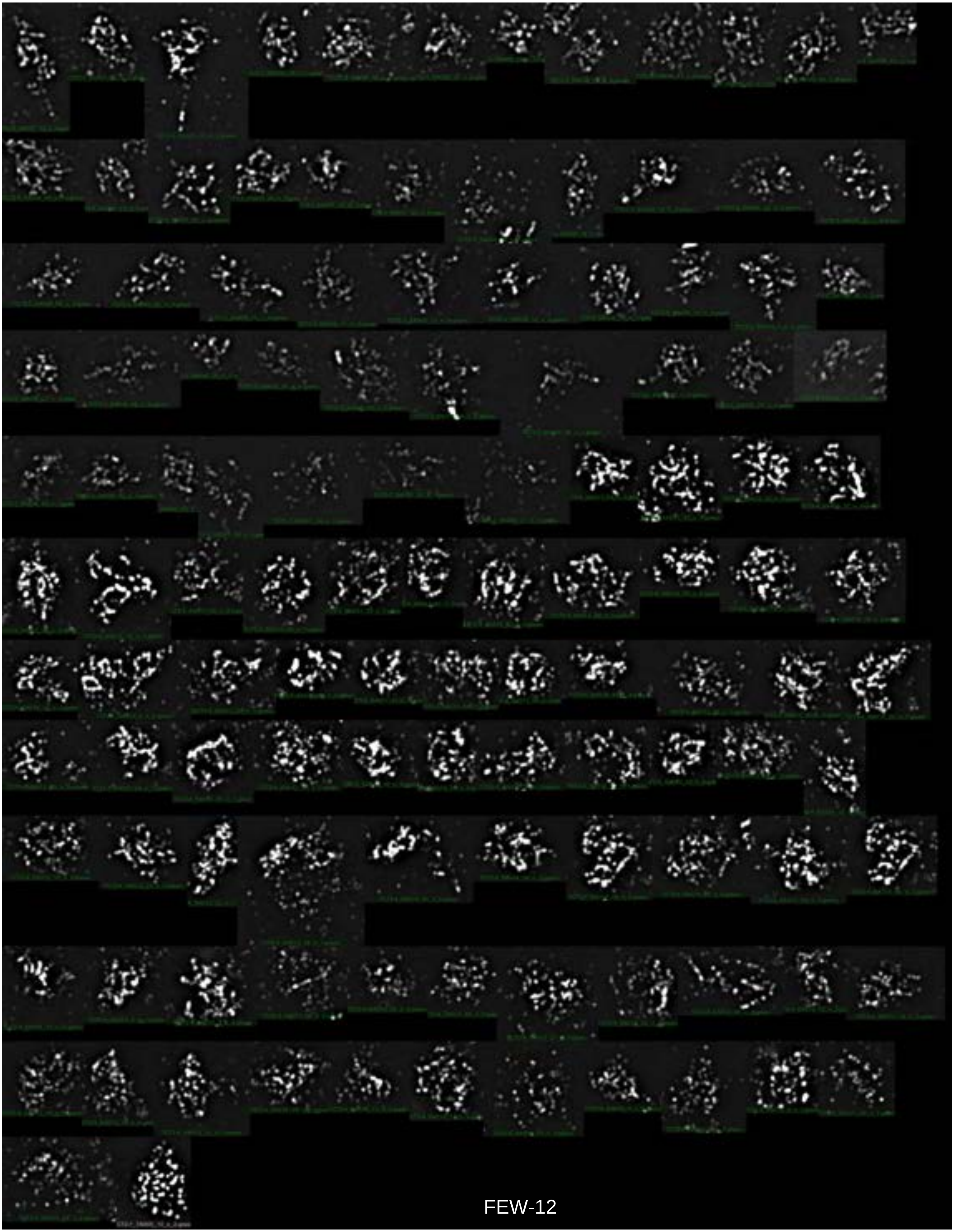

FEW-12

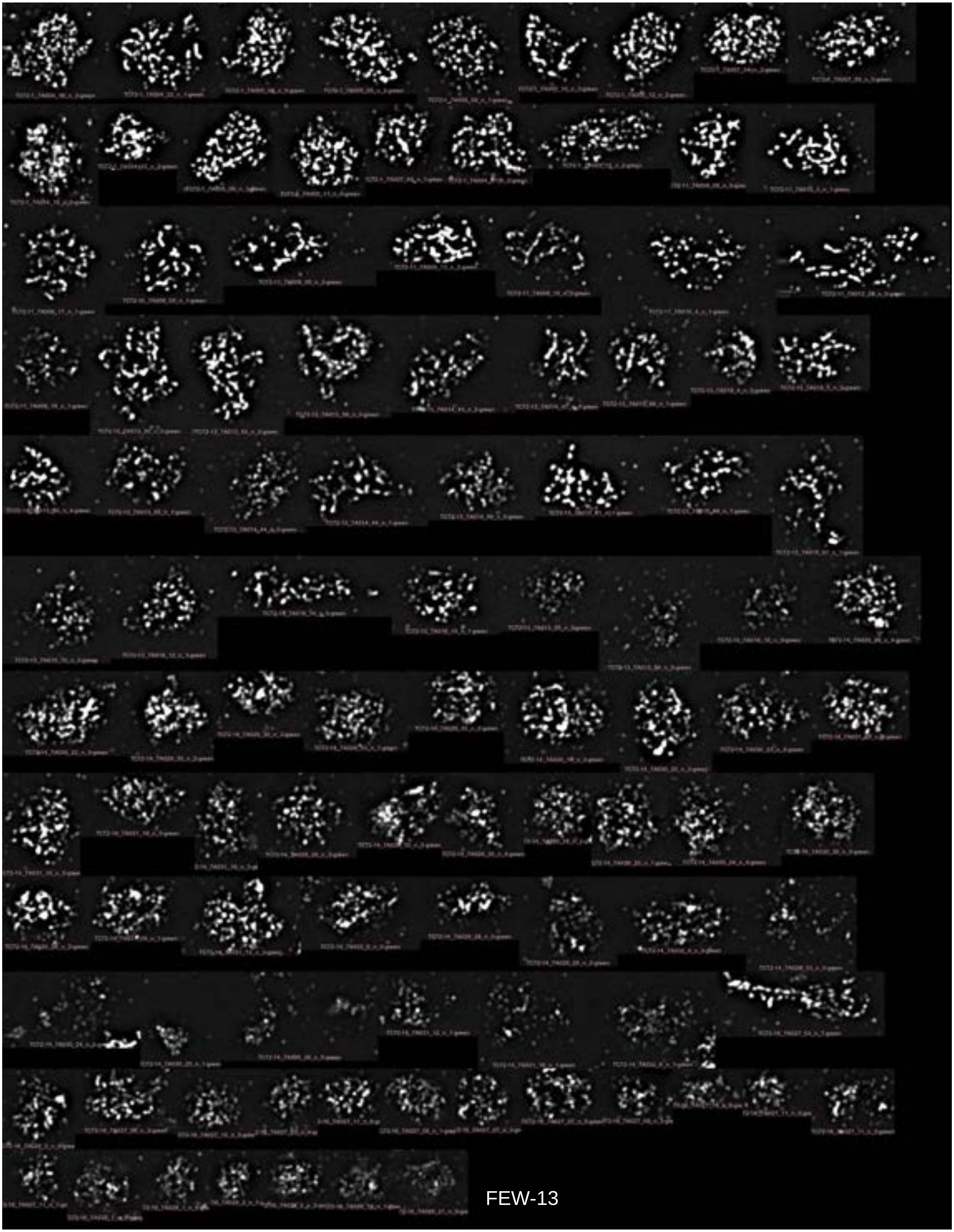

FEW-13

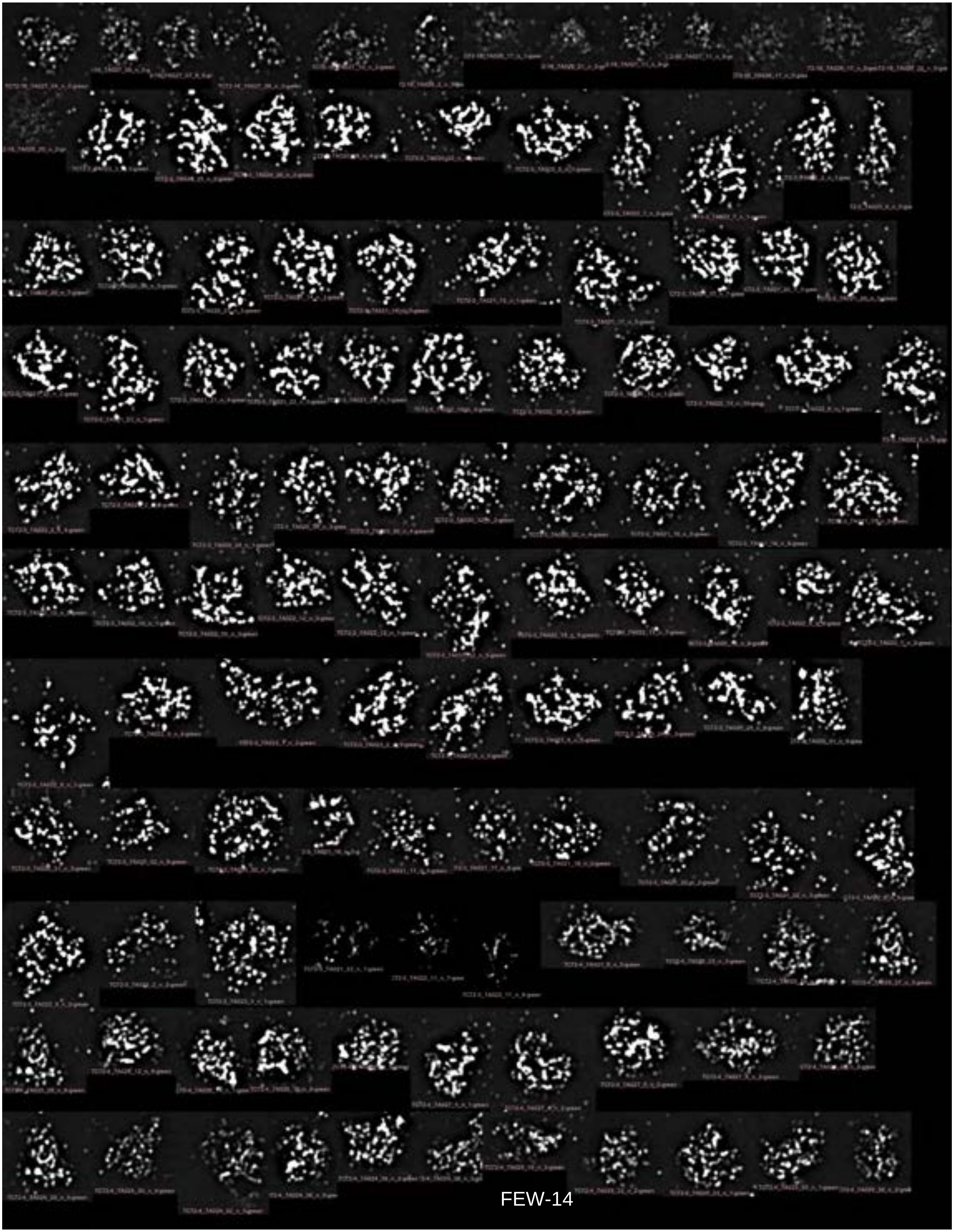

FEW-14

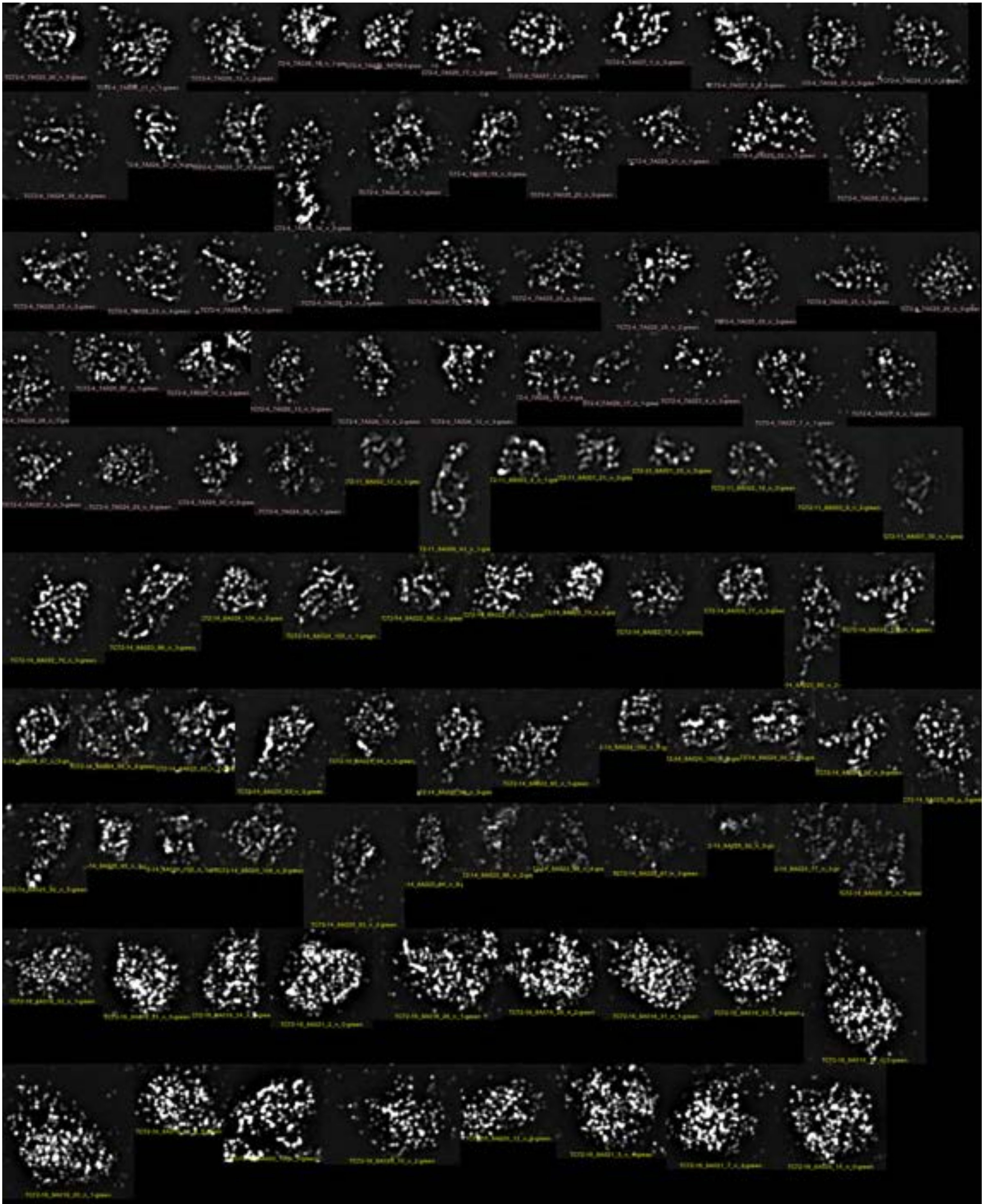

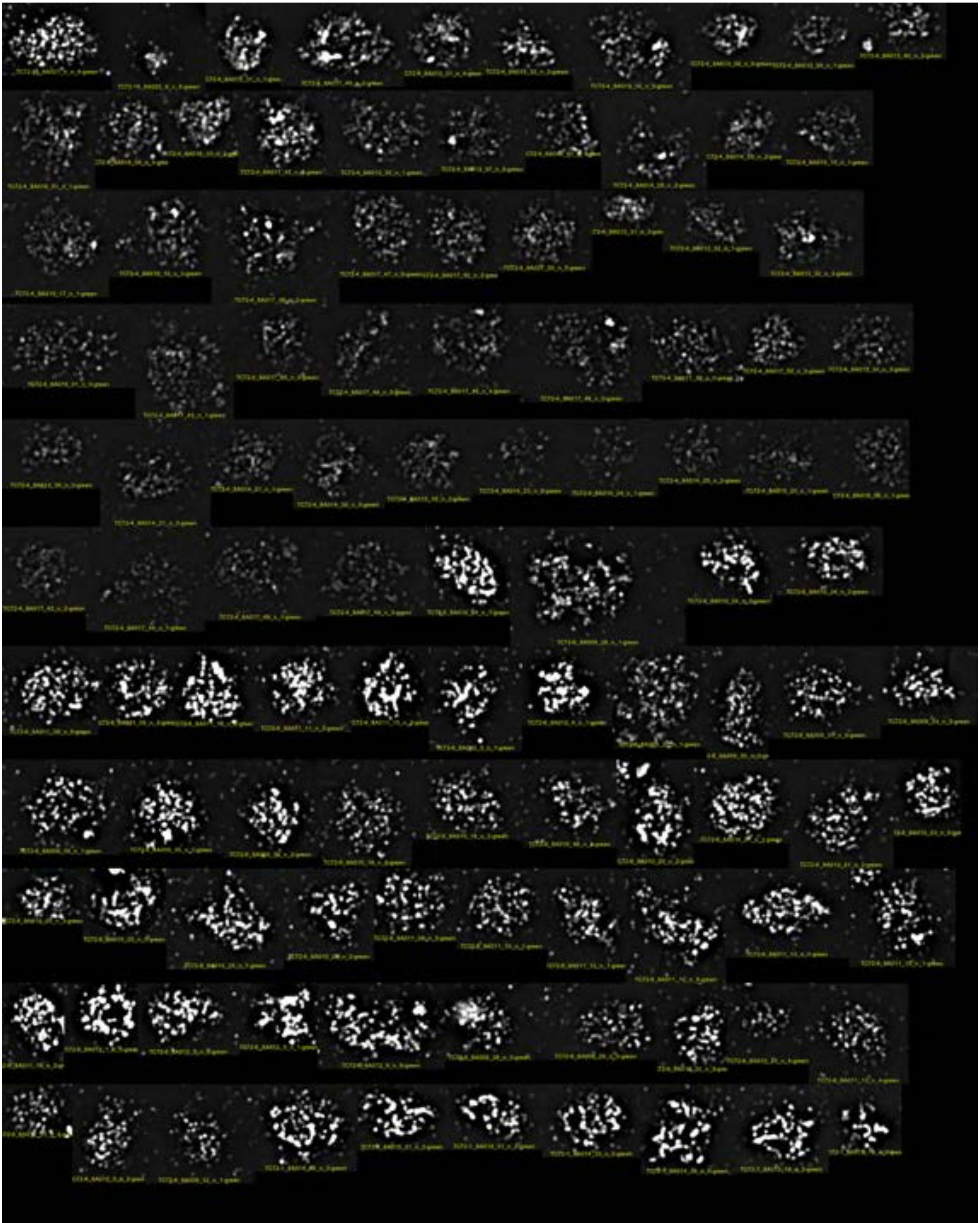

FEW-16

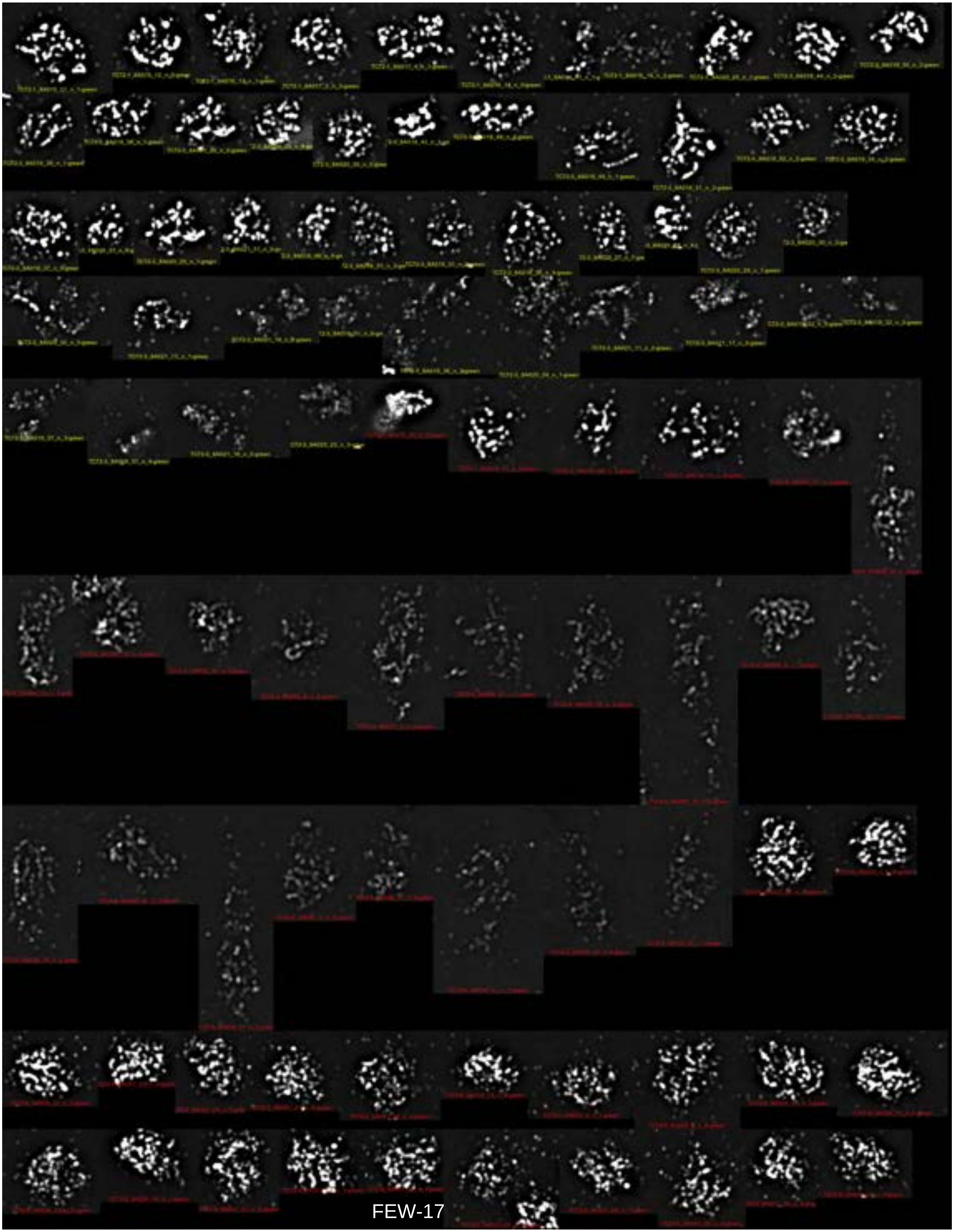

FEW-17

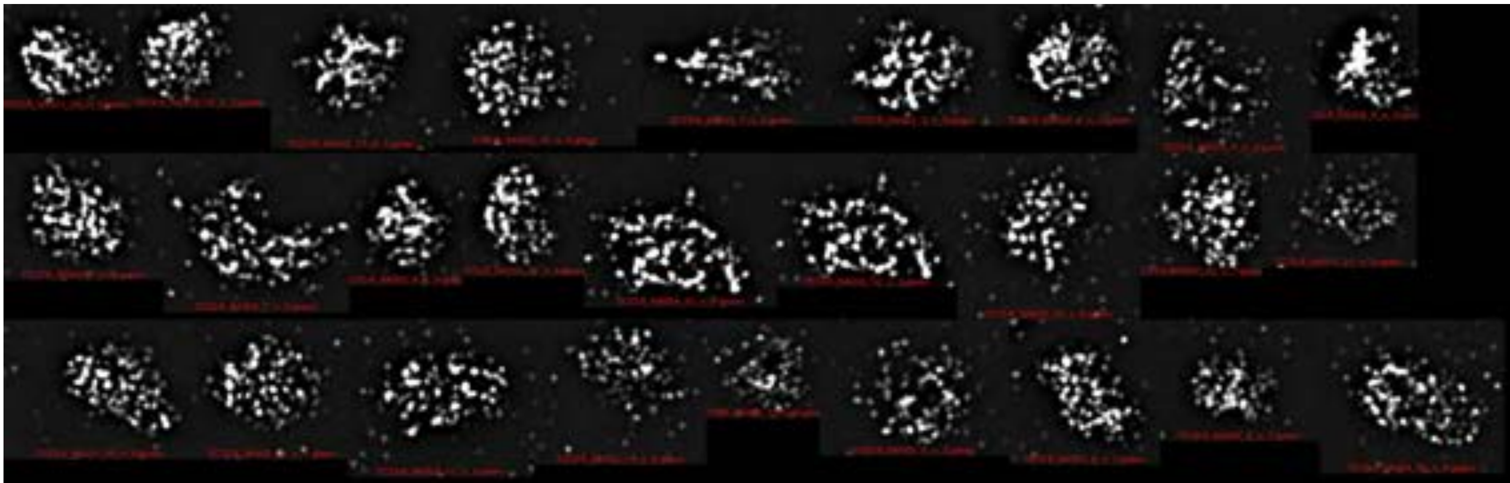

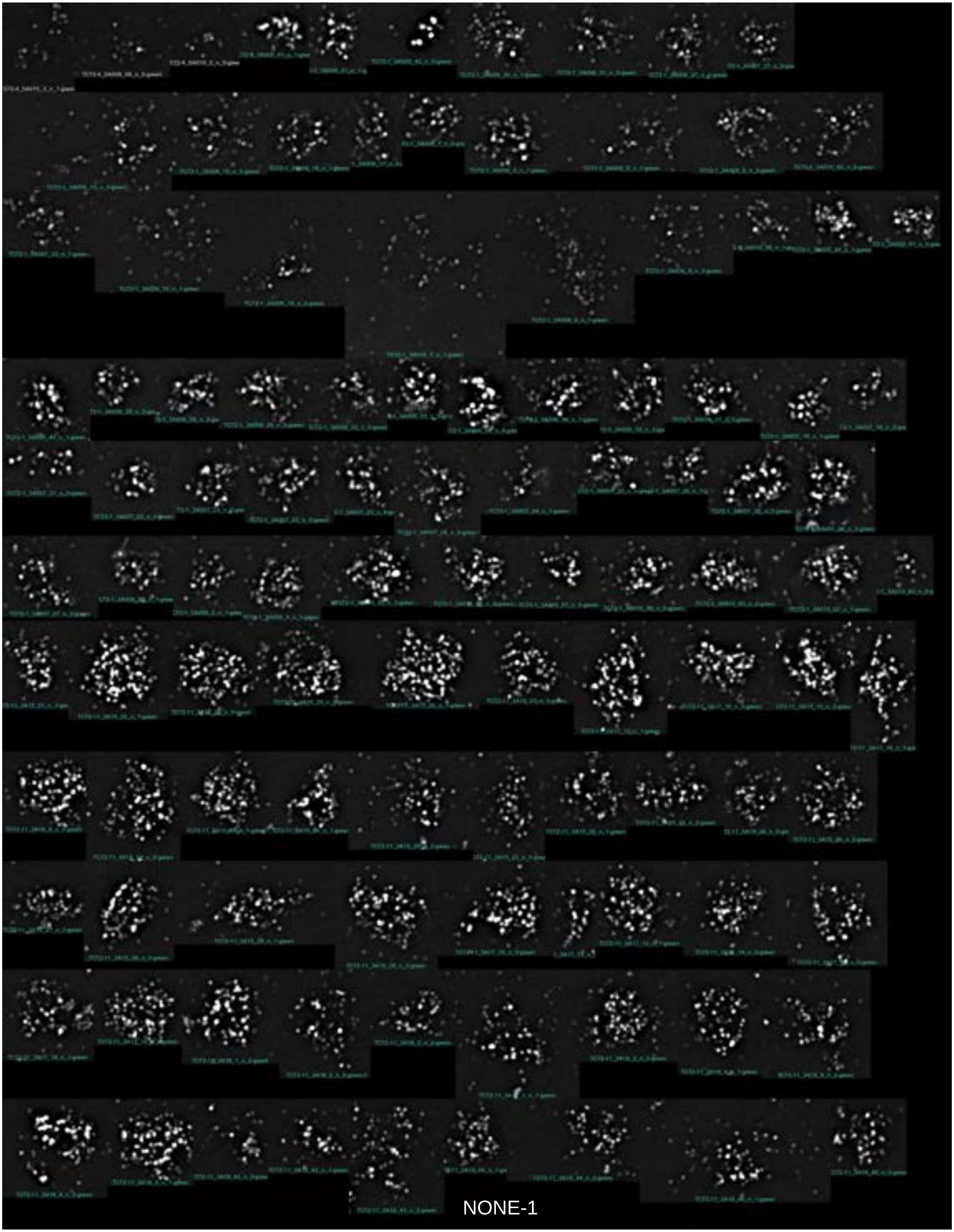

NONE-1

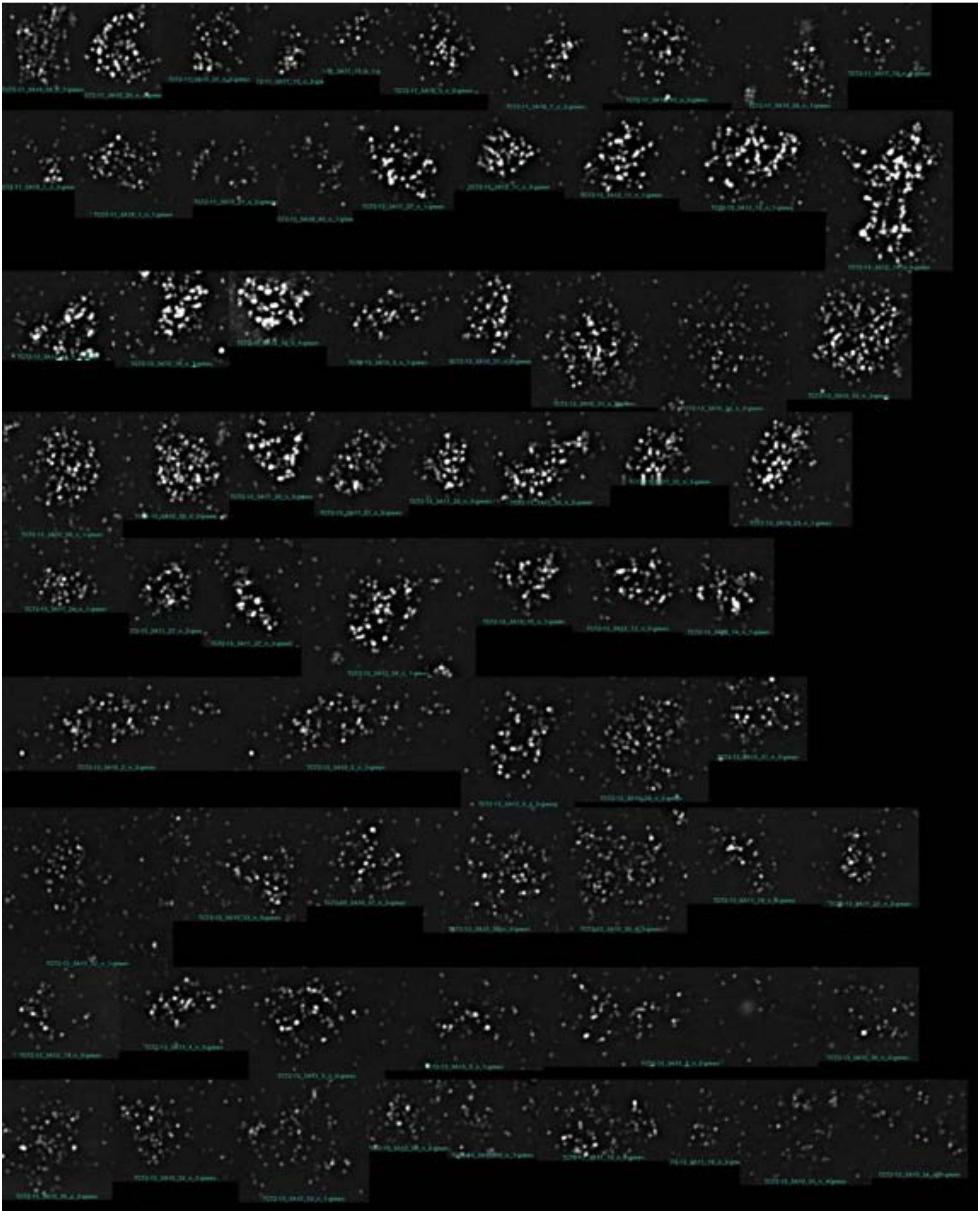

NONE-2

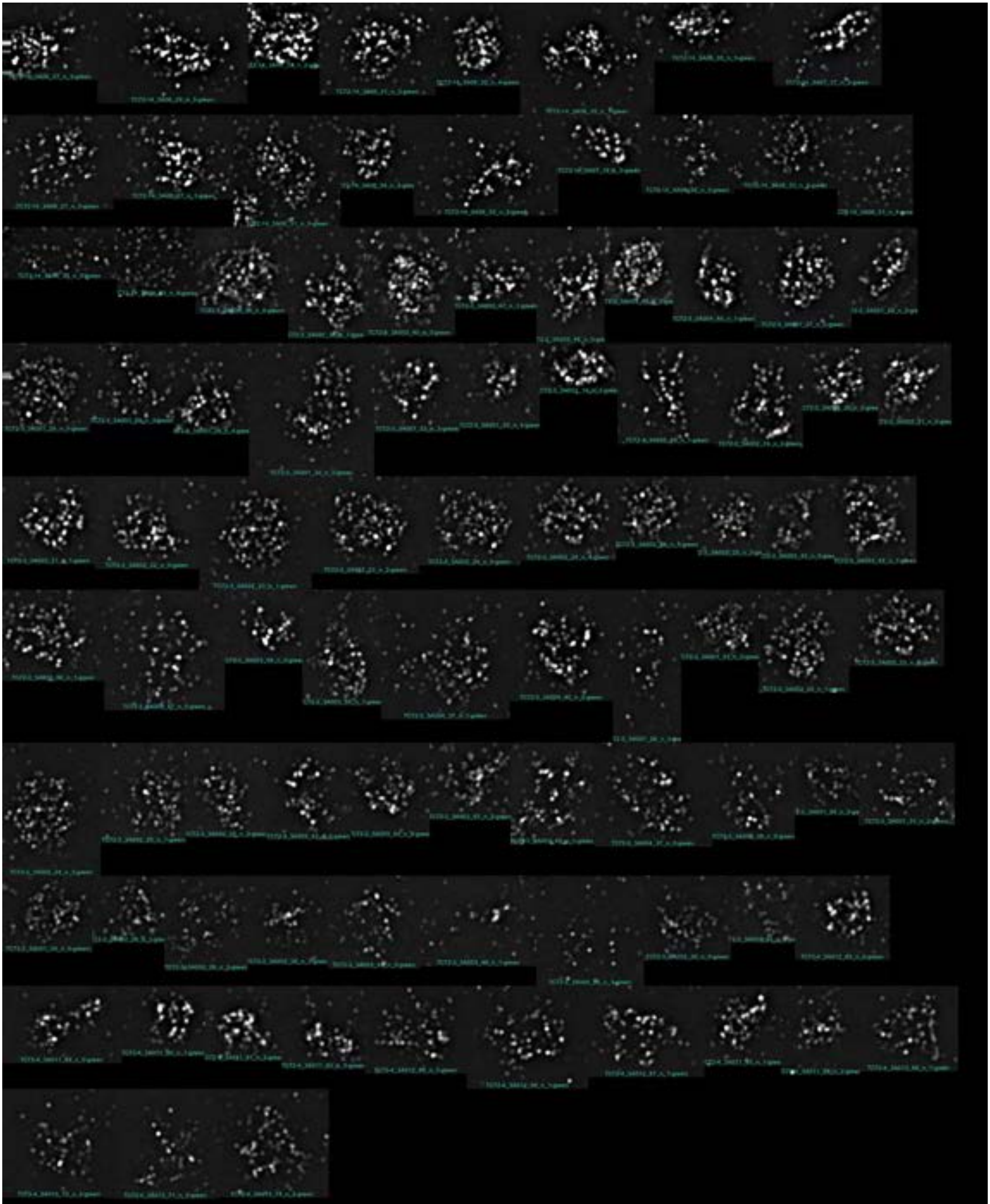

NONE-3

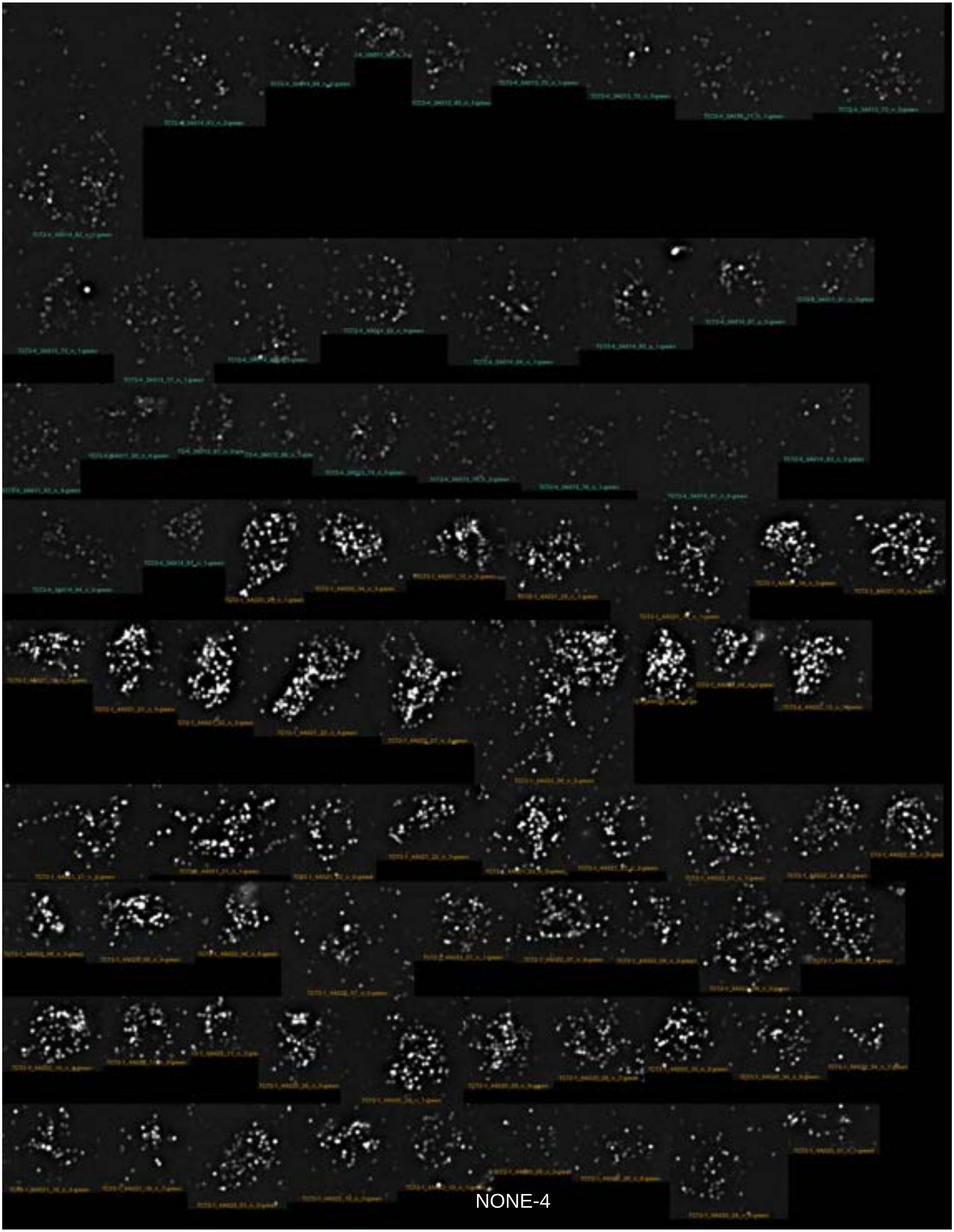

NONE-4

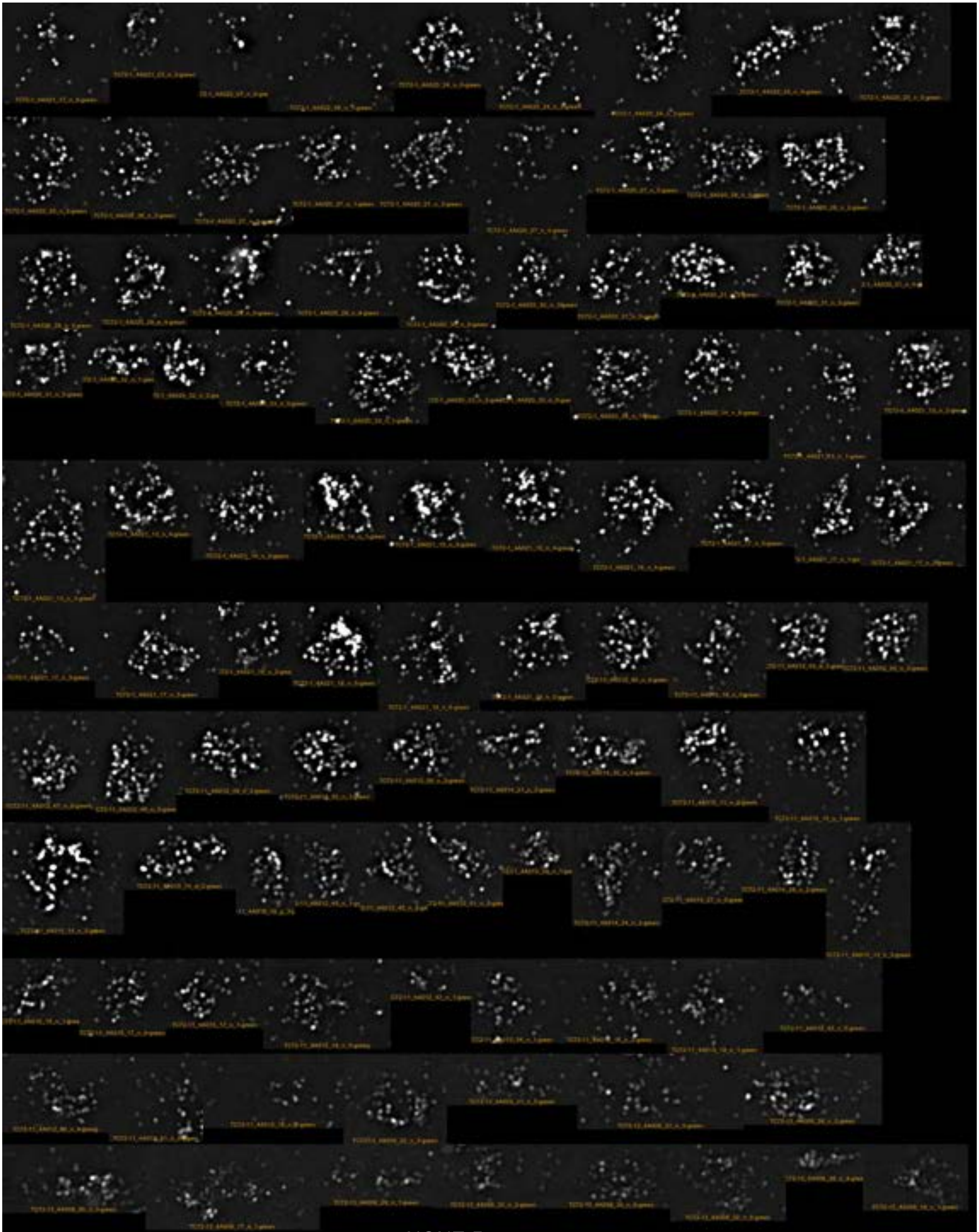

NONE-5

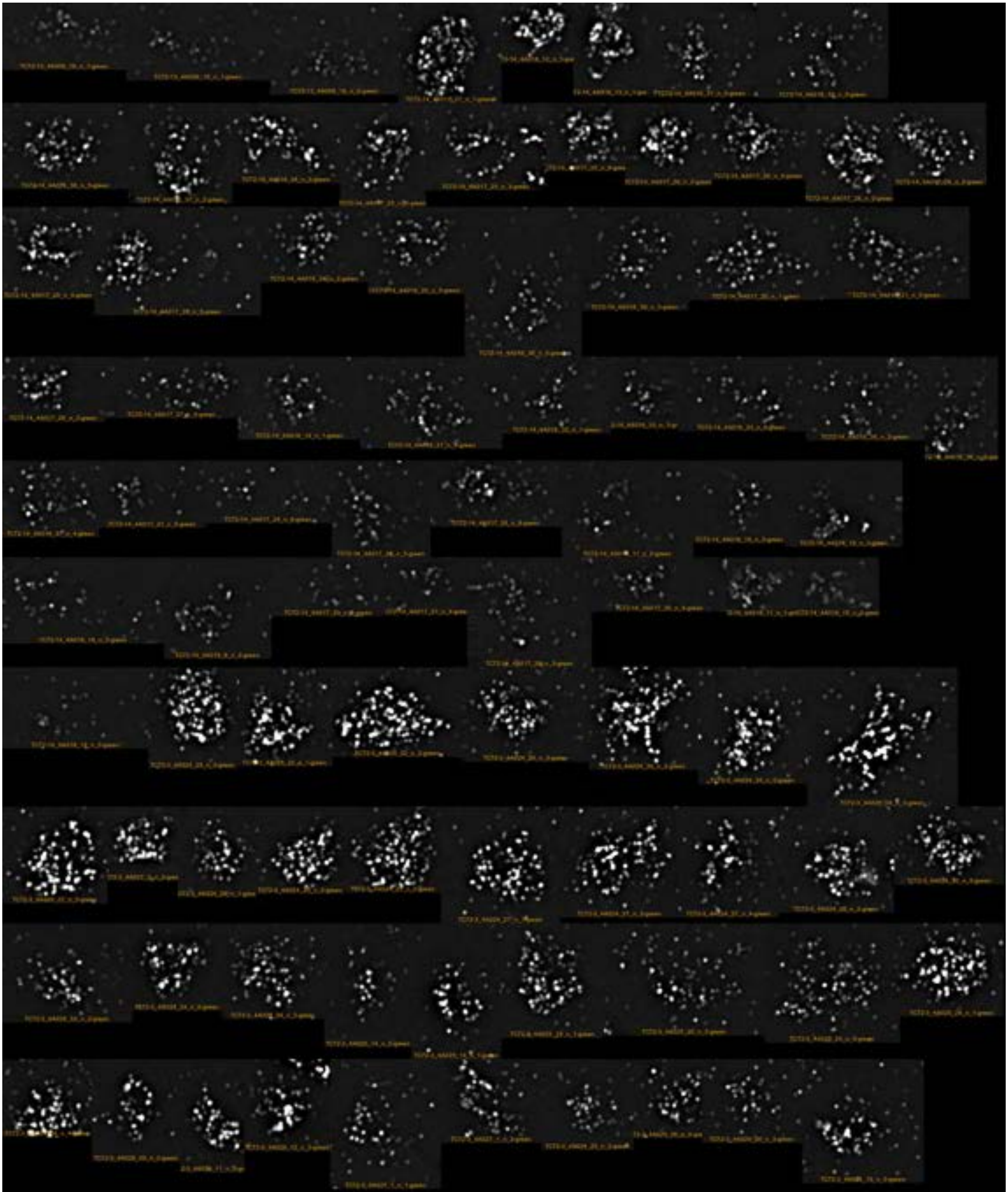

NONE-6

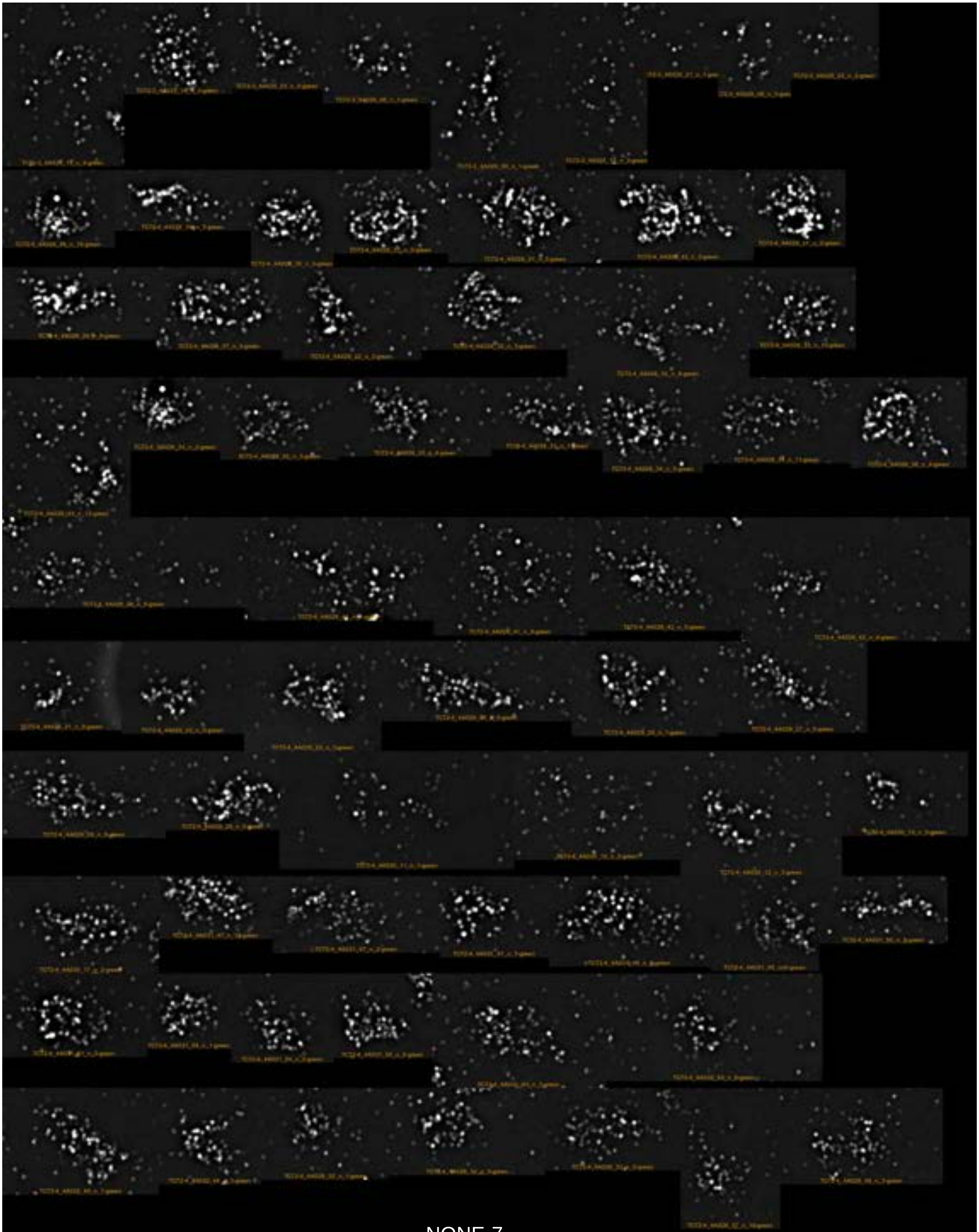

NONE-7

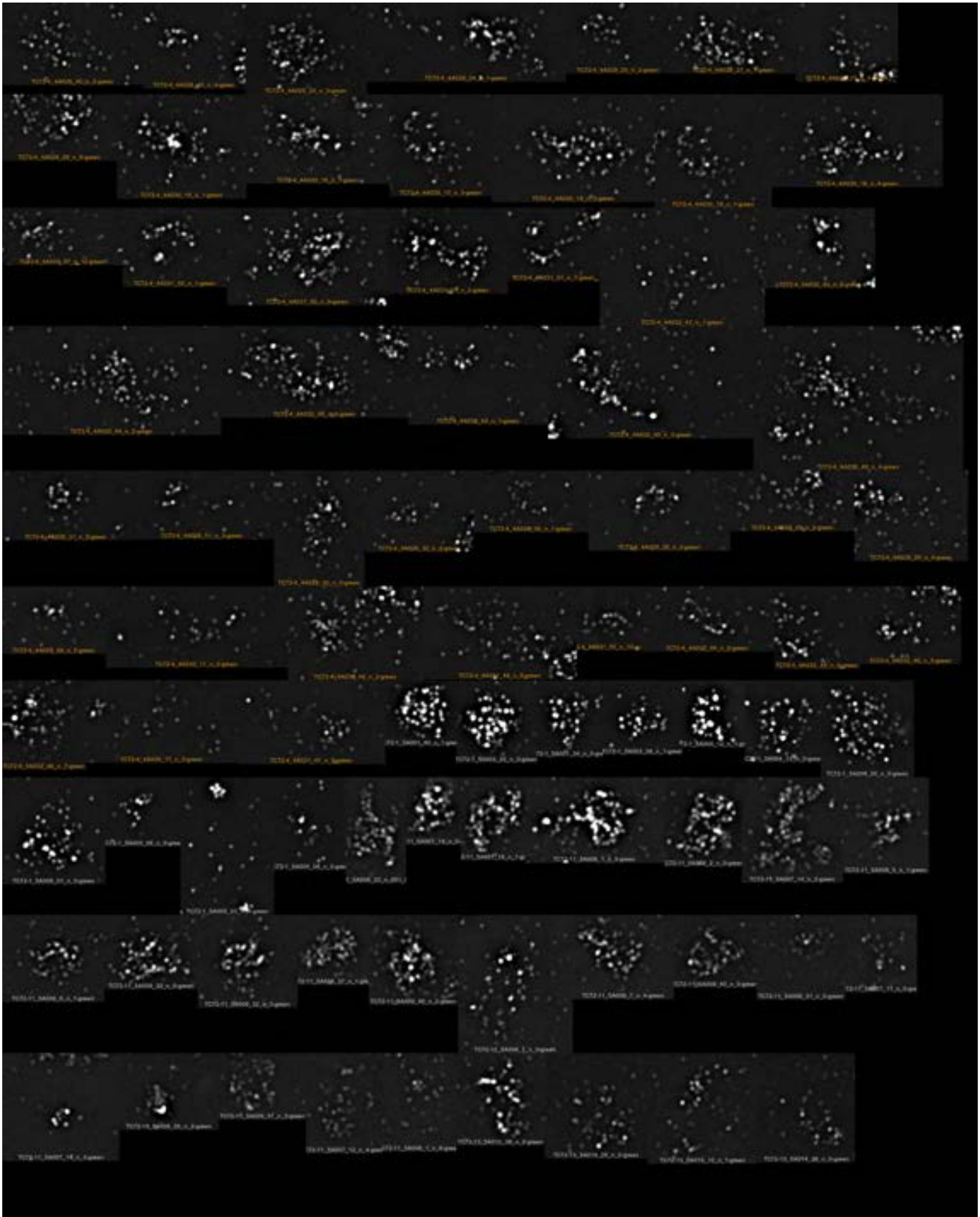

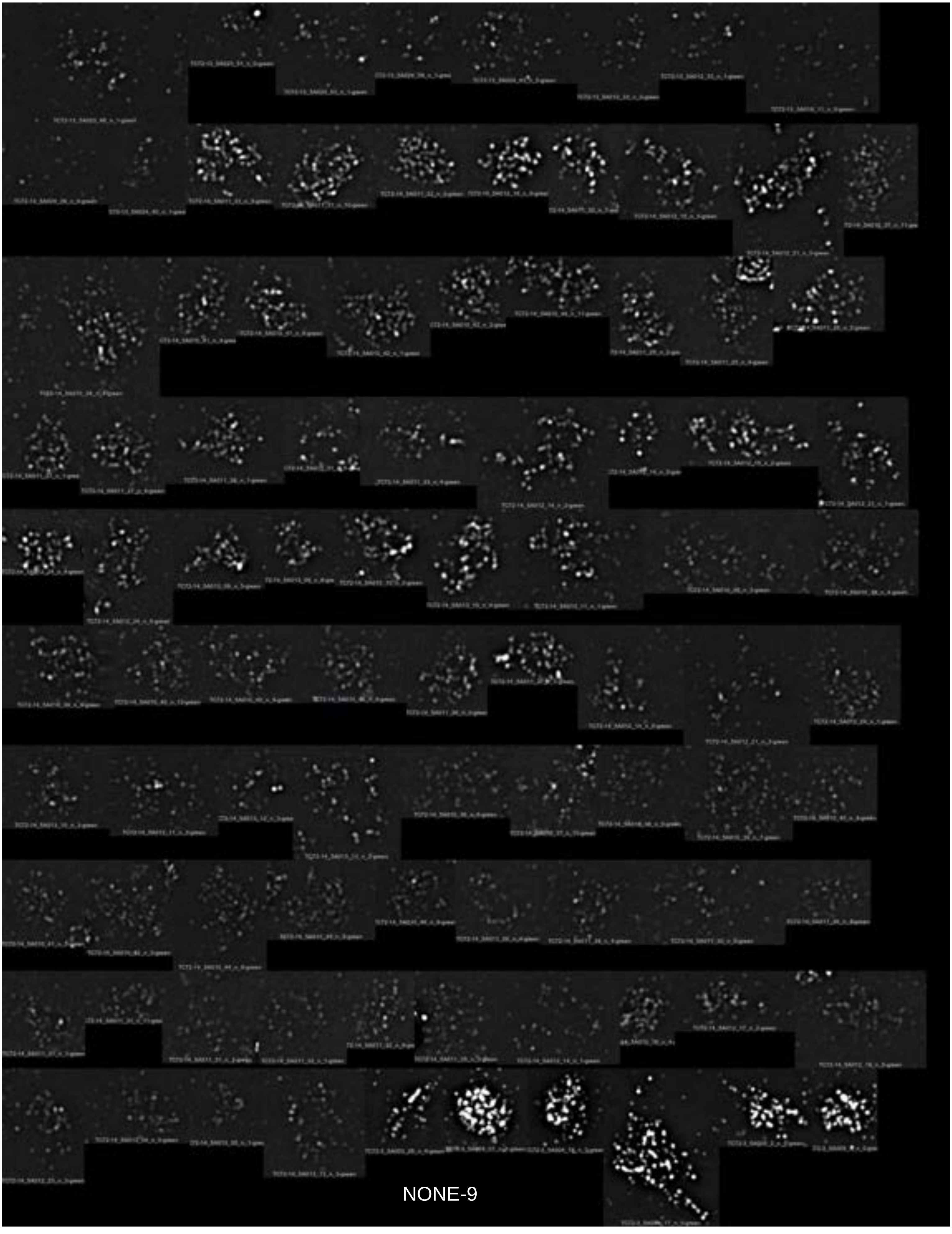

NONE-9

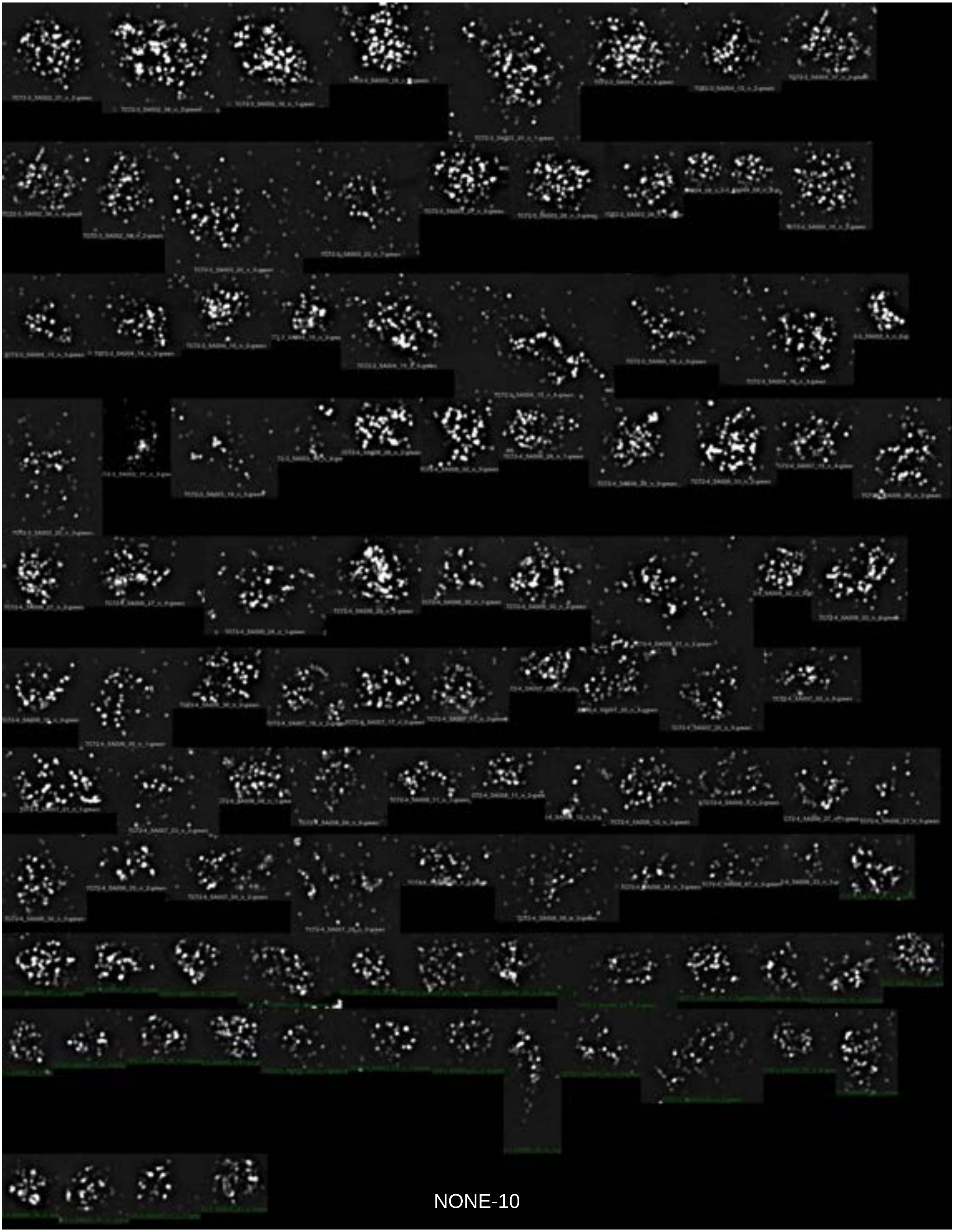

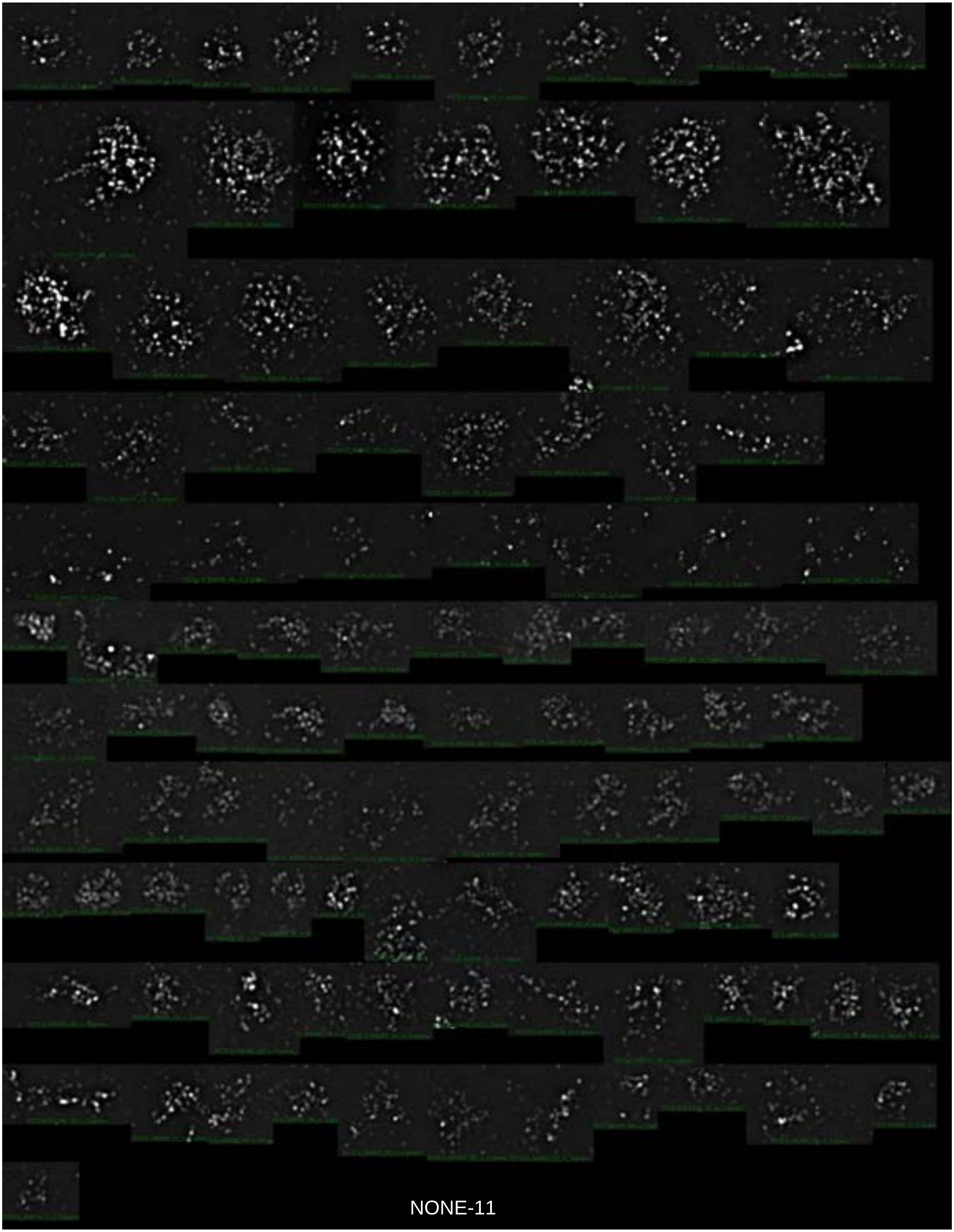

NONE-11

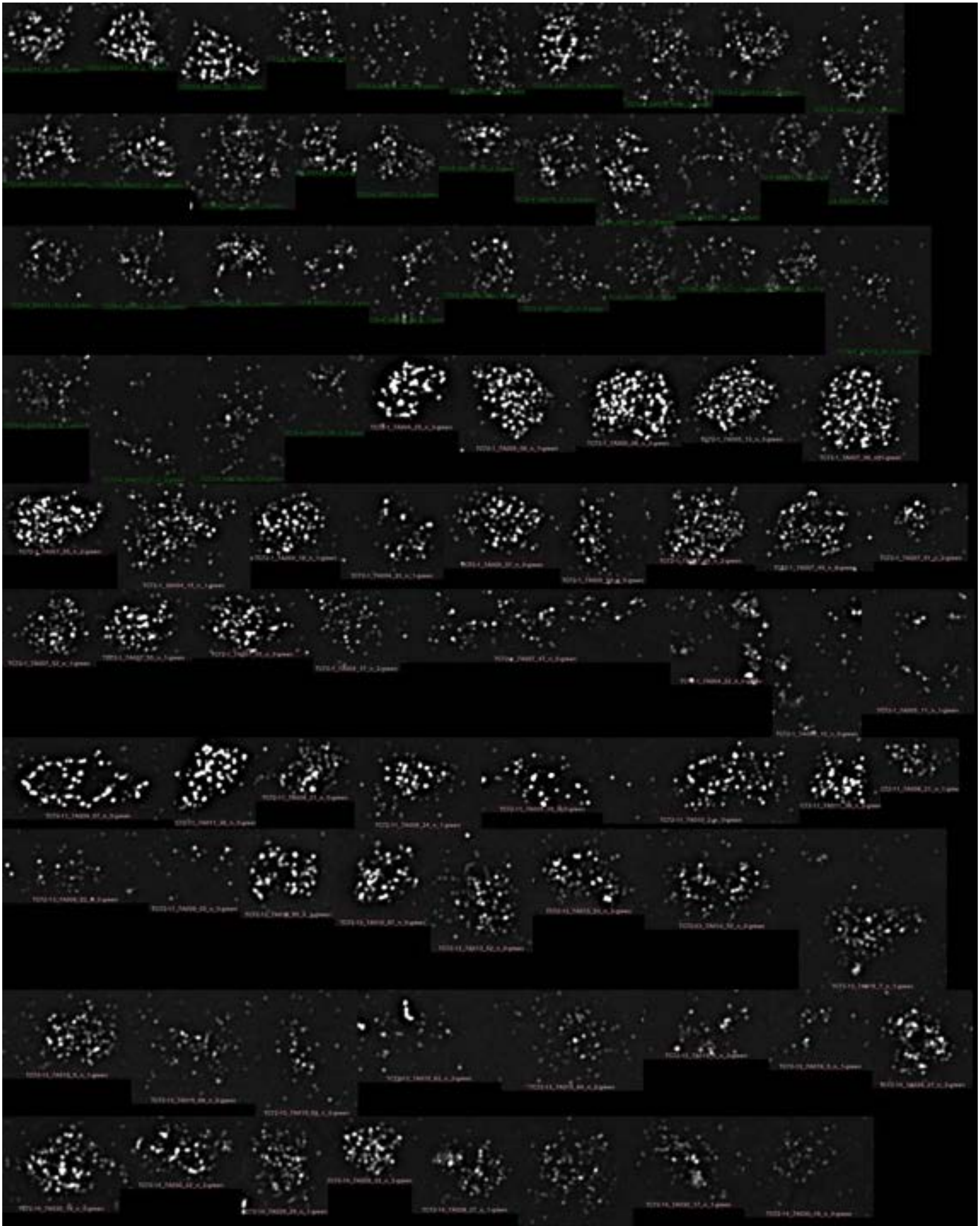

NONE-12

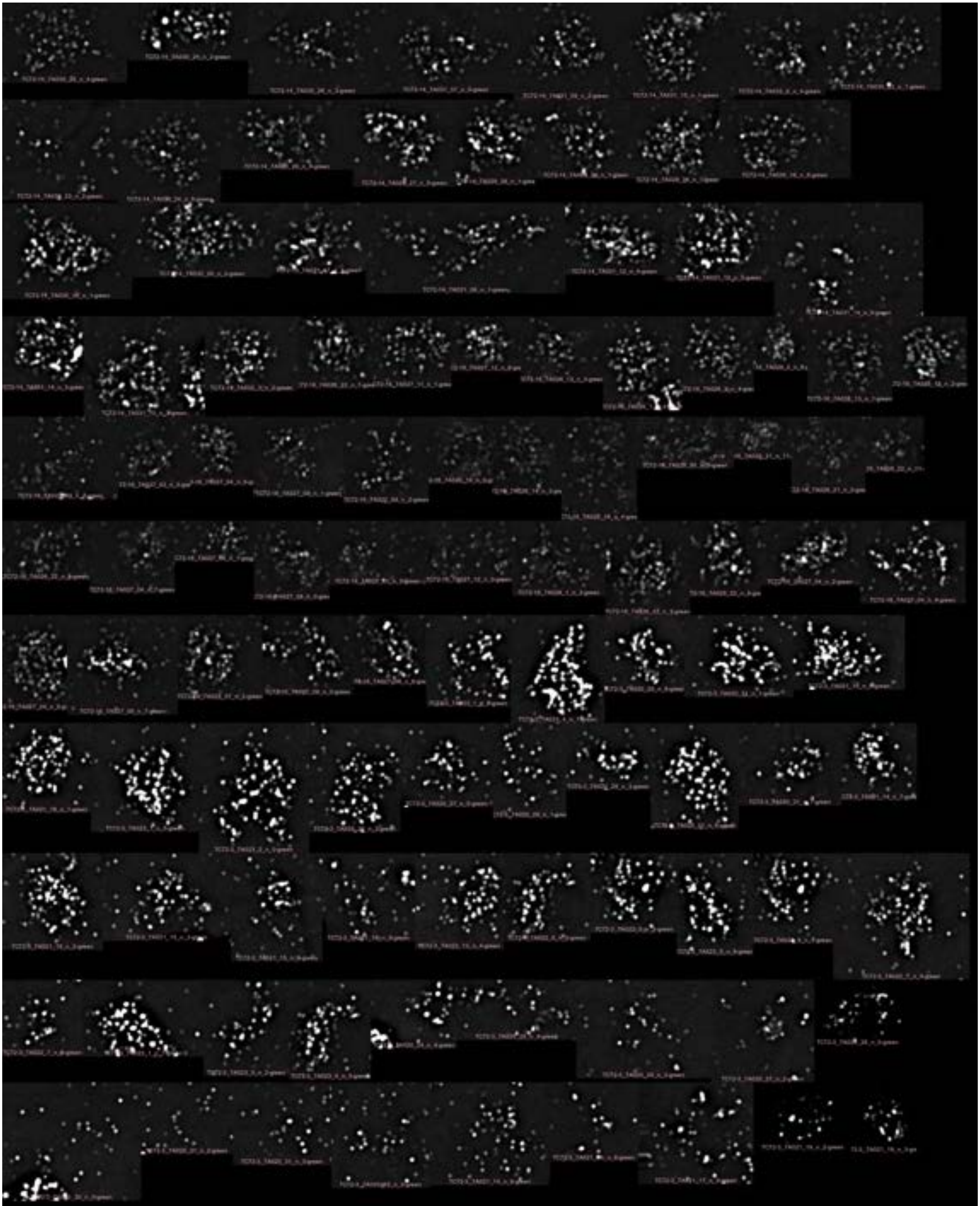

NONE-13

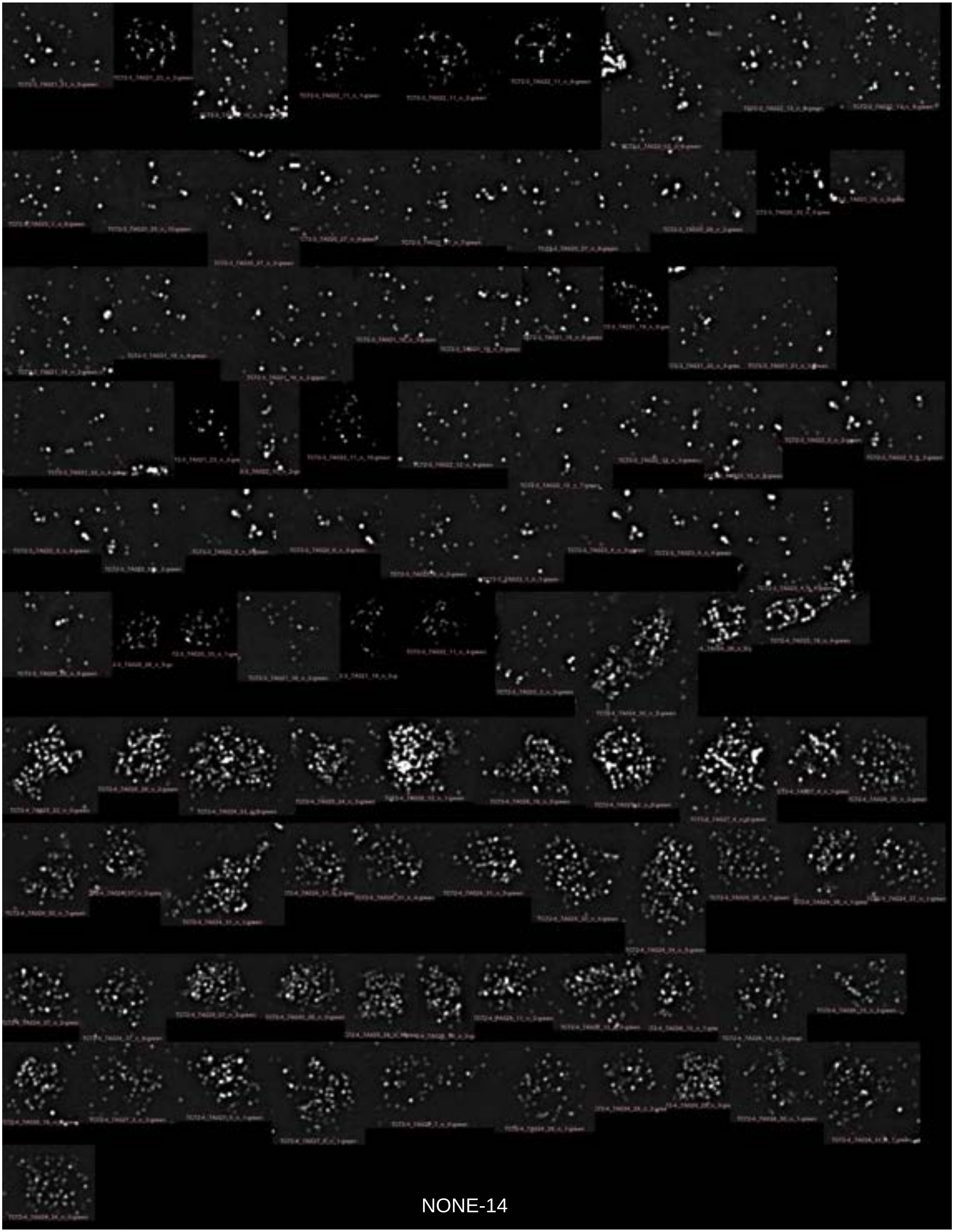

NONE-14

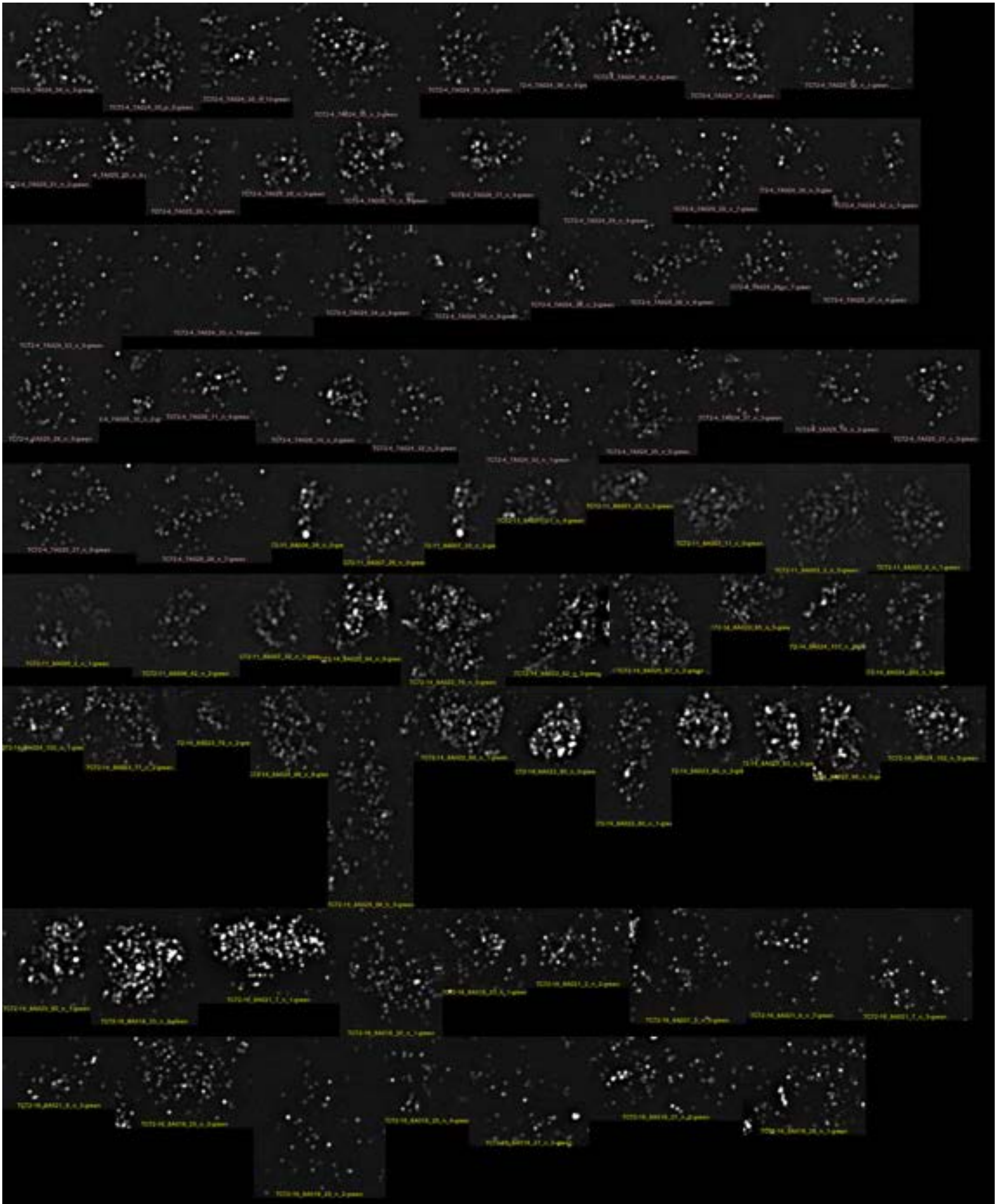

NONE-15

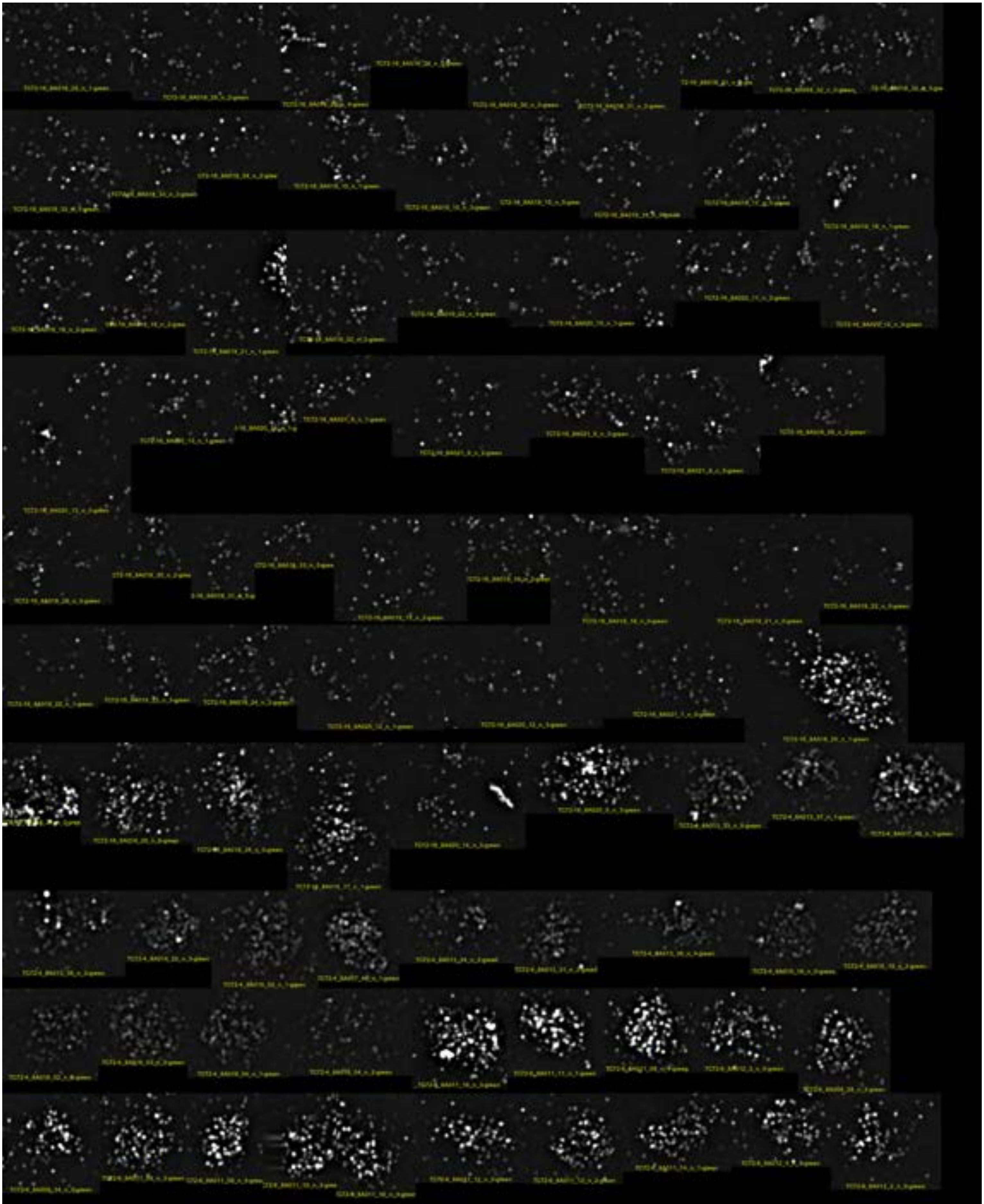

NONE-16

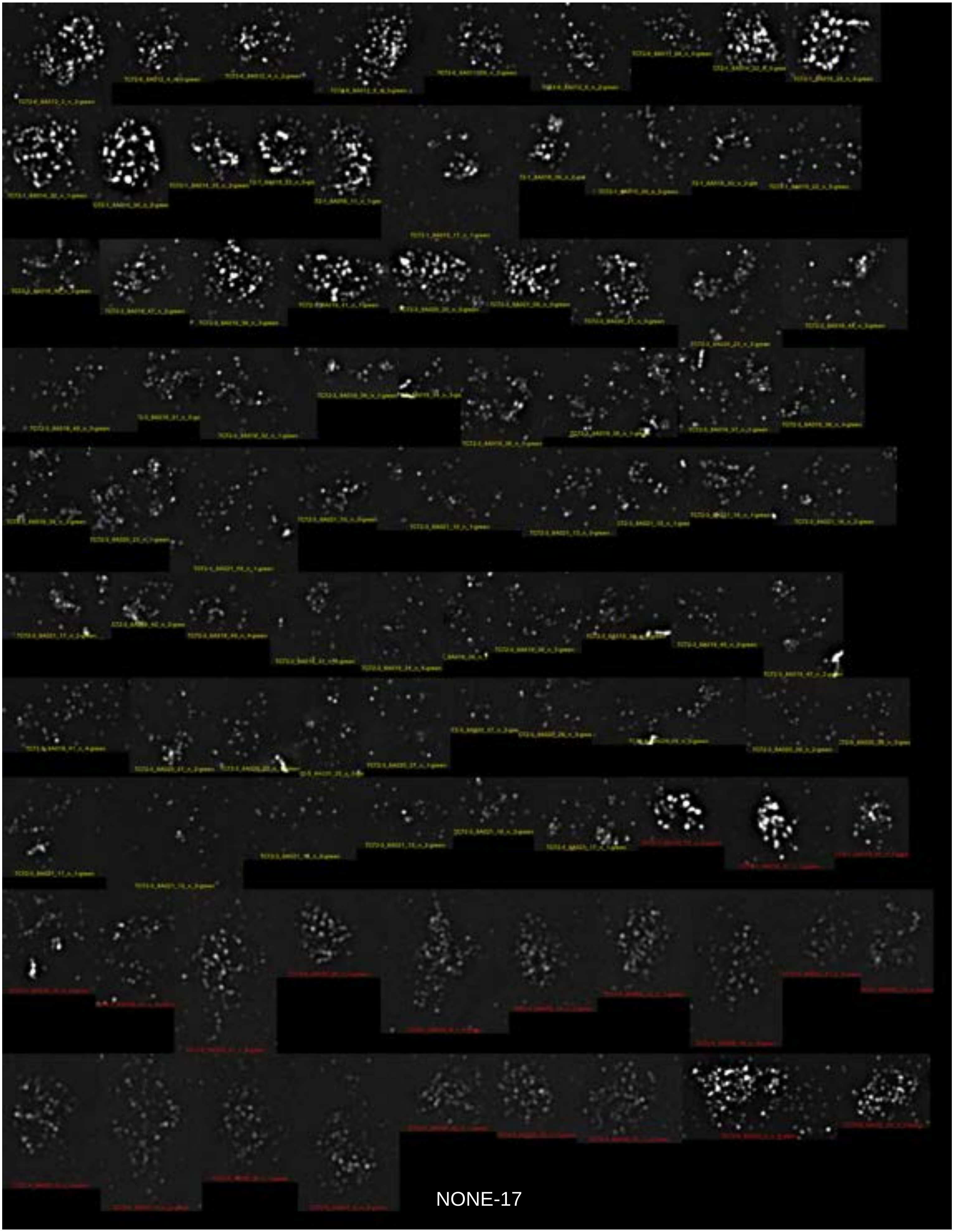

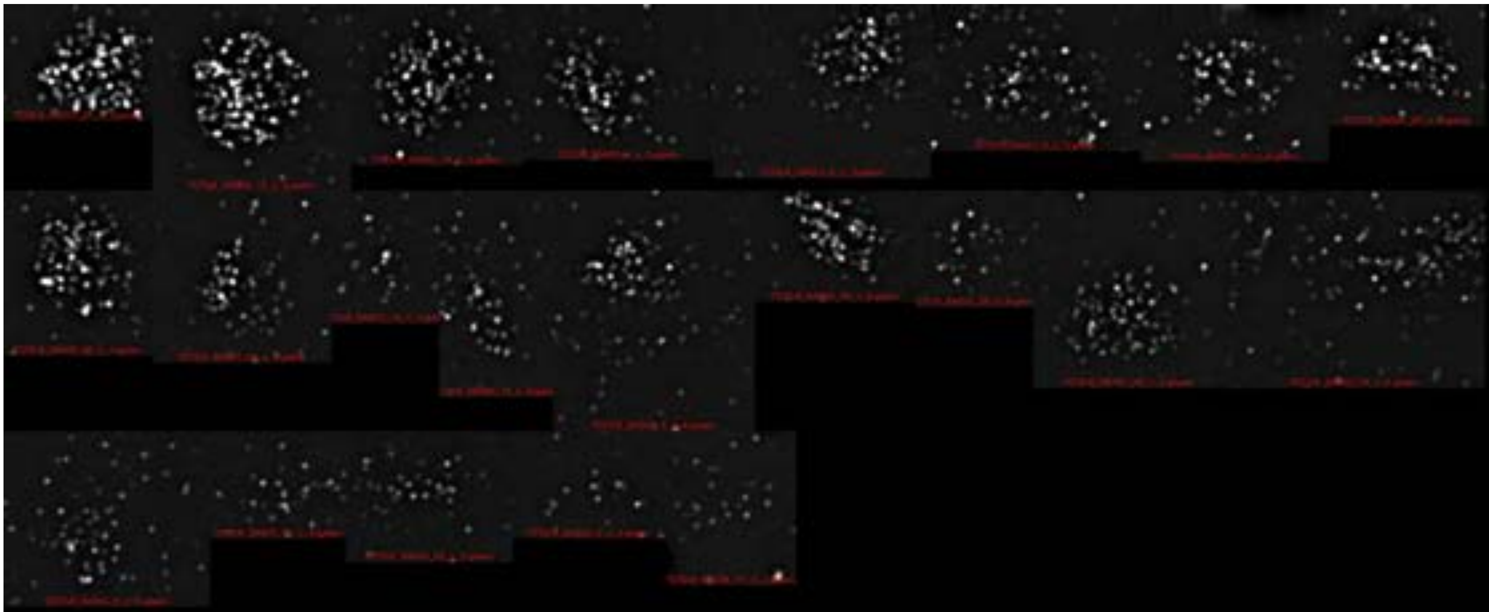

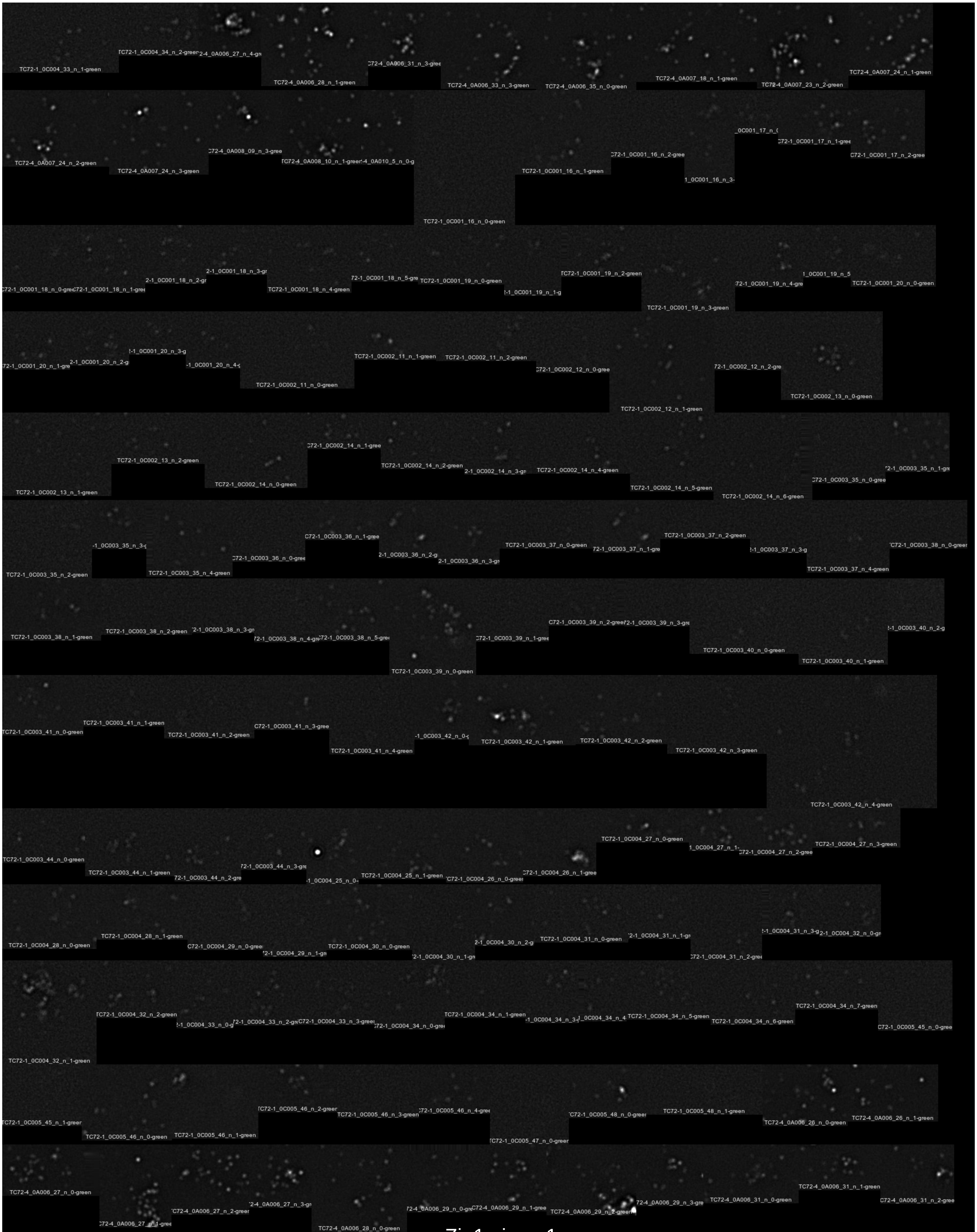
TC72-1_0C004_33_n_1-green
TC72-1_0C004_34_n_2-green
2-4_0A006_27_n_4-gre
TC72-4_0A006_28_n_1-green
C72-4_0A006_31_n_3-gree
TC72-4_0A006_33_n_3-green
TC72-4_0A006_35_n_0-green
TC72-4_0A007_18_n_1-green
TC72-4_0A007_23_n_2-green
TC72-4_0A007_24_n_1-green
TC72-4_0A007_24_n_2-green
TC72-4_0A007_24_n_3-green
C72-4_0A008_09_n_3-gree
TC72-4_0A008_10_n_1-green
-4_0A010_5_n_0-g
TC72-1_0C001_16_n_0-green
TC72-1_0C001_16_n_1-green
C72-1_0C001_16_n_2-gree
1_0C001_16_n_3-
_0C001_17_n_0
C72-1_0C001_17_n_1-gree
TC72-1_0C001_17_n_2-gree
C72-1_0C001_18_n_0-gree
C72-1_0C001_18_n_1-gree
2-1_0C001_18_n_2-gr
2-1_0C001_18_n_3-gr
TC72-1_0C001_18_n_4-green
72-1_0C001_18_n_5-gre
TC72-1_0C001_19_n_0-green
2-1_0C001_19_n_1-g
TC72-1_0C001_19_n_2-green
TC72-1_0C001_19_n_3-green
72-1_0C001_19_n_4-gre
1_0C001_19_n_5
TC72-1_0C001_20_n_0-green
72-1_0C001_20_n_1-gre
2-1_0C001_20_n_2-g
2-1_0C001_20_n_3-g
-1_0C001_20_n_4-g
TC72-1_0C002_11_n_0-green
TC72-1_0C002_11_n_1-green
TC72-1_0C002_11_n_2-green
C72-1_0C002_12_n_0-gree
TC72-1_0C002_12_n_1-green
72-1_0C002_12_n_2-gre
TC72-1_0C002_13_n_0-green
TC72-1_0C002_13_n_1-green
TC72-1_0C002_13_n_2-green
TC72-1_0C002_14_n_0-green
C72-1_0C002_14_n_1-gree
TC72-1_0C002_14_n_2-green
2-1_0C002_14_n_3-gr
TC72-1_0C002_14_n_4-green
TC72-1_0C002_14_n_5-green
TC72-1_0C002_14_n_6-green
C72-1_0C003_35_n_0-gree
2-1_0C003_35_n_1-gre
TC72-1_0C003_35_n_2-green
-1_0C003_35_n_3-g
TC72-1_0C003_35_n_4-green
C72-1_0C003_36_n_0-gree
C72-1_0C003_36_n_1-gree
2-1_0C003_36_n_2-g
2-1_0C003_36_n_3-gr
TC72-1_0C003_37_n_0-green
72-1_0C003_37_n_1-gre
TC72-1_0C003_37_n_2-green
2-1_0C003_37_n_3-g
TC72-1_0C003_37_n_4-green
C72-1_0C003_38_n_0-green
TC72-1_0C003_38_n_1-green
TC72-1_0C003_38_n_2-green
2-1_0C003_38_n_3-gr
2-1_0C003_38_n_4-gr
C72-1_0C003_38_n_5-gre
TC72-1_0C003_39_n_0-green
C72-1_0C003_39_n_1-gree
C72-1_0C003_39_n_2-gree
72-1_0C003_39_n_3-gre
TC72-1_0C003_40_n_0-green
TC72-1_0C003_40_n_1-green
2-1_0C003_40_n_2-g
C72-1_0C003_41_n_0-green
TC72-1_0C003_41_n_1-green
TC72-1_0C003_41_n_2-green
C72-1_0C003_41_n_3-gree
TC72-1_0C003_41_n_4-green
-1_0C003_42_n_0-g
TC72-1_0C003_42_n_1-green
TC72-1_0C003_42_n_2-green
TC72-1_0C003_42_n_3-green
TC72-1_0C003_42_n_4-green
TC72-1_0C003_44_n_0-green
TC72-1_0C003_44_n_1-green
72-1_0C003_44_n_2-gre
2-1_0C003_44_n_3-gre
-1_0C004_25_n_0-
TC72-1_0C004_25_n_1-green
C72-1_0C004_26_n_0-gree
C72-1_0C004_26_n_1-gree
TC72-1_0C004_27_n_0-green
1_0C004_27_n_1-
C72-1_0C004_27_n_2-green
TC72-1_0C004_27_n_3-green
TC72-1_0C004_28_n_0-green
TC72-1_0C004_28_n_1-green
C72-1_0C004_29_n_0-gree
2-1_0C004_29_n_1-gr
TC72-1_0C004_30_n_0-green
2-1_0C004_30_n_1-gr
2-1_0C004_30_n_2-g
TC72-1_0C004_31_n_0-green
2-1_0C004_31_n_1-gr
C72-1_0C004_31_n_2-gree
2-1_0C004_31_n_3-g
2-1_0C004_32_n_0-gr
TC72-1_0C004_32_n_1-green
TC72-1_0C004_32_n_2-green
2-1_0C004_33_n_0-g
2-1_0C004_33_n_2-gr
C72-1_0C004_33_n_3-gree
72-1_0C004_34_n_0-gre
TC72-1_0C004_34_n_1-green
-1_0C004_34_n_3-
_0C004_34_n_4
TC72-1_0C004_34_n_5-green
TC72-1_0C004_34_n_6-green
TC72-1_0C004_34_n_7-green
C72-1_0C005_45_n_0-gree
C72-1_0C005_45_n_1-green
TC72-1_0C005_46_n_0-green
TC72-1_0C005_46_n_1-green
C72-1_0C005_46_n_2-green
TC72-1_0C005_46_n_3-green
C72-1_0C005_46_n_4-gree
C72-1_0C005_47_n_0-green
C72-1_0C005_48_n_0-green
TC72-1_0C005_48_n_1-green
TC72-4_0A006_26_n_0-green
TC72-4_0A006_26_n_1-green
TC72-4_0A006_27_n_0-green
C72-4_0A006_27_n_1-gree
C72-4_0A006_27_n_2-green
2-4_0A006_27_n_3-gr
TC72-4_0A006_28_n_0-green
2-4_0A006_29_n_0-gre
C72-4_0A006_29_n_1-gree
TC72-4_0A006_29_n_2-green
2-4_0A006_29_n_3-gre
TC72-4_0A006_31_n_0-green
TC72-4_0A006_31_n_1-green
C72-4_0A006_31_n_2-gree

Zip1minus-1

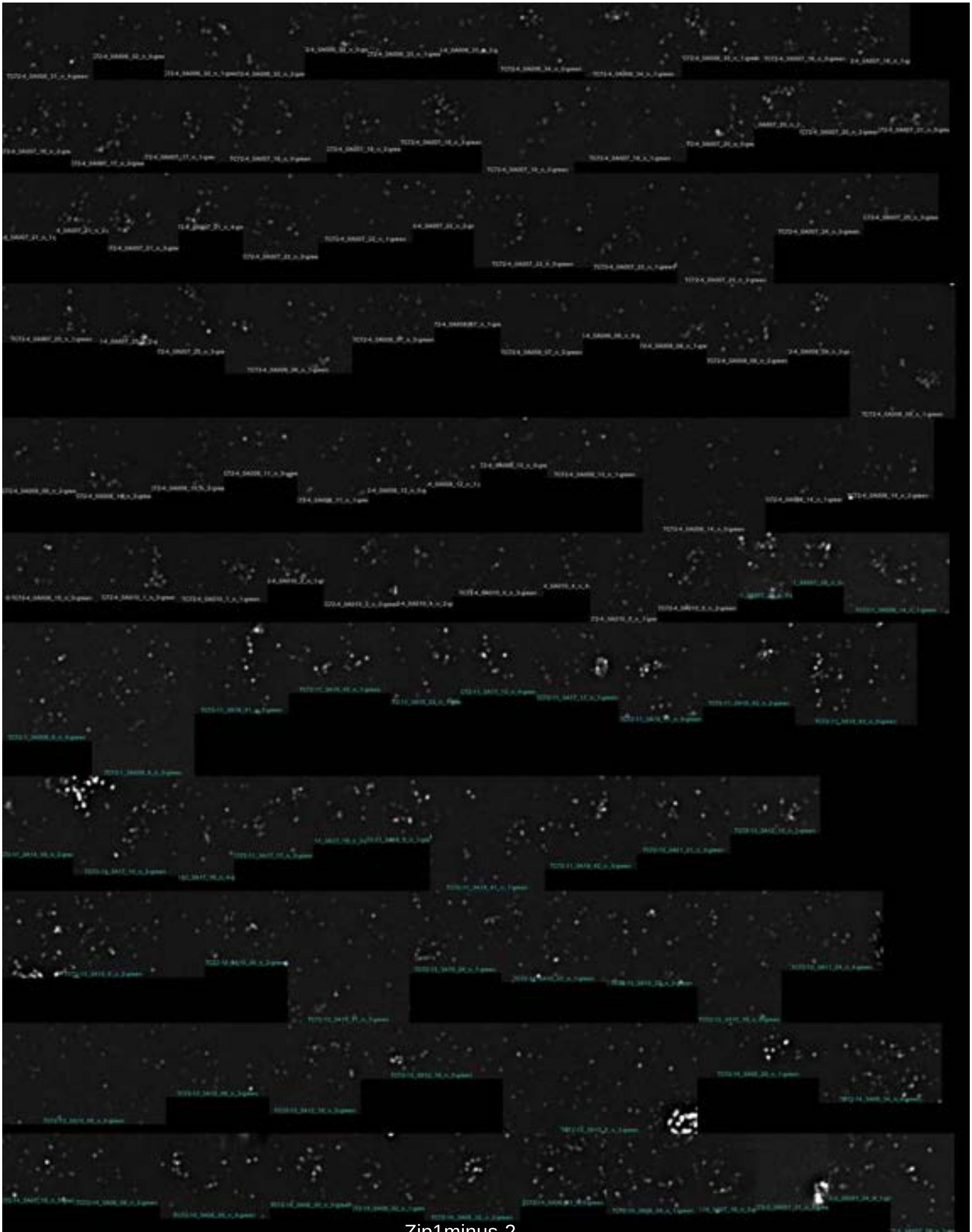

Zip1minus-2

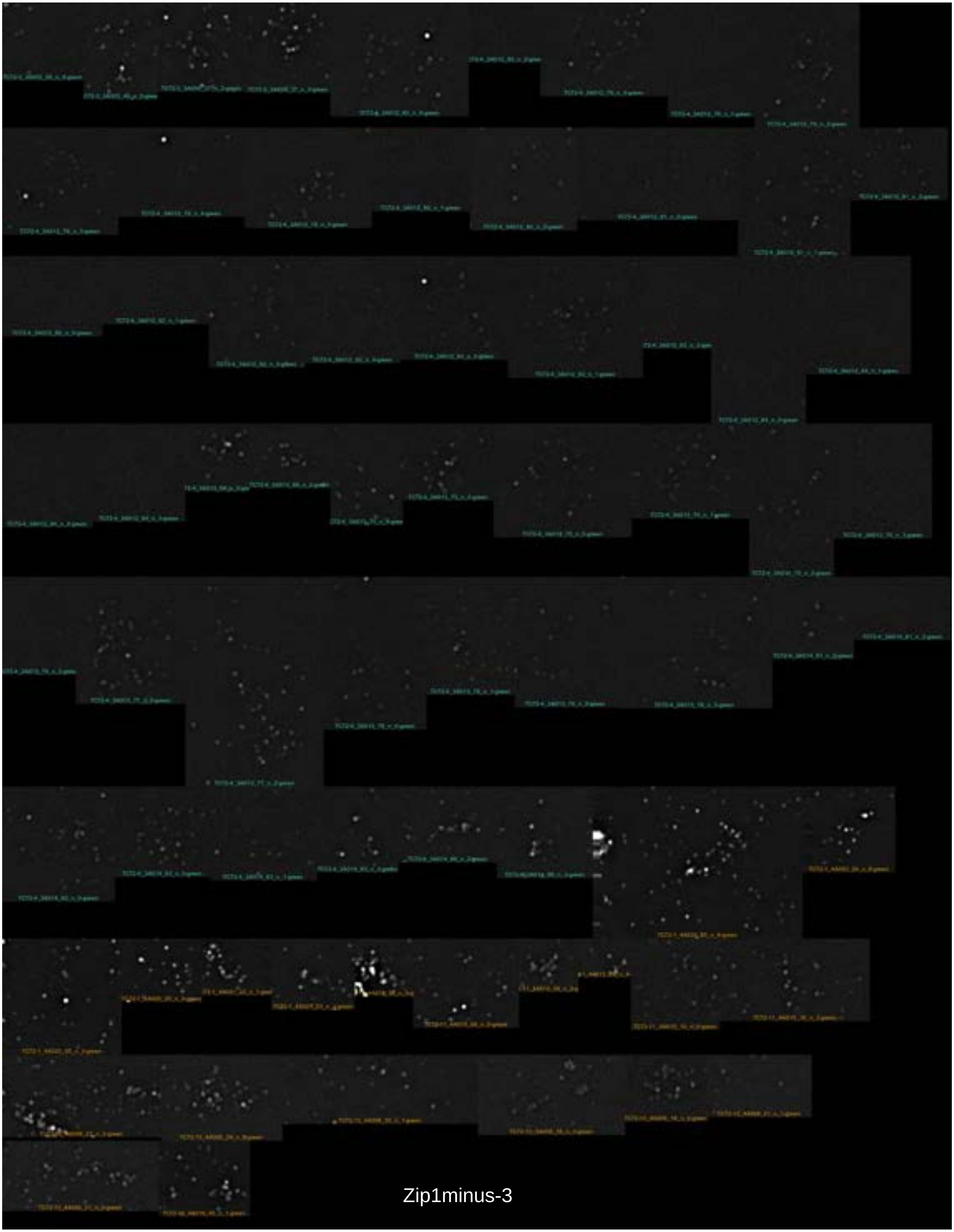

Zip1minus-3

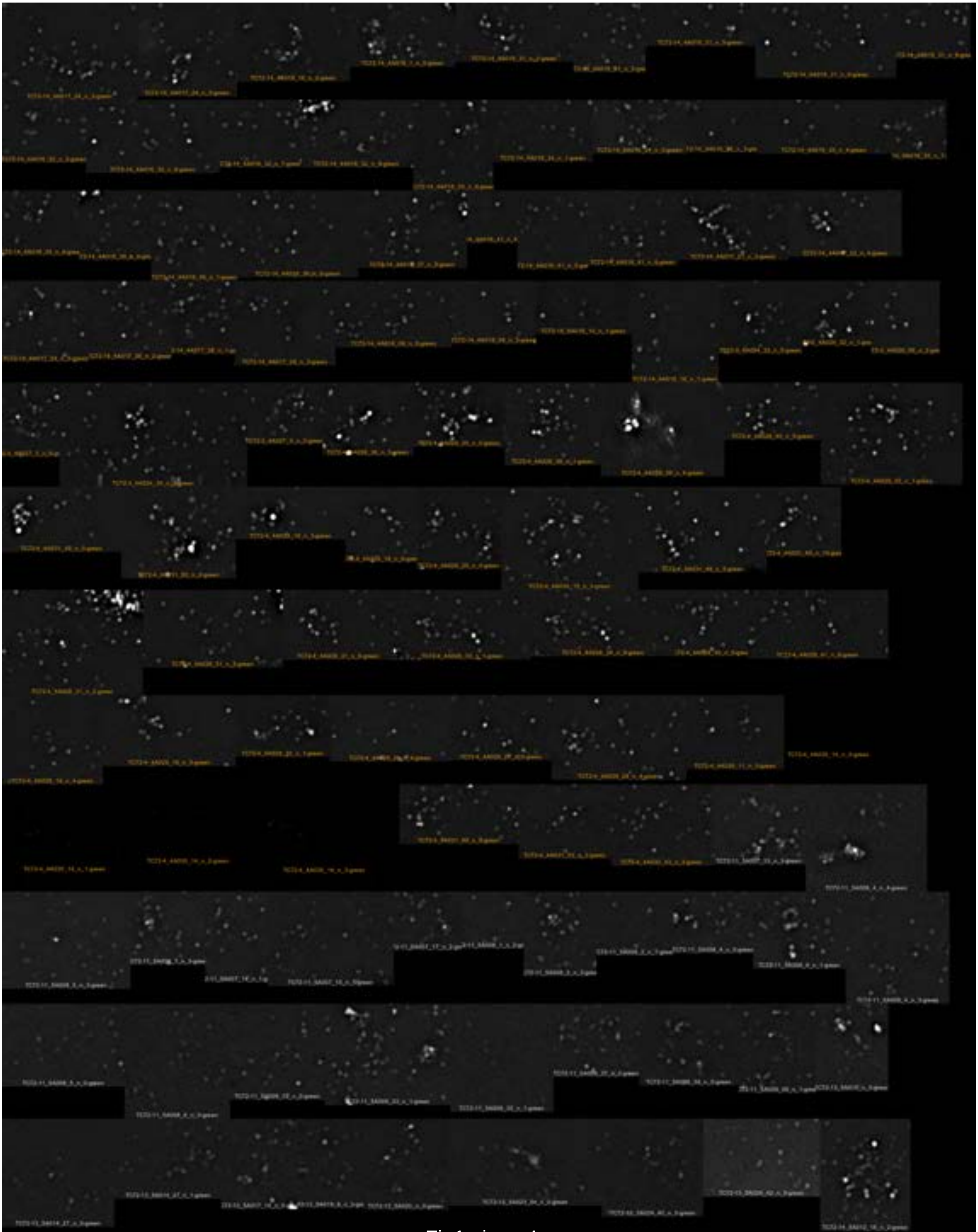

Zip1minus-4

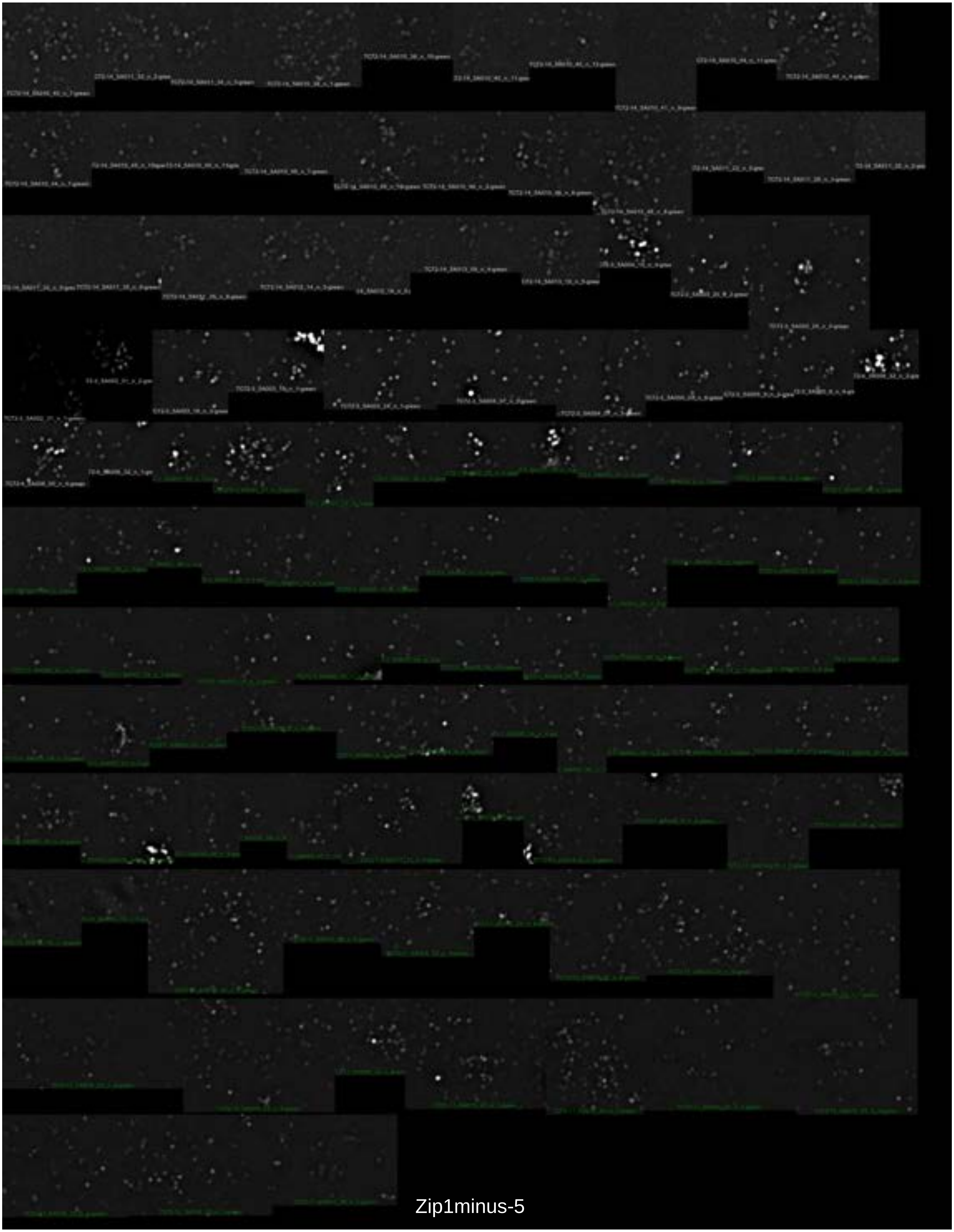

Zip1minus-5

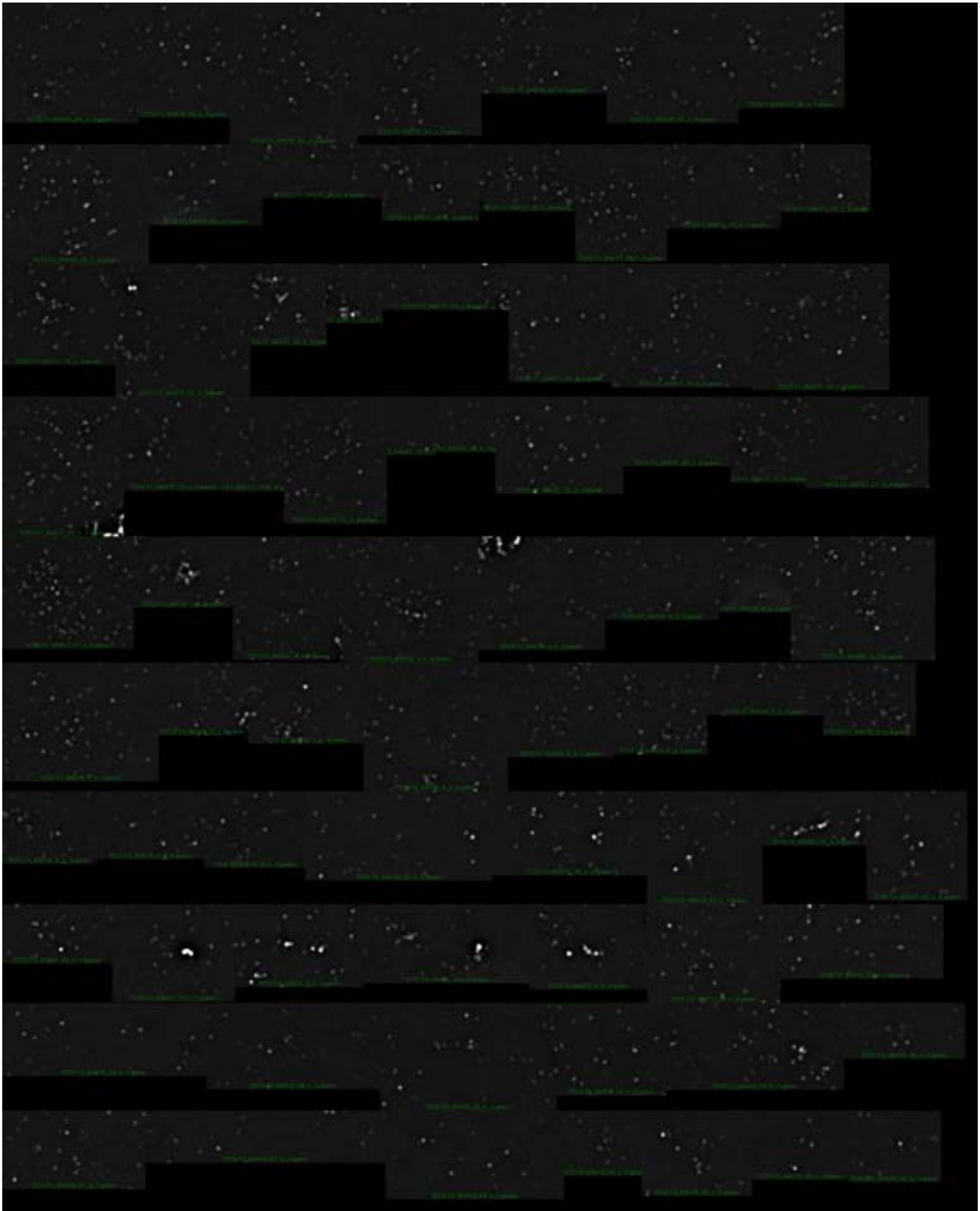

Zip1minus-6

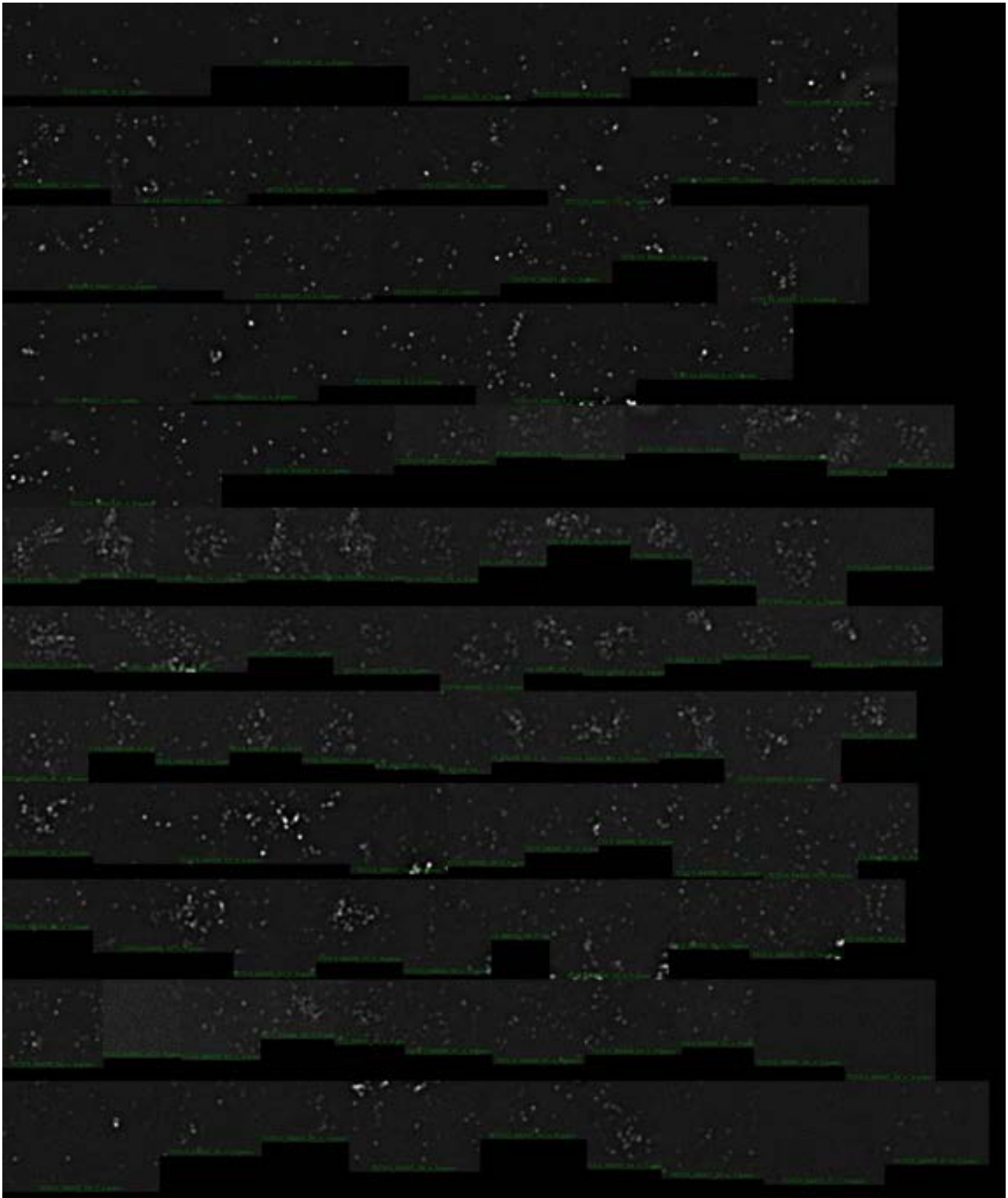

Zip1minus-7

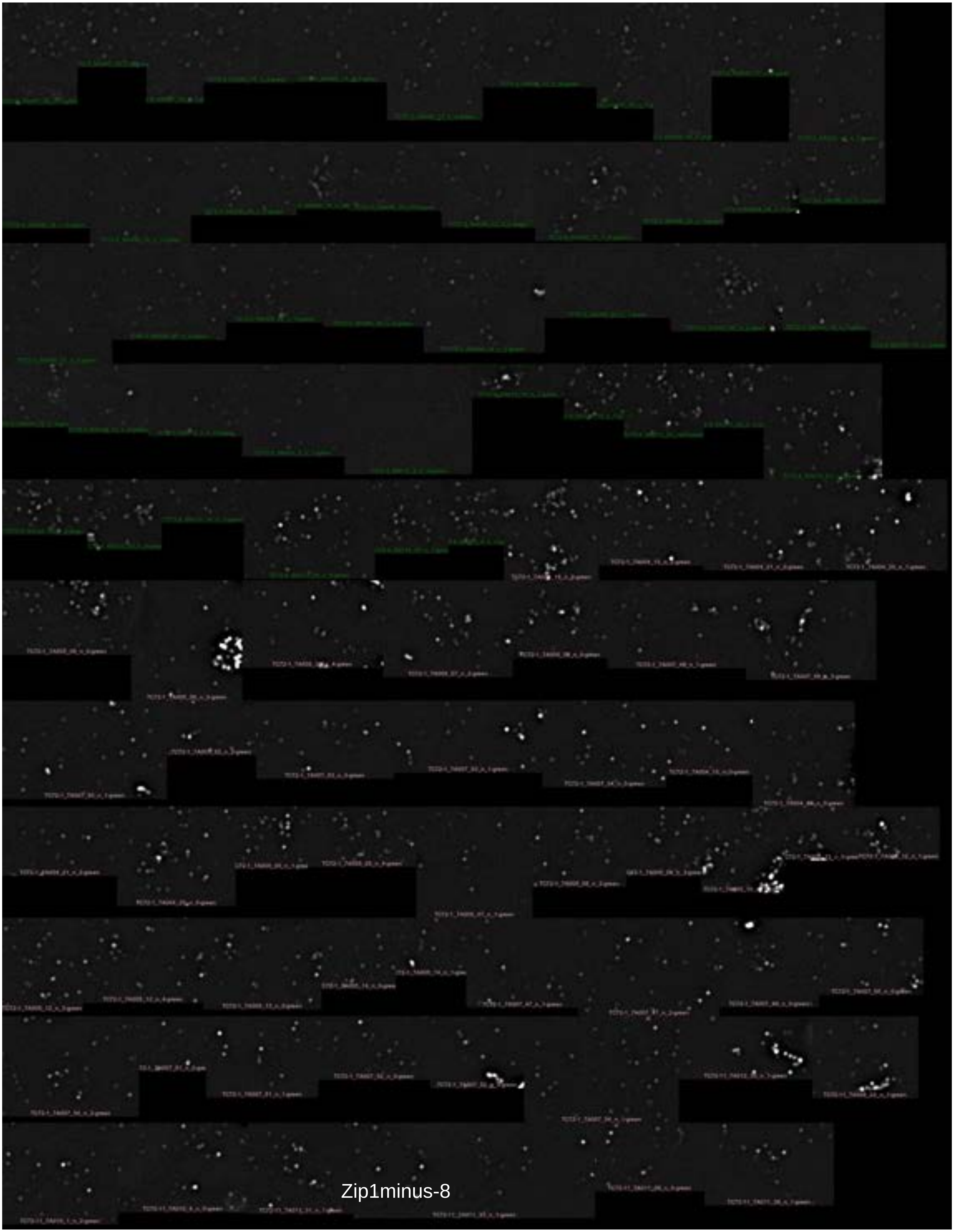

Zip1minus-8

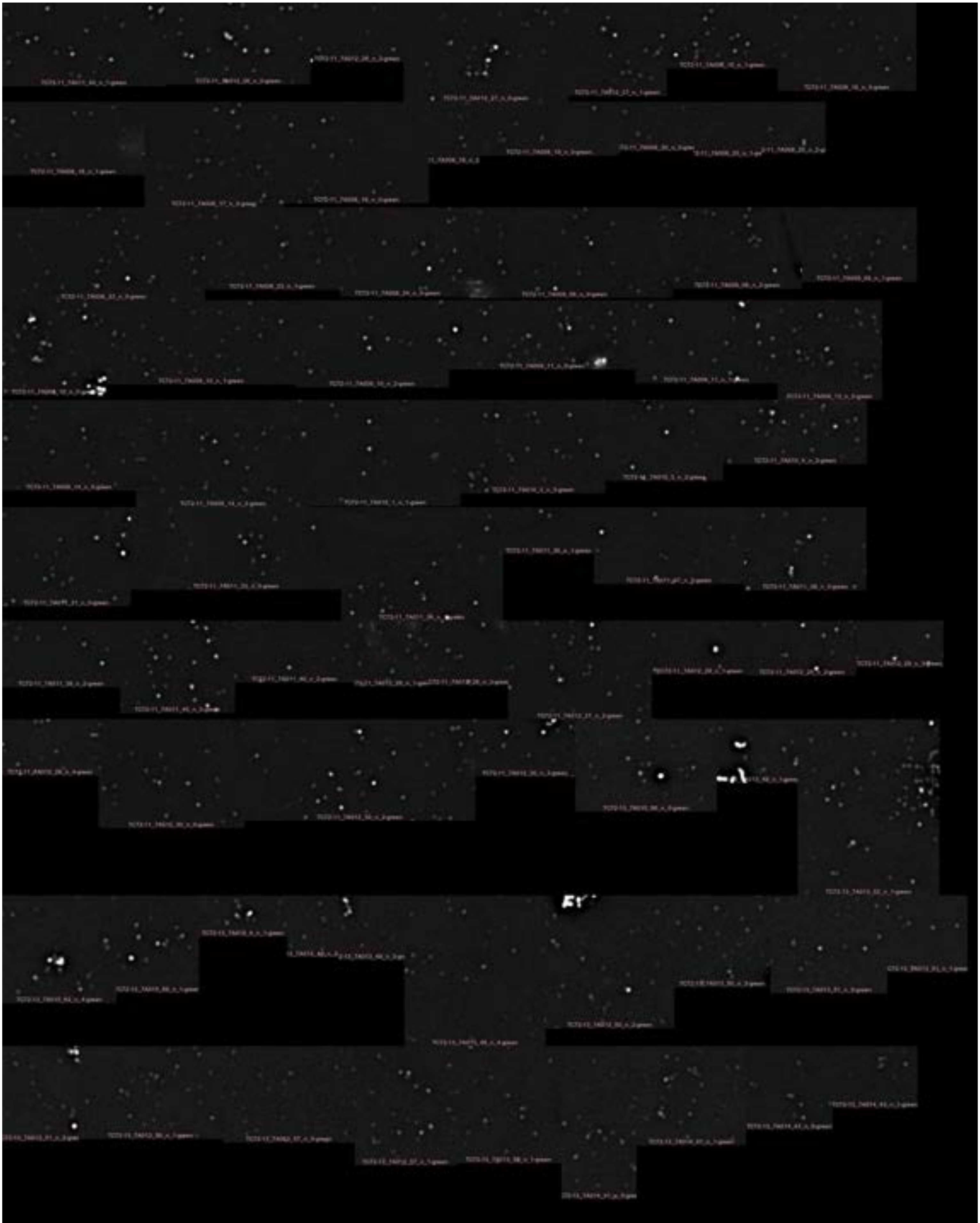

Zip1minus-9

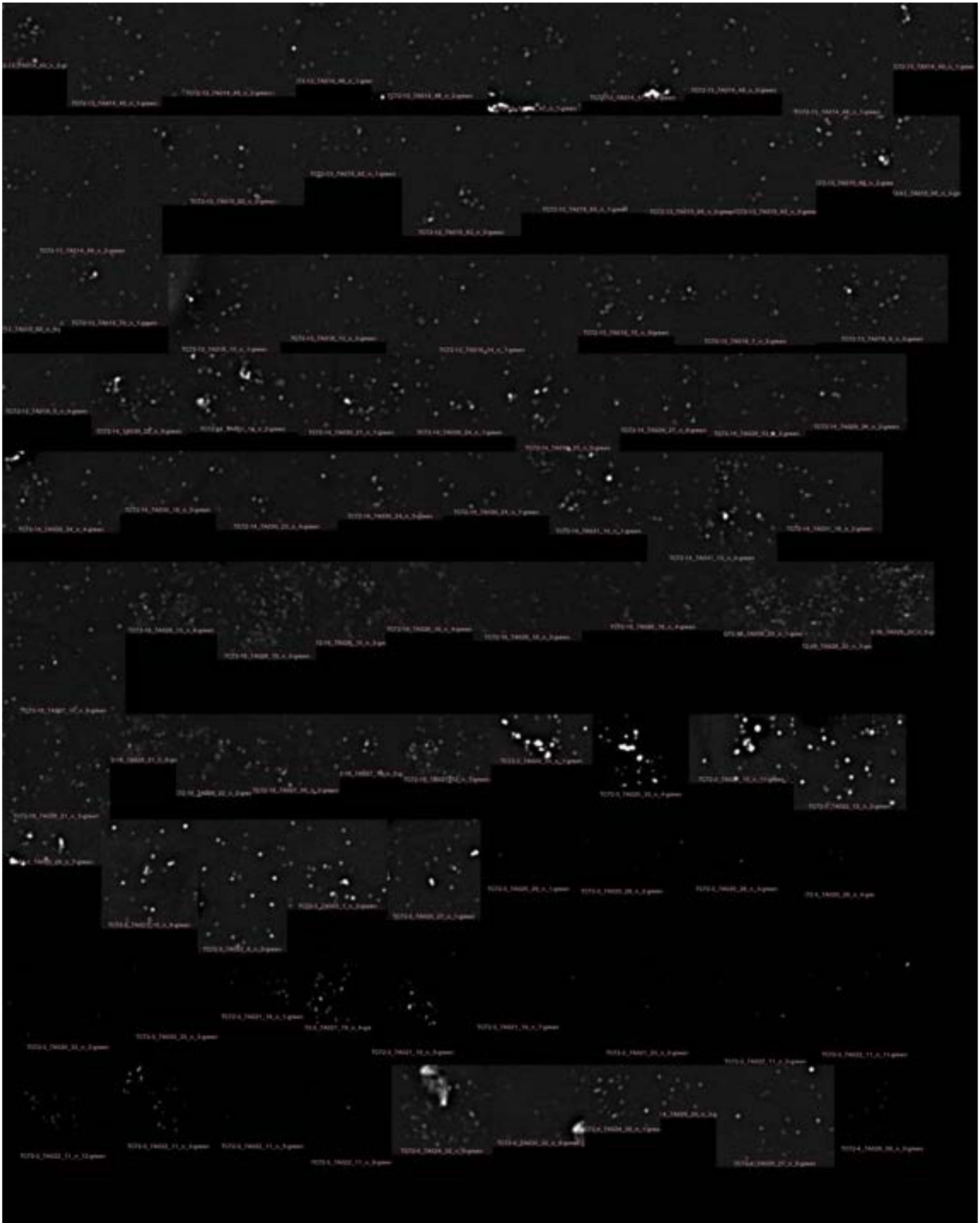

Zip1minus-10

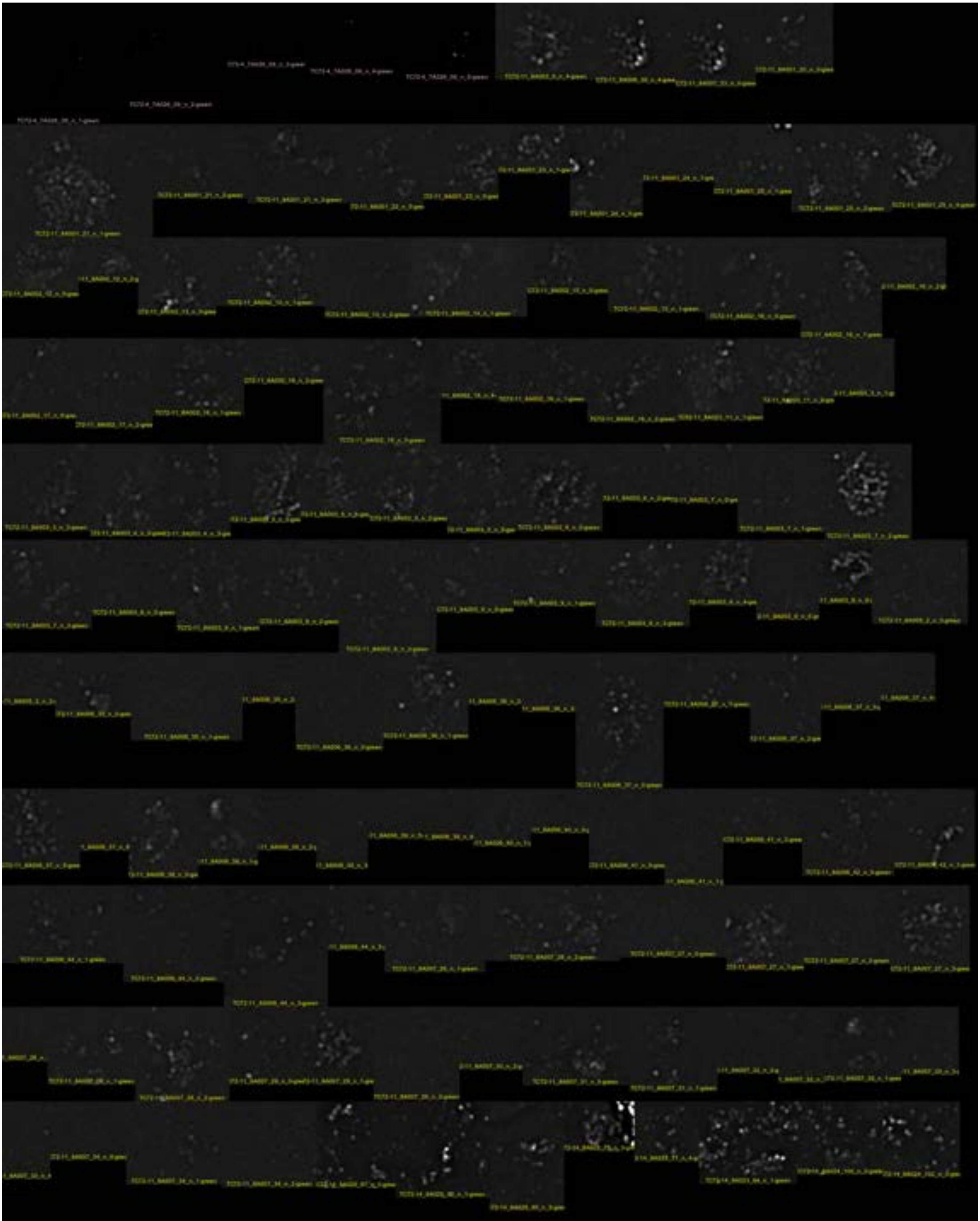

Zip1minus-11

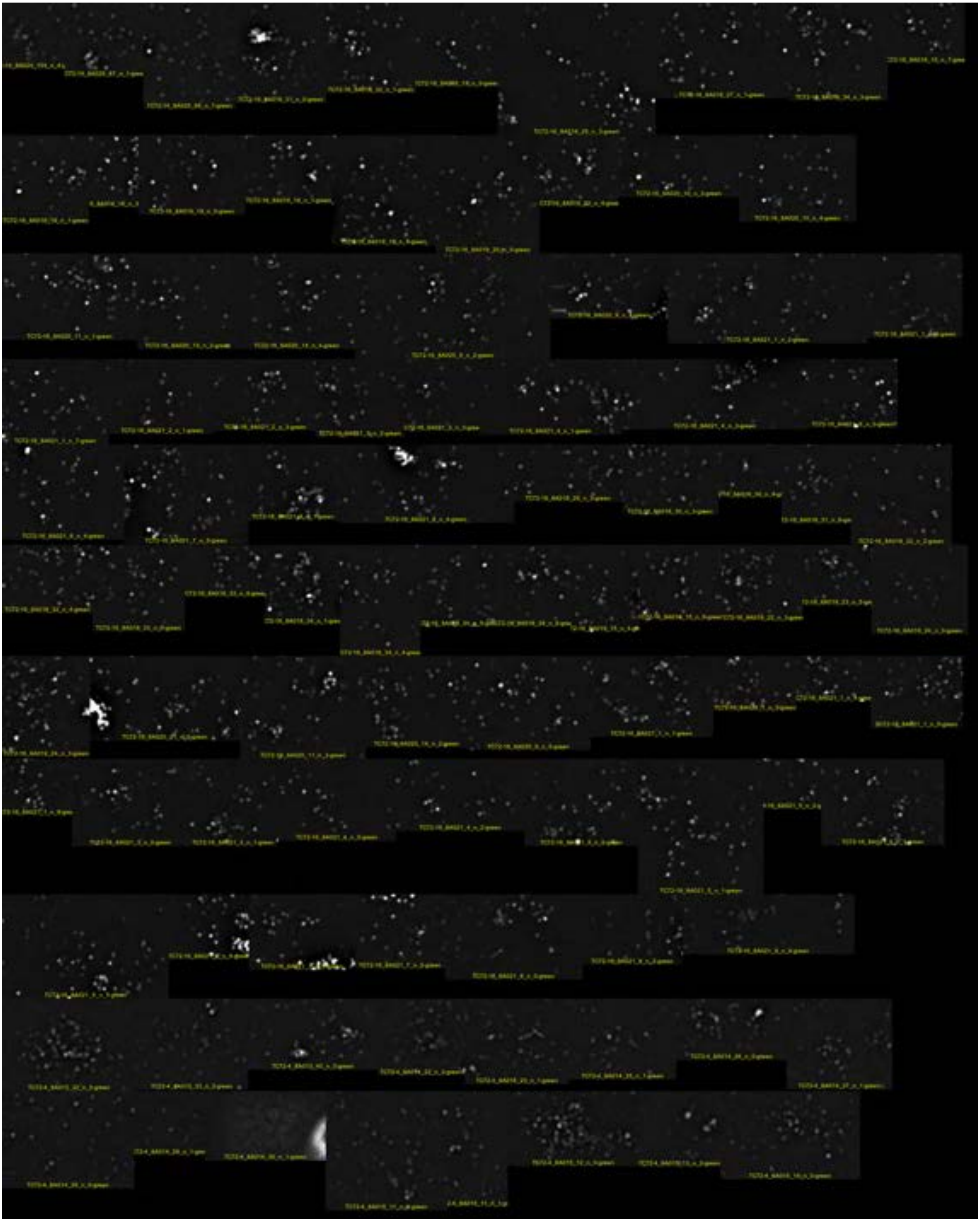

Zip1minus-12

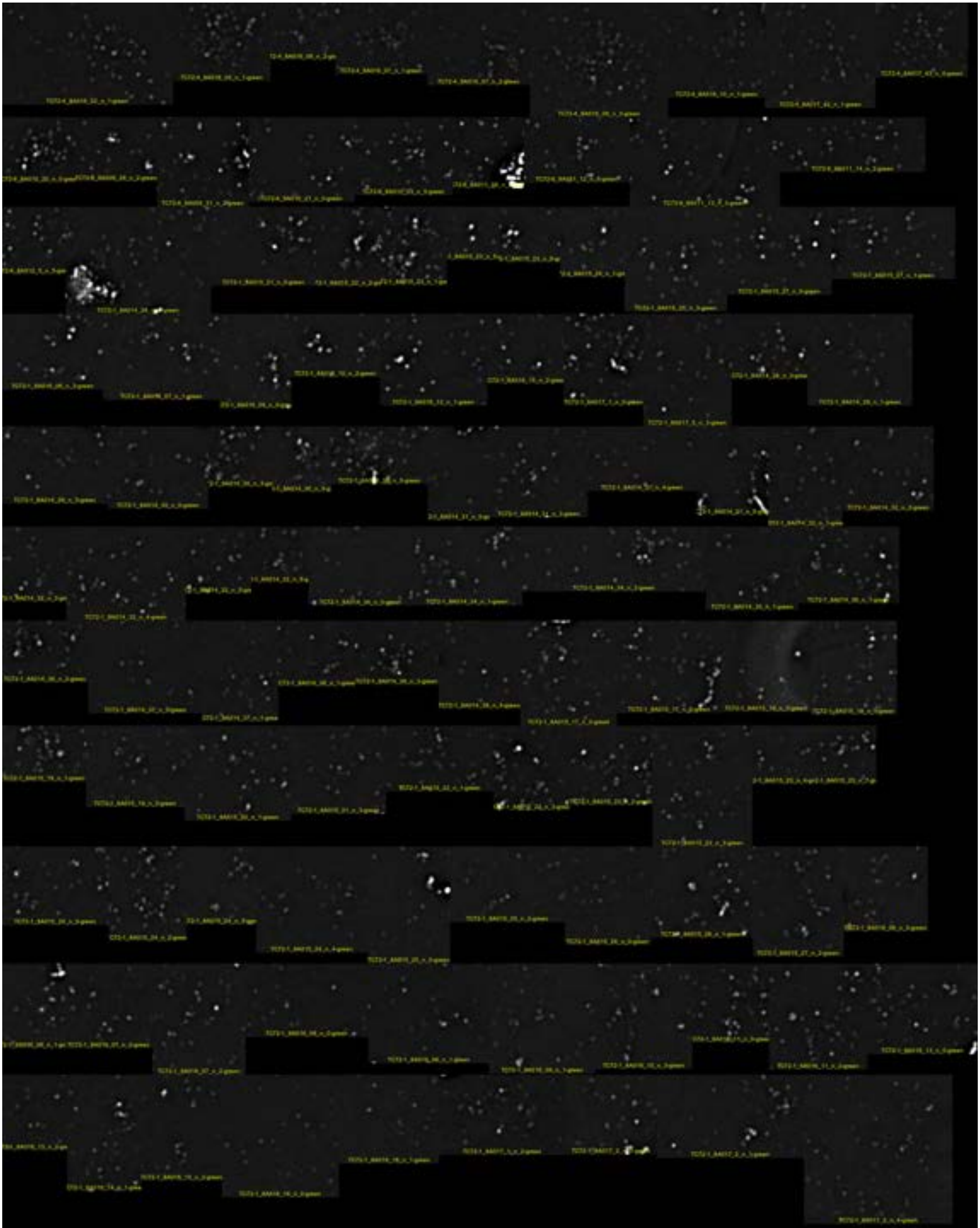

Zip1minus-13

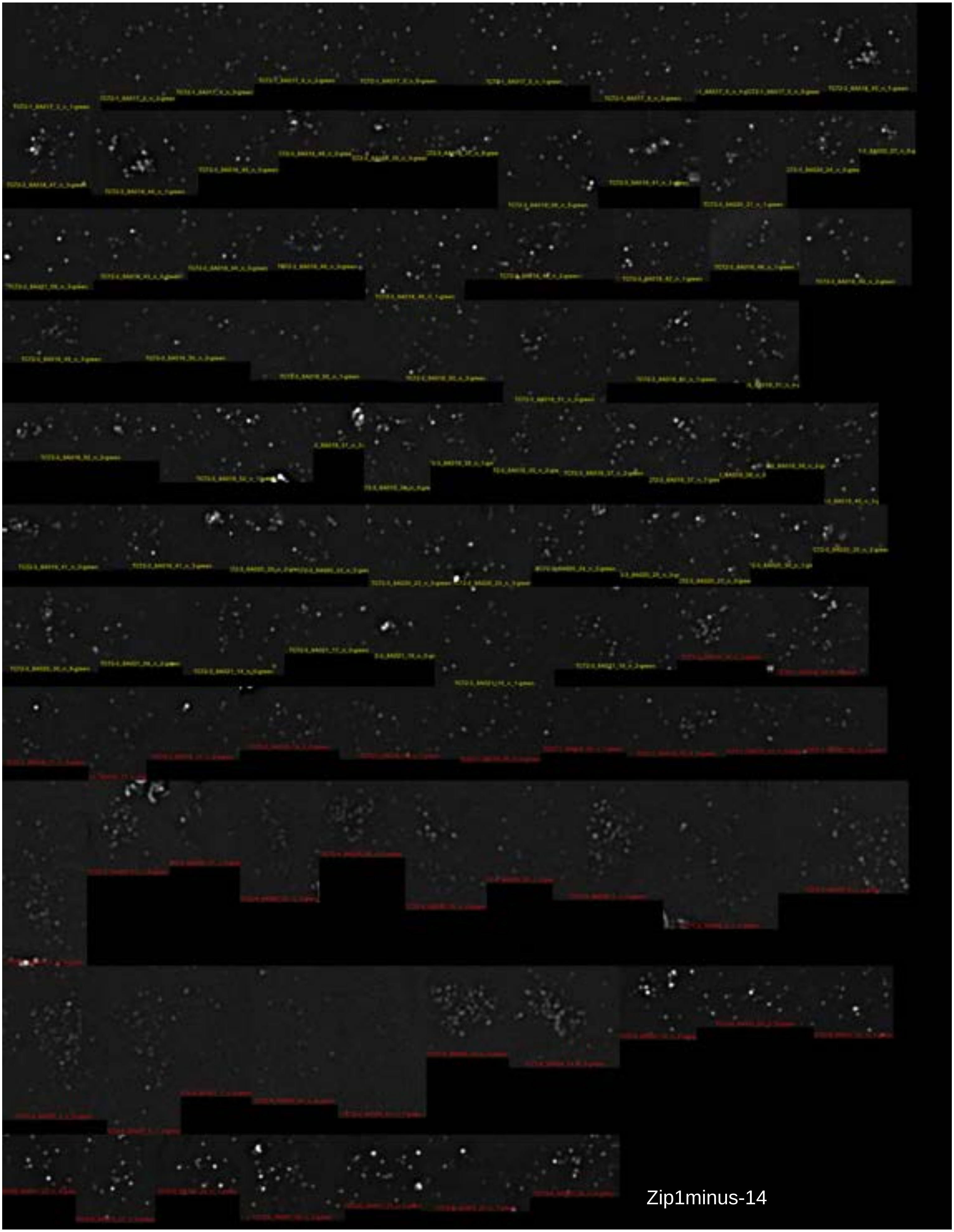
Zip1minus-14

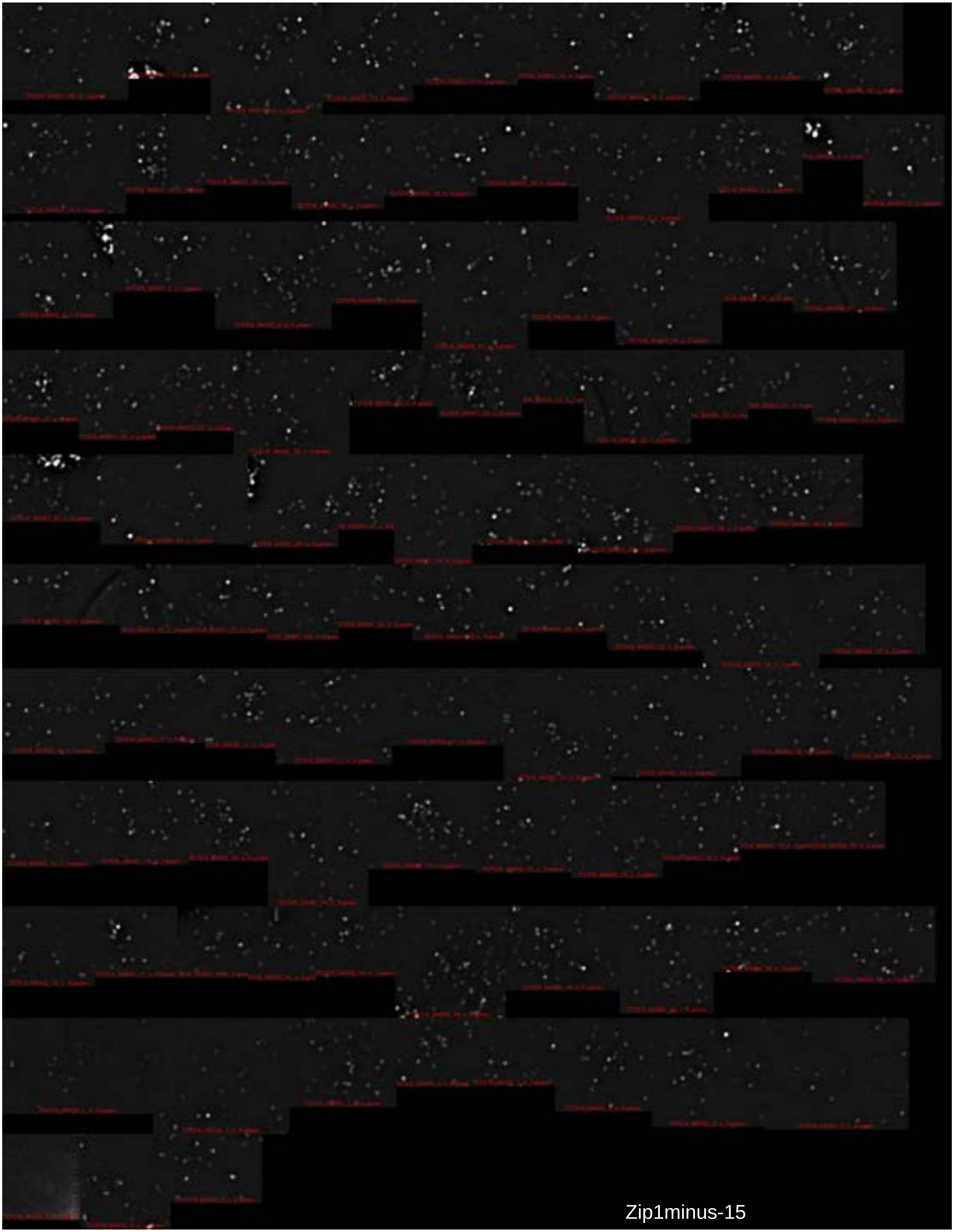

Zip1minus-15

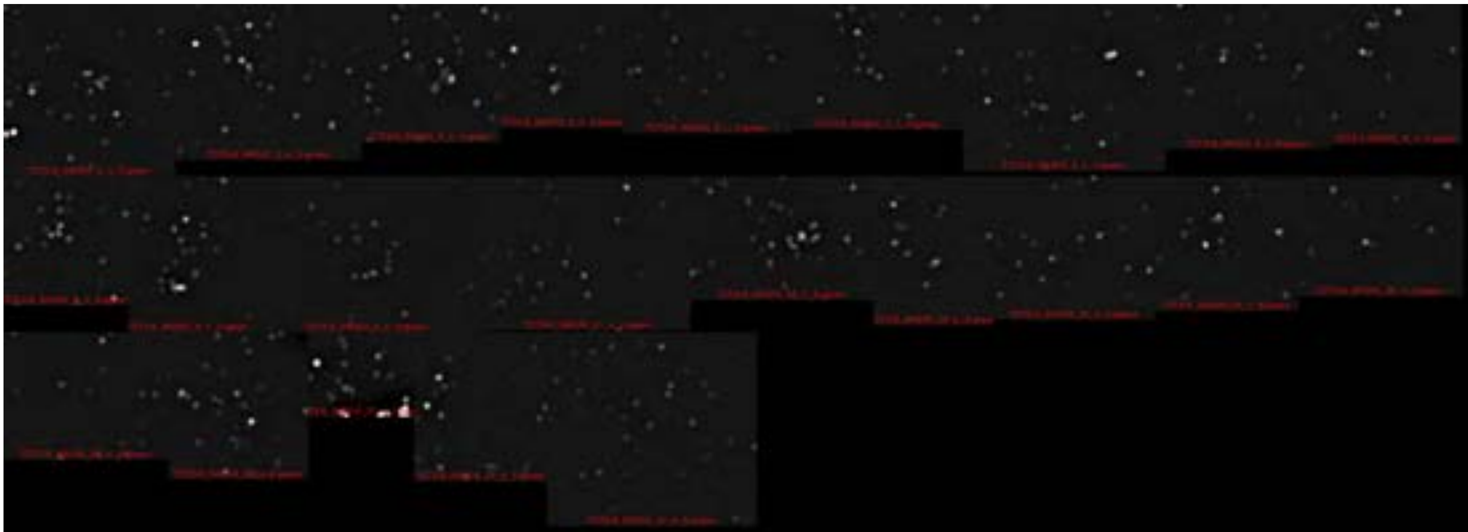

Zip1minus-16

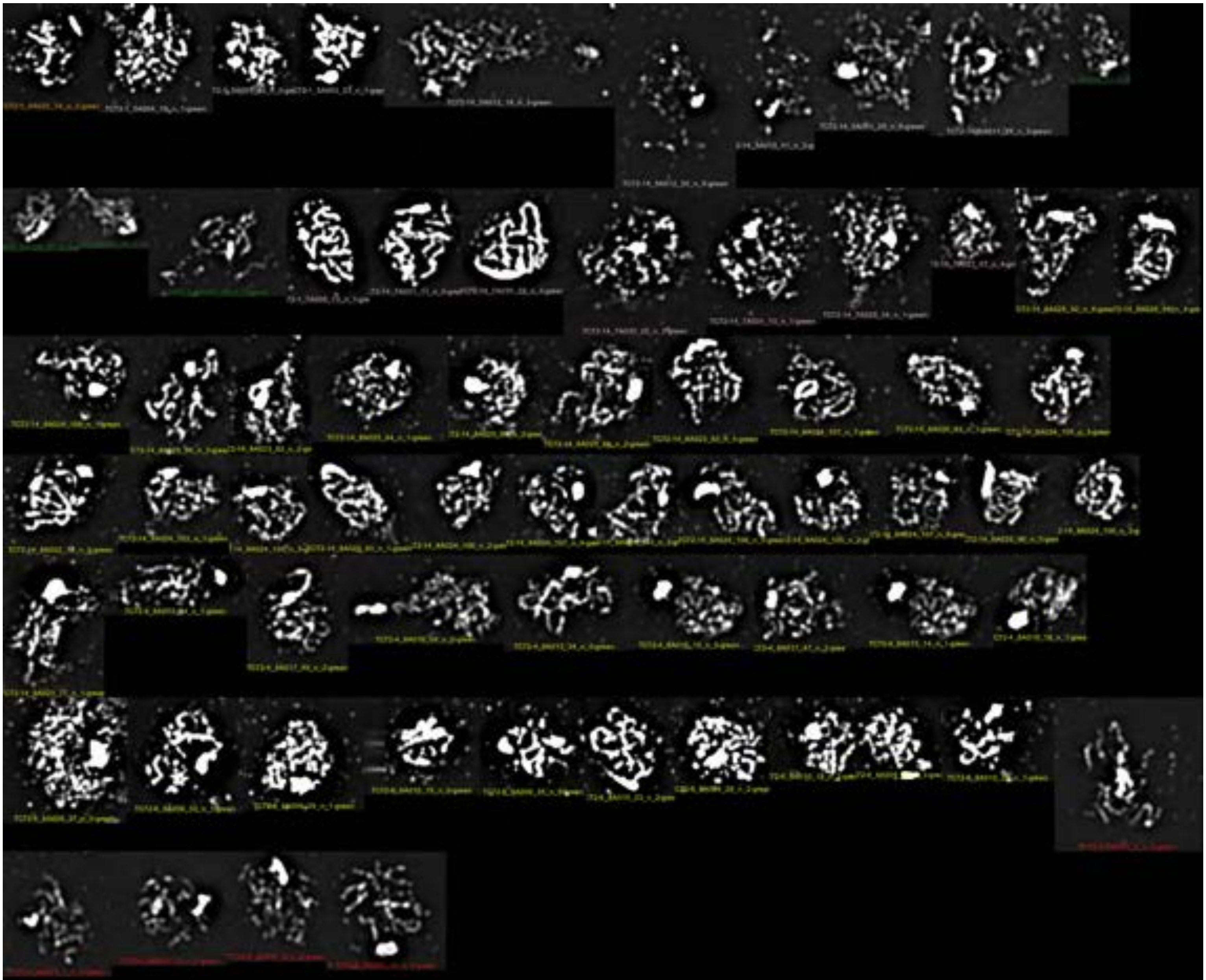

PC+CONTINUOUS-1

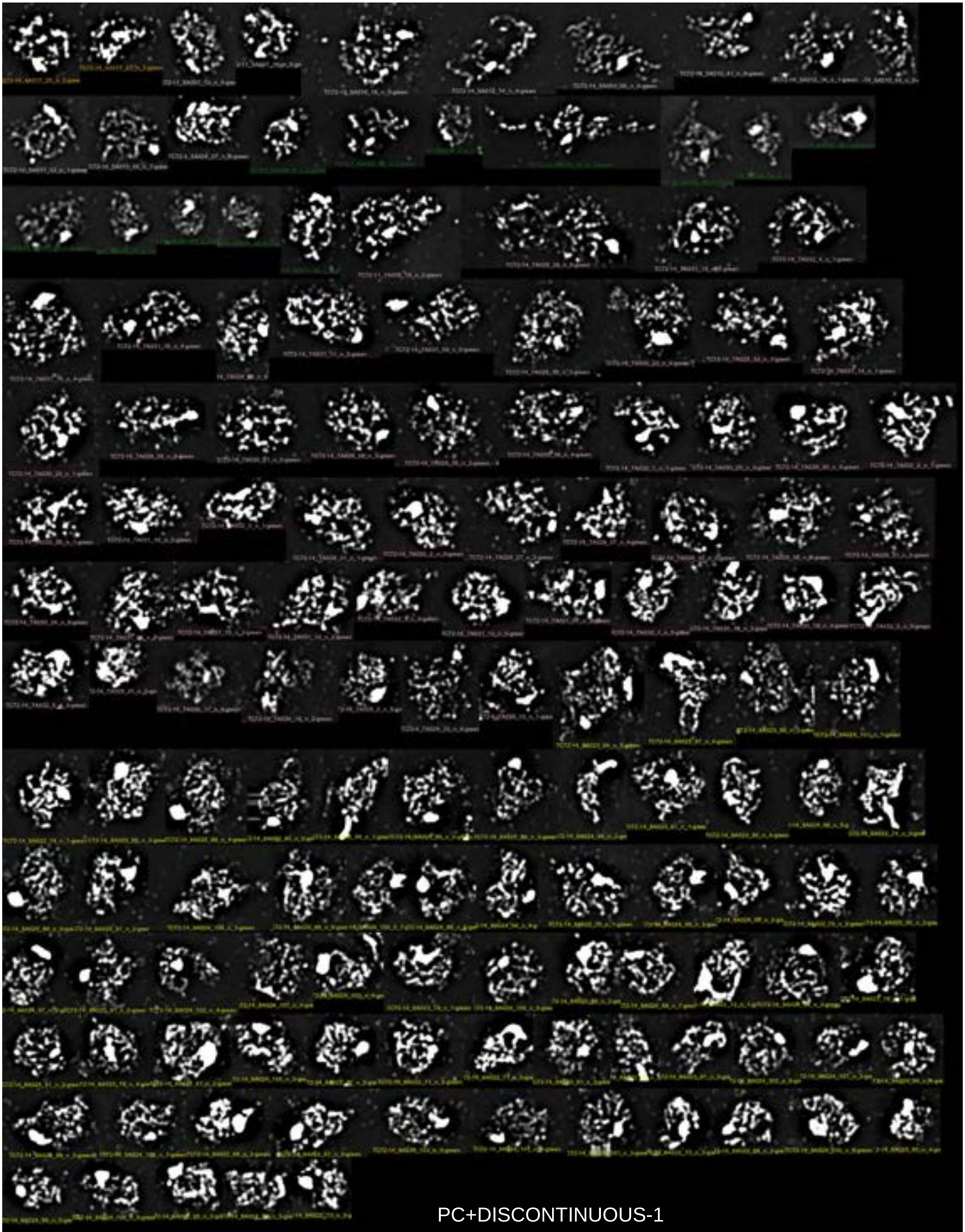

PC+DISCONTINUOUS-1

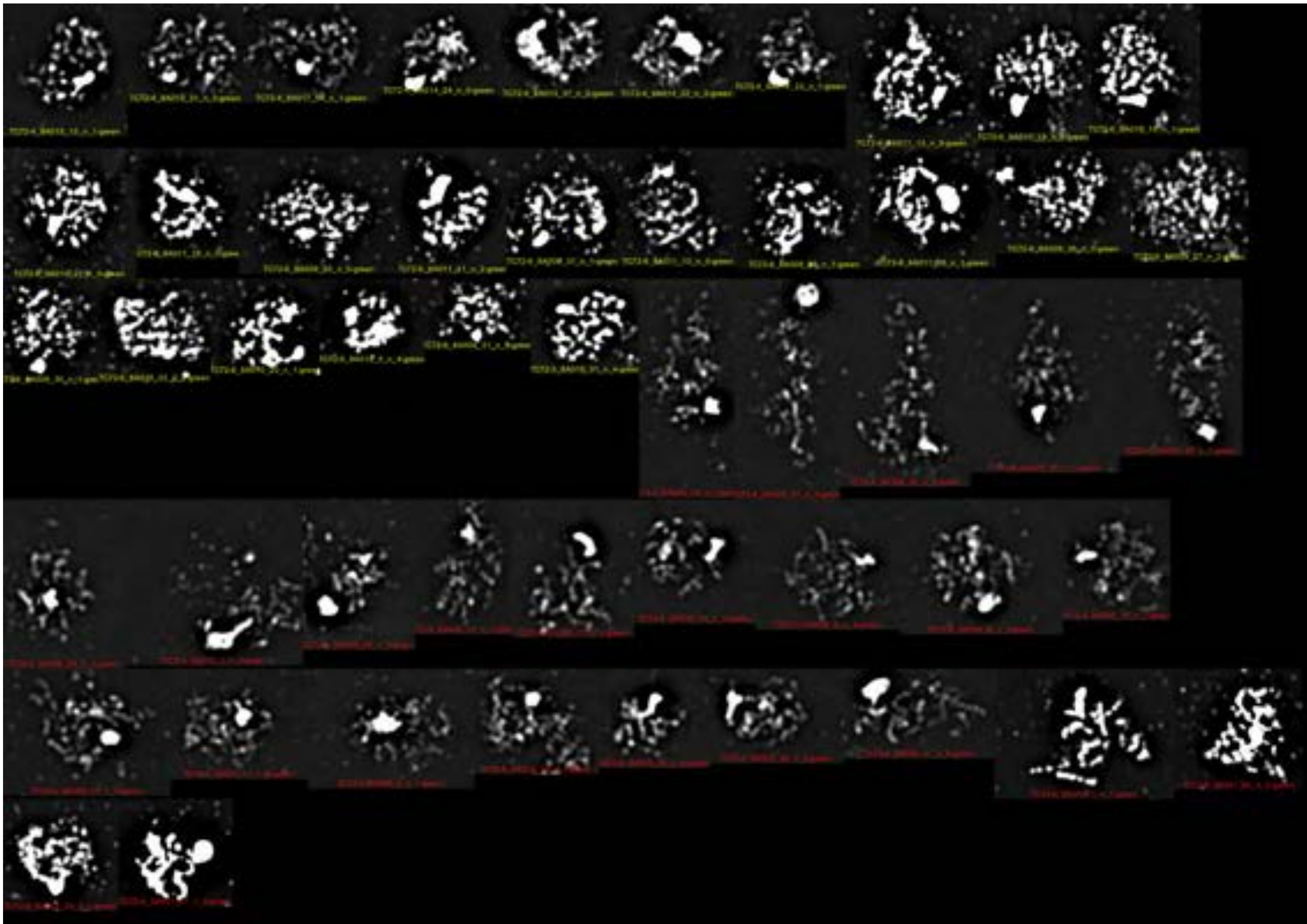

PC+DISCONTINUOUS-2

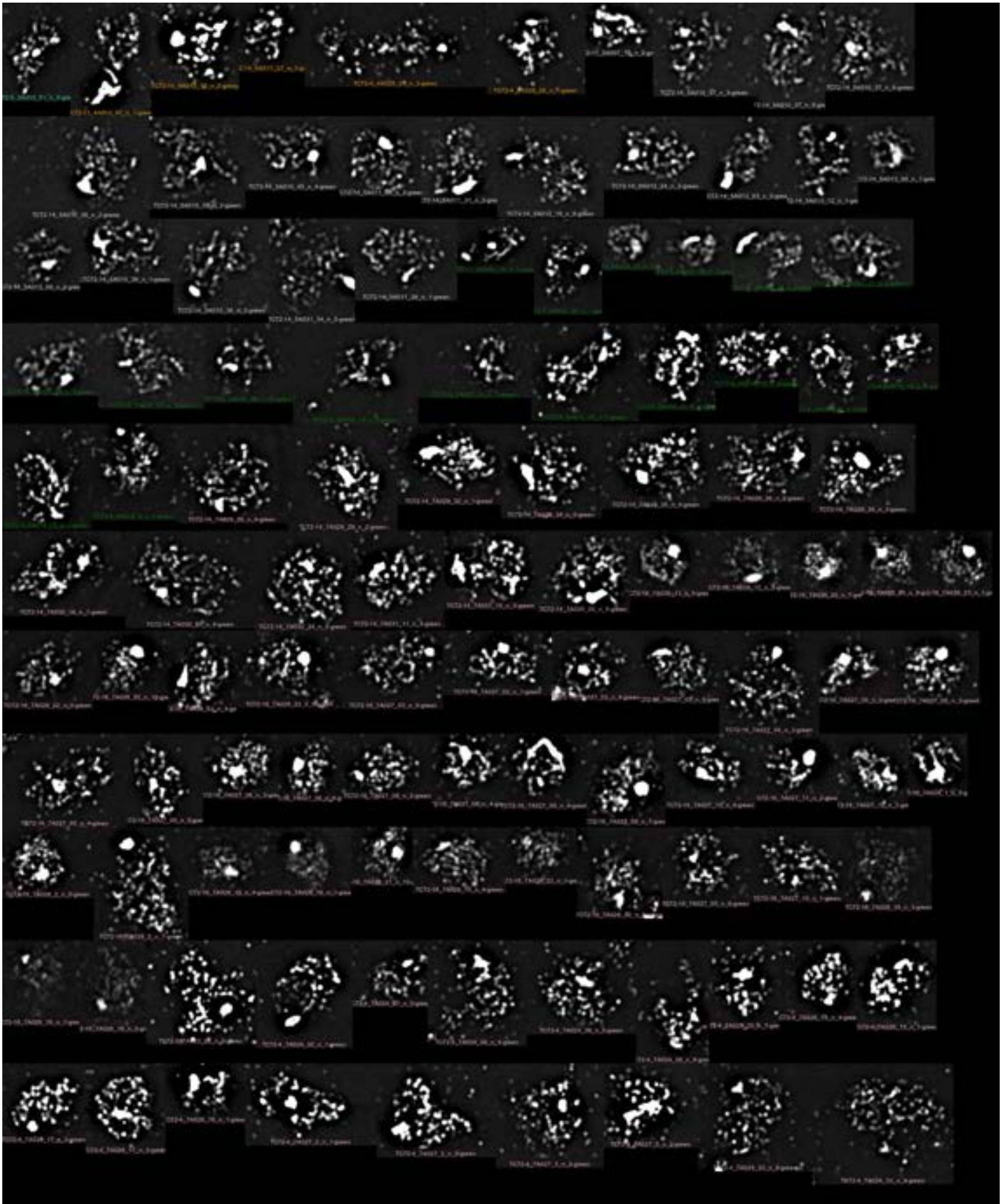

PC+FEW-1

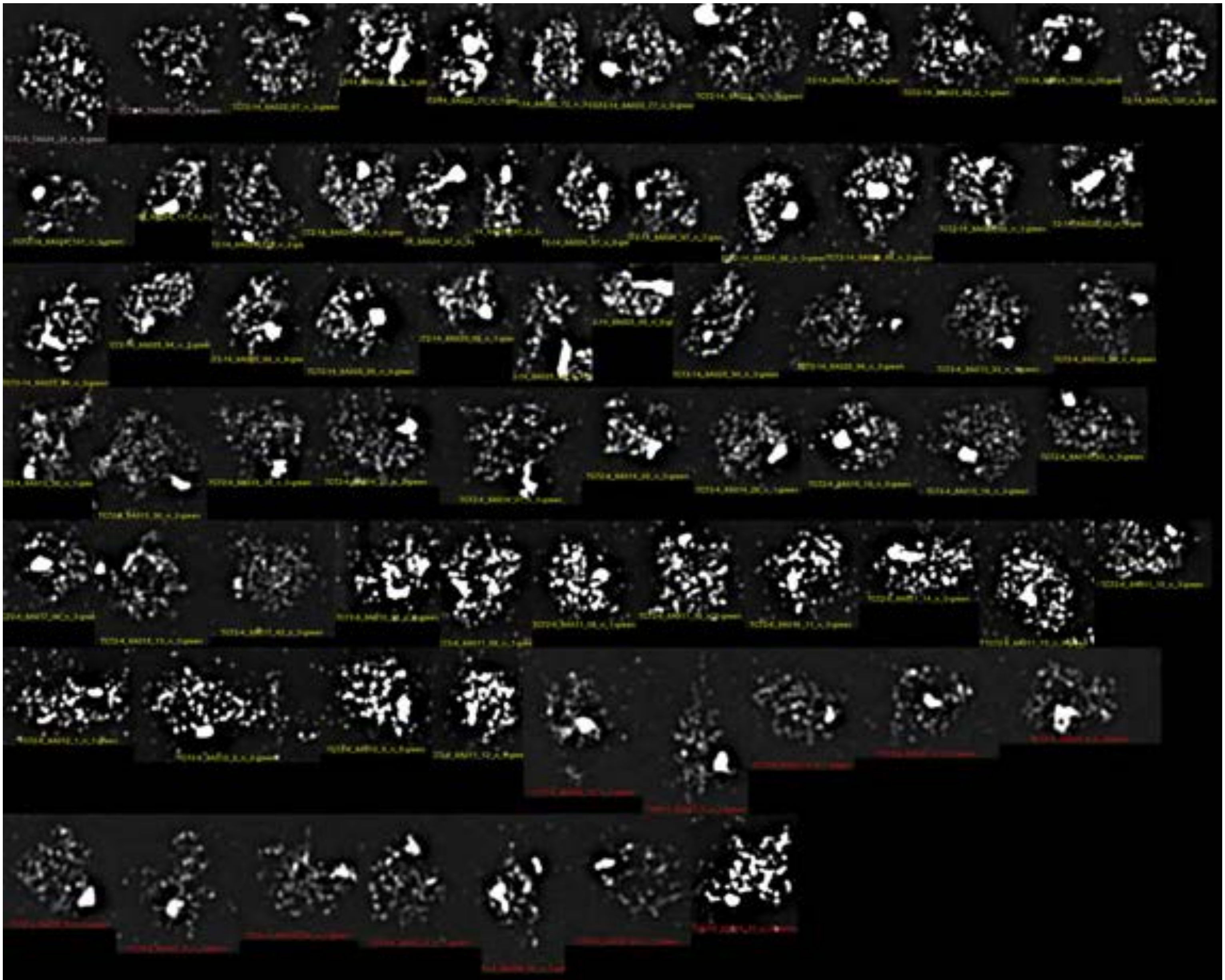

PC+FEW-2

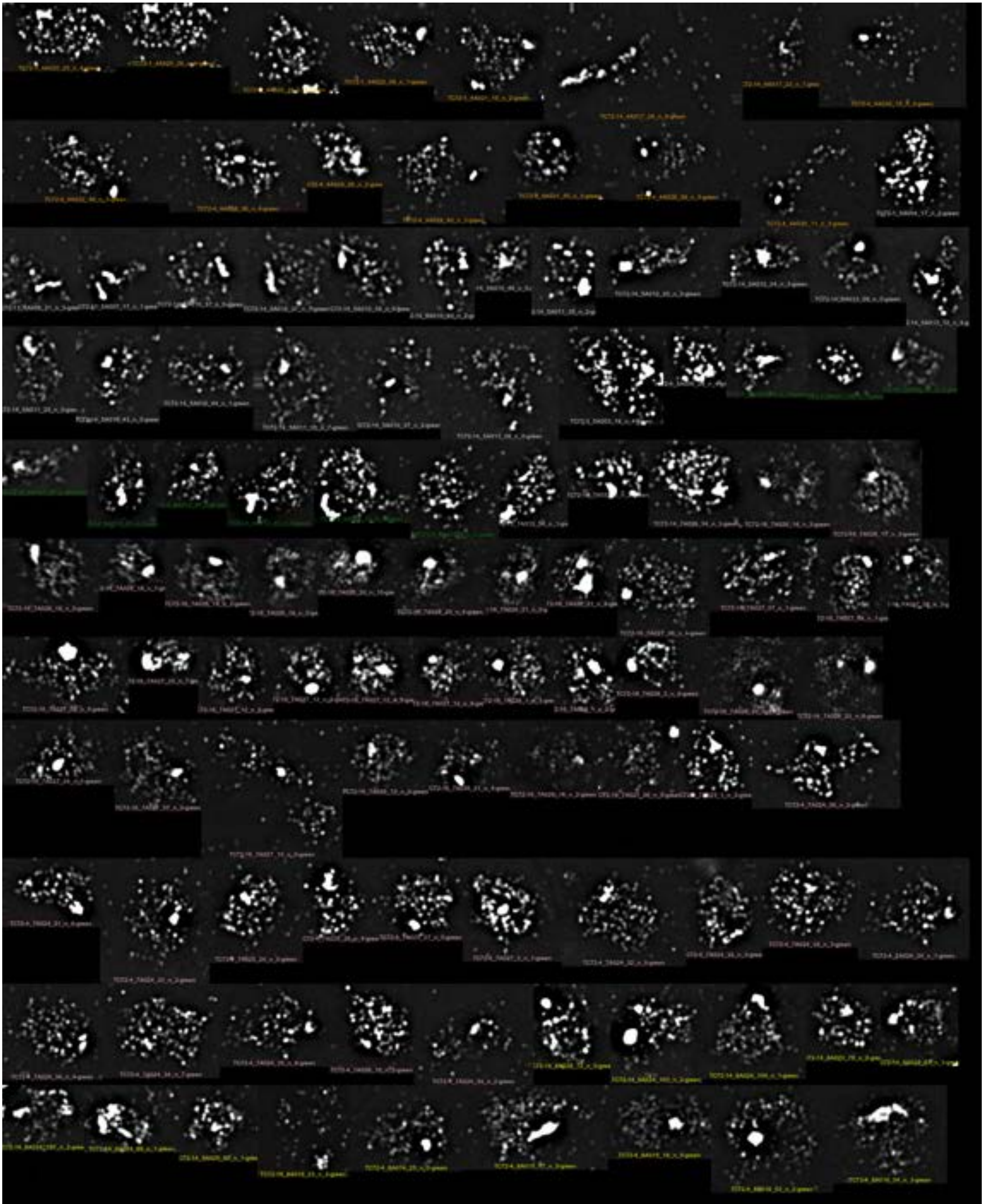

PC+NONE-1

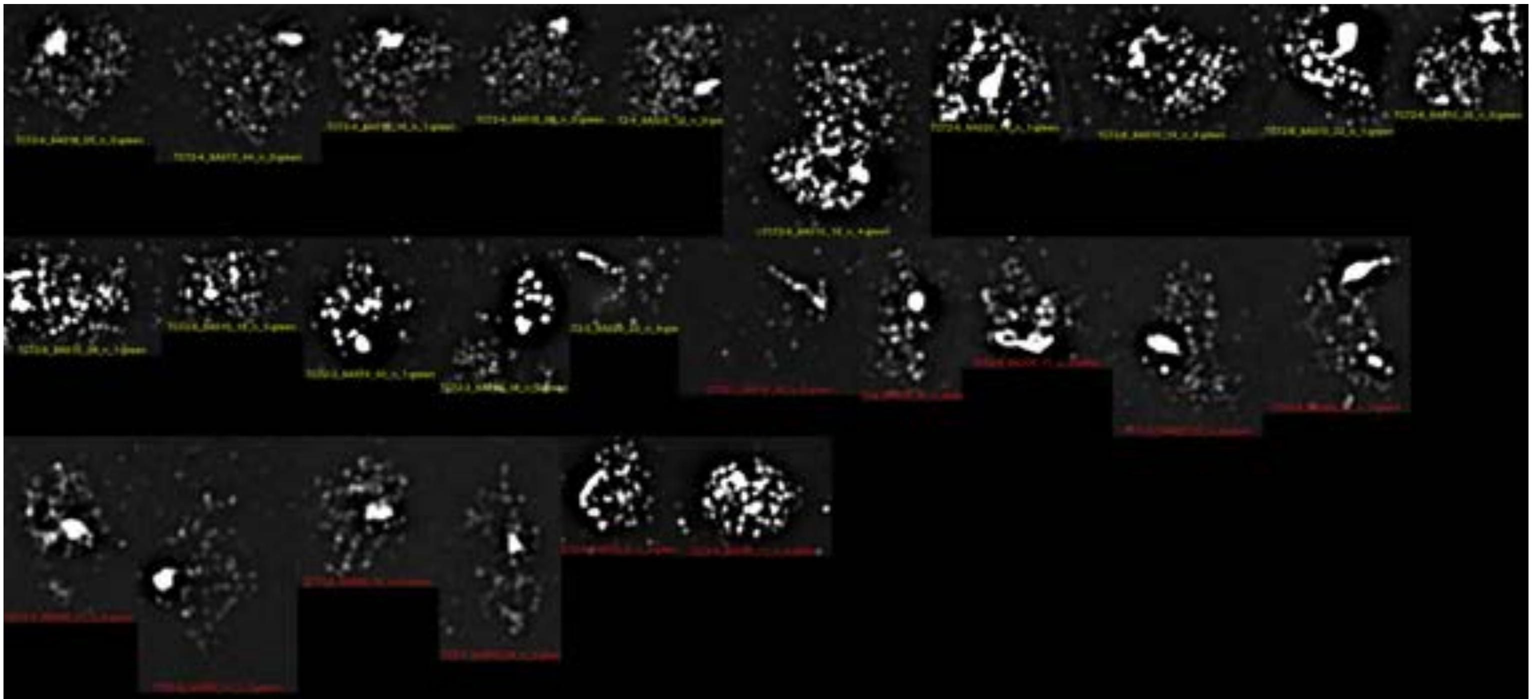

PC+NONE-2

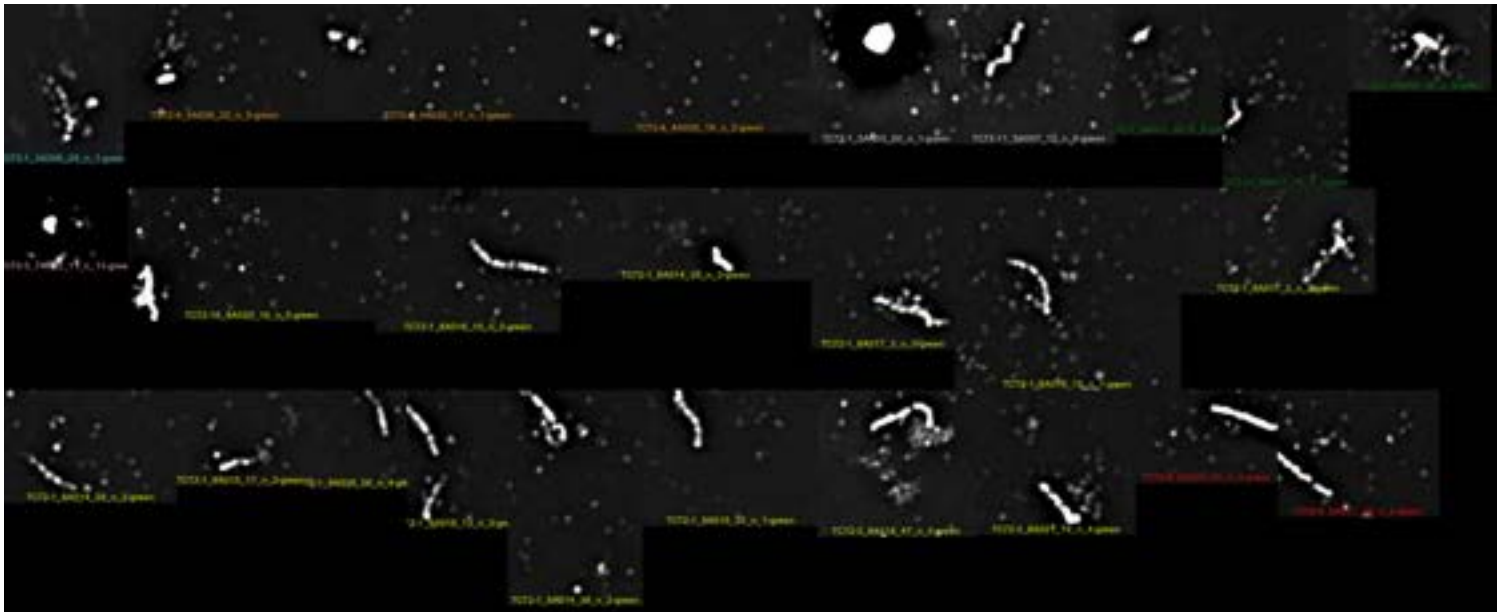

PC+Zip1minus-1

Supplemental Excel S1

### I. Data Input

**For Data Input:** Go to the **"Kinetics"** tab, click the "Load Sample Data" button and type the name of the sheet to be loaded:

a. **"MyData (1)" sheet:** Load your own data.

This worksheet allows you to store your original data and re-load it for a clean start.

b. **"SampleData" sheet:** Load data from Sandhu et al. 2020.

c. **"DemoData" sheet:** Load a sample that demonstrates input format.

Loading this worksheet will show you how to enter your data directly into the "Kinetics" worksheet and walk you through the data set up.

| Start Row | End Row |
|---|---|
| 4 | 46 |

You can find a 'Star Row' and 'End Row' input cells in the worksheets mentioned above. They allow you to limit the analysis to a portion of a data set. In the 'MyData (1)' worksheet, these values indicate the portion of a data set transferred to the 'Kinetics' tab using the 'Load Sample Data' button. In the 'Kinetics' worksheet, these values indicate the range of a data set included in the calculations.

| Event | Measurements | | Total # Events | |
|---|---|---|---|---|
| | Time (Hrs) = X | Units = Y | Units | |
| Event A (exp 1) | 0 | YA0 | Total class A events | 1 |
| | 1 | YA1 | | |
| | n | YAn | | |
| | Leave ≥ 1 row blank | | | |
| Event B (exp 1) | 0 | YB0 | Total class B events | |
| | 1 | YB1 | | |
| | n | Ybn | | |
| | Leave ≥ 1 row blank | | | |
| Event A (exp 2) | 0 | YA0 | Total class A events | 2 |
| | 1 | YA1 | | |
| | n | YAn | | |
| End | | | | |

### I. Data Input

**Notes:**

**1.** The Macro can be used to analyze data from multiple experiments at the same time.

**2.** Event (A) should be available for each experiment and should be usable as alternative reference point (i.e. instead of t = 0). If possible, Event (A) should be intrinsically cumulative.

**3.** Time points should start at t = 0.

**4.** If appropriate, set first and last event unit measurements (Y) manually to 0 (e.g. by background subtraction) .

**5.** For intrinsically cumulative events, the last measured value may be less than the total # events. However, the value measured at the last timepoint should be highest. Alternatively, the event should be categorized as cumulative by adding "(Cum)" at the end of its name.

**6.** Leave ≥ one row blank between (i) different events and (ii) different experimental sets.

**7.** For each event class "Total events" need to be entered, in same units as unit measurements. "Total events" for a given event may vary between different experiments.

**8.** Event labels must contain a common letter sequence to generate averages from different experiments.

**9.** In the "Averages" section, column headings "Event A to F" should be replaced with your specific event labels. These values will serve as identifiers for each event type. The total number of averages should be entered next to the headings **(These can be as many as needed beyod "F").**

**10.** You may choose to perform the calculations on only a subset of the data. To do this, type the number of the first row in the "Row start" section above the input section and "End" below the last row to be included in the analysis.

### II. Cumulative calculations

A. The **"Calculate"** button triggers the following Macro operations:

1. **Lifespan calculation:** The area under the curve of transient events is determined by adding all trapezoids and dividing them by the total number of the specific events (see Figure 1).

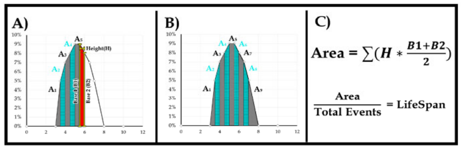

Figure 1.

2. **Conversion from transient to cumulative data:**
   a. On curve of measured values, values are interpolated at one-lifespan intervals in both directions from the peak (Figure 2, yellow asterisks). Starting lifespan interpolations at peak center ensures that cumulative plots reach the maximum.
   b. The cumulative curve is calculated as follows (Figure 2):
      i. All points before the first non-zero point are the same as non-cumulative measured values i.e. y(0).
      ii. The 1st non-zero point is [y(0)s + y(1st)].
      iii. The 2nd non-zero point at t(1st+1 lifespan) is y(2nd) plus [y(0) + y(1st)].
      iv. The nth point on the cumulative curve is the sum of y(nth) plus all interpolated y-values at preceding one-lifespan increments.

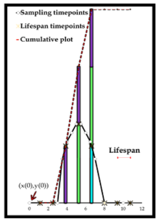

Figure 2.

3. **Options:**
   a. **Do you want to use an alternate reference event?**
   To account for experiment-to-experiment variations, you may set the 50% entry point of an arbitrary reference event (Event A) to x = 0.
   b. **"Store Data" :** Stores your input along with Macro output on separate worksheet. This can be the included MyOutput worksheets or any other user generated worksheet.

### II. Cumulative calculations

c. **Do you want to extrapolate the first time point ?**
The slope between the first two measured values > 0 can be used to extrapolate the time point where y = 0. This option is recommended if there is a large time gap between the 1st (e.g. t = 0 h) and 2nd (t > 0 h) value measurement (Figure 3. B). It avoids an overestimation of the event life span.

Example: If the first non-zero measurement was taken at t = 3.5 h (instead of t = 3h), this erroneously increases the calculated lifespan (Figure 3).

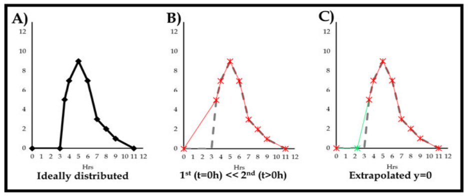

Figure 3.

**B. Graphing options:**

The **"Graph Data"** button gives the following options:

1. **Percent of maximum:** Plots cumulative data as percent of their respective maximum values facilitating comparison between event classes with different total events.

2. **Unaltered input measurements:** Plots entered data .

3. **Cumulative data:**
   a. At interpolated time points: Returns cumulative plots showing calculated values at one-lifespan increments.
   b. At entered time points: Returns cumulative plots showing calculated values at actual times of measurements.

### III. Average calculations

**"Average" :** Calculates and plots averages for each class including error bars (standard deviations).

**Worksheet navigation:**

**"Top" / "Bottom" :** Navigate to First/ Last data row.

**[ Magnifying glass]:** Fits worksheet to window.

**"Clear Output" :** Clears all calculations, keeps input data.

**"Clear All" :** Clears input and output, keeps stored worksheet.

| Input | | | | LifeSpan | Cumulative Values: | | | | Lifespan @ the 50 percentile | | Percentage of maximum | | In 10% increments | |
|---|---|---|---|---|---|---|---|---|---|---|---|---|---|---|
| | Measurements | | Total # Events | Time (Hrs) | at interpolated timepoints | | at taken timepoints | | 0.50 | 0.50 | Time (Hrs) | Units | Time (Hrs) | Units |
| Event Class | Time (Hrs) | Units | Units | | Time (Hrs) | Units | Time (Hrs) | Units | Entry | Exit | | | | |
| **CO-72-avg1.+3. WT (23C)** | 0.00 | 0.00 | 0.22 | | 0.00 | 0.00 | 0.00 | 2% | 6.71 | | 0.00 | 0.02 | 5.37 | 0.10 |
| | 3.00 | 0.01 | | | 3.00 | 0.01 | 3.00 | 3% | | | 3.00 | 0.03 | 6.02 | 0.20 |
| | 4.00 | 0.01 | | | 4.00 | 0.01 | 4.00 | 4% | | | 4.00 | 0.04 | 6.25 | 0.30 |
| | 5.00 | 0.01 | | | 5.00 | 0.01 | 5.00 | 5% | | | 5.00 | 0.05 | 6.48 | 0.40 |
| | 6.00 | 0.04 | | | 6.00 | 0.04 | 6.00 | 19% | | | 6.00 | 0.19 | 6.71 | 0.50 |
| | 7.00 | 0.14 | | | 7.00 | 0.14 | 7.00 | 63% | | | 7.00 | 0.63 | 6.94 | 0.60 |
| | 8.00 | 0.19 | | | 8.00 | 0.19 | 8.00 | 87% | | | 8.00 | 0.87 | 7.30 | 0.70 |
| | 9.00 | 0.22 | | | 9.00 | 0.22 | 9.00 | 98% | | | 9.00 | 0.98 | 7.72 | 0.80 |
| | 11.00 | 0.22 | | | 11.00 | 0.22 | 11.00 | 100% | | | 11.00 | 1.00 | | |
| | 12.00 | 0.22 | | | 12.00 | 0.22 | 12.00 | 100% | | | 12.00 | 1.00 | | |
| **DSB-72-avg1.+3. WT (23C)** | 0.00 | 0.00 | 0.23 | 1.23 | 0.00 | 0.00 | 0.00 | 0% | 4.34 | 5.57 | 0.00 | 0.00 | 2.75 | 0.10 |
| | 3.00 | 0.01 | | | 0.08 | 0.00 | 3.00 | 15% | | | 0.08 | 0.00 | 3.23 | 0.20 |
| | 4.00 | 0.07 | | | 1.31 | 0.00 | 4.00 | 39% | | | 1.31 | 0.02 | 3.71 | 0.30 |
| | 5.00 | 0.09 | | | 2.54 | 0.01 | 5.00 | 72% | | | 2.54 | 0.06 | 4.04 | 0.40 |
| | 6.00 | 0.06 | | | 3.77 | 0.07 | 6.00 | 90% | | | 3.77 | 0.31 | 4.34 | 0.50 |
| | 7.00 | 0.02 | | | 5.00 | 0.16 | 7.00 | 99% | | | 5.00 | 0.72 | 4.65 | 0.60 |
| | 8.00 | 0.01 | | | 6.23 | 0.21 | 8.00 | 102% | | | 6.23 | 0.94 | 4.95 | 0.70 |
| | 9.00 | 0.00 | | | 7.46 | 0.23 | 9.00 | 103% | | | 7.46 | 1.01 | 5.46 | 0.80 |
| | 11.00 | 0.00 | | | 8.69 | 0.23 | | | | | 8.69 | 1.03 | | |
| | | | | | 9.92 | 0.23 | | | | | 9.92 | 1.04 | | |
| **SEI-72-avg1.+3. WT (23C)** | 0.00 | 0.00 | 0.11 | 0.26 | 0.00 | 0.00 | 0.00 | 1% | 5.83 | 609% | 0.00 | 0.01 | 3.94 | 0.10 |
| | 3.00 | 0.00 | | | 0.06 | 0.00 | 3.00 | 5% | | | 0.06 | 0.01 | 4.62 | 0.20 |
| | 4.00 | 0.00 | | | 0.32 | 0.00 | 4.00 | 11% | | | 0.32 | 0.02 | 5.05 | 0.30 |
| | 5.00 | 0.01 | | | 0.58 | 0.00 | 5.00 | 29% | | | 0.58 | 0.02 | 5.45 | 0.40 |
| | 6.00 | 0.01 | | | 0.84 | 0.00 | 6.00 | 54% | | | 0.84 | 0.03 | 5.83 | 0.50 |
| | 7.00 | 0.00 | | | 1.09 | 0.00 | 7.00 | 74% | | | 1.09 | 0.03 | 6.23 | 0.60 |
| | 8.00 | 0.00 | | | 1.35 | 0.00 | 8.00 | 85% | | | 1.35 | 0.04 | 6.73 | 0.70 |
| | 9.00 | 0.00 | | | 1.61 | 0.00 | 9.00 | 91% | | | 1.61 | 0.04 | 7.46 | 0.80 |
| | 11.00 | 0.00 | | | 1.87 | 0.00 | | | | | 1.87 | 0.04 | | |
| | | | | | 2.13 | 0.01 | | | | | 2.13 | 0.05 | | |
| | | | | | 2.39 | 0.01 | | | | | 2.39 | 0.05 | | |
| | | | | | 2.64 | 0.01 | | | | | 2.64 | 0.05 | | |
| | | | | | 2.90 | 0.01 | | | | | 2.90 | 0.05 | | |
| | | | | | 3.16 | 0.01 | | | | | 3.16 | 0.06 | | |
| | | | | | 3.42 | 0.01 | | | | | 3.42 | 0.07 | | |
| | | | | | 3.68 | 0.01 | | | | | 3.68 | 0.08 | | |
| | | | | | 3.93 | 0.01 | | | | | 3.93 | 0.10 | | |
| | | | | | 4.19 | 0.01 | | | | | 4.19 | 0.13 | | |
| | | | | | 4.45 | 0.02 | | | | | 4.45 | 0.17 | | |
| | | | | | 4.71 | 0.02 | | | | | 4.71 | 0.22 | | |
| | | | | | 4.97 | 0.03 | | | | | 4.97 | 0.28 | | |
| | | | | | 5.23 | 0.04 | | | | | 5.23 | 0.34 | | |
| | | | | | 5.48 | 0.04 | | | | | 5.48 | 0.41 | | |
| | | | | | 5.74 | 0.05 | | | | | 5.74 | 0.48 | | |
| | | | | | 6.00 | 0.06 | | | | | 6.00 | 0.54 | | |
| | | | | | 6.26 | 0.07 | | | | | 6.26 | 0.61 | | |
| | | | | | 6.52 | 0.07 | | | | | 6.52 | 0.66 | | |
| | | | | | 6.77 | 0.08 | | | | | 6.77 | 0.71 | | |
| | | | | | 7.03 | 0.08 | | | | | 7.03 | 0.75 | | |
| | | | | | 7.29 | 0.09 | | | | | 7.29 | 0.78 | | |
| | | | | | 7.55 | 0.09 | | | | | 7.55 | 0.81 | | |
| | | | | | 7.81 | 0.09 | | | | | 7.81 | 0.83 | | |
| | | | | | 8.07 | 0.09 | | | | | 8.07 | 0.85 | | |
| | | | | | 8.32 | 0.10 | | | | | 8.32 | 0.87 | | |
| | | | | | 8.58 | 0.10 | | | | | 8.58 | 0.89 | | |
| | | | | | 8.84 | 0.10 | | | | | 8.84 | 0.91 | | |
| | | | | | 9.10 | 0.10 | | | | | 9.10 | 0.92 | | |
| | | | | | 9.36 | 0.10 | | | | | 9.36 | 0.94 | | |
| | | | | | 9.61 | 0.10 | | | | | 9.61 | 0.95 | | |
| | | | | | 9.87 | 0.11 | | | | | 9.87 | 0.96 | | |
| | | | | | 10.13 | 0.11 | | | | | 10.13 | 0.98 | | |
| | | | | | 10.39 | 0.11 | | | | | 10.39 | 0.99 | | |
| | | | | | 10.65 | 0.11 | | | | | 10.65 | 1.00 | | |
| | | | | | 10.91 | 0.11 | | | | | 10.91 | 1.01 | | |
| **IH-dHJs-72-avg1.+3. WT (23C)** | 0.00 | 0.00 | 0.22 | 0.13 | 0.00 | 0.00 | 0.00 | 0% | 6.06 | 6.19 | 0.00 | 0.00 | 4.32 | 0.10 |
| | 3.00 | 0.00 | | | 0.08 | 0.00 | 3.00 | 3% | | | 0.08 | 0.00 | 4.90 | 0.20 |
| | 4.00 | 0.00 | | | 0.21 | 0.00 | 4.00 | 7% | | | 0.21 | 0.00 | 5.32 | 0.30 |
| | 5.00 | 0.01 | | | 0.34 | 0.00 | 5.00 | 22% | | | 0.34 | 0.01 | 5.71 | 0.40 |
| | 6.00 | 0.01 | | | 0.47 | 0.00 | 6.00 | 48% | | | 0.47 | 0.01 | 6.06 | 0.50 |
| | 7.00 | 0.00 | | | 0.60 | 0.00 | 7.00 | 70% | | | 0.60 | 0.01 | 6.47 | 0.60 |
| | 8.00 | 0.00 | | | 0.72 | 0.00 | 8.00 | 81% | | | 0.72 | 0.01 | 7.02 | 0.70 |
| | 9.00 | 0.00 | | | 0.85 | 0.00 | 9.00 | 89% | | | 0.85 | 0.01 | 7.85 | 0.80 |
| | 11.00 | 0.00 | | | 0.98 | 0.00 | | | | | 0.98 | 0.01 | | |
| | | | | | 1.11 | 0.00 | | | | | 1.11 | 0.01 | | |
| | | | | | 1.24 | 0.00 | | | | | 1.24 | 0.01 | | |
| | | | | | 1.37 | 0.00 | | | | | 1.37 | 0.02 | | |
| | | | | | 1.50 | 0.00 | | | | | 1.50 | 0.02 | | |
| | | | | | 1.62 | 0.00 | | | | | 1.62 | 0.02 | | |
| | | | | | 1.75 | 0.00 | | | | | 1.75 | 0.02 | | |
| | | | | | 1.88 | 0.00 | | | | | 1.88 | 0.02 | | |
| | | | | | 2.01 | 0.00 | | | | | 2.01 | 0.02 | | |
| | | | | | 2.14 | 0.01 | | | | | 2.14 | 0.02 | | |
| | | | | | 2.27 | 0.01 | | | | | 2.27 | 0.02 | | |
| | | | | | 2.40 | 0.01 | | | | | 2.40 | 0.03 | | |
| | | | | | 2.53 | 0.01 | | | | | 2.53 | 0.03 | | |
| | | | | | 2.65 | 0.01 | | | | | 2.65 | 0.03 | | |
| | | | | | 2.78 | 0.01 | | | | | 2.78 | 0.03 | | |
| | | | | | 2.91 | 0.01 | | | | | 2.91 | 0.03 | | |
| | | | | | 3.04 | 0.01 | | | | | 3.04 | 0.03 | | |
| | | | | | 3.17 | 0.01 | | | | | 3.17 | 0.03 | | |
| | | | | | 3.30 | 0.01 | | | | | 3.30 | 0.04 | | |
| | | | | | 3.43 | 0.01 | | | | | 3.43 | 0.04 | | |
| | | | | | 3.55 | 0.01 | | | | | 3.55 | 0.05 | | |
| | | | | | 3.68 | 0.01 | | | | | 3.68 | 0.05 | | |
| | | | | | 3.81 | 0.01 | | | | | 3.81 | 0.06 | | |
| | | | | | 3.94 | 0.01 | | | | | 3.94 | 0.07 | | |
| | | | | | 4.07 | 0.02 | | | | | 4.07 | 0.07 | | |
| | | | | | 4.20 | 0.02 | | | | | 4.20 | 0.09 | | |
| | | | | | 4.33 | 0.02 | | | | | 4.33 | 0.10 | | |
| | | | | | 4.46 | 0.03 | | | | | 4.46 | 0.12 | | |
| | | | | | 4.58 | 0.03 | | | | | 4.58 | 0.14 | | |
| | | | | | 4.71 | 0.04 | | | | | 4.71 | 0.16 | | |

| | | | | | | | | | | | | | | |
|---|---|---|---|---|---|---|---|---|---|---|---|---|---|---|
| | | | | | 4.84 | 0.04 | | | | | 4.84 | 0.19 | | |
| | | | | | 4.97 | 0.05 | | | | | 4.97 | 0.22 | | |
| | | | | | 5.10 | 0.05 | | | | | 5.10 | 0.25 | | |
| | | | | | 5.23 | 0.06 | | | | | 5.23 | 0.28 | | |
| | | | | | 5.36 | 0.07 | | | | | 5.36 | 0.31 | | |
| | | | | | 5.49 | 0.08 | | | | | 5.49 | 0.34 | | |
| | | | | | 5.61 | 0.08 | | | | | 5.61 | 0.38 | | |
| | | | | | 5.74 | 0.09 | | | | | 5.74 | 0.41 | | |
| | | | | | 5.87 | 0.10 | | | | | 5.87 | 0.45 | | |
| | | | | | 6.00 | 0.11 | | | | | 6.00 | 0.48 | | |
| | | | | | 6.13 | 0.11 | | | | | 6.13 | 0.52 | | |
| | | | | | 6.26 | 0.12 | | | | | 6.26 | 0.55 | | |
| | | | | | 6.39 | 0.13 | | | | | 6.39 | 0.58 | | |
| | | | | | 6.51 | 0.13 | | | | | 6.51 | 0.61 | | |
| | | | | | 6.64 | 0.14 | | | | | 6.64 | 0.64 | | |
| | | | | | 6.77 | 0.15 | | | | | 6.77 | 0.66 | | |
| | | | | | 6.90 | 0.15 | | | | | 6.90 | 0.68 | | |
| | | | | | 7.03 | 0.15 | | | | | 7.03 | 0.70 | | |
| | | | | | 7.16 | 0.16 | | | | | 7.16 | 0.72 | | |
| | | | | | 7.29 | 0.16 | | | | | 7.29 | 0.74 | | |
| | | | | | 7.42 | 0.17 | | | | | 7.42 | 0.75 | | |
| | | | | | 7.54 | 0.17 | | | | | 7.54 | 0.77 | | |
| | | | | | 7.67 | 0.17 | | | | | 7.67 | 0.78 | | |
| | | | | | 7.80 | 0.18 | | | | | 7.80 | 0.80 | | |
| | | | | | 7.93 | 0.18 | | | | | 7.93 | 0.81 | | |
| | | | | | 8.06 | 0.18 | | | | | 8.06 | 0.82 | | |
| | | | | | 8.19 | 0.18 | | | | | 8.19 | 0.83 | | |
| | | | | | 8.32 | 0.18 | | | | | 8.32 | 0.84 | | |
| | | | | | 8.45 | 0.19 | | | | | 8.45 | 0.85 | | |
| | | | | | 8.57 | 0.19 | | | | | 8.57 | 0.86 | | |
| | | | | | 8.70 | 0.19 | | | | | 8.70 | 0.87 | | |
| | | | | | 8.83 | 0.19 | | | | | 8.83 | 0.88 | | |
| | | | | | 8.96 | 0.20 | | | | | 8.96 | 0.89 | | |
| | | | | | 9.09 | 0.20 | | | | | 9.09 | 0.90 | | |
| | | | | | 9.22 | 0.20 | | | | | 9.22 | 0.90 | | |
| | | | | | 9.35 | 0.20 | | | | | 9.35 | 0.91 | | |
| | | | | | 9.47 | 0.20 | | | | | 9.47 | 0.92 | | |
| | | | | | 9.60 | 0.20 | | | | | 9.60 | 0.93 | | |
| | | | | | 9.73 | 0.21 | | | | | 9.73 | 0.94 | | |
| | | | | | 9.86 | 0.21 | | | | | 9.86 | 0.94 | | |
| | | | | | 9.99 | 0.21 | | | | | 9.99 | 0.95 | | |
| | | | | | 10.12 | 0.21 | | | | | 10.12 | 0.96 | | |
| | | | | | 10.25 | 0.21 | | | | | 10.25 | 0.97 | | |
| | | | | | 10.38 | 0.21 | | | | | 10.38 | 0.97 | | |
| | | | | | 10.50 | 0.22 | | | | | 10.50 | 0.98 | | |
| | | | | | 10.63 | 0.22 | | | | | 10.63 | 0.99 | | |
| | | | | | 10.76 | 0.22 | | | | | 10.76 | 0.99 | | |
| | | | | | 10.89 | 0.22 | | | | | 10.89 | 1.00 | | |
| **5 Zip1 lines** | 0.00 | 0.00 | 0.98 | 0.19 | 0.00 | 0.00 | 0.00 | 0% | 5.69 | 5.88 | 0.00 | 0.00 | 3.56 | 0.10 |
| | 3.00 | 0.01 | | | 0.17 | 0.00 | 3.00 | 8% | | | 0.17 | 0.00 | 4.66 | 0.20 |
| | 4.00 | 0.01 | | | 0.35 | 0.00 | 4.00 | 12% | | | 0.35 | 0.00 | 5.09 | 0.30 |
| | 5.00 | 0.04 | | | 0.54 | 0.00 | 5.00 | 28% | | | 0.54 | 0.00 | 5.42 | 0.40 |
| | 6.00 | 0.08 | | | 0.73 | 0.01 | 6.00 | 63% | | | 0.73 | 0.01 | 5.69 | 0.50 |
| | 7.00 | 0.02 | | | 0.92 | 0.01 | 7.00 | 89% | | | 0.92 | 0.01 | 5.93 | 0.60 |
| | 8.00 | 0.01 | | | 1.11 | 0.01 | 8.00 | 98% | | | 1.11 | 0.01 | 6.18 | 0.70 |
| | 9.00 | 0.00 | | | 1.29 | 0.01 | | | | | 1.29 | 0.02 | 6.52 | 0.80 |
| | | | | | 1.48 | 0.02 | | | | | 1.48 | 0.02 | | |
| | | | | | 1.67 | 0.02 | | | | | 1.67 | 0.02 | | |
| | | | | | 1.86 | 0.03 | | | | | 1.86 | 0.03 | | |
| | | | | | 2.05 | 0.04 | | | | | 2.05 | 0.04 | | |
| | | | | | 2.24 | 0.04 | | | | | 2.24 | 0.04 | | |
| | | | | | 2.42 | 0.05 | | | | | 2.42 | 0.05 | | |
| | | | | | 2.61 | 0.06 | | | | | 2.61 | 0.06 | | |
| | | | | | 2.80 | 0.06 | | | | | 2.80 | 0.07 | | |
| | | | | | 2.99 | 0.07 | | | | | 2.99 | 0.07 | | |
| | | | | | 3.18 | 0.08 | | | | | 3.18 | 0.08 | | |
| | | | | | 3.36 | 0.09 | | | | | 3.36 | 0.09 | | |
| | | | | | 3.55 | 0.10 | | | | | 3.55 | 0.10 | | |
| | | | | | 3.74 | 0.11 | | | | | 3.74 | 0.11 | | |
| | | | | | 3.93 | 0.11 | | | | | 3.93 | 0.11 | | |
| | | | | | 4.12 | 0.12 | | | | | 4.12 | 0.13 | | |
| | | | | | 4.31 | 0.14 | | | | | 4.31 | 0.14 | | |
| | | | | | 4.49 | 0.17 | | | | | 4.49 | 0.17 | | |
| | | | | | 4.68 | 0.20 | | | | | 4.68 | 0.20 | | |
| | | | | | 4.87 | 0.24 | | | | | 4.87 | 0.24 | | |
| | | | | | 5.06 | 0.28 | | | | | 5.06 | 0.29 | | |
| | | | | | 5.25 | 0.34 | | | | | 5.25 | 0.34 | | |
| | | | | | 5.44 | 0.40 | | | | | 5.44 | 0.40 | | |
| | | | | | 5.62 | 0.46 | | | | | 5.62 | 0.47 | | |
| | | | | | 5.81 | 0.54 | | | | | 5.81 | 0.55 | | |
| | | | | | 6.00 | 0.62 | | | | | 6.00 | 0.63 | | |
| | | | | | 6.19 | 0.69 | | | | | 6.19 | 0.70 | | |
| | | | | | 6.38 | 0.75 | | | | | 6.38 | 0.76 | | |
| | | | | | 6.56 | 0.80 | | | | | 6.56 | 0.81 | | |
| | | | | | 6.75 | 0.83 | | | | | 6.75 | 0.85 | | |
| | | | | | 6.94 | 0.86 | | | | | 6.94 | 0.88 | | |
| | | | | | 7.13 | 0.88 | | | | | 7.13 | 0.90 | | |
| | | | | | 7.32 | 0.90 | | | | | 7.32 | 0.92 | | |
| | | | | | 7.51 | 0.92 | | | | | 7.51 | 0.94 | | |
| | | | | | 7.69 | 0.94 | | | | | 7.69 | 0.95 | | |
| | | | | | 7.88 | 0.95 | | | | | 7.88 | 0.97 | | |
| | | | | | 8.07 | 0.96 | | | | | 8.07 | 0.98 | | |
| | | | | | 8.26 | 0.97 | | | | | 8.26 | 0.99 | | |
| | | | | | 8.45 | 0.98 | | | | | 8.45 | 1.00 | | |
| | | | | | 8.64 | 0.98 | | | | | 8.64 | 1.00 | | |
| | | | | | 8.82 | 0.98 | | | | | 8.82 | 1.00 | | |
| **4 Zip1 partial lines** | 0.00 | 0.00 | 0.98 | 0.85 | 0.00 | 0.00 | 0.00 | 0% | 4.97 | 5.82 | 0.00 | 0.00 | 2.81 | 0.10 |
| | 3.00 | 0.05 | | | 0.75 | 0.01 | 3.00 | 11% | | | 0.75 | 0.01 | 3.94 | 0.20 |
| | 4.00 | 0.06 | | | 1.60 | 0.04 | 4.00 | 21% | | | 1.60 | 0.04 | 4.38 | 0.30 |
| | 5.00 | 0.28 | | | 2.45 | 0.08 | 5.00 | 51% | | | 2.45 | 0.08 | 4.67 | 0.40 |
| | 6.00 | 0.25 | | | 3.30 | 0.13 | 6.00 | 80% | | | 3.30 | 0.13 | 4.97 | 0.50 |
| | 7.00 | 0.10 | | | 4.15 | 0.22 | 7.00 | 95% | | | 4.15 | 0.22 | 5.29 | 0.60 |
| | 8.00 | 0.04 | | | 5.00 | 0.50 | 8.00 | 101% | | | 5.00 | 0.51 | 5.62 | 0.70 |
| | 9.00 | 0.00 | | | 5.85 | 0.76 | | | | | 5.85 | 0.77 | 6.01 | 0.80 |
| | | | | | 6.70 | 0.90 | | | | | 6.70 | 0.92 | | |

| | | | | | | | | | | | | | | |
|---|---|---|---|---|---|---|---|---|---|---|---|---|---|---|
| | | | | | 7.55 | 0.97 | | | | | 7.55 | 0.99 | | |
| | | | | | 8.40 | 1.00 | | | | | 8.40 | 1.02 | | |
| **3 Zip1 few lines** | 0.00 | 0.00 | 0.98 | 2.18 | 0.00 | 0.00 | 0.00 | 0% | 3.15 | 5.33 | 0.00 | 0.00 | 0.77 | 0.10 |
| | 3.00 | 0.37 | | | 0.65 | 0.08 | 3.00 | 47% | | | 0.65 | 0.08 | 1.38 | 0.20 |
| | 4.00 | 0.34 | | | 2.82 | 0.42 | 4.00 | 68% | | | 2.82 | 0.43 | 2.00 | 0.30 |
| | 5.00 | 0.44 | | | 5.00 | 0.86 | 5.00 | 88% | | | 5.00 | 0.88 | 2.62 | 0.40 |
| | 6.00 | 0.26 | | | 7.18 | 1.09 | 6.00 | 99% | | | 7.18 | 1.11 | 3.15 | 0.50 |
| | 7.00 | 0.25 | | | | | 7.00 | 109% | | | | | 3.63 | 0.60 |
| | 8.00 | 0.11 | | | | | 8.00 | | | | | | 4.12 | 0.70 |
| | 9.00 | 0.00 | | | | | | | | | | | 4.60 | 0.80 |
| **2 Zip1 no lines** | 0.00 | 0.00 | 0.98 | 2.42 | 0.00 | 0.00 | 0.00 | 0% | 2.52 | 4.94 | 0.00 | 0.00 | 0.57 | 0.10 |
| | 3.00 | 0.52 | | | 1.58 | 0.27 | 3.00 | 61% | | | 1.58 | 0.28 | 1.13 | 0.20 |
| | 4.00 | 0.56 | | | 4.00 | 0.83 | 4.00 | 85% | | | 4.00 | 0.85 | 1.67 | 0.30 |
| | 5.00 | 0.18 | | | 6.42 | 1.05 | 5.00 | 94% | | | 6.42 | 1.07 | 2.09 | 0.40 |
| | 6.00 | 0.15 | | | 8.84 | 1.07 | 6.00 | 103% | | | 8.84 | 1.09 | 2.52 | 0.50 |
| | 7.00 | 0.31 | | | | | 7.00 | 107% | | | | | 2.95 | 0.60 |
| | 8.00 | 0.13 | | | | | 8.00 | 108% | | | | | 3.37 | 0.70 |
| | 9.00 | 0.00 | | | | | | | | | | | 3.80 | 0.80 |
| **2+3+4+5 Zip1 lines+partial lines** | 0.00 | 0.00 | 0.98 | 3.39 | 0.00 | 0.00 | 0.00 | 0% | 1.66 | 5.05 | 0.00 | 0.00 | 0.31 | 0.10 |
| | 3.00 | 0.94 | | | 0.61 | 0.19 | 3.00 | 89% | | | 0.61 | 0.20 | 0.62 | 0.20 |
| | 4.00 | 0.96 | | | 4.00 | 1.15 | 4.00 | 117% | | | 4.00 | 1.17 | 0.94 | 0.30 |
| | 5.00 | 0.95 | | | | | | | | | | | 1.25 | 0.40 |
| **4+5 Zip1 lines+partial lines** | 0.00 | 0.00 | 0.98 | 1.04 | 0.00 | 0.00 | 0.00 | 0% | 4.98 | 6.02 | 0.00 | 0.00 | 2.75 | 0.10 |
| | 3.00 | 0.06 | | | 0.81 | 0.02 | 3.00 | 11% | | | 0.81 | 0.02 | 4.02 | 0.20 |
| | 4.00 | 0.06 | | | 1.85 | 0.05 | 4.00 | 19% | | | 1.85 | 0.05 | 4.34 | 0.30 |
| | 5.00 | 0.33 | | | 2.89 | 0.10 | 5.00 | 51% | | | 2.89 | 0.11 | 4.66 | 0.40 |
| | 6.00 | 0.33 | | | 3.92 | 0.17 | 6.00 | 83% | | | 3.92 | 0.17 | 4.98 | 0.50 |
| | 7.00 | 0.13 | | | 4.96 | 0.48 | 7.00 | 95% | | | 4.96 | 0.49 | 5.29 | 0.60 |
| | 8.00 | 0.06 | | | 6.00 | 0.81 | 8.00 | 101% | | | 6.00 | 0.83 | 5.59 | 0.70 |
| | 9.00 | 0.00 | | | 7.04 | 0.94 | | | | | 7.04 | 0.96 | 5.90 | 0.80 |
| | | | | | 8.08 | 0.99 | | | | | 8.08 | 1.01 | | |
| **3+4+5 Zip1 lines+partial lines** | 0.00 | 0.00 | 0.98 | 3.22 | 0.00 | 0.00 | 0.00 | 0% | 2.78 | 6.00 | 0.00 | 0.00 | 0.69 | 0.10 |
| | 3.00 | 0.42 | | | 1.78 | 0.25 | 3.00 | 55% | | | 1.78 | 0.26 | 1.39 | 0.20 |
| | 4.00 | 0.40 | | | 5.00 | 1.02 | 4.00 | 80% | | | 5.00 | 1.04 | 1.96 | 0.30 |
| | 5.00 | 0.77 | | | 8.22 | 1.15 | 5.00 | 104% | | | 8.22 | 1.17 | 2.37 | 0.40 |
| | 6.00 | 0.59 | | | | | 6.00 | 108% | | | | | 2.78 | 0.50 |
| | 7.00 | 0.38 | | | | | 7.00 | 112% | | | | | 3.19 | 0.60 |
| | 8.00 | 0.17 | | | | | 8.00 | 116% | | | | | 3.60 | 0.70 |
| | 9.00 | 0.00 | | | | | | | | | | | 4.01 | 0.80 |
| **2+3+4+5 Zip1 lines+partial lines** | 0.00 | 0.00 | 0.98 | 5.63 | 0.00 | 0.00 | 0.00 | 0% | 2.05 | 7.68 | 0.00 | 0.00 | 0.41 | 0.10 |
| | 3.00 | 0.94 | | | 4.00 | 0.96 | 3.00 | 73% | | | 4.00 | 0.98 | 0.82 | 0.20 |
| | 4.00 | 0.96 | | | | | 4.00 | 98% | | | | | 1.23 | 0.30 |
| | 5.00 | 0.95 | | | | | 5.00 | | | | | | 1.64 | 0.40 |
| | 6.00 | 0.74 | | | | | 6.00 | | | | | | 2.05 | 0.50 |
| | 7.00 | 0.69 | | | | | 7.00 | | | | | | 2.46 | 0.60 |
| | 8.00 | 0.30 | | | | | 8.00 | | | | | | 2.87 | 0.70 |
| | 9.00 | 0.00 | | | | | | | | | | | 3.27 | 0.80 |
| **Entry 2,3,4,5 Zip1** | 0.00 | 0.00 | 0.96 | 3.46 | 0.00 | 0.00 | 0.00 | 0% | 1.66 | 5.12 | 0.00 | 0.00 | 0.31 | 0.10 |
| | 3.00 | 0.94 | | | 0.54 | 0.17 | 3.00 | 89% | | | 0.54 | 0.18 | 0.61 | 0.20 |
| | 4.00 | 0.96 | | | 4.00 | 1.13 | 4.00 | 118% | | | 4.00 | 1.18 | 0.92 | 0.30 |
| | 5.00 | 0.95 | | | | | | | | | | | 1.22 | 0.40 |
| **MI tc72-avg1.+3. WT (23C)** | 0.00 | 0.00 | 0.98 | | 0.00 | 0.00 | 0.00 | 0% | 8.57 | | 0.00 | 0.00 | 6.75 | 0.10 |
| | 3.00 | 0.00 | | | 3.00 | 0.00 | 3.00 | 0% | | | 3.00 | 0.00 | 7.27 | 0.20 |
| | 4.00 | 0.00 | | | 4.00 | 0.00 | 4.00 | 0% | | | 4.00 | 0.00 | 7.69 | 0.30 |
| | 5.00 | 0.00 | | | 5.00 | 0.00 | 5.00 | 0% | | | 5.00 | 0.00 | 8.11 | 0.40 |
| | 6.00 | 0.00 | | | 6.00 | 0.00 | 6.00 | 0% | | | 6.00 | 0.00 | 8.57 | 0.50 |
| | 7.00 | 0.13 | | | 7.00 | 0.13 | 7.00 | 13% | | | 7.00 | 0.13 | 9.05 | 0.60 |
| | 8.00 | 0.37 | | | 8.00 | 0.37 | 8.00 | 38% | | | 8.00 | 0.38 | 9.84 | 0.70 |
| | 9.00 | 0.58 | | | 9.00 | 0.58 | 9.00 | 59% | | | 9.00 | 0.59 | 10.62 | 0.80 |
| | 11.00 | 0.83 | | | 11.00 | 0.83 | 11.00 | 85% | | | 11.00 | 0.85 | | |
| | 12.00 | 0.91 | | | 12.00 | 0.91 | 12.00 | 93% | | | 12.00 | 0.93 | | |
| | 13.00 | 0.98 | | | 13.00 | 0.98 | 13.00 | 100% | | | 13.00 | 1.00 | | |
| **End** | | | | | | | | | | | | | | |

| Input | | | | LifeSpan | Cumulative Values: | | | | Lifespan @ the 50 percentile | | Percentage of maximum | | In 10% increments | |
|---|---|---|---|---|---|---|---|---|---|---|---|---|---|---|
| Event Class | Measurements Time (Hrs) | Measurements Units | Total # Events Units | Time (Hrs) | at interpolated timepoints Time (Hrs) | at interpolated timepoints Units | at taken timepoints Time (Hrs) | at taken timepoints Units | 0.50 Entry | 0.50 Exit | Time (Hrs) | Units | Time (Hrs) | Units |
| CO-72-avg11.+13. WT (30C) | 0.00 | 0.00 | 0.20 | | 0.00 | 0.00 | 0.00 | 2% | 5.36 | | 0.00 | 0.02 | 4.11 | 0.10 |
| | 3.00 | 0.01 | | | 3.00 | 0.01 | 3.00 | 4% | | | 3.00 | 0.04 | 4.42 | 0.20 |
| | 4.00 | 0.01 | | | 4.00 | 0.01 | 4.00 | 7% | | | 4.00 | 0.07 | 4.74 | 0.30 |
| | 5.00 | 0.08 | | | 5.00 | 0.08 | 5.00 | 38% | | | 5.00 | 0.38 | 5.05 | 0.40 |
| | 6.00 | 0.14 | | | 6.00 | 0.14 | 6.00 | 71% | | | 6.00 | 0.71 | 5.36 | 0.50 |
| | 7.00 | 0.18 | | | 7.00 | 0.18 | 7.00 | 89% | | | 7.00 | 0.89 | 5.67 | 0.60 |
| | 8.00 | 0.19 | | | 8.00 | 0.19 | 8.00 | 96% | | | 8.00 | 0.96 | 5.98 | 0.70 |
| | 9.00 | 0.19 | | | 9.00 | 0.19 | 9.00 | 97% | | | 9.00 | 0.97 | 6.52 | 0.80 |
| DSB-72-avg11.+13. WT (30C) | 0.00 | 0.00 | 0.20 | 0.81 | 0.00 | 0.00 | 0.00 | 0% | 3.13 | 3.95 | 0.00 | 0.00 | 1.14 | 0.10 |
| | 3.00 | 0.04 | | | 0.76 | 0.01 | 3.00 | 47% | | | 0.76 | 0.05 | 1.80 | 0.20 |
| | 4.00 | 0.04 | | | 1.57 | 0.03 | 4.00 | 72% | | | 1.57 | 0.15 | 2.32 | 0.30 |
| | 5.00 | 0.03 | | | 2.38 | 0.06 | 5.00 | 90% | | | 2.38 | 0.31 | 2.73 | 0.40 |
| | 6.00 | 0.01 | | | 3.19 | 0.10 | 6.00 | 98% | | | 3.19 | 0.51 | 3.13 | 0.50 |
| | 7.00 | 0.00 | | | 4.00 | 0.14 | 7.00 | 100% | | | 4.00 | 0.72 | 3.52 | 0.60 |
| | 8.00 | 0.00 | | | 4.81 | 0.18 | | | | | 4.81 | 0.88 | 3.91 | 0.70 |
| | | | | | 5.62 | 0.19 | | | | | 5.62 | 0.96 | 4.40 | 0.80 |
| | | | | | 6.43 | 0.20 | | | | | 6.43 | 1.00 | | |
| | | | | | 7.24 | 0.20 | | | | | 7.24 | 1.00 | | |
| SEI-72-avg11.+13. WT (30C) | 0.00 | 0.00 | 0.10 | 0.27 | 0.00 | 0.00 | 0.00 | 1% | 4.87 | 513% | 0.00 | 0.01 | 3.06 | 0.10 |
| | 3.00 | 0.00 | | | 0.22 | 0.00 | 3.00 | 10% | | | 0.22 | 0.01 | 3.77 | 0.20 |
| | 4.00 | 0.01 | | | 0.48 | 0.00 | 4.00 | 25% | | | 0.48 | 0.02 | 4.18 | 0.30 |
| | 5.00 | 0.01 | | | 0.75 | 0.00 | 5.00 | 54% | | | 0.75 | 0.03 | 4.54 | 0.40 |
| | 6.00 | 0.00 | | | 1.01 | 0.00 | 6.00 | 75% | | | 1.01 | 0.04 | 4.87 | 0.50 |
| | 7.00 | 0.00 | | | 1.28 | 0.00 | 7.00 | 87% | | | 1.28 | 0.04 | 5.21 | 0.60 |
| | 8.00 | 0.00 | | | 1.55 | 0.01 | 8.00 | 95% | | | 1.55 | 0.05 | 5.67 | 0.70 |
| | 9.00 | 0.00 | | | 1.81 | 0.01 | | | | | 1.81 | 0.06 | 6.35 | 0.80 |
| | | | | | 2.08 | 0.01 | | | | | 2.08 | 0.07 | | |
| | | | | | 2.34 | 0.01 | | | | | 2.34 | 0.07 | | |
| | | | | | 2.61 | 0.01 | | | | | 2.61 | 0.08 | | |
| | | | | | 2.87 | 0.01 | | | | | 2.87 | 0.09 | | |
| | | | | | 3.14 | 0.01 | | | | | 3.14 | 0.10 | | |
| | | | | | 3.41 | 0.01 | | | | | 3.41 | 0.13 | | |
| | | | | | 3.67 | 0.02 | | | | | 3.67 | 0.18 | | |
| | | | | | 3.94 | 0.02 | | | | | 3.94 | 0.24 | | |
| | | | | | 4.20 | 0.03 | | | | | 4.20 | 0.30 | | |
| | | | | | 4.47 | 0.04 | | | | | 4.47 | 0.38 | | |
| | | | | | 4.73 | 0.05 | | | | | 4.73 | 0.46 | | |
| | | | | | 5.00 | 0.05 | | | | | 5.00 | 0.54 | | |
| | | | | | 5.27 | 0.06 | | | | | 5.27 | 0.61 | | |
| | | | | | 5.53 | 0.07 | | | | | 5.53 | 0.68 | | |
| | | | | | 5.80 | 0.07 | | | | | 5.80 | 0.72 | | |
| | | | | | 6.06 | 0.08 | | | | | 6.06 | 0.76 | | |
| | | | | | 6.33 | 0.08 | | | | | 6.33 | 0.80 | | |
| | | | | | 6.59 | 0.08 | | | | | 6.59 | 0.83 | | |
| | | | | | 6.86 | 0.09 | | | | | 6.86 | 0.86 | | |
| | | | | | 7.13 | 0.09 | | | | | 7.13 | 0.88 | | |
| | | | | | 7.39 | 0.09 | | | | | 7.39 | 0.90 | | |
| | | | | | 7.66 | 0.09 | | | | | 7.66 | 0.93 | | |
| | | | | | 7.92 | 0.09 | | | | | 7.92 | 0.94 | | |
| | | | | | 8.19 | 0.10 | | | | | 8.19 | 0.96 | | |
| | | | | | 8.45 | 0.10 | | | | | 8.45 | 0.98 | | |
| | | | | | 8.72 | 0.10 | | | | | 8.72 | 1.00 | | |
| | | | | | 8.99 | 0.10 | | | | | 8.99 | 1.02 | | |
| IH-dHJ-72-avg11.+13. WT (30C) | 0.00 | 0.00 | 0.20 | 0.12 | 0.00 | 0.00 | 0.00 | 0% | 5.13 | 524% | 0.00 | 0.00 | 3.65 | 0.10 |
| | 3.00 | 0.00 | | | 0.01 | 0.00 | 3.00 | 4% | | | 0.01 | 0.00 | 4.16 | 0.20 |
| | 4.00 | 0.00 | | | 0.13 | 0.00 | 4.00 | 16% | | | 0.13 | 0.00 | 4.53 | 0.30 |
| | 5.00 | 0.01 | | | 0.25 | 0.00 | 5.00 | 46% | | | 0.25 | 0.01 | 4.84 | 0.40 |
| | 6.00 | 0.00 | | | 0.37 | 0.00 | 6.00 | 73% | | | 0.37 | 0.01 | 5.13 | 0.50 |
| | 7.00 | 0.00 | | | 0.49 | 0.00 | 7.00 | 87% | | | 0.49 | 0.01 | 5.45 | 0.60 |
| | 8.00 | 0.00 | | | 0.61 | 0.00 | 8.00 | 95% | | | 0.61 | 0.01 | 5.86 | 0.70 |
| | 9.00 | 0.00 | | | 0.72 | 0.00 | | | | | 0.72 | 0.01 | 6.43 | 0.80 |
| | | | | | 0.84 | 0.00 | | | | | 0.84 | 0.01 | | |
| | | | | | 0.96 | 0.00 | | | | | 0.96 | 0.02 | | |
| | | | | | 1.08 | 0.00 | | | | | 1.08 | 0.02 | | |
| | | | | | 1.20 | 0.00 | | | | | 1.20 | 0.02 | | |
| | | | | | 1.32 | 0.00 | | | | | 1.32 | 0.02 | | |
| | | | | | 1.44 | 0.00 | | | | | 1.44 | 0.02 | | |
| | | | | | 1.56 | 0.00 | | | | | 1.56 | 0.02 | | |
| | | | | | 1.67 | 0.00 | | | | | 1.67 | 0.02 | | |
| | | | | | 1.79 | 0.01 | | | | | 1.79 | 0.03 | | |
| | | | | | 1.91 | 0.01 | | | | | 1.91 | 0.03 | | |
| | | | | | 2.03 | 0.01 | | | | | 2.03 | 0.03 | | |
| | | | | | 2.15 | 0.01 | | | | | 2.15 | 0.03 | | |
| | | | | | 2.27 | 0.01 | | | | | 2.27 | 0.03 | | |
| | | | | | 2.39 | 0.01 | | | | | 2.39 | 0.03 | | |
| | | | | | 2.51 | 0.01 | | | | | 2.51 | 0.04 | | |
| | | | | | 2.62 | 0.01 | | | | | 2.62 | 0.04 | | |
| | | | | | 2.74 | 0.01 | | | | | 2.74 | 0.04 | | |
| | | | | | 2.86 | 0.01 | | | | | 2.86 | 0.04 | | |
| | | | | | 2.98 | 0.01 | | | | | 2.98 | 0.04 | | |
| | | | | | 3.10 | 0.01 | | | | | 3.10 | 0.05 | | |
| | | | | | 3.22 | 0.01 | | | | | 3.22 | 0.05 | | |
| | | | | | 3.34 | 0.01 | | | | | 3.34 | 0.06 | | |
| | | | | | 3.46 | 0.01 | | | | | 3.46 | 0.07 | | |
| | | | | | 3.57 | 0.02 | | | | | 3.57 | 0.09 | | |
| | | | | | 3.69 | 0.02 | | | | | 3.69 | 0.11 | | |
| | | | | | 3.81 | 0.03 | | | | | 3.81 | 0.13 | | |
| | | | | | 3.93 | 0.03 | | | | | 3.93 | 0.15 | | |
| | | | | | 4.05 | 0.03 | | | | | 4.05 | 0.17 | | |
| | | | | | 4.17 | 0.04 | | | | | 4.17 | 0.20 | | |
| | | | | | 4.29 | 0.05 | | | | | 4.29 | 0.23 | | |
| | | | | | 4.41 | 0.05 | | | | | 4.41 | 0.26 | | |
| | | | | | 4.52 | 0.06 | | | | | 4.52 | 0.30 | | |
| | | | | | 4.64 | 0.07 | | | | | 4.64 | 0.33 | | |
| | | | | | 4.76 | 0.07 | | | | | 4.76 | 0.37 | | |
| | | | | | 4.88 | 0.08 | | | | | 4.88 | 0.41 | | |
| | | | | | 5.00 | 0.09 | | | | | 5.00 | 0.46 | | |
| | | | | | 5.12 | 0.10 | | | | | 5.12 | 0.50 | | |
| | | | | | 5.24 | 0.11 | | | | | 5.24 | 0.54 | | |
| | | | | | 5.36 | 0.11 | | | | | 5.36 | 0.57 | | |
| | | | | | 5.48 | 0.12 | | | | | 5.48 | 0.61 | | |
| | | | | | 5.59 | 0.13 | | | | | 5.59 | 0.64 | | |
| | | | | | 5.71 | 0.13 | | | | | 5.71 | 0.67 | | |
| | | | | | 5.83 | 0.14 | | | | | 5.83 | 0.69 | | |
| | | | | | 5.95 | 0.14 | | | | | 5.95 | 0.72 | | |

| | | | | | | | | | | | | | | |
|---|---|---|---|---|---|---|---|---|---|---|---|---|---|---|
| | | | | | 6.07 | 0.15 | | | | | 6.07 | 0.74 | | |
| | | | | | 6.19 | 0.15 | | | | | 6.19 | 0.76 | | |
| | | | | | 6.31 | 0.16 | | | | | 6.31 | 0.78 | | |
| | | | | | 6.43 | 0.16 | | | | | 6.43 | 0.80 | | |
| | | | | | 6.54 | 0.16 | | | | | 6.54 | 0.82 | | |
| | | | | | 6.66 | 0.17 | | | | | 6.66 | 0.83 | | |
| | | | | | 6.78 | 0.17 | | | | | 6.78 | 0.85 | | |
| | | | | | 6.90 | 0.17 | | | | | 6.90 | 0.86 | | |
| | | | | | 7.02 | 0.17 | | | | | 7.02 | 0.87 | | |
| | | | | | 7.14 | 0.18 | | | | | 7.14 | 0.88 | | |
| | | | | | 7.26 | 0.18 | | | | | 7.26 | 0.89 | | |
| | | | | | 7.38 | 0.18 | | | | | 7.38 | 0.91 | | |
| | | | | | 7.49 | 0.18 | | | | | 7.49 | 0.92 | | |
| | | | | | 7.61 | 0.18 | | | | | 7.61 | 0.92 | | |
| | | | | | 7.73 | 0.19 | | | | | 7.73 | 0.93 | | |
| | | | | | 7.85 | 0.19 | | | | | 7.85 | 0.94 | | |
| | | | | | 7.97 | 0.19 | | | | | 7.97 | 0.95 | | |
| | | | | | 8.09 | 0.19 | | | | | 8.09 | 0.96 | | |
| | | | | | 8.21 | 0.19 | | | | | 8.21 | 0.97 | | |
| | | | | | 8.33 | 0.19 | | | | | 8.33 | 0.97 | | |
| | | | | | 8.44 | 0.20 | | | | | 8.44 | 0.98 | | |
| | | | | | 8.56 | 0.20 | | | | | 8.56 | 0.98 | | |
| | | | | | 8.68 | 0.20 | | | | | 8.68 | 0.99 | | |
| | | | | | 8.80 | 0.20 | | | | | 8.80 | 1.00 | | |
| | | | | | 8.92 | 0.20 | | | | | 8.92 | 1.00 | | |
| **5 Zip1 lines** | 0.00 | 0.00 | 0.95 | 0.34 | 0.00 | 0.00 | 0.00 | 0% | 4.75 | 5.09 | 0.00 | 0.00 | 3.45 | 0.10 |
| | 3.00 | 0.00 | | | 0.24 | 0.00 | 3.00 | 2% | | | 0.24 | 0.00 | 3.77 | 0.20 |
| | 4.00 | 0.13 | | | 0.59 | 0.00 | 4.00 | 29% | | | 0.59 | 0.00 | 4.03 | 0.30 |
| | 5.00 | 0.06 | | | 0.93 | 0.00 | 5.00 | 55% | | | 0.93 | 0.00 | 4.34 | 0.40 |
| | 6.00 | 0.07 | | | 1.27 | 0.00 | 6.00 | 75% | | | 1.27 | 0.00 | 4.75 | 0.50 |
| | 7.00 | 0.05 | | | 1.61 | 0.01 | 7.00 | 92% | | | 1.61 | 0.01 | 5.26 | 0.60 |
| | 8.00 | 0.01 | | | 1.95 | 0.01 | 8.00 | 99% | | | 1.95 | 0.01 | 5.75 | 0.70 |
| | 9.00 | 0.00 | | | 2.29 | 0.01 | | | | | 2.29 | 0.01 | 6.25 | 0.80 |
| | | | | | 2.63 | 0.02 | | | | | 2.63 | 0.02 | | |
| | | | | | 2.98 | 0.02 | | | | | 2.98 | 0.02 | | |
| | | | | | 3.32 | 0.06 | | | | | 3.32 | 0.07 | | |
| | | | | | 3.66 | 0.15 | | | | | 3.66 | 0.16 | | |
| | | | | | 4.00 | 0.27 | | | | | 4.00 | 0.29 | | |
| | | | | | 4.34 | 0.38 | | | | | 4.34 | 0.40 | | |
| | | | | | 4.68 | 0.46 | | | | | 4.68 | 0.49 | | |
| | | | | | 5.02 | 0.53 | | | | | 5.02 | 0.55 | | |
| | | | | | 5.37 | 0.59 | | | | | 5.37 | 0.62 | | |
| | | | | | 5.71 | 0.66 | | | | | 5.71 | 0.69 | | |
| | | | | | 6.05 | 0.72 | | | | | 6.05 | 0.76 | | |
| | | | | | 6.39 | 0.79 | | | | | 6.39 | 0.83 | | |
| | | | | | 6.73 | 0.84 | | | | | 6.73 | 0.89 | | |
| | | | | | 7.07 | 0.89 | | | | | 7.07 | 0.93 | | |
| | | | | | 7.41 | 0.92 | | | | | 7.41 | 0.97 | | |
| | | | | | 7.76 | 0.94 | | | | | 7.76 | 0.99 | | |
| | | | | | 8.10 | 0.95 | | | | | 8.10 | 1.00 | | |
| | | | | | 8.44 | 0.95 | | | | | 8.44 | 1.00 | | |
| | | | | | 8.78 | 0.95 | | | | | 8.78 | 1.00 | | |
| **4 Zip1 partial lines** | 0.00 | 0.00 | 0.95 | 0.67 | 0.00 | 0.00 | 0.00 | 0% | 4.13 | 4.80 | 0.00 | 0.00 | 2.26 | 0.10 |
| | 3.00 | 0.06 | | | 0.66 | 0.01 | 3.00 | 19% | | | 0.66 | 0.01 | 3.07 | 0.20 |
| | 4.00 | 0.21 | | | 1.33 | 0.04 | 4.00 | 46% | | | 1.33 | 0.04 | 3.50 | 0.30 |
| | 5.00 | 0.17 | | | 2.00 | 0.08 | 5.00 | 73% | | | 2.00 | 0.08 | 3.81 | 0.40 |
| | 6.00 | 0.08 | | | 2.66 | 0.13 | 6.00 | 88% | | | 2.66 | 0.13 | 4.13 | 0.50 |
| | 7.00 | 0.04 | | | 3.33 | 0.23 | 7.00 | 95% | | | 3.33 | 0.24 | 4.48 | 0.60 |
| | 8.00 | 0.03 | | | 4.00 | 0.44 | 8.00 | 100% | | | 4.00 | 0.46 | 4.88 | 0.70 |
| | 9.00 | 0.00 | | | 4.67 | 0.62 | | | | | 4.67 | 0.65 | 5.34 | 0.80 |
| | | | | | 5.34 | 0.76 | | | | | 5.34 | 0.80 | | |
| | | | | | 6.00 | 0.84 | | | | | 6.00 | 0.88 | | |
| | | | | | 6.67 | 0.89 | | | | | 6.67 | 0.93 | | |
| | | | | | 7.34 | 0.92 | | | | | 7.34 | 0.97 | | |
| | | | | | 8.01 | 0.95 | | | | | 8.01 | 1.00 | | |
| | | | | | 8.68 | 0.96 | | | | | 8.68 | 1.01 | | |
| **3 Zip1 few lines** | 0.00 | 0.00 | 0.95 | 1.86 | 0.00 | 0.00 | 0.00 | 0% | 3.00 | 4.85 | 0.00 | 0.00 | 0.83 | 0.10 |
| | 3.00 | 0.34 | | | 1.29 | 0.15 | 3.00 | 50% | | | 1.29 | 0.16 | 1.51 | 0.20 |
| | 4.00 | 0.42 | | | 3.14 | 0.50 | 4.00 | 74% | | | 3.14 | 0.53 | 2.01 | 0.30 |
| | 5.00 | 0.44 | | | 5.00 | 0.94 | 5.00 | 99% | | | 5.00 | 0.99 | 2.50 | 0.40 |
| | 6.00 | 0.11 | | | 6.86 | 1.03 | 6.00 | 104% | | | 6.86 | 1.09 | 3.00 | 0.50 |
| | 7.00 | 0.09 | | | 8.71 | 1.04 | 7.00 | 109% | | | 8.71 | 1.09 | 3.43 | 0.60 |
| | 8.00 | 0.02 | | | | | 8.00 | 109% | | | | | 3.83 | 0.70 |
| | 9.00 | 0.00 | | | | | | | | | | | 4.23 | 0.80 |
| **2 Zip1 no lines** | 0.00 | 0.00 | 0.95 | 1.53 | 0.00 | 0.00 | 0.00 | 0% | 2.27 | 3.80 | 0.00 | 0.00 | 0.61 | 0.10 |
| | 3.00 | 0.47 | | | 1.47 | 0.23 | 3.00 | 74% | | | 1.47 | 0.24 | 1.21 | 0.20 |
| | 4.00 | 0.20 | | | 3.00 | 0.70 | 4.00 | 85% | | | 3.00 | 0.74 | 1.65 | 0.30 |
| | 5.00 | 0.15 | | | 4.53 | 0.87 | 5.00 | 94% | | | 4.53 | 0.92 | 1.96 | 0.40 |
| | 6.00 | 0.09 | | | 6.06 | 0.96 | 6.00 | 100% | | | 6.06 | 1.01 | 2.27 | 0.50 |
| | 7.00 | 0.06 | | | 7.59 | 0.99 | 7.00 | 103% | | | 7.59 | 1.05 | 2.58 | 0.60 |
| | 8.00 | 0.02 | | | | | 8.00 | | | | | | 2.89 | 0.70 |
| | 9.00 | 0.00 | | | | | | | | | | | 3.54 | 0.80 |
| **2+3+4+5 Zip1 lines+partial lines** | 0.00 | 0.00 | 0.95 | | 0.00 | 0.00 | 0.00 | 0% | 1.63 | | 0.00 | 0.00 | 0.33 | 0.10 |
| | 3.00 | 0.87 | | | 3.00 | 0.87 | 3.00 | 92% | | | 3.00 | 0.92 | 0.65 | 0.20 |
| | 4.00 | 0.95 | | | 4.00 | 0.95 | 4.00 | 100% | | | 4.00 | 1.00 | 0.98 | 0.30 |
| | | | | | | | | | | | | | 1.30 | 0.40 |
| | | | | | | | | | | | | | 1.63 | 0.50 |
| | | | | | | | | | | | | | 1.96 | 0.60 |
| | | | | | | | | | | | | | 2.28 | 0.70 |
| | | | | | | | | | | | | | 2.61 | 0.80 |
| | | | | | | | | | | | | | 0.00 | |
| **4+5 Zip1 lines+partial lines** | 0.00 | 0.00 | 0.95 | 1.01 | 0.00 | 0.00 | 0.00 | 0% | 4.09 | 5.10 | 0.00 | 0.00 | 2.57 | 0.10 |
| | 3.00 | 0.06 | | | 0.97 | 0.02 | 3.00 | 13% | | | 0.97 | 0.02 | 3.20 | 0.20 |
| | 4.00 | 0.33 | | | 1.98 | 0.06 | 4.00 | 48% | | | 1.98 | 0.06 | 3.49 | 0.30 |
| | 5.00 | 0.23 | | | 2.99 | 0.12 | 5.00 | 72% | | | 2.99 | 0.13 | 3.77 | 0.40 |
| | 6.00 | 0.15 | | | 4.00 | 0.45 | 6.00 | 87% | | | 4.00 | 0.48 | 4.09 | 0.50 |
| | 7.00 | 0.09 | | | 5.01 | 0.69 | 7.00 | 96% | | | 5.01 | 0.72 | 4.50 | 0.60 |
| | 8.00 | 0.04 | | | 6.02 | 0.83 | 8.00 | 100% | | | 6.02 | 0.87 | 4.92 | 0.70 |
| | 9.00 | 0.00 | | | 7.03 | 0.92 | | | | | 7.03 | 0.96 | 5.53 | 0.80 |
| | | | | | 8.04 | 0.95 | | | | | 8.04 | 1.00 | | |
| **3+4+5 Zip1 lines+partial lines** | 0.00 | 0.00 | 0.95 | 2.87 | 0.00 | 0.00 | 0.00 | 0% | 2.36 | 5.22 | 0.00 | 0.00 | 0.70 | 0.10 |
| | 3.00 | 0.40 | | | 1.13 | 0.15 | 3.00 | 68% | | | 1.13 | 0.16 | 1.27 | 0.20 |

| | | | | | | | | | | | | | | |
|---|---|---|---|---|---|---|---|---|---|---|---|---|---|---|
| | 4.00 | 0.75 | | | 4.00 | 0.91 | 4.00 | 95% | | | 4.00 | 0.95 | 1.64 | 0.30 |
| | 5.00 | 0.67 | | | 6.87 | 1.09 | 5.00 | 102% | | | 6.87 | 1.15 | 2.00 | 0.40 |
| | 6.00 | 0.26 | | | | | 6.00 | 109% | | | | | 2.36 | 0.50 |
| | 7.00 | 0.17 | | | | | 7.00 | | | | | | 2.72 | 0.60 |
| | 8.00 | 0.05 | | | | | 8.00 | | | | | | 3.08 | 0.70 |
| | 9.00 | 0.00 | | | | | | | | | | | 3.44 | 0.80 |
| **2+3+4+5 Zip1 lines+partial lines** | 0.00 | 0.00 | 0.95 | 4.39 | 0.00 | 0.00 | 0.00 | 0% | 2.00 | 6.39 | 0.00 | 0.00 | 0.40 | 0.10 |
| | 3.00 | 0.87 | | | 4.00 | 0.95 | 3.00 | 75% | | | 4.00 | 1.00 | 0.80 | 0.20 |
| | 4.00 | 0.95 | | | 8.39 | 1.00 | 4.00 | 100% | | | 8.39 | 1.05 | 1.20 | 0.30 |
| | 5.00 | 0.82 | | | | | 5.00 | 101% | | | | | 1.60 | 0.40 |
| | 6.00 | 0.34 | | | | | 6.00 | 102% | | | | | 2.00 | 0.50 |
| | 7.00 | 0.23 | | | | | 7.00 | 103% | | | | | 2.40 | 0.60 |
| | 8.00 | 0.08 | | | | | 8.00 | 104% | | | | | 2.80 | 0.70 |
| | 9.00 | 0.00 | | | | | | | | | | | 3.20 | 0.80 |
| **MI tc72-avg11.+13. WT (30C)** | 0.00 | 0.00 | 0.95 | | 0.00 | 0.00 | 0.00 | 0% | 6.92 | | 0.00 | 0.00 | 5.17 | 0.10 |
| | 3.00 | 0.00 | | | 3.00 | 0.00 | 3.00 | 0% | | | 3.00 | 0.00 | 5.58 | 0.20 |
| | 4.00 | 0.00 | | | 4.00 | 0.00 | 4.00 | 0% | | | 4.00 | 0.00 | 5.99 | 0.30 |
| | 5.00 | 0.06 | | | 5.00 | 0.06 | 5.00 | 6% | | | 5.00 | 0.06 | 6.45 | 0.40 |
| | 6.00 | 0.29 | | | 6.00 | 0.29 | 6.00 | 30% | | | 6.00 | 0.30 | 6.92 | 0.50 |
| | 7.00 | 0.49 | | | 7.00 | 0.49 | 7.00 | 52% | | | 7.00 | 0.52 | 7.38 | 0.60 |
| | 8.00 | 0.70 | | | 8.00 | 0.70 | 8.00 | 74% | | | 8.00 | 0.74 | 7.84 | 0.70 |
| | 9.00 | 0.86 | | | 9.00 | 0.86 | 9.00 | 91% | | | 9.00 | 0.91 | 8.37 | 0.80 |
| | 11.00 | 0.88 | | | 11.00 | 0.88 | 11.00 | 93% | | | 11.00 | 0.93 | | |
| | 12.00 | 0.91 | | | 12.00 | 0.91 | 12.00 | 96% | | | 12.00 | 0.96 | | |
| | 13.00 | 0.95 | | | 13.00 | 0.95 | 13.00 | 100% | | | 13.00 | 1.00 | | |
| **End** | | | | | | | | | | | | | | |

| | Input | | | LifeSpan | Cumulative Values: | | | | Lifespan @ the 50 percentile | | Percentage of maximum | | In 10% increments | |
|---|---|---|---|---|---|---|---|---|---|---|---|---|---|---|
| | Measurements | | Total # Events | | at interpolated timepoints | | at taken timepoints | | 0.50 | 0.50 | | | | |
| Event Class | Time (Hrs) | Units | Units | Time (Hrs) | Time (Hrs) | Units | Time (Hrs) | Units | Entry | Exit | Time (Hrs) | Units | Time (Hrs) | Units |
| CO-tc55-1(pdr5D)-see notes | 0.00 | 0.00 | 0.15 | | 0.00 | 0.00 | 0.00 | 0% | 7.01 | | 0.00 | 0.00 | 5.39 | 0.10 |
| | 3.00 | 0.00 | | | 3.00 | 0.00 | 3.00 | 1% | | | 3.00 | 0.01 | 5.77 | 0.20 |
| | 4.00 | 0.00 | | | 4.00 | 0.00 | 4.00 | 0% | | | 4.00 | 0.00 | 6.17 | 0.30 |
| | 5.00 | 0.00 | | | 5.00 | 0.00 | 5.00 | 0% | | | 5.00 | 0.00 | 6.59 | 0.40 |
| | 6.00 | 0.04 | | | 6.00 | 0.04 | 6.00 | 26% | | | 6.00 | 0.26 | 7.01 | 0.50 |
| | 7.00 | 0.07 | | | 7.00 | 0.07 | 7.00 | 50% | | | 7.00 | 0.50 | 7.43 | 0.60 |
| | 8.50 | 0.12 | | | 8.50 | 0.12 | 8.50 | 81% | | | 8.50 | 0.81 | 7.85 | 0.70 |
| | | | | | | | | | | | | | 8.26 | 0.80 |
| DSB-tc55-1 | 0.00 | 0.00 | 0.13 | 0.85 | 0.00 | 0.00 | 0.00 | 0% | 4.51 | 5.37 | 0.00 | 0.00 | 1.76 | 0.10 |
| | 3.00 | 0.01 | | | 0.73 | 0.00 | 3.00 | 24% | | | 0.73 | 0.03 | 2.67 | 0.20 |
| | 4.00 | 0.02 | | | 1.59 | 0.01 | 4.00 | 40% | | | 1.59 | 0.08 | 3.39 | 0.30 |
| | 5.00 | 0.02 | | | 2.44 | 0.02 | 5.00 | 60% | | | 2.44 | 0.17 | 3.98 | 0.40 |
| | 6.00 | 0.02 | | | 3.29 | 0.04 | 6.00 | 76% | | | 3.29 | 0.28 | 4.51 | 0.50 |
| | 7.00 | 0.01 | | | 4.15 | 0.06 | 7.00 | 87% | | | 4.15 | 0.43 | 5.03 | 0.60 |
| | 8.50 | 0.01 | | | 5.00 | 0.08 | 8.50 | 98% | | | 5.00 | 0.60 | 5.62 | 0.70 |
| | 10.00 | 0.00 | | | 5.85 | 0.10 | | | | | 5.85 | 0.74 | 6.34 | 0.80 |
| | | | | | 6.71 | 0.11 | | | | | 6.71 | 0.85 | | |
| | | | | | 7.56 | 0.12 | | | | | 7.56 | 0.92 | | |
| | | | | | 8.41 | 0.13 | | | | | 8.41 | 0.98 | | |
| | | | | | 9.27 | 0.13 | | | | | 9.27 | 1.00 | | |
| I-tc55-1 (1/3 of IH-dHJ)-ok'd27Dec | 0.00 | 0.00 | 0.08 | 0.35 | 0.00 | 0.00 | 0.00 | 0% | 6.36 | 6.72 | 0.00 | 0.00 | 3.63 | 0.10 |
| | 3.00 | 0.00 | | | 0.34 | 0.00 | 3.00 | 5% | | | 0.34 | 0.00 | 4.45 | 0.20 |
| | 4.00 | 0.00 | | | 0.69 | 0.00 | 4.00 | 15% | | | 0.69 | 0.00 | 5.37 | 0.30 |
| | 5.00 | 0.00 | | | 1.04 | 0.00 | 5.00 | 25% | | | 1.04 | 0.01 | 5.89 | 0.40 |
| | 6.00 | 0.01 | | | 1.40 | 0.00 | 6.00 | 42% | | | 1.40 | 0.01 | 6.36 | 0.50 |
| | 7.00 | 0.01 | | | 1.75 | 0.00 | 7.00 | 63% | | | 1.75 | 0.02 | 6.86 | 0.60 |
| | 8.50 | 0.00 | | | 2.11 | 0.00 | 8.50 | 90% | | | 2.11 | 0.02 | 7.37 | 0.70 |
| | 10.00 | 0.00 | | | 2.46 | 0.00 | | | | | 2.46 | 0.03 | 7.91 | 0.80 |
| | | | | | 2.81 | 0.00 | | | | | 2.81 | 0.04 | | |
| | | | | | 3.17 | 0.00 | | | | | 3.17 | 0.06 | | |
| | | | | | 3.52 | 0.01 | | | | | 3.52 | 0.09 | | |
| | | | | | 3.88 | 0.01 | | | | | 3.88 | 0.13 | | |
| | | | | | 4.23 | 0.01 | | | | | 4.23 | 0.18 | | |
| | | | | | 4.58 | 0.02 | | | | | 4.58 | 0.21 | | |
| | | | | | 4.94 | 0.02 | | | | | 4.94 | 0.24 | | |
| | | | | | 5.29 | 0.02 | | | | | 5.29 | 0.29 | | |
| | | | | | 5.65 | 0.03 | | | | | 5.65 | 0.35 | | |
| | | | | | 6.00 | 0.03 | | | | | 6.00 | 0.42 | | |
| | | | | | 6.35 | 0.04 | | | | | 6.35 | 0.50 | | |
| | | | | | 6.71 | 0.04 | | | | | 6.71 | 0.57 | | |
| | | | | | 7.06 | 0.05 | | | | | 7.06 | 0.64 | | |
| | | | | | 7.42 | 0.05 | | | | | 7.42 | 0.71 | | |
| | | | | | 7.77 | 0.06 | | | | | 7.77 | 0.77 | | |
| | | | | | 8.12 | 0.06 | | | | | 8.12 | 0.84 | | |
| | | | | | 8.48 | 0.07 | | | | | 8.48 | 0.90 | | |
| | | | | | 8.83 | 0.07 | | | | | 8.83 | 0.95 | | |
| | | | | | 9.19 | 0.07 | | | | | 9.19 | 0.98 | | |
| | | | | | 9.54 | 0.07 | | | | | 9.54 | 1.00 | | |
| | | | | | 9.89 | 0.08 | | | | | 9.89 | 1.00 | | |
| IH-dHJ-tc55-1-ok'd27Dec20 | 0.00 | 0.00 | 0.15 | 0.24 | 0.00 | 0.00 | 0.00 | 0% | 6.73 | 6.96 | 0.00 | 0.00 | 3.82 | 0.10 |
| | 3.00 | 0.00 | | | 0.16 | 0.00 | 3.00 | 2% | | | 0.16 | 0.00 | 4.54 | 0.20 |
| | 4.00 | 0.01 | | | 0.40 | 0.00 | 4.00 | 13% | | | 0.40 | 0.00 | 5.64 | 0.30 |
| | 5.00 | 0.00 | | | 0.63 | 0.00 | 5.00 | 23% | | | 0.63 | 0.01 | 6.23 | 0.40 |
| | 6.00 | 0.01 | | | 0.87 | 0.00 | 6.00 | 36% | | | 0.87 | 0.01 | 6.73 | 0.50 |
| | 7.00 | 0.01 | | | 1.11 | 0.00 | 7.00 | 56% | | | 1.11 | 0.01 | 7.19 | 0.60 |
| | 8.50 | 0.01 | | | 1.34 | 0.00 | 8.50 | 87% | | | 1.34 | 0.01 | 7.65 | 0.70 |
| | 10.00 | 0.00 | | | 1.58 | 0.00 | | | | | 1.58 | 0.01 | 8.14 | 0.80 |
| | | | | | 1.81 | 0.00 | | | | | 1.81 | 0.01 | | |
| | | | | | 2.05 | 0.00 | | | | | 2.05 | 0.01 | | |
| | | | | | 2.28 | 0.00 | | | | | 2.28 | 0.01 | | |
| | | | | | 2.52 | 0.00 | | | | | 2.52 | 0.02 | | |
| | | | | | 2.76 | 0.00 | | | | | 2.76 | 0.02 | | |
| | | | | | 2.99 | 0.00 | | | | | 2.99 | 0.02 | | |
| | | | | | 3.23 | 0.00 | | | | | 3.23 | 0.03 | | |
| | | | | | 3.46 | 0.01 | | | | | 3.46 | 0.05 | | |
| | | | | | 3.70 | 0.01 | | | | | 3.70 | 0.08 | | |
| | | | | | 3.93 | 0.02 | | | | | 3.93 | 0.12 | | |
| | | | | | 4.17 | 0.02 | | | | | 4.17 | 0.16 | | |
| | | | | | 4.41 | 0.03 | | | | | 4.41 | 0.19 | | |
| | | | | | 4.64 | 0.03 | | | | | 4.64 | 0.21 | | |
| | | | | | 4.88 | 0.03 | | | | | 4.88 | 0.23 | | |
| | | | | | 5.11 | 0.04 | | | | | 5.11 | 0.24 | | |
| | | | | | 5.35 | 0.04 | | | | | 5.35 | 0.26 | | |
| | | | | | 5.59 | 0.04 | | | | | 5.59 | 0.29 | | |
| | | | | | 5.82 | 0.05 | | | | | 5.82 | 0.33 | | |
| | | | | | 6.06 | 0.06 | | | | | 6.06 | 0.37 | | |
| | | | | | 6.29 | 0.06 | | | | | 6.29 | 0.41 | | |
| | | | | | 6.53 | 0.07 | | | | | 6.53 | 0.46 | | |
| | | | | | 6.76 | 0.08 | | | | | 6.76 | 0.51 | | |
| | | | | | 7.00 | 0.08 | | | | | 7.00 | 0.56 | | |
| | | | | | 7.24 | 0.09 | | | | | 7.24 | 0.61 | | |
| | | | | | 7.47 | 0.10 | | | | | 7.47 | 0.66 | | |
| | | | | | 7.71 | 0.11 | | | | | 7.71 | 0.71 | | |
| | | | | | 7.94 | 0.11 | | | | | 7.94 | 0.76 | | |
| | | | | | 8.18 | 0.12 | | | | | 8.18 | 0.81 | | |
| | | | | | 8.41 | 0.13 | | | | | 8.41 | 0.86 | | |
| | | | | | 8.65 | 0.13 | | | | | 8.65 | 0.90 | | |
| | | | | | 8.89 | 0.14 | | | | | 8.89 | 0.93 | | |
| | | | | | 9.12 | 0.14 | | | | | 9.12 | 0.96 | | |
| | | | | | 9.36 | 0.15 | | | | | 9.36 | 0.98 | | |
| | | | | | 9.59 | 0.15 | | | | | 9.59 | 1.00 | | |
| | | | | | 9.83 | 0.15 | | | | | 9.83 | 1.00 | | |
| Zip3 (>35)-tc55-1 | 0.00 | 0.00 | 0.95 | 2.12 | 0.00 | 0.00 | 0.00 | 0% | 4.38 | 6.50 | 0.00 | 0.00 | 2.96 | 0.10 |
| | 3.00 | 0.06 | | | 0.76 | 0.02 | 3.00 | 11% | | | 0.76 | 0.02 | 3.32 | 0.20 |
| | 4.00 | 0.17 | | | 2.88 | 0.07 | 4.00 | 39% | | | 2.88 | 0.08 | 3.67 | 0.30 |
| | 5.00 | 0.57 | | | 5.00 | 0.64 | 5.00 | 67% | | | 5.00 | 0.67 | 4.03 | 0.40 |
| | 6.00 | 0.52 | | | 7.12 | 1.00 | 6.00 | 85% | | | 7.12 | 1.06 | 4.38 | 0.50 |
| | 7.00 | 0.38 | | | 9.24 | 1.05 | 7.00 | 103% | | | 9.24 | 1.11 | 4.73 | 0.60 |
| | 8.50 | 0.10 | | | | | 8.50 | 109% | | | | | 5.14 | 0.70 |
| | 10.00 | 0.00 | | | | | | | | | | | 5.70 | 0.80 |

| | | | | | | | | | | | | | | |
|---|---|---|---|---|---|---|---|---|---|---|---|---|---|---|
| **Zip3 (22-35)-tc55-1** | 0.00 | 0.00 | 0.95 | 1.77 | 0.00 | 0.00 | 0.00 | 0% | 3.96 | 5.72 | 0.00 | 0.00 | 1.14 | 0.10 |
| | 3.00 | 0.25 | | | 1.47 | 0.12 | 3.00 | 35% | | | 1.47 | 0.13 | 1.95 | 0.20 |
| | 4.00 | 0.24 | | | 3.23 | 0.37 | 4.00 | 51% | | | 3.23 | 0.39 | 2.63 | 0.30 |
| | 5.00 | 0.26 | | | 5.00 | 0.63 | 5.00 | 66% | | | 5.00 | 0.66 | 3.31 | 0.40 |
| | 6.00 | 0.22 | | | 6.77 | 0.76 | 6.00 | 74% | | | 6.77 | 0.80 | 3.96 | 0.50 |
| | 7.00 | 0.11 | | | 8.53 | 0.97 | 7.00 | 83% | | | 8.53 | 1.03 | 4.61 | 0.60 |
| | 8.50 | 0.22 | | | | | 8.50 | 102% | | | | | 5.49 | 0.70 |
| | 10.00 | 0.00 | | | | | | | | | | | 6.74 | 0.80 |
| **Zip3 (10-21)-tc55-1** | 0.00 | 0.02 | 0.95 | 1.81 | 0.00 | 0.02 | 0.00 | 2% | 2.20 | 4.01 | 0.00 | 0.02 | 0.46 | 0.10 |
| | 3.00 | 0.46 | | | 1.19 | 0.22 | 3.00 | 72% | | | 1.19 | 0.23 | 1.03 | 0.20 |
| | 4.00 | 0.38 | | | 3.00 | 0.68 | 4.00 | 81% | | | 3.00 | 0.72 | 1.46 | 0.30 |
| | 5.00 | 0.10 | | | 4.81 | 0.83 | 5.00 | 89% | | | 4.81 | 0.88 | 1.83 | 0.40 |
| | 6.00 | 0.07 | | | 6.62 | 0.90 | 6.00 | 93% | | | 6.62 | 0.95 | 2.20 | 0.50 |
| | 7.00 | 0.06 | | | 8.44 | 0.99 | 7.00 | 97% | | | 8.44 | 1.04 | 2.57 | 0.60 |
| | 8.50 | 0.08 | | | | | 8.50 | | | | | | 2.94 | 0.70 |
| | 10.00 | 0.00 | | | | | | | | | | | 3.94 | 0.80 |
| **ALL-Z3>21** | 0.00 | 0.00 | 0.95 | 3.89 | 0.00 | 0.00 | 0.00 | 0% | 2.80 | 6.69 | 0.00 | 0.00 | 0.92 | 0.10 |
| | 3.00 | 0.31 | | | 1.11 | 0.11 | 3.00 | 54% | | | 1.11 | 0.12 | 1.47 | 0.20 |
| | 4.00 | 0.41 | | | 5.00 | 0.94 | 4.00 | 77% | | | 5.00 | 0.99 | 1.91 | 0.30 |
| | 5.00 | 0.83 | | | 8.89 | 1.18 | 5.00 | 99% | | | 8.89 | 1.24 | 2.36 | 0.40 |
| | 6.00 | 0.75 | | | | | 6.00 | 106% | | | | | 2.80 | 0.50 |
| | 7.00 | 0.49 | | | | | 7.00 | 112% | | | | | 3.25 | 0.60 |
| | 8.50 | 0.32 | | | | | 8.50 | 121% | | | | | 3.70 | 0.70 |
| | 10.00 | 0.00 | | | | | | | | | | | 4.14 | 0.80 |
| **ALL-Z3>9** | 0.00 | 0.02 | 0.95 | 5.70 | 0.00 | 0.02 | 0.00 | 2% | 2.45 | 8.14 | 0.00 | 0.02 | 0.40 | 0.10 |
| | 3.00 | 0.77 | | | 5.00 | 0.95 | 3.00 | 61% | | | 5.00 | 1.00 | 0.92 | 0.20 |
| | 4.00 | 0.79 | | | | | 4.00 | 80% | | | | | 1.43 | 0.30 |
| | 5.00 | 0.93 | | | | | 5.00 | 100% | | | | | 1.94 | 0.40 |
| | 6.00 | 0.82 | | | | | 6.00 | | | | | | 2.45 | 0.50 |
| | 7.00 | 0.56 | | | | | 7.00 | | | | | | 2.96 | 0.60 |
| | 8.50 | 0.40 | | | | | 8.50 | | | | | | 3.47 | 0.70 |
| | 10.00 | 0.00 | | | | | | | | | | | 3.98 | 0.80 |
| **ALL-Z3>9** | 0.00 | 0.02 | 0.95 | | 0.00 | 0.02 | 0.00 | 2% | 1.81 | | 0.00 | 0.02 | 0.30 | 0.10 |
| | 3.00 | 0.77 | | | 3.00 | 0.77 | 3.00 | 81% | | | 3.00 | 0.81 | 0.68 | 0.20 |
| | 4.00 | 0.79 | | | 4.00 | 0.79 | 4.00 | 83% | | | 4.00 | 0.83 | 1.06 | 0.30 |
| | 5.00 | 0.93 | | | 5.00 | 0.93 | 5.00 | 98% | | | 5.00 | 0.98 | 1.43 | 0.40 |
| | | | | | | | | | | | | | 1.81 | 0.50 |
| | | | | | | | | | | | | | 2.19 | 0.60 |
| | | | | | | | | | | | | | 2.57 | 0.70 |
| | | | | | | | | | | | | | 2.95 | 0.80 |
| **6. Rec8 lines** | 0.00 | 0.00 | 0.95 | 1.85 | 0.00 | 0.00 | 0.00 | 0% | 5.33 | 7.18 | 0.00 | 0.00 | 2.34 | 0.10 |
| | 3.00 | 0.10 | | | 0.45 | 0.01 | 3.00 | 16% | | | 0.45 | 0.02 | 3.38 | 0.20 |
| | 4.00 | 0.15 | | | 2.30 | 0.09 | 4.00 | 26% | | | 2.30 | 0.10 | 4.29 | 0.30 |
| | 5.00 | 0.29 | | | 4.15 | 0.26 | 5.00 | 44% | | | 4.15 | 0.27 | 4.81 | 0.40 |
| | 6.00 | 0.34 | | | 6.00 | 0.60 | 6.00 | 63% | | | 6.00 | 0.63 | 5.33 | 0.50 |
| | 7.00 | 0.27 | | | 7.85 | 0.88 | 7.00 | 79% | | | 7.85 | 0.93 | 5.85 | 0.60 |
| | 8.50 | 0.30 | | | 9.70 | 0.94 | 8.50 | 95% | | | 9.70 | 0.99 | 6.45 | 0.70 |
| | 10.00 | 0.00 | | | | | | | | | | | 7.06 | 0.80 |
| **5+6 Rec8** | 0.00 | 0.00 | 0.95 | 3.20 | 0.00 | 0.00 | 0.00 | 0% | 3.94 | 7.15 | 0.00 | 0.00 | 1.02 | 0.10 |
| | 3.00 | 0.28 | | | 2.80 | 0.26 | 3.00 | 31% | | | 2.80 | 0.27 | 2.04 | 0.20 |
| | 4.00 | 0.45 | | | 6.00 | 0.86 | 4.00 | 51% | | | 6.00 | 0.90 | 2.93 | 0.30 |
| | 5.00 | 0.50 | | | 9.20 | 1.04 | 5.00 | 71% | | | 9.20 | 1.09 | 3.43 | 0.40 |
| | 6.00 | 0.60 | | | | | 6.00 | 90% | | | | | 3.94 | 0.50 |
| | 7.00 | 0.34 | | | | | 7.00 | 96% | | | | | 4.45 | 0.60 |
| | 8.50 | 0.34 | | | | | 8.50 | 105% | | | | | 4.96 | 0.70 |
| | 10.00 | 0.00 | | | | | | | | | | | 5.47 | 0.80 |
| **4+5+6 Rec8** | 0.00 | 0.00 | 0.95 | 4.09 | 0.00 | 0.00 | 0.00 | 0% | 3.25 | 7.35 | 0.00 | 0.00 | 0.75 | 0.10 |
| | 3.00 | 0.38 | | | 1.91 | 0.24 | 3.00 | 45% | | | 1.91 | 0.25 | 1.50 | 0.20 |
| | 4.00 | 0.62 | | | 6.00 | 0.95 | 4.00 | 64% | | | 6.00 | 1.00 | 2.25 | 0.30 |
| | 5.00 | 0.65 | | | | | 5.00 | 82% | | | | | 3.00 | 0.40 |
| | 6.00 | 0.71 | | | | | 6.00 | 100% | | | | | 3.75 | 0.50 |
| | 7.00 | 0.42 | | | | | 7.00 | | | | | | 4.50 | 0.60 |
| | 8.50 | 0.42 | | | | | 8.50 | | | | | | 5.25 | 0.70 |
| | 10.00 | 0.00 | | | | | | | | | | | 6.00 | 0.80 |
| **5 Zip1 lines** | 0.00 | 0.00 | 0.95 | 0.64 | 0.00 | 0.00 | 0.00 | 0% | 5.92 | 6.56 | 0.00 | 0.00 | 4.85 | 0.10 |
| | 3.00 | 0.00 | | | 0.23 | 0.00 | 3.00 | 0% | | | 0.23 | 0.00 | 5.25 | 0.20 |
| | 4.00 | 0.01 | | | 0.87 | 0.00 | 4.00 | 1% | | | 0.87 | 0.00 | 5.51 | 0.30 |
| | 5.00 | 0.07 | | | 1.52 | 0.00 | 5.00 | 14% | | | 1.52 | 0.00 | 5.71 | 0.40 |
| | 6.00 | 0.30 | | | 2.16 | 0.00 | 6.00 | 54% | | | 2.16 | 0.00 | 5.92 | 0.50 |
| | 7.00 | 0.16 | | | 2.80 | 0.00 | 7.00 | 84% | | | 2.80 | 0.00 | 6.17 | 0.60 |
| | 8.50 | 0.02 | | | 3.44 | 0.00 | 8.50 | 101% | | | 3.44 | 0.00 | 6.46 | 0.70 |
| | 10.00 | 0.00 | | | 4.08 | 0.02 | | | | | 4.08 | 0.02 | 6.81 | 0.80 |
| | | | | | 4.72 | 0.07 | | | | | 4.72 | 0.07 | | |
| | | | | | 5.36 | 0.21 | | | | | 5.36 | 0.23 | | |
| | | | | | 6.00 | 0.51 | | | | | 6.00 | 0.54 | | |
| | | | | | 6.64 | 0.72 | | | | | 6.64 | 0.76 | | |
| | | | | | 7.28 | 0.86 | | | | | 7.28 | 0.90 | | |
| | | | | | 7.92 | 0.94 | | | | | 7.92 | 0.98 | | |
| | | | | | 8.56 | 0.96 | | | | | 8.56 | 1.01 | | |
| | | | | | 9.20 | 0.97 | | | | | 9.20 | 1.02 | | |
| | | | | | 9.84 | 0.97 | | | | | 9.84 | 1.02 | | |
| **4 Zip1 partial lines** | 0.00 | 0.00 | 0.95 | 0.66 | 0.00 | 0.00 | 0.00 | 0% | 6.04 | 6.70 | 0.00 | 0.00 | 4.15 | 0.10 |
| | 3.00 | 0.00 | | | 0.40 | 0.00 | 3.00 | 0% | | | 0.40 | 0.00 | 4.62 | 0.20 |
| | 4.00 | 0.05 | | | 1.06 | 0.00 | 4.00 | 8% | | | 1.06 | 0.00 | 5.00 | 0.30 |
| | 5.00 | 0.16 | | | 1.72 | 0.00 | 5.00 | 30% | | | 1.72 | 0.00 | 5.50 | 0.40 |
| | 6.00 | 0.10 | | | 2.37 | 0.00 | 6.00 | 49% | | | 2.37 | 0.00 | 6.04 | 0.50 |
| | 7.00 | 0.13 | | | 3.03 | 0.00 | 7.00 | 69% | | | 3.03 | 0.00 | 6.56 | 0.60 |
| | 8.50 | 0.10 | | | 3.69 | 0.03 | 8.50 | 94% | | | 3.69 | 0.04 | 7.06 | 0.70 |
| | 10.00 | 0.00 | | | 4.34 | 0.12 | | | | | 4.34 | 0.13 | 7.60 | 0.80 |
| | | | | | 5.00 | 0.28 | | | | | 5.00 | 0.30 | | |
| | | | | | 5.66 | 0.41 | | | | | 5.66 | 0.43 | | |
| | | | | | 6.31 | 0.52 | | | | | 6.31 | 0.55 | | |
| | | | | | 6.97 | 0.65 | | | | | 6.97 | 0.68 | | |
| | | | | | 7.63 | 0.77 | | | | | 7.63 | 0.81 | | |
| | | | | | 8.28 | 0.87 | | | | | 8.28 | 0.91 | | |
| | | | | | 8.94 | 0.94 | | | | | 8.94 | 0.99 | | |
| | | | | | 9.60 | 0.96 | | | | | 9.60 | 1.02 | | |
| **3 Zip1 dots+fe lines** | 0.00 | 0.04 | 0.95 | 1.26 | 0.00 | 0.04 | 0.00 | 4% | 4.46 | 5.71 | 0.00 | 0.04 | 1.22 | 0.10 |
| | 3.00 | 0.08 | | | 1.23 | 0.10 | 3.00 | 25% | | | 1.23 | 0.10 | 2.64 | 0.20 |
| | 4.00 | 0.18 | | | 2.49 | 0.17 | 4.00 | 40% | | | 2.49 | 0.18 | 3.41 | 0.30 |
| | 5.00 | 0.26 | | | 3.74 | 0.33 | 5.00 | 62% | | | 3.74 | 0.34 | 4.00 | 0.40 |
| | 6.00 | 0.13 | | | 5.00 | 0.59 | 6.00 | 73% | | | 5.00 | 0.62 | 4.46 | 0.50 |

| | | | | | | | | | | | | | | |
|---|---|---|---|---|---|---|---|---|---|---|---|---|---|---|
| | 7.00 | 0.13 | | | 6.26 | 0.72 | 7.00 | 84% | | | 6.26 | 0.76 | 4.91 | 0.60 |
| | 8.50 | 0.16 | | | 7.51 | 0.86 | 8.50 | 101% | | | 7.51 | 0.90 | 5.73 | 0.70 |
| | 10.00 | 0.00 | | | 8.77 | 0.99 | | | | | 8.77 | 1.04 | 6.61 | 0.80 |
| **2 Zip1 dots** | 0.00 | 0.08 | 0.95 | 2.39 | 0.00 | 0.08 | 0.00 | 8% | 2.12 | 4.51 | 0.00 | 0.08 | 0.04 | 0.10 |
| | 3.00 | 0.42 | | | 0.23 | 0.18 | 3.00 | 66% | | | 0.23 | 0.19 | 0.29 | 0.20 |
| | 4.00 | 0.42 | | | 2.61 | 0.55 | 4.00 | 85% | | | 2.61 | 0.58 | 0.90 | 0.30 |
| | 5.00 | 0.44 | | | 5.00 | 0.99 | 5.00 | 105% | | | 5.00 | 1.05 | 1.51 | 0.40 |
| | 6.00 | 0.18 | | | 7.39 | 1.10 | 6.00 | 109% | | | 7.39 | 1.16 | 2.12 | 0.50 |
| | 7.00 | 0.11 | | | 9.77 | 1.11 | 7.00 | 114% | | | 9.77 | 1.17 | 2.71 | 0.60 |
| | 8.50 | 0.09 | | | | | 8.50 | 116% | | | | | 3.22 | 0.70 |
| | 10.00 | 0.00 | | | | | | | | | | | 3.73 | 0.80 |
| **4+5 Zip1 lines+partial lines** | 0.00 | 0.00 | 0.95 | 1.30 | 0.00 | 0.00 | 0.00 | 0% | 5.59 | 6.88 | 0.00 | 0.00 | 3.94 | 0.10 |
| | 3.00 | 0.00 | | | 0.81 | 0.00 | 3.00 | 2% | | | 0.81 | 0.00 | 4.63 | 0.20 |
| | 4.00 | 0.05 | | | 2.11 | 0.00 | 4.00 | 11% | | | 2.11 | 0.00 | 4.98 | 0.30 |
| | 5.00 | 0.23 | | | 3.40 | 0.02 | 5.00 | 31% | | | 3.40 | 0.02 | 5.28 | 0.40 |
| | 6.00 | 0.40 | | | 4.70 | 0.20 | 6.00 | 64% | | | 4.70 | 0.21 | 5.59 | 0.50 |
| | 7.00 | 0.29 | | | 6.00 | 0.60 | 7.00 | 84% | | | 6.00 | 0.64 | 5.89 | 0.60 |
| | 8.50 | 0.12 | | | 7.30 | 0.86 | 8.50 | 102% | | | 7.30 | 0.90 | 6.31 | 0.70 |
| | 10.00 | 0.00 | | | 8.60 | 0.97 | | | | | 8.60 | 1.03 | 6.79 | 0.80 |
| | | | | | 9.89 | 0.98 | | | | | 9.89 | 1.03 | | |
| **3+4+5 Zip1 lines+partial lines** | 0.00 | 0.04 | 0.95 | 2.55 | 0.00 | 0.04 | 0.00 | 4% | 4.55 | 7.11 | 0.00 | 0.04 | 0.97 | 0.10 |
| | 3.00 | 0.08 | | | 0.89 | 0.09 | 3.00 | 23% | | | 0.89 | 0.09 | 2.58 | 0.20 |
| | 4.00 | 0.23 | | | 3.45 | 0.24 | 4.00 | 38% | | | 3.45 | 0.25 | 3.65 | 0.30 |
| | 5.00 | 0.49 | | | 6.00 | 0.78 | 5.00 | 60% | | | 6.00 | 0.82 | 4.10 | 0.40 |
| | 6.00 | 0.54 | | | 8.55 | 1.05 | 6.00 | 82% | | | 8.55 | 1.10 | 4.55 | 0.50 |
| | 7.00 | 0.42 | | | | | 7.00 | 93% | | | | | 5.00 | 0.60 |
| | 8.50 | 0.28 | | | | | 8.50 | 110% | | | | | 5.45 | 0.70 |
| | 10.00 | 0.00 | | | | | | | | | | | 5.90 | 0.80 |
| **2+3+4+5 Zip1 lines+partial lines** | 0.00 | 0.12 | 0.95 | 4.94 | 0.00 | 0.12 | 0.00 | 12% | 1.31 | 6.25 | 0.00 | 0.12 | | 0.10 |
| | 3.00 | 0.50 | | | 0.06 | 0.24 | 3.00 | 84% | | | 0.06 | 0.25 | 0.04 | 0.20 |
| | 4.00 | 0.66 | | | 5.00 | 1.17 | 4.00 | 104% | | | 5.00 | 1.23 | 0.31 | 0.30 |
| | 5.00 | 0.93 | | | 9.94 | 1.19 | 5.00 | 123% | | | 9.94 | 1.25 | 0.81 | 0.40 |
| | 6.00 | 0.72 | | | | | 6.00 | 124% | | | | | 1.31 | 0.50 |
| | 7.00 | 0.52 | | | | | 7.00 | 124% | | | | | 1.81 | 0.60 |
| | 8.50 | 0.37 | | | | | 8.50 | 125% | | | | | 2.32 | 0.70 |
| | 10.00 | 0.00 | | | | | | | | | | | 2.82 | 0.80 |
| **2+3+4+5 Zip1 lines+partial lines** | 0.00 | 0.12 | 0.95 | | 0.00 | 0.12 | 0.00 | 12% | 2.81 | | 0.00 | 0.12 | | 0.10 |
| | 3.00 | 0.50 | | | 3.00 | 0.50 | 3.00 | 53% | | | 3.00 | 0.53 | 0.58 | 0.20 |
| | 4.00 | 0.66 | | | 4.00 | 0.66 | 4.00 | 69% | | | 4.00 | 0.69 | 1.32 | 0.30 |
| | 5.00 | 0.93 | | | 5.00 | 0.93 | 5.00 | 98% | | | 5.00 | 0.98 | 2.06 | 0.40 |
| | | | | | | | | | | | | | 2.81 | 0.50 |
| | | | | | | | | | | | | | 3.45 | 0.60 |
| | | | | | | | | | | | | | 4.06 | 0.70 |
| | | | | | | | | | | | | | 4.66 | 0.80 |
| **MI/II-tc55-1** | 0.00 | 0.00 | 0.95 | | 0.00 | 0.00 | 0.00 | 0% | 5.12 | | 0.00 | 0.00 | 5.02 | 0.10 |
| | 3.00 | 0.00 | | | 3.00 | 0.00 | 3.00 | 0% | | | 3.00 | 0.00 | 5.05 | 0.20 |
| | 4.00 | 0.00 | | | 4.00 | 0.00 | 4.00 | 0% | | | 4.00 | 0.00 | 5.07 | 0.30 |
| | 5.00 | 0.00 | | | 5.00 | 0.00 | 5.00 | 0% | | | 5.00 | 0.00 | 5.09 | 0.40 |
| | 6.00 | 4.10 | | | 6.00 | 4.10 | 6.00 | 431% | | | 6.00 | 4.31 | 5.12 | 0.50 |
| | 7.00 | 12.40 | | | 7.00 | 12.40 | 7.00 | 1305% | | | 7.00 | 13.05 | 5.14 | 0.60 |
| | 8.50 | 32.28 | | | 8.50 | 32.28 | 8.50 | 3398% | | | 8.50 | 33.98 | 5.16 | 0.70 |
| | 11.00 | 84.82 | | | 11.00 | 84.82 | 11.00 | 8929% | | | 11.00 | 89.29 | 5.19 | 0.80 |
| | 13.00 | 100.00 | | | 13.00 | 100.00 | 13.00 | 10526% | | | 13.00 | 105.26 | 5.00 | |
| **End** | | | | | | | | | | | | | | |

| Input | | | | LifeSpan | Cumulative Values: | | | | Lifespan @ the 50 percentile | | Percentage of maximum | | In 10% increments | |
|---|---|---|---|---|---|---|---|---|---|---|---|---|---|---|
| | Measurements | | Total # Events | Time (Hrs) | at interpolated timepoints | | at taken timepoints | | 0.50 | 0.50 | Time (Hrs) | Units | Time (Hrs) | Units |
| Event Class | Time (Hrs) | Units | Units | | Time (Hrs) | Units | Time (Hrs) | Units | Entry | Exit | | | | |
| CO-PADMORE-30C+500mMKCl | 0.00 | 0.00 | 14.00 | | 0.00 | 0.00 | 0.00 | 0% | 7.12 | | 0.00 | 0.00 | 4.81 | 0.10 |
| | 2.00 | 0.00 | | | 2.00 | 0.00 | 2.00 | 0% | | | 2.00 | 0.00 | 5.76 | 0.20 |
| | 3.00 | 0.00 | | | 3.00 | 0.00 | 3.00 | 0% | | | 3.00 | 0.00 | 6.19 | 0.30 |
| | 3.50 | 0.00 | | | 3.50 | 0.00 | 3.50 | 0% | | | 3.50 | 0.00 | 6.74 | 0.40 |
| | 4.00 | 0.00 | | | 4.00 | 0.00 | 4.00 | 0% | | | 4.00 | 0.00 | 7.12 | 0.50 |
| | 4.50 | 0.00 | | | 4.50 | 0.00 | 4.50 | 0% | | | 4.50 | 0.00 | 7.37 | 0.60 |
| | 5.00 | 2.25 | | | 5.00 | 2.25 | 5.00 | 16% | | | 5.00 | 0.16 | 7.63 | 0.70 |
| | 5.50 | 1.80 | | | 5.50 | 1.80 | 5.50 | 13% | | | 5.50 | 0.13 | 7.90 | 0.80 |
| | 6.00 | 3.75 | | | 6.00 | 3.75 | 6.00 | 27% | | | 6.00 | 0.27 | | |
| | 6.50 | 4.95 | | | 6.50 | 4.95 | 6.50 | 35% | | | 6.50 | 0.35 | | |
| | 7.00 | 6.30 | | | 7.00 | 6.30 | 7.00 | 45% | | | 7.00 | 0.45 | | |
| | 7.50 | 9.15 | | | 7.50 | 9.15 | 7.50 | 65% | | | 7.50 | 0.65 | | |
| | 8.00 | 11.70 | | | 8.00 | 11.70 | 8.00 | 84% | | | 8.00 | 0.84 | | |
| | 9.00 | 13.80 | | | 9.00 | 13.80 | 9.00 | 99% | | | 9.00 | 0.99 | | |
| CO as percent of 92% max | 0.00 | 0.00 | 92.00 | | 0.00 | 0.00 | 0.00 | 0% | 7.11 | | 0.00 | 0.00 | 4.81 | 0.10 |
| | 2.00 | 0.00 | | | 2.00 | 0.00 | 2.00 | 0% | | | 2.00 | 0.00 | 5.75 | 0.20 |
| | 3.00 | 0.00 | | | 3.00 | 0.00 | 3.00 | 0% | | | 3.00 | 0.00 | 6.16 | 0.30 |
| | 3.50 | 0.00 | | | 3.50 | 0.00 | 3.50 | 0% | | | 3.50 | 0.00 | 6.71 | 0.40 |
| | 4.00 | 0.00 | | | 4.00 | 0.00 | 4.00 | 0% | | | 4.00 | 0.00 | 7.11 | 0.50 |
| | 4.50 | 0.00 | | | 4.50 | 0.00 | 4.50 | 0% | | | 4.50 | 0.00 | 7.35 | 0.60 |
| | 5.00 | 15.00 | | | 5.00 | 15.00 | 5.00 | 16% | | | 5.00 | 0.16 | 7.60 | 0.70 |
| | 5.50 | 12.00 | | | 5.50 | 12.00 | 5.50 | 13% | | | 5.50 | 0.13 | 7.87 | 0.80 |
| | 6.00 | 25.00 | | | 6.00 | 25.00 | 6.00 | 27% | | | 6.00 | 0.27 | | |
| | 6.50 | 33.00 | | | 6.50 | 33.00 | 6.50 | 36% | | | 6.50 | 0.36 | | |
| | 7.00 | 42.00 | | | 7.00 | 42.00 | 7.00 | 46% | | | 7.00 | 0.46 | | |
| | 7.50 | 61.00 | | | 7.50 | 61.00 | 7.50 | 66% | | | 7.50 | 0.66 | | |
| | 8.00 | 78.00 | | | 8.00 | 78.00 | 8.00 | 85% | | | 8.00 | 0.85 | | |
| | 9.00 | 92.00 | | | 9.00 | 92.00 | 9.00 | 100% | | | 9.00 | 1.00 | | |
| DSB-PADMORE | 0.00 | 0.00 | 15.00 | 1.04 | 0.00 | 0.00 | 0.00 | 0% | 3.79 | 4.83 | 0.00 | 0.00 | 2.03 | 0.10 |
| | 2.00 | 0.50 | | | 0.33 | 0.08 | 2.00 | 10% | | | 0.33 | 0.01 | 2.64 | 0.20 |
| | 3.00 | 3.40 | | | 1.38 | 0.43 | 3.00 | 29% | | | 1.38 | 0.03 | 3.04 | 0.30 |
| | 3.50 | 4.00 | | | 2.42 | 2.14 | 3.50 | 42% | | | 2.42 | 0.14 | 3.44 | 0.40 |
| | 4.00 | 4.20 | | | 3.46 | 6.09 | 4.00 | 56% | | | 3.46 | 0.41 | 3.79 | 0.50 |
| | 4.50 | 4.50 | | | 4.50 | 10.59 | 4.50 | 71% | | | 4.50 | 0.71 | 4.13 | 0.60 |
| | 5.00 | 3.50 | | | 5.54 | 12.89 | 5.00 | 78% | | | 5.54 | 0.86 | 4.48 | 0.70 |
| | 5.50 | 2.30 | | | 6.58 | 15.19 | 5.50 | 85% | | | 6.58 | 1.01 | 5.14 | 0.80 |
| | 6.00 | 2.40 | | | 7.63 | 15.67 | 6.00 | 93% | | | 7.63 | 1.04 | | |
| | 6.50 | 2.70 | | | 8.67 | 15.70 | 6.50 | 100% | | | 8.67 | 1.05 | | |
| | 7.00 | 0.30 | | | 9.71 | 15.70 | 7.00 | 103% | | | 9.71 | 1.05 | | |
| | 7.50 | 0.60 | | | | | 7.50 | 104% | | | | | | |
| | 8.00 | 0.10 | | | | | 8.00 | 105% | | | | | | |
| | 9.00 | 0.00 | | | | | 9.00 | 105% | | | | | | |
| | 10.00 | 0.00 | | | | | | | | | | | | |
| Full SC | 0.00 | 0.00 | 82.00 | 1.46 | 0.00 | 0.00 | 0.00 | 0% | 5.21 | 6.67 | 0.00 | 0.00 | 3.26 | 0.10 |
| | 2.00 | 0.00 | | | 1.37 | 0.00 | 2.00 | 2% | | | 1.37 | 0.00 | 4.13 | 0.20 |
| | 3.00 | 5.00 | | | 2.83 | 4.15 | 3.00 | 7% | | | 2.83 | 0.05 | 4.56 | 0.30 |
| | 3.50 | 5.00 | | | 4.29 | 17.88 | 3.50 | 13% | | | 4.29 | 0.22 | 4.89 | 0.40 |
| | 4.00 | 12.00 | | | 5.75 | 54.38 | 4.00 | 18% | | | 5.75 | 0.66 | 5.21 | 0.50 |
| | 4.50 | 15.00 | | | 7.21 | 74.71 | 4.50 | 28% | | | 7.21 | 0.91 | 5.54 | 0.60 |
| | 5.00 | 20.00 | | | 8.67 | 83.37 | 5.00 | 43% | | | 8.67 | 1.02 | 5.97 | 0.70 |
| | 5.50 | 40.00 | | | | | 5.50 | 59% | | | | | 6.56 | 0.80 |
| | 6.00 | 33.00 | | | | | 6.00 | 71% | | | | | | |
| | 6.50 | 39.00 | | | | | 6.50 | 79% | | | | | | |
| | 7.00 | 30.00 | | | | | 7.00 | 88% | | | | | | |
| | 7.50 | 7.00 | | | | | 7.50 | 93% | | | | | | |
| | 8.00 | 10.00 | | | | | 8.00 | 97% | | | | | | |
| | 9.00 | 8.00 | | | | | 9.00 | | | | | | | |
| | 10.00 | 0.00 | | | | | | | | | | | | |
| SC pre A | 0.00 | 0.00 | 82.00 | 0.63 | 0.00 | 0.00 | 0.00 | 0% | 3.94 | 4.56 | 0.00 | 0.00 | 2.66 | 0.10 |
| | 2.00 | 0.00 | | | 0.61 | 0.00 | 2.00 | 1% | | | 0.61 | 0.00 | 3.07 | 0.20 |
| | 3.00 | 10.00 | | | 1.24 | 0.00 | 3.00 | 18% | | | 1.24 | 0.00 | 3.37 | 0.30 |
| | 3.50 | 20.00 | | | 1.87 | 0.00 | 3.50 | 34% | | | 1.87 | 0.00 | 3.66 | 0.40 |
| | 4.00 | 16.00 | | | 2.49 | 4.94 | 4.00 | 52% | | | 2.49 | 0.06 | 3.94 | 0.50 |
| | 4.50 | 20.00 | | | 3.12 | 17.38 | 4.50 | 69% | | | 3.12 | 0.21 | 4.21 | 0.60 |
| | 5.00 | 10.00 | | | 3.75 | 35.38 | 5.00 | 78% | | | 3.75 | 0.43 | 4.57 | 0.70 |
| | 5.50 | 0.00 | | | 4.38 | 54.40 | 5.50 | 80% | | | 4.38 | 0.66 | 5.39 | 0.80 |
| | 6.00 | 8.00 | | | 5.01 | 64.28 | 6.00 | 84% | | | 5.01 | 0.78 | | |
| | 6.50 | 1.00 | | | 5.63 | 66.43 | 6.50 | 89% | | | 5.63 | 0.81 | | |
| | 7.00 | 7.00 | | | 6.26 | 70.76 | 7.00 | 94% | | | 6.26 | 0.86 | | |
| | 7.50 | 3.00 | | | 6.89 | 76.44 | 7.50 | 97% | | | 6.89 | 0.93 | | |
| | 8.00 | 2.00 | | | 7.52 | 79.40 | 8.00 | 98% | | | 7.52 | 0.97 | | |
| | 9.00 | 0.00 | | | 8.15 | 81.11 | 9.00 | 99% | | | 8.15 | 0.99 | | |
| | 10.00 | 0.00 | | | 8.77 | 81.56 | | | | | 8.77 | 0.99 | | |
| | | | | | 9.40 | 81.56 | | | | | 9.40 | 0.99 | | |
| SC pre B | 0.00 | 0.00 | 82.00 | 0.72 | 0.00 | 0.00 | 0.00 | 0% | 4.38 | 5.10 | 0.00 | 0.00 | 2.87 | 0.10 |
| | 2.00 | 0.00 | | | 0.45 | 0.00 | 2.00 | 1% | | | 0.45 | 0.00 | 3.38 | 0.20 |
| | 3.00 | 7.00 | | | 1.17 | 0.00 | 3.00 | 12% | | | 1.17 | 0.00 | 3.74 | 0.30 |
| | 3.50 | 13.00 | | | 1.88 | 0.00 | 3.50 | 23% | | | 1.88 | 0.00 | 4.09 | 0.40 |
| | 4.00 | 16.00 | | | 2.60 | 4.20 | 4.00 | 37% | | | 2.60 | 0.05 | 4.38 | 0.50 |
| | 4.50 | 20.00 | | | 3.32 | 15.01 | 4.50 | 54% | | | 3.32 | 0.18 | 4.68 | 0.60 |
| | 5.00 | 20.00 | | | 4.03 | 31.28 | 5.00 | 70% | | | 4.03 | 0.38 | 4.99 | 0.70 |
| | 5.50 | 18.00 | | | 4.75 | 51.28 | 5.50 | 85% | | | 4.75 | 0.63 | 5.32 | 0.80 |
| | 6.00 | 8.00 | | | 5.47 | 69.41 | 6.00 | 90% | | | 5.47 | 0.85 | | |
| | 6.50 | 1.00 | | | 6.18 | 74.85 | 6.50 | 94% | | | 6.18 | 0.91 | | |
| | 7.00 | 7.00 | | | 6.90 | 80.64 | 7.00 | 99% | | | 6.90 | 0.98 | | |
| | 7.50 | 4.00 | | | 7.62 | 83.72 | 7.50 | 101% | | | 7.62 | 1.02 | | |
| | 8.00 | 0.00 | | | 8.33 | 83.72 | 8.00 | 102% | | | 8.33 | 1.02 | | |
| | 9.00 | 0.00 | | | 9.05 | 83.72 | 9.00 | 102% | | | 9.05 | 1.02 | | |
| | 10.00 | 0.00 | | | 9.77 | 83.72 | | | | | 9.77 | 1.02 | | |
| Full SC+preSC-B | 0.00 | 0.00 | 82.00 | 2.18 | 0.00 | 0.00 | 0.00 | 0% | 4.52 | 6.70 | 0.00 | 0.00 | 2.31 | 0.10 |
| | 2.00 | 0.00 | | | 1.40 | 0.00 | 2.00 | 7% | | | 1.40 | 0.00 | 3.23 | 0.20 |
| | 3.00 | 12.00 | | | 3.57 | 19.46 | 3.00 | 17% | | | 3.57 | 0.24 | 3.80 | 0.30 |
| | 3.50 | 18.00 | | | 5.75 | 68.96 | 3.50 | 23% | | | 5.75 | 0.84 | 4.16 | 0.40 |
| | 4.00 | 28.00 | | | 7.93 | 79.11 | 4.00 | 36% | | | 7.93 | 0.96 | 4.52 | 0.50 |
| | 4.50 | 35.00 | | | | | 4.50 | 49% | | | | | 4.88 | 0.60 |
| | 5.00 | 40.00 | | | | | 5.00 | 63% | | | | | 5.24 | 0.70 |
| | 5.50 | 58.00 | | | | | 5.50 | 77% | | | | | 5.60 | 0.80 |
| | 6.00 | 41.00 | | | | | 6.00 | 86% | | | | | | |
| | 6.50 | 40.00 | | | | | 6.50 | 88% | | | | | | |
| | 7.00 | 37.00 | | | | | 7.00 | 91% | | | | | | |
| | 7.50 | 11.00 | | | | | 7.50 | 94% | | | | | | |
| | 8.00 | 10.00 | | | | | 8.00 | | | | | | | |
| | 9.00 | 8.00 | | | | | 9.00 | | | | | | | |
| | 10.00 | 0.00 | | | | | | | | | | | | |

| | | | | | | | | | | | | | | |
|---|---|---|---|---|---|---|---|---|---|---|---|---|---|---|
| **Full SC+preSC-B+pre-SC-A** | 0.00 | 0.00 | 82.00 | 2.80 | 0.00 | 0.00 | 0.00 | 0% | 4.00 | 6.81 | 0.00 | 0.00 | 1.25 | 0.10 |
| | 2.00 | 0.00 | | | 0.14 | 0.00 | 2.00 | 17% | | | 0.14 | 0.00 | 2.35 | 0.20 |
| | 3.00 | 22.00 | | | 2.95 | 20.79 | 3.00 | 27% | | | 2.95 | 0.25 | 3.14 | 0.30 |
| | 3.50 | 38.00 | | | 5.75 | 74.29 | 3.50 | 38% | | | 5.75 | 0.91 | 3.57 | 0.40 |
| | 4.00 | 44.00 | | | 8.55 | 84.07 | 4.00 | 50% | | | 8.55 | 1.03 | 4.00 | 0.50 |
| | 4.50 | 55.00 | | | | | 4.50 | 62% | | | | | 4.43 | 0.60 |
| | 5.00 | 50.00 | | | | | 5.00 | 73% | | | | | 4.86 | 0.70 |
| | 5.50 | 58.00 | | | | | 5.50 | 85% | | | | | 5.29 | 0.80 |
| | 6.00 | 49.00 | | | | | 6.00 | 92% | | | | | | |
| | 6.50 | 41.00 | | | | | 6.50 | 94% | | | | | | |
| | 7.00 | 44.00 | | | | | 7.00 | 96% | | | | | | |
| | 7.50 | 14.00 | | | | | 7.50 | 98% | | | | | | |
| | 8.00 | 12.00 | | | | | 8.00 | 100% | | | | | | |
| | 9.00 | 8.00 | | | | | 9.00 | | | | | | | |
| | 10.00 | 0.00 | | | | | | | | | | | | |
| **Meiosis I+/-II** | 0.00 | 0.00 | 82.00 | | 0.00 | 0.00 | 0.00 | 0% | 7.87 | | 0.00 | 0.00 | 6.02 | 0.10 |
| | 2.00 | 0.00 | | | 2.00 | 0.00 | 2.00 | 0% | | | 2.00 | 0.00 | 7.06 | 0.20 |
| | 3.00 | 0.00 | | | 3.00 | 0.00 | 3.00 | 0% | | | 3.00 | 0.00 | 7.40 | 0.30 |
| | 3.50 | 0.00 | | | 3.50 | 0.00 | 3.50 | 0% | | | 3.50 | 0.00 | 7.65 | 0.40 |
| | 4.00 | 0.00 | | | 4.00 | 0.00 | 4.00 | 0% | | | 4.00 | 0.00 | 7.87 | 0.50 |
| | 4.50 | 0.00 | | | 4.50 | 0.00 | 4.50 | 0% | | | 4.50 | 0.00 | 8.20 | 0.60 |
| | 5.00 | 2.00 | | | 5.00 | 2.00 | 5.00 | 2% | | | 5.00 | 0.02 | 8.71 | 0.70 |
| | 5.50 | 2.00 | | | 5.50 | 2.00 | 5.50 | 2% | | | 5.50 | 0.02 | 9.45 | 0.80 |
| | 6.00 | 8.00 | | | 6.00 | 8.00 | 6.00 | 10% | | | 6.00 | 0.10 | | |
| | 6.50 | 13.00 | | | 6.50 | 13.00 | 6.50 | 16% | | | 6.50 | 0.16 | | |
| | 7.00 | 15.00 | | | 7.00 | 15.00 | 7.00 | 18% | | | 7.00 | 0.18 | | |
| | 7.50 | 27.00 | | | 7.50 | 27.00 | 7.50 | 33% | | | 7.50 | 0.33 | | |
| | 8.00 | 46.00 | | | 8.00 | 46.00 | 8.00 | 56% | | | 8.00 | 0.56 | | |
| | 9.00 | 62.00 | | | 9.00 | 62.00 | 9.00 | 76% | | | 9.00 | 0.76 | | |
| | 10.00 | 70.00 | | | 10.00 | 70.00 | 10.00 | 85% | | | 10.00 | 0.85 | | |
| | 11.00 | 82.00 | | | 11.00 | 82.00 | 11.00 | 100% | | | 11.00 | 1.00 | | |
| **End** | | | | | | | | | | | | | | |

| Input |  |  |  | LifeSpan | Cumulative Values: |  |  |  | Lifespan @ the 50 percentile |  | Percentage of maximum |  | In 10% increments |  |
|---|---|---|---|---|---|---|---|---|---|---|---|---|---|---|
|  | Measurements |  | Total # Events |  | at interpolated timepoints |  | at taken timepoints |  | 0.50 | 0.50 |  |  |  |  |
| Event Class | Time (Hrs) | Units | Units | Time (Hrs) | Time (Hrs) | Units | Time (Hrs) | Units | Entry | Exit | Time (Hrs) | Units | Time (Hrs) | Units |
| O-Hunter_Kleckner-NKY3230-TC2 | 0.00 | 0.00 | 23.60 |  | 0.00 | 0.00 | 0.00 | 0% | 5.52 |  | -5.52 | 0.00 | -1.46 | 0.10 |
|  | 2.50 | 0.00 |  |  | 2.50 | 0.00 | 2.50 | 0% | f CO set to 'zero' |  | -3.02 | 0.00 | -1.20 | 0.20 |
|  | 3.00 | 0.00 |  |  | 3.00 | 0.00 | 3.00 | 0% | 0.00 |  | -2.52 | 0.00 | -0.90 | 0.30 |
|  | 3.50 | 0.00 |  |  | 3.50 | 0.00 | 3.50 | 0% |  |  | -2.02 | 0.00 | -0.51 | 0.40 |
|  | 4.00 | 1.80 |  |  | 4.00 | 1.80 | 4.00 | 8% |  |  | -1.52 | 0.08 | 0.00 | 0.50 |
|  | 4.50 | 6.30 |  |  | 4.50 | 6.30 | 4.50 | 27% |  |  | -1.02 | 0.27 | 0.51 | 0.60 |
|  | 5.00 | 9.40 |  |  | 5.00 | 9.40 | 5.00 | 40% |  |  | -0.52 | 0.40 | 0.98 | 0.70 |
|  | 6.00 | 14.00 |  |  | 6.00 | 14.00 | 6.00 | 59% |  |  | 0.48 | 0.59 | 1.45 | 0.80 |
|  | 7.50 | 21.50 |  |  | 7.50 | 21.50 | 7.50 | 91% |  |  | 1.98 | 0.91 |  |  |
|  | 10.00 | 23.60 |  |  | 10.00 | 23.60 | 10.00 | 100% |  |  | 4.48 | 1.00 |  |  |
| DSB-TC20 | 0.00 | 0.00 | 25.00 | 1.23 | 0.00 | 0.00 | 0.00 | 0% | 3.80 | 5.03 | -5.52 | 0.00 | -3.96 | 0.10 |
|  | 2.50 | 3.30 |  |  | 0.30 | 0.40 | 2.50 | 22% | f CO set to 'zero' |  | -5.22 | 0.02 | -3.18 | 0.20 |
|  | 3.00 | 4.50 |  |  | 1.54 | 2.43 | 3.00 | 31% | -1.72 | -0.49 | -3.99 | 0.10 | -2.56 | 0.30 |
|  | 3.50 | 6.00 |  |  | 2.77 | 6.37 | 3.50 | 43% |  |  | -2.75 | 0.25 | -2.14 | 0.40 |
|  | 4.00 | 7.30 |  |  | 4.00 | 13.67 | 4.00 | 55% |  |  | -1.52 | 0.55 | -1.72 | 0.50 |
|  | 4.50 | 7.20 |  |  | 5.23 | 19.02 | 4.50 | 63% |  |  | -0.29 | 0.76 | -1.22 | 0.60 |
|  | 5.00 | 5.90 |  |  | 6.46 | 21.87 | 5.00 | 72% |  |  | 0.94 | 0.87 | -0.64 | 0.70 |
|  | 6.00 | 3.50 |  |  | 7.70 | 23.23 | 6.00 | 83% |  |  | 2.17 | 0.93 | 0.14 | 0.80 |
|  | 7.50 | 1.40 |  |  | 8.93 | 24.41 | 7.50 | 92% |  |  | 3.41 | 0.98 |  |  |
|  | 10.00 | 1.00 |  |  | 10.16 | 25.25 | 10.00 | 101% |  |  | 4.64 | 1.01 |  |  |
|  | 11.00 | 0.00 |  |  |  |  |  |  |  |  |  |  |  |  |
| SEI (TC20) | 0.00 | 0.00 | 11.80 | 1.03 | 0.00 | 0.00 | 0.00 | 0% | 4.43 | 5.46 | -5.52 | 0.00 | -2.49 | 0.10 |
|  | 2.50 | 0.15 |  |  | 0.40 | 0.02 | 2.50 | 3% | f CO set to 'zero' |  | -5.13 | 0.00 | -1.94 | 0.20 |
|  | 3.00 | 0.80 |  |  | 1.42 | 0.11 | 3.00 | 10% | -1.09 | -0.06 | -4.10 | 0.01 | -1.65 | 0.30 |
|  | 3.50 | 1.68 |  |  | 2.45 | 0.26 | 3.50 | 17% |  |  | -3.07 | 0.02 | -1.37 | 0.40 |
|  | 4.00 | 4.20 |  |  | 3.47 | 1.89 | 4.00 | 35% |  |  | -2.05 | 0.16 | -1.09 | 0.50 |
|  | 4.50 | 4.30 |  |  | 4.50 | 6.19 | 4.50 | 52% |  |  | -1.02 | 0.52 | -0.60 | 0.60 |
|  | 5.00 | 2.10 |  |  | 5.53 | 8.37 | 5.00 | 61% |  |  | 0.00 | 0.71 | -0.05 | 0.70 |
|  | 6.00 | 2.25 |  |  | 6.55 | 10.09 | 6.00 | 78% |  |  | 1.03 | 0.85 | 0.64 | 0.80 |
|  | 7.50 | 0.81 |  |  | 7.58 | 10.88 | 7.50 | 92% |  |  | 2.06 | 0.92 |  |  |
|  | 10.00 | 0.22 |  |  | 8.60 | 11.43 | 10.00 | 100% |  |  | 3.08 | 0.97 |  |  |
|  | 11.00 | 0.00 |  |  | 9.63 | 11.74 |  |  |  |  | 4.11 | 0.99 |  |  |
|  |  |  |  |  | 10.66 | 11.81 |  |  |  |  | 5.13 | 1.00 |  |  |
| IH-dHJ (tc20) | 0.00 | 0.00 | 23.60 | 0.40 | 0.00 | 0.00 | 0.00 | 0% | 5.34 | 5.75 | -5.52 | 0.00 | -1.81 | 0.10 |
|  | 2.50 | 0.00 |  |  | 0.08 | 0.00 | 2.50 | 0% | f CO set to 'zero' |  | -5.44 | 0.00 | -1.41 | 0.20 |
|  | 3.00 | 0.25 |  |  | 0.48 | 0.00 | 3.00 | 2% | -0.18 | 0.22 | -5.04 | 0.00 | -1.09 | 0.30 |
|  | 3.50 | 0.89 |  |  | 0.88 | 0.00 | 3.50 | 7% |  |  | -4.64 | 0.00 | -0.67 | 0.40 |
|  | 4.00 | 2.24 |  |  | 1.29 | 0.00 | 4.00 | 17% |  |  | -4.24 | 0.00 | -0.18 | 0.50 |
|  | 4.50 | 2.95 |  |  | 1.69 | 0.00 | 4.50 | 32% |  |  | -3.83 | 0.00 | 0.35 | 0.60 |
|  | 5.00 | 1.95 |  |  | 2.09 | 0.00 | 5.00 | 43% |  |  | -3.43 | 0.00 | 0.96 | 0.70 |
|  | 6.00 | 1.77 |  |  | 2.49 | 0.00 | 6.00 | 62% |  |  | -3.03 | 0.00 | 1.79 | 0.80 |
|  | 7.50 | 0.91 |  |  | 2.89 | 0.20 | 7.50 | 82% |  |  | -2.63 | 0.01 |  |  |
|  | 10.00 | 0.47 |  |  | 3.29 | 0.82 | 10.00 | 99% |  |  | -2.23 | 0.03 |  |  |
|  | 11.00 | 0.00 |  |  | 3.70 | 2.24 |  |  |  |  | -1.83 | 0.10 |  |  |
|  |  |  |  |  | 4.10 | 4.62 |  |  |  |  | -1.42 | 0.20 |  |  |
|  |  |  |  |  | 4.50 | 7.57 |  |  |  |  | -1.02 | 0.32 |  |  |
|  |  |  |  |  | 4.90 | 9.72 |  |  |  |  | -0.62 | 0.41 |  |  |
|  |  |  |  |  | 5.30 | 11.61 |  |  |  |  | -0.22 | 0.49 |  |  |
|  |  |  |  |  | 5.71 | 13.44 |  |  |  |  | 0.18 | 0.57 |  |  |
|  |  |  |  |  | 6.11 | 15.15 |  |  |  |  | 0.59 | 0.64 |  |  |
|  |  |  |  |  | 6.51 | 16.62 |  |  |  |  | 0.99 | 0.70 |  |  |
|  |  |  |  |  | 6.91 | 17.87 |  |  |  |  | 1.39 | 0.76 |  |  |
|  |  |  |  |  | 7.31 | 18.89 |  |  |  |  | 1.79 | 0.80 |  |  |
|  |  |  |  |  | 7.71 | 19.76 |  |  |  |  | 2.19 | 0.84 |  |  |
|  |  |  |  |  | 8.12 | 20.56 |  |  |  |  | 2.59 | 0.87 |  |  |
|  |  |  |  |  | 8.52 | 21.29 |  |  |  |  | 3.00 | 0.90 |  |  |
|  |  |  |  |  | 8.92 | 21.95 |  |  |  |  | 3.40 | 0.93 |  |  |
|  |  |  |  |  | 9.32 | 22.54 |  |  |  |  | 3.80 | 0.96 |  |  |
|  |  |  |  |  | 9.72 | 23.06 |  |  |  |  | 4.20 | 0.98 |  |  |
|  |  |  |  |  | 10.13 | 23.47 |  |  |  |  | 4.60 | 0.99 |  |  |
|  |  |  |  |  | 10.53 | 23.70 |  |  |  |  | 5.01 | 1.00 |  |  |
|  |  |  |  |  | 10.93 | 23.73 |  |  |  |  | 5.41 | 1.01 |  |  |
| Meiosis divisions (TC20) | 0.00 | 0.00 | 94.00 |  | 0.00 | 0.00 | 0.00 | 0% | 6.80 |  | -5.52 | 0.00 | -0.44 | 0.10 |
|  | 2.50 | 0.00 |  |  | 2.50 | 0.00 | 2.50 | 0% | f CO set to 'zero' |  | -3.02 | 0.00 | 0.11 | 0.20 |
|  | 3.00 | 0.00 |  |  | 3.00 | 0.00 | 3.00 | 0% | 1.28 |  | -2.52 | 0.00 | 0.60 | 0.30 |
|  | 3.50 | 0.00 |  |  | 3.50 | 0.00 | 3.50 | 0% |  |  | -2.02 | 0.00 | 0.94 | 0.40 |
|  | 4.00 | 0.00 |  |  | 4.00 | 0.00 | 4.00 | 0% |  |  | -1.52 | 0.00 | 1.28 | 0.50 |
|  | 4.50 | 2.00 |  |  | 4.50 | 2.00 | 4.50 | 2% |  |  | -1.02 | 0.02 | 1.63 | 0.60 |
|  | 5.00 | 8.00 |  |  | 5.00 | 8.00 | 5.00 | 9% |  |  | -0.52 | 0.09 | 1.97 | 0.70 |
|  | 6.00 | 25.00 |  |  | 6.00 | 25.00 | 6.00 | 27% |  |  | 0.48 | 0.27 | 3.02 | 0.80 |
|  | 7.50 | 66.00 |  |  | 7.50 | 66.00 | 7.50 | 70% |  |  | 1.98 | 0.70 |  |  |
|  | 10.00 | 88.00 |  |  | 10.00 | 88.00 | 10.00 | 94% |  |  | 4.48 | 0.94 |  |  |
|  | 12.00 | 94.00 |  |  | 12.00 | 94.00 | 12.00 | 100% |  |  | 6.48 | 1.00 |  |  |
| End |  |  |  |  |  |  |  |  |  |  |  |  |  |  |

| Input | | | | LifeSpan | Cumulative Values: | | | | Lifespan @ the 50 percentile | | Percentage of maximum | | In 10% increments | |
|---|---|---|---|---|---|---|---|---|---|---|---|---|---|---|
| Event Class | Measurements | | Total # Events | | at interpolated timepoints | | at taken timepoints | | 0.50 | 0.50 | | | | |
| | Time (Hrs) | Units | Units | Time (Hrs) | Time (Hrs) | Units | Time (Hrs) | Units | Entry | Exit | Time (Hrs) | Units | Time (Hrs) | Units |
| tc42-#3-201-CO | 0 | 0.00 | 0.18 | | 0.00 | 0.00 | 0.00 | 3% | 8.02 | | 0.00 | 0.03 | 4.54 | 0.10 |
| | 2.5 | 0.00 | | | 2.50 | 0.00 | 2.50 | 1% | | | 2.50 | 0.01 | 6.68 | 0.20 |
| | 4 | 0.01 | | | 4.00 | 0.01 | 4.00 | 7% | | | 4.00 | 0.07 | 7.31 | 0.30 |
| | 5 | 0.02 | | | 5.00 | 0.02 | 5.00 | 13% | | | 5.00 | 0.13 | 7.67 | 0.40 |
| | 6 | 0.03 | | | 6.00 | 0.03 | 6.00 | 17% | | | 6.00 | 0.17 | 8.02 | 0.50 |
| | 7 | 0.04 | | | 7.00 | 0.04 | 7.00 | 21% | | | 7.00 | 0.21 | 8.38 | 0.60 |
| | 8.5 | 0.12 | | | 8.50 | 0.12 | 8.50 | 64% | | | 8.50 | 0.64 | 8.98 | 0.70 |
| | 10 | 0.15 | | | 10.00 | 0.15 | 10.00 | 84% | | | 10.00 | 0.84 | 9.71 | 0.80 |
| | 11 | 0.15 | | | 11.00 | 0.15 | 11.00 | 84% | | | 11.00 | 0.84 | | |
| | 14 | 0.18 | | | 14.00 | 0.18 | 14.00 | 100% | | | 14.00 | 1.00 | | |
| tc42-#3-201-DSB | 0 | 0.00 | 0.225 | 1.15 | 0.00 | 0.00 | 0.00 | 0% | 4.53 | 5.68 | 0.00 | 0.00 | 2.88 | 0.10 |
| | 2.5 | 0.00 | | | 0.55 | 0.00 | 2.50 | 7% | | | 0.55 | 0.00 | 3.27 | 0.20 |
| | 4 | 0.07 | | | 1.70 | 0.00 | 4.00 | 38% | | | 1.70 | 0.01 | 3.67 | 0.30 |
| | 5 | 0.06 | | | 2.85 | 0.02 | 5.00 | 60% | | | 2.85 | 0.09 | 4.08 | 0.40 |
| | 6 | 0.05 | | | 4.00 | 0.09 | 6.00 | 78% | | | 4.00 | 0.38 | 4.53 | 0.50 |
| | 7 | 0.04 | | | 5.15 | 0.14 | 7.00 | 91% | | | 5.15 | 0.63 | 4.99 | 0.60 |
| | 8.5 | 0.01 | | | 6.30 | 0.19 | 8.50 | 101% | | | 6.30 | 0.83 | 5.53 | 0.70 |
| | 10 | 0.00 | | | 7.45 | 0.22 | 10.00 | 102% | | | 7.45 | 0.96 | 6.11 | 0.80 |
| | 11 | 0.00 | | | 8.60 | 0.23 | | | | | 8.60 | 1.01 | | |
| | | | | | 9.76 | 0.23 | | | | | 9.76 | 1.02 | | |
| | | | | | 10.91 | 0.23 | | | | | 10.91 | 1.02 | | |
| tc42-#3-201-SEI | 0 | 0.00 | 0.11 | 0.64 | 0.00 | 0.00 | 0.00 | 0% | 6.07 | 671% | 0.00 | 0.00 | 3.85 | 0.10 |
| | 2.5 | 0.00 | | | 0.21 | 0.00 | 2.50 | 2% | | | 0.21 | 0.00 | 4.68 | 0.20 |
| | 4 | 0.005428962 | | | 0.85 | 0.00 | 4.00 | 11% | | | 0.85 | 0.00 | 5.22 | 0.30 |
| | 5 | 0.011 | | | 1.50 | 0.00 | 5.00 | 26% | | | 1.50 | 0.01 | 5.66 | 0.40 |
| | 6 | 0.018 | | | 2.14 | 0.00 | 6.00 | 48% | | | 2.14 | 0.02 | 6.07 | 0.50 |
| | 7 | 0.015 | | | 2.78 | 0.00 | 7.00 | 70% | | | 2.78 | 0.03 | 6.51 | 0.60 |
| | 8.5 | 0.008 | | | 3.43 | 0.01 | 8.50 | 92% | | | 3.43 | 0.07 | 6.99 | 0.70 |
| | 10 | 0.002 | | | 4.07 | 0.01 | 10.00 | 100% | | | 4.07 | 0.12 | 7.57 | 0.80 |
| | 11 | 0.000 | | | 4.71 | 0.02 | | | | | 4.71 | 0.20 | | |
| | | | | | 5.36 | 0.04 | | | | | 5.36 | 0.33 | | |
| | | | | | 6.00 | 0.05 | | | | | 6.00 | 0.48 | | |
| | | | | | 6.64 | 0.07 | | | | | 6.64 | 0.63 | | |
| | | | | | 7.29 | 0.08 | | | | | 7.29 | 0.76 | | |
| | | | | | 7.93 | 0.09 | | | | | 7.93 | 0.86 | | |
| | | | | | 8.57 | 0.10 | | | | | 8.57 | 0.92 | | |
| | | | | | 9.22 | 0.11 | | | | | 9.22 | 0.97 | | |
| | | | | | 9.86 | 0.11 | | | | | 9.86 | 0.99 | | |
| | | | | | 10.50 | 0.11 | | | | | 10.50 | 1.00 | | |
| tc42-#3-201-IH-dHJ-COinWT | 0 | 0.000 | 0.22 | 0.83 | 0.00 | 0.00 | 0.00 | 0% | 6.44 | 7.27 | 0.00 | 0.00 | 4.30 | 0.10 |
| | 2.5 | 0.001 | | | 0.37 | 0.00 | 2.50 | 1% | | | 0.37 | 0.00 | 5.10 | 0.20 |
| | 4 | 0.008 | | | 1.20 | 0.00 | 4.00 | 7% | | | 1.20 | 0.00 | 5.65 | 0.30 |
| | 5 | 0.022 | | | 2.03 | 0.00 | 5.00 | 19% | | | 2.03 | 0.01 | 6.13 | 0.40 |
| | 6 | 0.034 | | | 2.86 | 0.00 | 6.00 | 37% | | | 2.86 | 0.02 | 6.44 | 0.50 |
| | 7 | 0.062 | | | 3.69 | 0.01 | 7.00 | 69% | | | 3.69 | 0.05 | 6.73 | 0.60 |
| | 8.5 | 0.020 | | | 4.52 | 0.03 | 8.50 | 94% | | | 4.52 | 0.12 | 7.05 | 0.70 |
| | 10 | 0.006 | | | 5.34 | 0.05 | 10.00 | 101% | | | 5.34 | 0.23 | 7.52 | 0.80 |
| | 11 | 0.000 | | | 6.17 | 0.09 | | | | | 6.17 | 0.41 | | |
| | | | | | 7.00 | 0.15 | | | | | 7.00 | 0.69 | | |
| | | | | | 7.83 | 0.19 | | | | | 7.83 | 0.87 | | |
| | | | | | 8.66 | 0.21 | | | | | 8.66 | 0.95 | | |
| | | | | | 9.48 | 0.22 | | | | | 9.48 | 1.00 | | |
| | | | | | 10.31 | 0.22 | | | | | 10.31 | 1.02 | | |
| tc42-#3-201-IH-dHJ-COin201 | 0 | 0.000 | 0.18 | 1.01 | 0.00 | 0.00 | 0.00 | 0% | 6.36 | 7.38 | 0.00 | 0.00 | 4.24 | 0.10 |
| | 2.5 | 0.001 | | | 0.93 | 0.00 | 2.50 | 2% | | | 0.93 | 0.00 | 5.05 | 0.20 |
| | 4 | 0.008 | | | 1.94 | 0.00 | 4.00 | 7% | | | 1.94 | 0.01 | 5.59 | 0.30 |
| | 5 | 0.021707361 | | | 2.95 | 0.00 | 5.00 | 19% | | | 2.95 | 0.02 | 6.07 | 0.40 |
| | 6 | 0.034 | | | 3.96 | 0.01 | 6.00 | 38% | | | 3.96 | 0.07 | 6.36 | 0.50 |
| | 7 | 0.062 | | | 4.98 | 0.03 | 7.00 | 72% | | | 4.98 | 0.19 | 6.66 | 0.60 |
| | 8.5 | 0.020 | | | 5.99 | 0.07 | 8.50 | 94% | | | 5.99 | 0.37 | 6.95 | 0.70 |
| | 10 | 0.006 | | | 7.00 | 0.13 | 10.00 | 102% | | | 7.00 | 0.72 | 7.46 | 0.80 |
| | 11 | 0.000 | | | 8.01 | 0.16 | | | | | 8.01 | 0.90 | | |
| | | | | | 9.02 | 0.18 | | | | | 9.02 | 0.99 | | |
| | | | | | 10.04 | 0.18 | | | | | 10.04 | 1.02 | | |
| tc42-#3-201-SEI-COin201 | 0 | 0.000 | 0.09 | 0.79 | 0.00 | 0.00 | 0.00 | 0% | 5.98 | 6.76 | 0.00 | 0.00 | 3.77 | 0.10 |
| | 2.5 | 0.001 | | | 0.50 | 0.00 | 2.50 | 3% | | | 0.50 | 0.00 | 4.58 | 0.20 |
| | 4 | 0.005 | | | 1.28 | 0.00 | 4.00 | 13% | | | 1.28 | 0.01 | 5.15 | 0.30 |
| | 5 | 0.011 | | | 2.07 | 0.00 | 5.00 | 27% | | | 2.07 | 0.02 | 5.57 | 0.40 |
| | 6 | 0.018 | | | 2.85 | 0.00 | 6.00 | 51% | | | 2.85 | 0.04 | 5.98 | 0.50 |
| | 7 | 0.015 | | | 3.64 | 0.01 | 7.00 | 72% | | | 3.64 | 0.09 | 6.42 | 0.60 |
| | 8.5 | 0.008 | | | 4.43 | 0.02 | 8.50 | 93% | | | 4.43 | 0.17 | 6.89 | 0.70 |
| | 10 | 0.002 | | | 5.21 | 0.03 | 10.00 | 100% | | | 5.21 | 0.31 | 7.46 | 0.80 |
| | 11 | 0.000 | | | 6.00 | 0.05 | | | | | 6.00 | 0.51 | | |
| | | | | | 6.79 | 0.06 | | | | | 6.79 | 0.68 | | |
| | | | | | 7.57 | 0.07 | | | | | 7.57 | 0.82 | | |
| | | | | | 8.36 | 0.08 | | | | | 8.36 | 0.92 | | |
| | | | | | 9.15 | 0.09 | | | | | 9.15 | 0.98 | | |
| | | | | | 9.93 | 0.09 | | | | | 9.93 | 1.00 | | |
| | | | | | 10.72 | 0.09 | | | | | 10.72 | 1.01 | | |
| tc42-#3-201-lines (5) | 0 | 0 | 0.95 | 0.63 | 0.00 | 0.00 | 0.00 | 0% | 6.33 | 6.96 | 0.00 | 0.00 | 2.87 | 0.10 |
| | 5 | 0.059 | | | 0.02 | 0.00 | 5.00 | 28% | | | 0.02 | 0.00 | 4.19 | 0.20 |
| | 6 | 0.078 | | | 0.66 | 0.01 | 6.00 | 43% | | | 0.66 | 0.01 | 5.19 | 0.30 |
| | 7 | 0.235 | | | 1.29 | 0.02 | 7.00 | 76% | | | 1.29 | 0.02 | 5.88 | 0.40 |
| | 8.5 | 0.035 | | | 1.92 | 0.05 | 8.50 | 99% | | | 1.92 | 0.05 | 6.33 | 0.50 |
| | 10 | 0.000 | | | 2.56 | 0.08 | | | | | 2.56 | 0.08 | 6.60 | 0.60 |
| | | | | | 3.19 | 0.11 | | | | | 3.19 | 0.12 | 6.86 | 0.70 |
| | | | | | 3.83 | 0.16 | | | | | 3.83 | 0.17 | 7.17 | 0.80 |
| | | | | | 4.46 | 0.21 | | | | | 4.46 | 0.22 | | |
| | | | | | 5.10 | 0.27 | | | | | 5.10 | 0.29 | | |
| | | | | | 5.73 | 0.35 | | | | | 5.73 | 0.37 | | |
| | | | | | 6.37 | 0.48 | | | | | 6.37 | 0.51 | | |
| | | | | | 7.00 | 0.72 | | | | | 7.00 | 0.76 | | |
| | | | | | 7.63 | 0.87 | | | | | 7.63 | 0.91 | | |
| | | | | | 8.27 | 0.93 | | | | | 8.27 | 0.98 | | |
| | | | | | 8.90 | 0.96 | | | | | 8.90 | 1.01 | | |
| | | | | | 9.54 | 0.97 | | | | | 9.54 | 1.02 | | |
| tc42-#3-201-4 | 0 | 0.000 | 0.95 | 1.63 | 0.00 | 0.00 | 0.00 | 0% | 4.63 | 6.25 | 0.00 | 0.00 | 1.54 | 0.10 |
| | 5 | 0.260 | | | 1.12 | 0.06 | 5.00 | 57% | | | 1.12 | 0.06 | 2.62 | 0.20 |
| | 6 | 0.300 | | | 2.75 | 0.20 | 6.00 | 77% | | | 2.75 | 0.21 | 3.35 | 0.30 |
| | 7 | 0.300 | | | 4.37 | 0.43 | 7.00 | 90% | | | 4.37 | 0.45 | 4.03 | 0.40 |
| | 8.5 | 0.060 | | | 6.00 | 0.73 | 8.50 | 99% | | | 6.00 | 0.77 | 4.63 | 0.50 |
| | 10 | 0.000 | | | 7.63 | 0.93 | | | | | 7.63 | 0.98 | 5.14 | 0.60 |
| | | | | | 9.25 | 0.96 | | | | | 9.25 | 1.01 | 5.66 | 0.70 |
| | | | | | | | | | | | | | 6.26 | 0.80 |
| tc42-#3-201-3 | 0 | 0 | 0.95 | 1.95 | 0.00 | 0.00 | 0.00 | 0% | 4.37 | 6.32 | 0.00 | 0.00 | 1.35 | 0.10 |

| | | | | | | | | | | | | | | |
|---|---|---|---|---|---|---|---|---|---|---|---|---|---|---|
| | 5 | 0.330 | | | 0.15 | 0.01 | 5.00 | 62% | | | 0.15 | 0.01 | 2.40 | 0.20 |
| | 6 | 0.360 | | | 2.10 | 0.15 | 6.00 | 82% | | | 2.10 | 0.16 | 3.10 | 0.30 |
| | 7 | 0.210 | | | 4.05 | 0.42 | 7.00 | 91% | | | 4.05 | 0.44 | 3.79 | 0.40 |
| | 8.5 | 0.160 | | | 6.00 | 0.78 | 8.50 | 101% | | | 6.00 | 0.82 | 4.37 | 0.50 |
| | 10 | 0.000 | | | 7.95 | 0.95 | | | | | 7.95 | 1.00 | 4.89 | 0.60 |
| | | | | | 9.90 | 0.96 | | | | | 9.90 | 1.02 | 5.40 | 0.70 |
| | | | | | | | | | | | | | 5.91 | 0.80 |
| tc42-#3-201-4-5 | 0 | 0.000 | 0.95 | 2.25 | 0.00 | 0.00 | 0.00 | 0% | 4.75 | 7.01 | 0.00 | 0.00 | 1.38 | 0.10 |
| | 5 | 0.317 | | | 0.24 | 0.01 | 5.00 | 56% | | | 0.24 | 0.02 | 2.62 | 0.20 |
| | 6 | 0.382 | | | 2.49 | 0.17 | 6.00 | 81% | | | 2.49 | 0.18 | 3.33 | 0.30 |
| | 7 | 0.534 | | | 4.75 | 0.47 | 7.00 | 106% | | | 4.75 | 0.50 | 4.04 | 0.40 |
| | 8.5 | 0.094 | | | 7.00 | 1.01 | 8.50 | 109% | | | 7.00 | 1.06 | 4.75 | 0.50 |
| | 10 | 0.000 | | | 9.25 | 1.05 | | | | | 9.25 | 1.11 | 5.15 | 0.60 |
| tc42-#3-201-3-5 | 0 | 0.000 | 0.95 | 4.20 | 0.00 | 0.00 | 0.00 | 0% | 3.17 | 7.37 | 0.00 | 0.00 | 0.74 | 0.10 |
| | 5 | 0.644 | | | 1.80 | 0.23 | 5.00 | 84% | | | 1.80 | 0.24 | 1.48 | 0.20 |
| | 6 | 0.745 | | | 6.00 | 0.98 | 6.00 | 103% | | | 6.00 | 1.03 | 2.21 | 0.30 |
| | 7 | 0.745098039 | | | | | 7.00 | | | | | | 2.95 | 0.40 |
| | 8.5 | 0.25 | | | | | 8.50 | | | | | | 3.69 | 0.50 |
| | 10 | 0.00 | | | | | | | | | | | 4.43 | 0.60 |
| tc42-#3-201-2-5 | 0 | 0.00 | 0.95 | 5.82 | 0.00 | 0.00 | 0.00 | 0% | 2.89 | 8.71 | 0.00 | 0.00 | 0.52 | 0.10 |
| | 5 | 0.91 | | | 0.18 | 0.03 | 5.00 | 86% | | | 0.18 | 0.03 | 1.04 | 0.20 |
| | 6 | 0.95 | | | 6.00 | 0.98 | 6.00 | 104% | | | 6.00 | 1.04 | 1.56 | 0.30 |
| | 7 | 0.86 | | | | | 7.00 | | | | | | 2.09 | 0.40 |
| | 8.5 | 0.51 | | | | | 8.50 | | | | | | 2.61 | 0.50 |
| | 10 | 0.00 | | | | | | | | | | | 3.13 | 0.60 |
| tc42-#3-201-2-5 | 0 | 0.00 | 0.95 | | 0.00 | 0.00 | 0.00 | 0% | 2.61 | | 0.00 | 0.00 | 0.52 | 0.10 |
| | 5 | 0.91 | | | 5.00 | 0.91 | 5.00 | 96% | | | 5.00 | 0.96 | 1.04 | 0.20 |
| | 6 | 0.95 | | | 6.00 | 0.95 | 6.00 | 100% | | | 6.00 | 1.00 | 1.56 | 0.30 |
| | | | | | | | | | | | | | 2.09 | 0.40 |
| | | | | | | | | | | | | | 2.61 | 0.50 |
| | | | | | | | | | | | | | 3.13 | 0.60 |
| | | | | | | | | | | | | | 3.65 | 0.70 |
| | | | | | | | | | | | | | 4.17 | 0.80 |
| | | | | | | | | | | | | | 0.00 | |
| End | | | | | | | | | | | | | | |
